# Multi-micron crisscross structures from combinatorially assembled DNA-origami slats

**DOI:** 10.1101/2022.01.06.475243

**Authors:** Christopher M. Wintersinger, Dionis Minev, Anastasia Ershova, Hiroshi M. Sasaki, Gokul Gowri, Jonathan F. Berengut, F. Eduardo Corea-Dilbert, Peng Yin, William M. Shih

**Author notes:** Denotes equal contribution (Christopher M. Wintersinger, Dionis Minev, Anastasia Ershova).

## Abstract

Living systems achieve robust self-assembly across length scales. Meanwhile, nanofabrication strategies such as DNA origami have enabled robust self-assembly of submicron-scale shapes.However, erroneous and missing linkages restrict the number of unique origami that can be practically combined into a single supershape. We introduce crisscross polymerization of DNA-origami slats for strictly seed-dependent growth of custom multi-micron shapes with user-defined nanoscale surface patterning. Using a library of ~2000 strands that can be combinatorially assembled to yield any of ~1e48 distinct DNA origami slats, we realize five-gigadalton structures composed of >1000 uniquely addressable slats, and periodic structures incorporating >10,000 slats. Thus crisscross growth provides a generalizable route for prototyping and scalable production of devices integrating thousands of unique components that each are sophisticated and molecularly precise.

**One-sentence summary:** Crisscross polymerization of DNA-origami slats can yield micron-scale structures with uniquely addressable nanoscale features.

## Introduction

In structural DNA nanotechnology^1^, the scaffolded DNA origami method affords robust self-assembly of arbitrary two- and three-dimensional nanoscale objects^2–7^. The oligonucleotide “staple” strands are designed to lack complementarity to each other, and folding is exactly controlled by a long single-stranded DNA (ssDNA) scaffold that is substoichiometric with respect to the staple strands, resulting in one-to-one conversion of scaffold particles into origami. This absolute scaffold dependence enables assembly over a broad range of temperatures and salt concentrations, while circumventing accumulation of incomplete assemblies or byproducts. Due to their robust folding performance, origami offer a user-friendly approach — suitable for specialists and non-specialists alike — for creating structures with addressable features. Applications have included plasmonic devices relying on placement of nanoparticles^8,9^, therapeutic devices with spatial control over cargos that sense the *in vivo* environment^10,11^, and research tools that place biomolecules in specified arrangements to deduce their biophysical properties^12,13^.

One limitation of DNA origami is that their maximal size is limited to about five megadaltons because the shape is bounded by the length of the scaffold DNA. While it is possible to use longer scaffold sequences, they are difficult to obtain in ssDNA form, and are delicate to handle because they are prone to shearing^14–16^. DNA bricks^17–20^ may instead be used to create structures significantly larger than a single origami, with as many as 30,000 unique monomers and a total mass of ~0.5 GDa per assembled particle. In contrast to origami “staple” strands, DNA tiles and bricks are complementary to each other thus eliminating the scaffold dependence of the assembly. However, spontaneous association of building blocks effectively limits the yield for such single-pot growth processes and increases the burden for post-assembly purification. Additionally, for the largest of such structures, current pricing on nanomole-scale oligonucleotide synthesis can be cost-prohibitive.

Hierarchical approaches can be used, however achieving assemblies containing more than a few distinct DNA nanostructures has been challenging^21–31^. In the most complex demonstration to date, structures consisting of 64 unique DNA-origami components were constructed using a method termed fractal assembly. To suppress off-target joining of DNA-origami monomers, three sequential steps were employed to build 4-component, then 16-component, and finally 64-component supershapes^32^. Fine tuning of monomer stoichiometries and reaction temperatures were required, nevertheless reported yields dropped from ~93% to ~48% to ~2% for the three steps, respectively. Despite only four sub-assemblies coming together per stage and precise care in preparation, unfinished, erroneous, and aggregated by-products led to rapidly diminishing yields as more unique monomers and assembly stages were added. Thus fractal assembly in this way appears effective for hierarchical constructions with dozens of components, but may face severe yield issues when larger numbers of parts are desired.

Previously we introduced crisscross polymerization of ssDNA slats for robust control over nucleation^33^; here we generalize this method to operate with DNA-origami slats — which are over two orders of magnitude larger than their ssDNA counterparts — for synchronous initiation of growth of target supershapes from relatively small seeds. Using six-helix bundle (6HB)^29,34,35^ and twelve-helix bundle (12HB) nanorods extending weak binding handles along their lengths, we created a diversity of finite and periodic assemblies. The largest finite supershape consisted of 1022 unique slats arranged into a sheet, a molecular mass of ~5.4 gigadaltons, and lateral dimensions of ~2 μm. The number of fully formed assemblies is controlled exactly by the amount of seed added, with the robustness of growth from hundreds of origami-based parts comparable to that of origami folding itself from similar numbers of much smaller components.

### Designing DNA-origami slats for crisscross assembly

In crisscross polymerization, an incoming slat must engage with a large number of other slats (up to either eight or sixteen in this study) for stable attachment to the edge of a growing structure; this requirement for a high level of coordination is the basis for the robustness of crisscross against unwanted spurious nucleation^33^ (Fig. S1). To meet this design criterion, each individual pairwise interaction must be quite weak at the desired temperature of growth, and ideally far below this temperature as well. For ssDNA slats, this was achieved with interactions that span just a half turn of DNA (i.e. 5 or 6 bp)^33^. We hypothesized that crisscross assembly could also be implemented for DNA-origami slats by engineering sufficiently weak binding handles. Then to bypass the nucleation barrier in a controlled fashion, pre-formed seeds could be employed that use much stronger binding interactions to capture an initial set of “nucleating” slats in an arrangement that resembles a critical nucleus (Fig. S1Bi–ii). Consequently, a cascade of energetically favorable downstream assembly steps could propagate growth (Fig. 1A, Fig. S1Biii–v).

**Figure 1.**
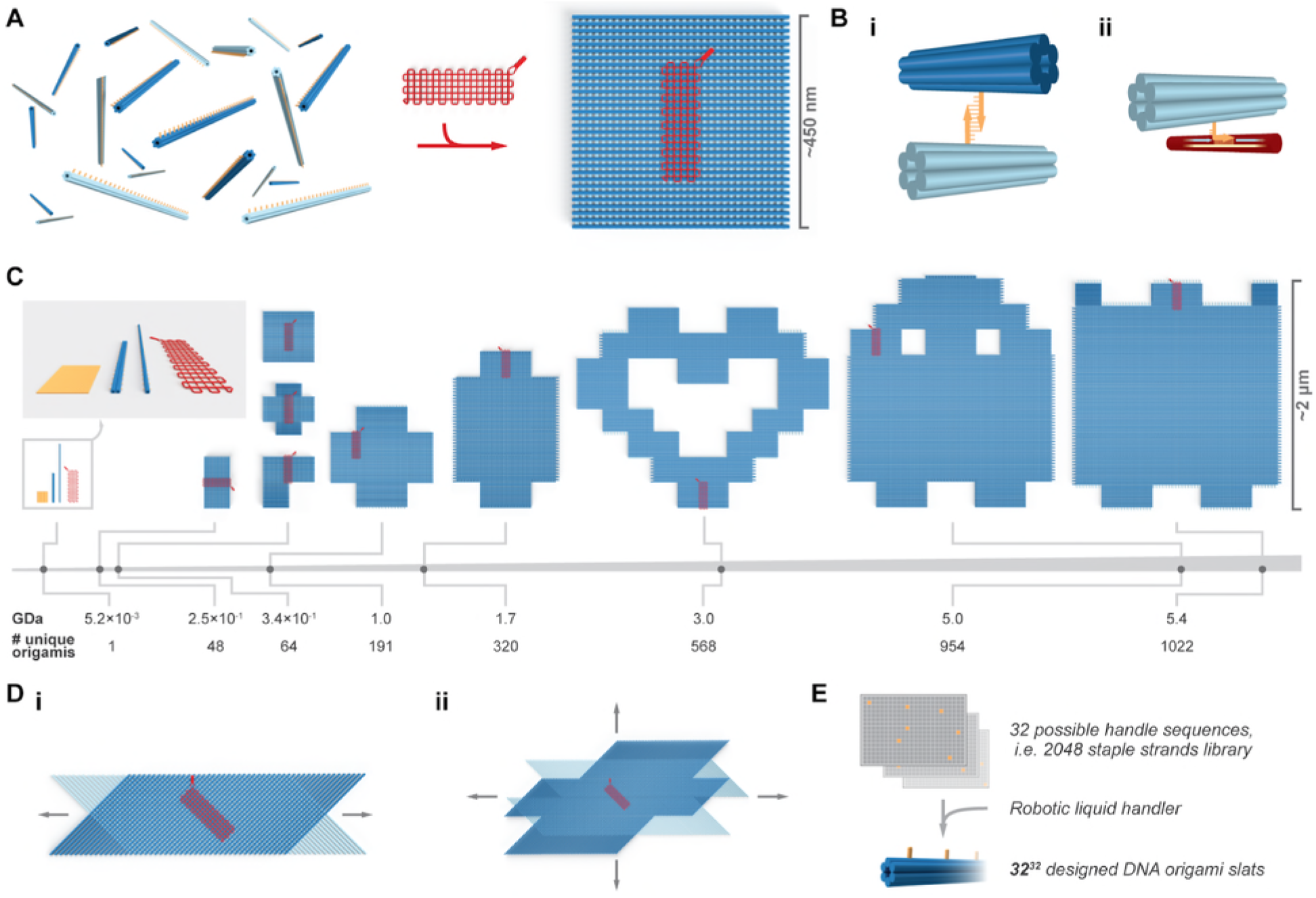
Overview of crisscross assembly of DNA-origami slats. **A**, left, a pool of 64 unique free 6HB slats. A 6HB slat functionally comprises a linear arrangement of 32 binding-site sequences, each selected from the same set of 32 distinct sequences; a square megastructure with 64 unique slats is triggered to form only when the gridiron-origami seed is added. **Bi,** binding of a pair of complementary weak 7-nt handles on two perpendicular 6HB slats. **Bii**, a strong 10-nt “plug” handle on a 6HB slat engaged with an exposed region of scaffold (i.e. a “socket”) on the gridiron seed. **C**, the breadth and relative scale of the megastructures tested, versus the leftward single DNA-origami slats, seed, and origami reference square. **Di–ii**, renderings of periodic one- and two-dimensionally growing ribbons and sheets. **E**, schematic for how subsets of the strand library are combinatorially collected to yield unique slats.

As a convenient shorthand, we refer to any designed DNA-based assembly that consists of roughly a million or more nucleotides as “megastructures”. To achieve crisscross growth of megastructures from DNA-origami building blocks, we designed 6HB and 12HB slats that assemble by nucleating upon a gridiron-origami^7^ seed (Fig. 1A, Fig. S2A). The 3’ ends of staple strands on the top and bottom helices of each slat were encoded with ssDNA binding handles to link the slats to one another, though each could alternatively be programmed as an addressable “node” that engages a desired cargo (Fig. 1Bi, Fig. S2Bi–ii). The 6HB is ~450 nm long and features 32 handle positions spaced ~14 nm apart along its length; the 12HB is ~225 nm long with 16 positions along its length (see Supplementary Text 1 and Fig. S3 for more discussion of the 12HB design). As depicted in red in Fig. 1A and in Fig. S2Aiii, the seed has 16 columns of five “sockets”, where each column captures an individual “nucleating” slat with five 10-nucleotide (nt) handles that each “plug” into its complementary socket (Fig. S2Biii, Fig. 1Bii). Spacings of the plug handles on a nucleating slat, at 42 bp intervals, are commensurate with spacing of seed sockets. We validated folding of 6HB slats, 12HB slats, and seeds by imaging with negative-stain transmission electron microscopy (TEM, Fig. S4–Fig. S5).

We tested unseeded formation of sample crisscross structures with 10-, 9-, 8-, 7-, or 6-nt handles to explore how handle length affects spurious nucleation as a function of temperature. The 10-, 9-, and 8-nt handles were found to yield significant unseeded assembly at relevant temperatures (Fig. S6A–Ci). However, this was not observed with either the 7- or 6-nt handles (Fig. S6Cii–iii). Hence, we narrowed our focus to 7-nt handles, affording greater thermal stability versus 6-nt handles, for creating origami-slat megastructures (Fig. 1C–D). The algorithm for designing the sequence handles, computed energies versus handle length, and need for poly-T linkers are described in Supplementary Text 2, Fig. S7, and Fig. S8, respectively.

### Diversity of finite megastructures from a small set of strands

We conceived of many finite and periodic megastructures that could be made from two perpendicular layers of the DNA slats (Fig. 1C–D). Initially, we considered creating these designs using unique handles at the slat intersections, but such an approach would have been cost prohibitive since a distinct set of staple strands would need to be purchased for each design. We hypothesized that with the same handle sequence appearing on distinct slats, or even on the same slat, any transient non-target interaction would readily dissociate except in cases where several matching handles could be engaged simultaneously. We selected 32 isoenergetic 7-nt handle sequences and purchased a library of 2048 staple strands (i.e. 32 positions × 32 handle sequences × 2 for complementary handles) that would allow any one of these handles to be encoded at any possible perpendicular slat intersection. In principle, subsets of staple strands could be selected from this library to create up to ^32^ (~10^48^) distinct 6HB (or 32^16^ ≈ 10^24^ half-length 12HB using an analogous library strategy) slats (Fig. 1E).

We tested this library for growth of finite megastructures to determine the extent to which the strands could be rearranged to make novel shapes. The relative area, molecular mass, and number of unique DNA origamis in each are shown in Fig. 1C. The smaller shapes with 48 and 64 slats have maximum lateral dimensions limited to 450 nm, the length of a 6HB slat (Fig. S9A). In order to achieve larger dimensions, we also designed assembly trajectories where the 6HB slats join in a zigzagging raster-fill pattern where for a typical step, sixteen parallel slats bind to each of two growth fronts that rotates 90 degrees clockwise or counterclockwise (Fig. S9B–D, Fig. S10). Using this raster-fill growth paradigm, we created larger megastructures including a 191-slat plus symbol, a 320-slat elongated plus symbol, a 568-slat heart, a ghost caricature with 954 slats, and a sheet with 1022 slats. The largest 1022-slat sheet has lateral dimensions ~2 μm and a molecular mass of ~5.4 gigadaltons, which is over an order of magnitude greater number of unique DNA origamis compared to fractal tiles as previously published^32^ (Fig. 1C). We juxtaposed a single DNA-origami square in Fig. 1C (i.e. the 85 nm × 85 nm shape as shown with leftmost orange render) to illustrate how origami crisscross overcomes the size limits attainable with single origamis.

We selected strands from the handle library for independent folding of each slat. We then combined the folded slats into pools with maximally 100 unique members, and concentrated each pool into a smaller volume (Fig. S11). Crisscross growth was initiated by mixing a seed with top- and bottom-layer slat pools. The larger finite shapes with rastering growth (Fig. S9C–D, Fig. S10) were assembled in several stages, where ~200 of the slats were added and incubated for ~60 hours before 2.5-fold dilution into a pool of the subsequent series of slats. We successfully assembled the panel of finite megastructures, as shown by TEM in Fig. 2. All the megastructures formed dispersed single particles in a seed-dependent fashion (Fig. S12–Fig. S15). The structures in Fig. 2Ai–v were formed exclusively with 6HB slats, with the exception of the 1022-slat sheet in Fig. 2Avi that also had 28 12HB slats. Half or more of the total slats in the Fig. 2B shapes were the shorter 12HB slats, allowing for the megastructures to have features finer than the length of a 6HB slat (Supplementary Text 1, Fig. S3).

**Figure 2.**
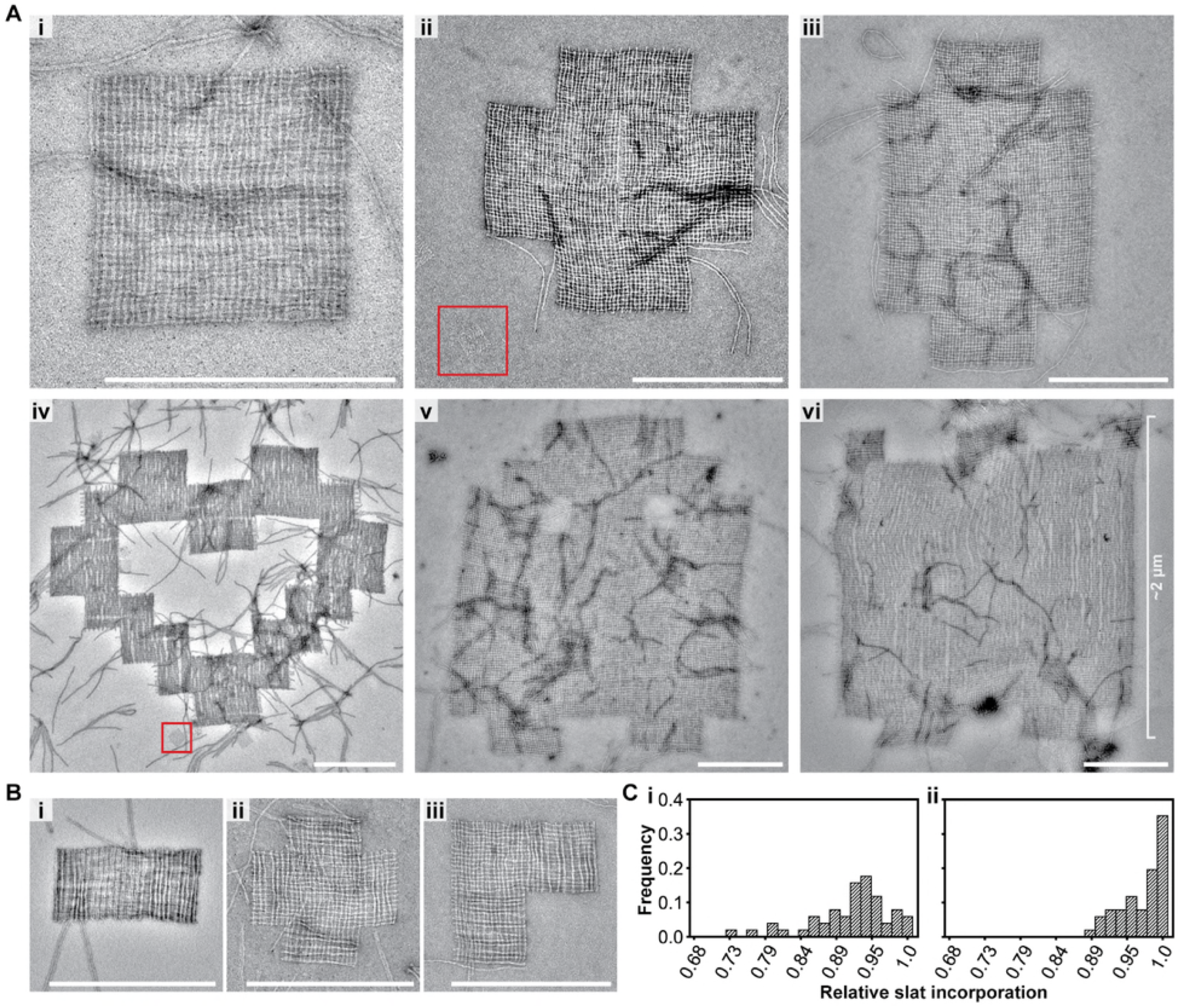
Assembly of finite megastructures from DNA-origami slats, where every slat is unique and addressable (see the designs in Fig. 1C). **A,** TEM images of megastructures composed entirely of 6HB slats, except for the 1022-slat rectangle that has 28 horizontal 12HB slats in *Avi*. The red boxed regions are a single origami reference square for size comparison, which is the largest area structure attainable with the same scaffold used for each slat. **B**, TEM images of finite megastructures where half or more of the slats are 12HBs. **C**, histogram for the number of slats counted in close-up TEM images of 50 randomly selected finite squares. The squares were assembled at 34°C overnight (*Ci*) versus assembled for an additional two days at room temperature ( *Cii*). Scale bars are 500 nm.

To assess the relative incorporation of the 6HB slats into the megastructures, we counted the number of slats in higher-magnification TEM images in 64-slat squares. We determined an average incorporation of ~90% of the slats after overnight isothermal assembly, increasing to ~97% after an additional two days at room temperature (Fig. 2Ci–ii), which is comparable to the ~80–90% full incorporation of a given staple strand into a typical DNA origami^36^. We also assessed the relative completion of the largest finite megastructures by concentrating the final samples and looking to see if the features of each shape (i.e. corners and middle sections of the shape were appropriately filled with slats) could be observed in lower-magnification TEM images. Greater than 22% of the megastructures for the 954-slat ghost and 1022-slat sheet were complete by the last assembly stage (N_954_ghost_ = 225, N_1022_sheet_ = 267, see Fig. S15). This suggests that over 75% of the assemblies at each stage were competent for continuing growth (i.e. 0.78^n^ = 0.225, where n = 6 growth stages, see caption of Fig. S15).

### Periodic ribbon and sheet megastructures

We used the strand library to create periodic 6HB-based crisscross ribbons and sheets (Fig. 3). We first explored the ribbons depicted in Fig. 3A, which grew bidirectionally from the first series of slats bound to the seed. In Fig. S16A–B, each slat added is staggered one binding site unit compared to the parallel slat that preceded it, as similar to previously established for ssDNA slats^33^. The ribbons in Fig. 3Ai versus Fig. S17 are termed version 16 (v16) and version 8 (v8), respectively. In v16, a given slat has 32 perpendicular slats bound to all of its 32 possible binding handles, versus v8 which only has 16 slats bound to every other of its 32 possible binding sites. As observed by TEM, v16 ribbons appeared as relatively uniform flat ribbons as expected. In contrast, v8 ribbons were much more flexible, and exhibited pronounced fluctuations in width along their lengths due to an accordion-style stretch. Furthermore, many v8 ribbons underwent full conversion to elongated spindles, although it is unclear from the images what the structure is (e.g. whether this a simple accordion-style stretch taken to an extreme, or instead these are twisted as well) (Fig. S18A–B). We also created ribbons with zigzag raster growth, where alternating clockwise then counter-clockwise sets of 16 slat additions creates jagged edges, while alternating two clockwise sets with two counter-clockwise sets creates flush edges (Fig. 3Aii versus Fig. 3Aiii, also see Fig. S16C–D and Fig. S18C–D). Ribbons of all three design types attained comparable mean lengths after 16 hours of isothermal incubation, despite the differences in programmed stagger (Fig. S19).

**Figure 3.**
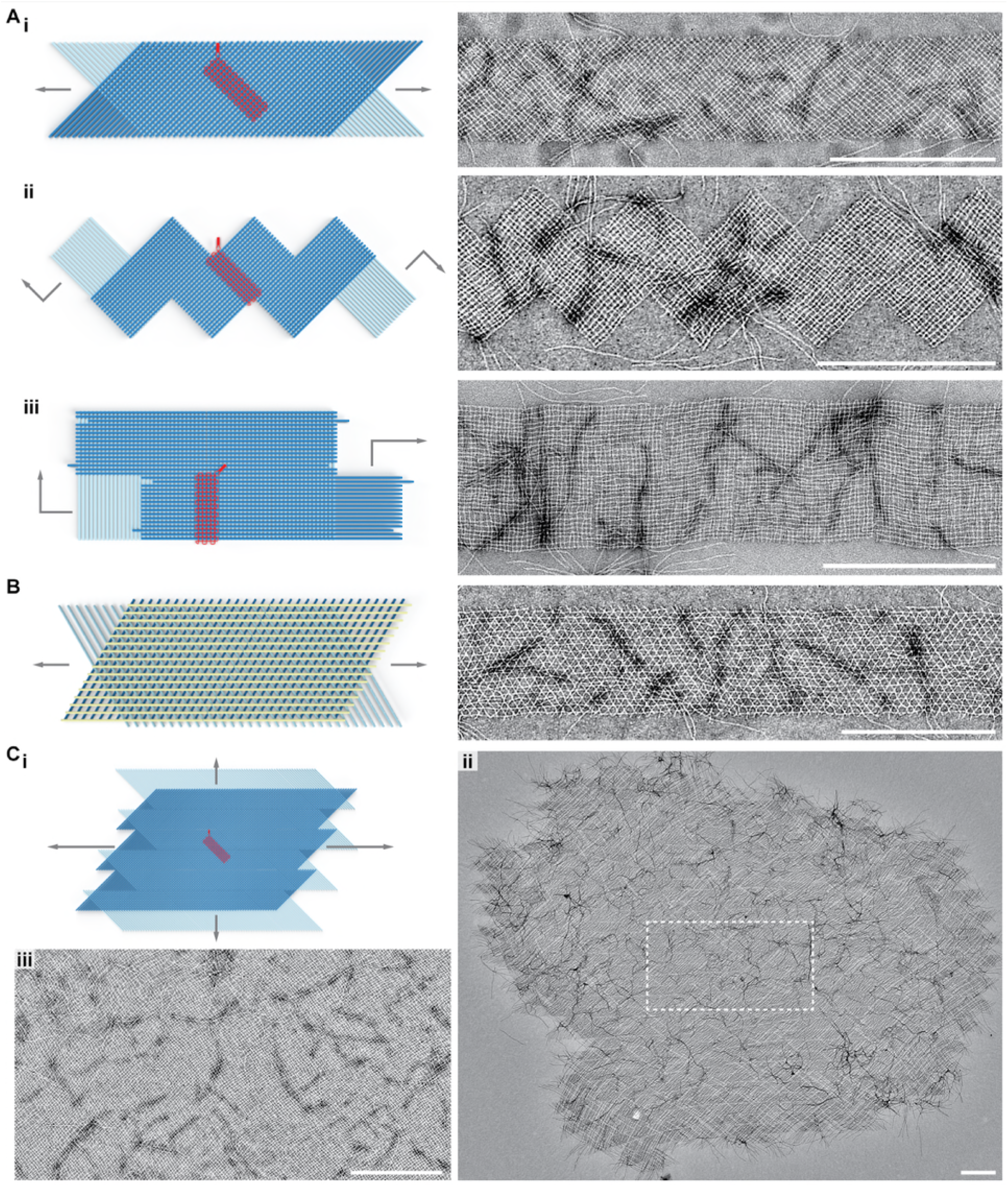
Assembly of periodic ribbons and sheets that grow with 6HB slats in one- and two-dimensions. **Ai**, v16 ribbon where top and bottom-layer slats are staggered such that addition occurs in alternating order. **Aii**, v16 ribbon where the slats are added in a zigzag raster-fill pattern that creates jagged edges. **Aiii**, v16 ribbon where the slats are added in a zigzag raster-fill pattern that creates flush edges. **B**, tri-layer arrangement of slats, where a top-layer of yellow slats rigidifies the otherwise flexible v8 ribbon. **Ci**, rendering of v16 growth from *Ai* where slats are shifted to enable formation of a sheet that grows in two dimensions. **Cii**, TEM image of a sheet, where the representative area from the **Ciii** close-up image is outlined with the dashed white box. Scale bars are 500 nm.

For all periodic designs, the size of the repeating set of slats was explored from 8 to 64 unique slats in top and bottom layers each (Fig. S20). However, most designs in this study were composed of 8 or 16 unique slats in the top and bottom layers each. We found that the apparent second-order rate constant for slat addition became progressively smaller as the overall slat concentration was increased to over 1 – 2 μM (Supplementary Text 3.1, Fig. S21, Fig. S22). Consideration of this limiting behavior motivated our strategy to grow our larger megastructures in multiple stages, sequentially adding subpools with only ~200 slats at a time to avoid lower than ~4 nM concentration of any one slat while maintaining total slat concentration close to 1 μM (Supplementary Text 3.2; rightward panels of Fig. S9C–D and Fig. S10; the megastructure in Fig. 2Aiii grew faster using the multi-stage protocol as shown in Fig. S14).

We also created periodic 2D structures with the 6HB slats. One approach was to incorporate an additional layer of slats to a v8 ribbon to show that megastructures with more than two layers are possible (Fig. 3B, Fig. S23). In this particular design, the bottom two layers of the v8 ribbon are locked into a rigid conformation where slats between layers are positioned 60° to one another. Next, we tested 2D sheets with two layers of slats (Fig. 3C). The design of the sheets is equivalent to the v16 staggered 1D ribbons shown in Fig. 3Ai, but with the bottom layer of slats shifted half of their lengths with respect to the top layer (Fig. S24). These sheets, in this case defined by 16 + 16 slat unit cells with lateral dimensions of ~320 nm and containing 512 addressable nodes, typically grew to significant dimensions after three days of isothermal growth; the rightward example in Fig. 3Cii is composed of ~10,000 slats, with a molecular mass exceeding 50 GDa and lateral dimensions of ~10 μm (also see additional sheets in Fig. S25). The higher-magnification TEM image Fig. 3Ciii shows a typical middle region of a sheet, with a fabric-like character where defects such as missing slats were infrequent. Both the 1D ribbons and 2D sheets were of sufficiently large molecular mass that they could be extracted from the excess unpolymerized slats by sedimentation into a pellet via centrifugation at 2500 RCF (Fig. S26). We also note that the ribbons were stable at room temperature for two days in MgCl2 as low as 4 mM (versus the 15 mM condition during growth, see Fig. S27).

### Addressability of megastructures

To illustrate that origami-crisscross megastructures can be functionalized as large addressable canvases, we designed the 1022-slat sheet and periodic sheets to display custom patterns of handles on their top faces (Fig. 4A–B). We assembled a 10 nm DNA nanocube^37^ contrast agent bearing a single complementary handle, incubated the patterned sheets with the purified nanocube, and visualized the resulting patterns using negative-stain TEM (Fig. S28). The 1022-slat sheets in Fig. 4A were observed to display the programmed patterning of intricate designs including a jigsaw puzzle piece, a happy face, and institutional logos for some of our affiliations. Each ~2.0 μm × ~1.8 μm DNA canvas contains 16,128 addressable nodes, spaced ~14 nm apart. The nanocube was further used to decorate patterned 2D sheets, with the left panel of Fig. 4B showing the smaller 512-node canvas and the right panel showing a TEM overview of a checkerboard, a polkadot sheet, and a continuous jagged line. We additionally characterized the periodic 2D sheets and 1D ribbons using DNA-PAINT. We resolved single handles when spaced ~57 and ~43 nm (i.e. 168 and 126 bp, respectively) between adjacent handles (Fig. 4C, Fig. S29, Fig. S30, Fig. S31).

**Figure 4.**
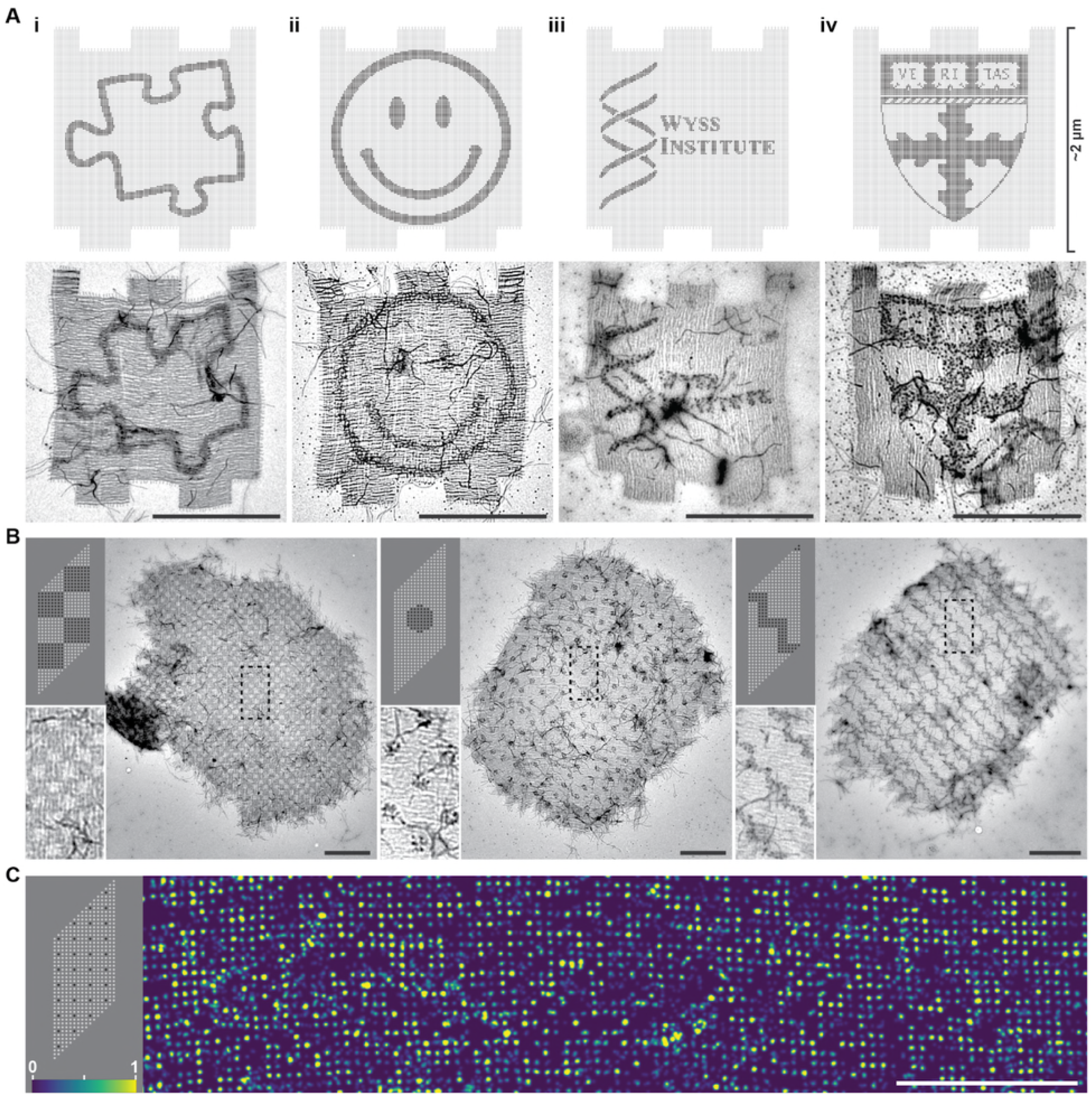
Finite and periodic crisscross megastructures as addressable DNA canvases to pattern arbitrary cargo. **A,** top row, designs of the finite 1022-slat sheet, with darker dots indicating sites that were programmed with a handle sequence to bind a DNA-nanocube contrast agent, with patterns including the outline of a jigsaw puzzle piece (**i**), a happy face (**ii**), and the crests for the Wyss Institute of Harvard University (**iii**) and the Harvard John A. Paulson School of Engineering and Applied Sciences (**iv**). Bottom row, TEM images of the patterned finite sheets. **B**, TEM images of 512-node periodic-sheet canvases patterned with DNA nanocubes, with the upper-left panels showing the designs. Boxed regions are shown more closely in the bottom-left panels. **C**, DNA-PAINT image of single handles on the sheets, as indicated in the top left design panel. Relative imager strand on-time is denoted by color as shown in bottom-left. Scale bars are 1 μm.

### Nucleation control, growth kinetics, and design principles

There was no observable formation of either finite or periodic megastructures in the absence of an added seed (Fig. 5A–B, Fig. S32). As a benchmark, we folded a single DNA-origami square with a constant concentration of staple strands (400 nM each) and variable concentrations of scaffold (0.5–50 nM), and observed a linear one-for-one dependence in the concentration of squares formed (Fig. 5Ci, Fig. S33). We proceeded to mix constant concentrations of slats with variable concentrations of seed and indeed observed a similar one-for-one stoichiometric dependence in the megastructures formed (Fig. 5ii–iii, Fig. S34).

**Figure 5.**
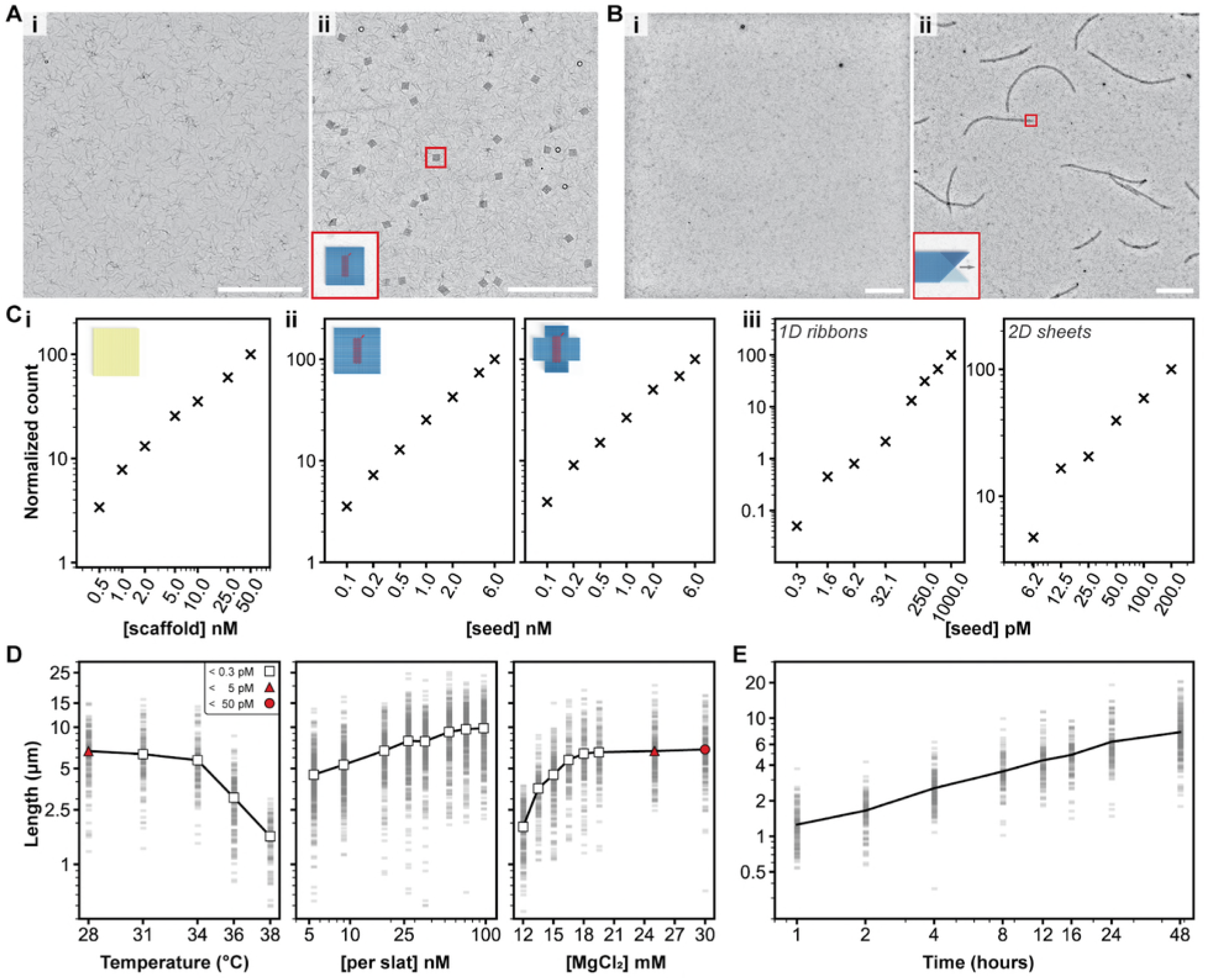
Characterization of growth versus reaction parameters. **A** and **B**, seed-controlled assembly of the finite 64-slat square and the periodic 1D ribbon in low magnification TEM images, with no seed versus seed in *i* and *ii*, respectively. **Ci,** the number of DNA-origami squares formed versus the amount of scaffold added, **Cii**, the number of finite squares (left) or plus symbols (right) versus amount of seed added, and **Ciii**, the number of periodic megastructures versus the amount of seed added. The relative number of particles per condition is shown with the ‘**×**’ marker. **D**, how the length of v16 1D ribbons with 7-nt binding sites varies as a function of temperature, concentration of slats, and concentration of MgCl_2_. Each faint gray bar is a single ribbon measurement. The white markers indicate no spontaneous assembly above the detection limit, versus red markers where spontaneous assembly was observed to the degree shown in the legend in the leftmost plot. **E**, the length of the v16 1D ribbons versus time, grown at 20 nM of each slat. Axes in all plots are on a log_10_ scale, with the exception of the temperature and MgCl_2_ scale in *D*. All scale bars are 5 μm, particle counts in *C* were determined by counting structures in ten low-magnification TEM images, and ~150 ribbons were measured per condition in *D* and *E*.

To quantitatively assess spontaneous nucleation under various reaction conditions, we compared seeded v16 7-nt ribbons to unseeded control reactions with variations to temperature, concentration of slats, or concentration of Mg^2+^ (Fig. 5D). Any unseeded control reactions where no ribbons were observed were presumed to have fewer than 0.3 pM ribbons, as per our limit of detection (Fig. S35, Fig. S36). No spontaneously formed ribbons were observed above 28°C (leftmost Fig. 5D). Growth could not be observed above 38°C, and fully formed ribbons were observed to shrink at temperatures above ~42°C (Fig. S37). Thus we selected 34°C as a reliable temperature for robust and fast seed-controlled growth. Spontaneous nucleation was also not evident at this temperature across a broad window of concentrations of the slats and Mg^2+^ (middle and rightmost Fig. 5D). Hence, growth of the v16 7-nt ribbons is strictly seed-dependent at temperatures well below where ribbons begin to shrink, using high concentrations of each slat and Mg^2+^ of up to 100 nM and ~20 mM, respectively. We further characterized relative spontaneous nucleation of v8 and v16 ribbons using weaker 6-nt and stronger 8-nt handles (Supplementary Text 4, Fig. S37–Fig. S39).

The apparent second-order rate constant for slat addition, which we estimated at ~10^6^ M^-1^s^-1^, was remarkably high (Supplementary Text 5, Fig. 5E, Fig. S40). It is notable that these observed kinetics are comparable to those for hybridization of DNA strands^38^, or up to two-orders of magnitude faster than some other approaches to assemble DNA origami, such as blunt-end stacking of shape-complementary features (although these carry the advantage of stronger penalization of non-cognate interactions)^39^. Finally, we explored principles for combinatorial design of slats — using our 2048 7-nt handle library allowing for 32 possible handles at each of 32 positions — that are sufficiently orthogonal to support robust growth of complex megastructures (Supplementary Text 6, Fig. S41–Fig. S45).

### Conclusion

We generalized crisscross polymerization to DNA-origami slats for growth of diverse finite shapes including an addressable canvas from 1022 unique slats that spans about two micrometers per side, periodic ribbons with several different extension patterns, and periodic sheets that attained lateral dimensions exceeding ten micrometers. Hierarchical self-assembly with these building blocks exhibited several features that are advantageous for rapid and robust nanoconstruction: (i) strict seed dependence of initiation, compatible with addition of slats at relatively high concentrations and in stages; (ii) relatively low defect rate in terms of missing slats and prematurely terminated megastructures, implying robustness in spite of inherent defects in the origami building blocks (e.g. unavailable handles)^36^, (iii) highly orthogonal behavior that enables hundreds of distinct slats to assemble correctly in a single pot; (iv) fast apparent growth kinetics (10^6^ M^-1^s^-1^) despite the 5 MDa size of the building blocks. Moreover, structural diversity was created by mixing and matching strands from a library of only 2048 staple strands, where each binding handle of the slats was encoded with one of 32 possible sequences. Therefore prototyping diverse megastructures in this way is cost effective.

In future studies, the design of the DNA-origami slats could be tailored to create a larger diversity of megastructure architectures^29^, and various routes to more three-dimensional structures should be accessible^4,40–42^. As with other tiling approaches, it may be possible to program growth with sophisticated algorithmic behaviors^43–45^. The resulting megastructures could provide access to templates for patterning of diverse guests, such as functional proteins and optically active nanoparticles, on length scales comparable to those of biological cells.

## Supporting information

caDNAno files in json format

sequences in Excel format

## Acknowledgments

The authors would like to thank the following individuals: Jocelyn (Josie) Kishi for suggesting and helping write a grant for the Echo Acoustic Liquid Handler that made this work possible; Serkan Cabi and Tao Zhang for helping test early designs of crisscross origamis; Prof. Vinothan Manoharan, Prof. Michael Brenner, Rasmus Sørensen, and Jaeseung Hahn for fruitful discussions.

## Author contributions

- Conceptualization: CMW, DM, AE, JFB, WMS
- Methodology: CMW, DM, AE, WMS
- Software: CMW, DM, AE, GG
- Validation: CMW, DM, AE
- Formal analysis: CMW, DM, AE, HMS, GG
- Investigation: CMW, DM, AE, HMS, GG, JFB, FECD
- Writing (Original Draft): CMW
- Writing (Review & Editing): CMW, DM, AE, HMS, GG, FECD, WMS
- Visualization: CMW, DM, AE, HMS, JFB
- Supervision: CMW, PY, WMS
- Funding acquisition: CMW, DM, AE, PY, WMS

## Data and materials availability

All raw TEM image data that was measured to determine growth and nucleation of origami crisscross megastructures, scripts that were used to make various assignments of handle staple oligonucleotides, and scripts that were used to measure Hamming distances of sequence assignments are available upon request from WMS.

## Supplementary Materials

### The Supporting Information document contains the following

Materials and Methods
Supplementary Text 1–6
Figs. S1–S45
Tables S1 to S8
Note: References (*46–52*) are only cited in the Supporting Information

### Supplementary Data S1 to S8 are in two compressed files which contain the following

S1–S4 Spreadsheets of sequences, finite assignments, periodic assignments, and nanocube/PAINT patterns, respectively.
S5–S8 DNA origami caDNAno v0 designs of the 6HB slat, 12HB slat, gridiron seed, andsingle reference square, respectively.

