## Supplementary Materials pdf for "Multi-micron crisscross structures from combinatorially assembled DNA-origami slats"

### Table of contents

Materials and methods

Supplementary Text 1: Design notes regarding the 12HB slat

1.1 Derivation of 12HB slats from 6HB slats

1.2 Yield differences and the consequence on downstream usage of the slats

Supplementary Text 2: Algorithm for selection of the handle sequences

2.1 Initial sequence generation

2.2 Optimizing for sequence orthogonality

2.3 Discussion about the final sequences and pertinence to megastructures

Supplementary Text 3: Relationship between the number of unique slats in a design versus the relative rate of growth

3.1 Growth slows when large numbers of unique slats are mixed simultaneously

3.2 Growth of finite megastructures with large numbers of unique slats can be accelerated by adding the slats in multiple steps

Supplementary Text 4: Detailed characterization of binding handles

4.1 How the number of binding sites (i.e. v16 or v8) versus temperature influences spontaneous nucleation

4.2 How the base-pairing strength of the binding site handles versus temperature influences growth rate and spontaneous nucleation

Supplementary Text 5: Kinetics of ribbon assembly

5.1 Calculation of the number of slats added to the ribbon

5.2 Approximation of growth using pseudo-first order kinetics

5.3 How the base-pairing strength of the binding-site handles influences growth rate

Supplementary Text 6: Sequence assignment to megastructures from the 2048-handle sequence library, optimization of the Hamming distances of slats within megastructure designs

6.1 Increased pairwise complementarity between slats (i.e. minimized Hamming distances) causes drastic slowdown of 1D ribbon growth

6.2 Possible mechanism for how complementarity between the slats causes growth slowdown

6.3 Discussion of the Hamming distances used in the finite and periodic megastructure designs

Supplementary Figures Fig. S1–Fig. S45

Fig. S1–Fig. S8: Slat and seed design and folding, qualitative energetics of crisscross assembly, selection of initial binding-site strength, energies of various binding-handle sets, and need for T-linkers

Fig. S9–Fig. S15: Models of finite megastructures, preparation time for each design, and additional TEM results of finite megastructures

Fig. S16–Fig. S27: Models of periodic megastructures, growth comparison of different ribbon designs, relationship between growth rate and number of unique slats, and additional TEM results of periodic megastructures

Fig. S28–Fig. S31: Model of the DNA nanocube, additional TEM of nanocube patterns, and DNA-PAINT results of 1D ribbons and 2D sheets

Fig. S32–Fig. S34: TEM results when no seed added, AGE results of single DNA origami square versus scaffold, representative TEM of megastructures versus concentration of seed

Fig. S35–Fig. S40: Standard curve of ribbons, melt temperature of v16 7-nt ribbons, temperature characterization and melt temperature of v8 7-nt, v16/v8 6-nt, and v16/v8 8-nt ribbons, growth versus time for 6-nt and 8-nt v16 ribbons

Fig. S41–Fig. S45: Hamming-distance analysis of 1D ribbon growth, mechanistic testing for growth changes to 1D ribbons vs. kinetic trapping of slats resulting from Hamming distance, and optimized Hamming distances of all the megastructure designs tested in this work

Table S1–Table S8: Core-strand sequences for 6HB, 12HB, seed, and single origami reference square; Sequences of the 6-, 7-, 8-, 9-, and 10-nt handles; Nanocube strand sequences

### Materials and methods

**Method 1 Design and purchasing of handle staple strands:** Binding sequences for the various handles were selected as explained in Supplementary Text 2. The handle sequences (and the complementary handle sequences) were appended to the 3′ end of either the top or bottom helix staple strands of the 6HB slat, where they were separated with a thymine (2T) linker. The binding handle sequences are reported in Table S6 and Table S7 (and also in Supplementary Data S1 as a spreadsheet), and the importance of the linker sequence is shown in Fig. S8. All strands were purchased dry at full yield at the 10 nmol scale from Integrated DNA Technologies (IDT). We preferred to purchase the strands on Echo 525 compatible source plates (Labcyte #PP-0200) directly from IDT, though sometimes ordered the strands on other plates and later transferred them to Echo 525 compatible source plates using a manual multichannel pipette. All strands were rehydrated in 50 µL of water, with their concentration assumed to be ~200 µM.

**Method 2 Design of sequence assignments (i.e. permutations) of the handles from the library:** See Supplementary Text 6 for an explanation of why care had to be taken in choosing permutations as described below, and Supplementary Data S2–S3 for the final permutations. Blank layouts of top- and bottom-layer slats were drawn out in Microsoft Excel sheets after the megastructure design was initially conceived. Python scripts were then used to populate each cell of the blank Excel sheet randomly with a number ranging from 1 to 32, corresponding to a specific handle sequence from the 2048 strand library. Next, the script converted the assigned sequences into a set of top-layer slats and bottom-layer slats, where each slat is defined as a one-dimensional list that is 32 numbers long. The Hamming distances between the slats were measured to determine the number of handles on a given slat from one layer that matched together with complementary handles with each slat in the other layer. The process of random assignment and measurement of the Hamming distances in the resulting slats was repeated until some arbitrary maximum threshold of allowable kinetic-trap strength (i.e. the minimum Hamming distance) was attained. We note that megastructure designs that were composed of larger numbers of unique slats tended to have more undesired matched complementarity compared to designs composed of smaller numbers of unique slats. One possible solution to further maximize the Hamming distances for a design would be to increase the size of the handle library (i.e. using > 32 different possible 7-nt handles at each slat intersection with a strand library that has > 2048 unique handle strands versus what was tested here). Nonetheless, we were able to attain satisfactory growth of megastructures that were composed with up to 1022 unique DNA slats that were generated from the 2048 7-nt handle library.

**Method 3 Pooling handle strands for each 6HB and 12HB slat and other folding details for preparation of slats on 96-well plates:** Liquid handling protocols for each design were written to a csv file using Python scripts referencing the Excel sheet of the megastructure design (see Method 2). Staple strands with or without handles were added into 96-well PCR plates (Eppendorf E0030129512) from the 384-well strand library using a Labcyte Echo 525 acoustic liquid handler, which read the csv file using Echo Cherry Pick v1.7.2. Each particular 6HB or 12HB slat respectively required transfer of 64 or 32 strands for the customizable top and bottom helices of the slat. We note that handle staple strands could also be pooled together manually using single or multichannel pipettes, though such preparations could require an inordinate amount of time depending on the number of slats in the megastructure. See Fig. S11 for a time comparison of such approaches. After strand transfer into the 96-well plates, any droplets along the rim of the wells were spun down, at which point a slat-folding mixture containing other core strands, scaffold, and buffer as specified in Method 4 was added to each well and mixed using a manual multichannel pipette. The 96-well plates were then thermally sealed with plastic films using an ABgene ALPS-300 microplate sealer and spun down one final time on a centrifuge, at which point plates were placed on a thermocycler and the origami folded using the temperature gradient in Method 4.

**Method 4 Design and folding of DNA origami:** All DNA origamis were designed using legacy caDNAno[46] v0. See Supplementary Data S5–S8 for caDNAno json files of the 6HB, 12HB, gridiron seed, and single origami reference square. Staple and scaffold DNA sequences are reported in Table S1–Table S5 (and also in Supplementary Data S1 as a spreadsheet). Unpurified dehydrated staple oligonucleotides were purchased from Integrated DNA Technologies (IDT) at 100 or 10 nmol scale. For the slats, each unpurified core staple strand (i.e. strands other than the 64 or 32 customizable handle strands for the top and bottom helices of the 6HB and 12HB slats, respectively) were rehydrated water at ~1 mM and pooled together with equal volumes of each strand. For the gridiron seed and origami reference square, each dehydrated strand was resuspended in water at ~100 µM and pooled together with equal volumes of each strand. The p8064 and p8634 scaffold strands were produced from M13 phage replication in *Escherichia coli*. All folding was done in 1x TE buffer (5 mM Tris pH 8.0, 1 mM EDTA) with the $MgCl_2$ as specified below. The gridiron seed was mixed with 40 nM p8634 scaffold, ~250 nM of each staple strand, 10 mM $MgCl_2$, and folded with a 12-hour temperature gradient: 94–86°C in 5 min steps less 4°C/step; 85–70°C in 5 min steps less 1°C/step; 70–40°C in 15 min steps less 1°C/step; 40–25°C in 10 min steps less 1°C/step; and 16°C thereafter until the sample was collected. The 6HB slats were mixed with 50 nM p8064 scaffold, ~500 nM of each staple strand, 6 mM $MgCl_2$, and folded with an 18 hour temperature gradient: 80°C/10 minutes in a single step; 60–45°C in 160 6.75 min steps less 0.1°C/step; 16°C thereafter until collection of the sample. The 12HB slats were mixed with 50 nM p8064 scaffold, ~500 nM of each staple strand, 8 mM $MgCl_2$, and folded with an 18 hour temperature gradient: 80°C/10 minutes in a single step; 75–45°C in 310 3.48 min steps less 0.1°C/step; 16°C thereafter until collection of the sample. The origami reference square was mixed with 40 nM p8064 scaffold, ~400 nM of each staple strand, 6 mM $MgCl_2$, and folded with an 18 hour temperature gradient: 80°C/10 minutes in a single step; 60–45°C in 160 6.75 min steps less 0.1°C/step; 16°C thereafter until collection of the sample. We note that there were special considerations for preparation of the slats in 96-well plates, as explained in Method 3.

**Method 5 Preparation of the DNA nanocube contrast agent:** Unpurified dehydrated nanocube oligonucleotides were purchased from Integrated DNA Technologies (IDT) at 10-nmol scale. Unpurified nanocube strands were rehydrated in water at ~100 µM each and pooled together with equal volumes per strand. Out of the 28 total nanocube strands as published previously[37], a single strand was selected and appended with a 4T linker and 16-nt handle to its 3′ end (see sequences in Table S8 or Supplementary Data S1). The nanocube was prepared with ~1 µM of each strand (with the handle-tagged strand at ~2 µM), 40 mM $MgCl_2$, and folded with a 42-hour temperature gradient: 80°C/10 min in a single step; 65–37°C in 290 8.69 min steps less 0.1°C/step; 16°C thereafter until collection of the sample. The folded nanocube was separated on an agarose gel and purified as described in Method 8, and bound to the megastructures without any further downstream assembly.

**Method 6 Agarose-gel electrophoresis:** Gel characterization of the DNA-origami gridiron seed, 6HB slats, or 12HB slats was performed using the Thermo Scientific™ Owl™ EasyCast™ B2 electrophoresis system. UltraPure agarose (Life Technologies, 16500500) was melted in 0.5x TBE (45 mM Tris, 45 mM boric acid, 0.78 mM EDTA, 11 mM $MgCl_2$) to a concentration of 1.0 % (w/v). The molten agarose was cooled to 65ºC and $6.25*10^{-5}$ % (w/v) ethidium bromide was added. To assess folding, ~50 fmol of DNA-origami sample was mixed in an excess of agarose-gel loading buffer (5 mM Tris, 1 mM EDTA, 30% w/v glycerol, 0.025% w/v xylene cyanol, 10 mM $MgCl_2$; with typically 4 µL loading buffer added to 1 µL of each sample that was folded with 50 nM scaffold). The mixed samples were loaded onto the gel and separated for 3–4 hours at 60 V at room temperature. Control samples for size and densitometry included one or both of the following: first, ~50 fmol of the same scaffold from which the DNA origami was folded; or second, ~0.5 µg of Gene Ruler 1kb Plus DNA Ladder (Thermo Scientific™ SM1331). Gel images were captured on a GE Typhoon FLA 9500 fluorescent imager using the ethidium-bromide parameters as given in the Typhoon control software. The photo-multiplier tube (PMT) was set to 500V. Densitometry to quantify relative assembly of DNA bands was performed with FIJI ImageJ (v1.53c)[47]. Background subtraction with a rolling ball radius of 30–60 pixels was performed on linear TIFF images. The GelAnalyzer plugin in ImageJ and wand tool was used to integrate total pixel intensities from lanes of interest. DNA-origami yields were determined by taking the ratio of the intensity of the band of interest with respect to all the species with a molecular weight larger than the excess staple strands.

**Method 7 PEG precipitation to concentrate pools of 6HB slats:** The slats as folded on 96-well PCR plates were collected and combined into pools using a manual multichannel pipette. Each pool maximally had ~100 slats, though the number of slats in a given pool was variable depending on the design of the megastructure. We generally kept the slats in one layer of the megastructure in a separate pool from the perpendicular slats in the other layer of the megastructure, so that slats with complementary handles would not be concentrated in the same mixture together. Our rationale was that spontaneous interactions between the slats during concentration could be deleterious to the yield, though we did not study this carefully to determine whether such care was necessary. The pooled slats were subsequently concentrated using two rounds of PEG precipitation, as adapted from as published previously[48]. The $Mg^{2+}$ in the slat pool was increased from 6 mM to 20 mM by adding the appropriate volume of 1 M $MgCl_2$. One volume of 2x PEG-purification buffer (5 mM Tris, 1 mM EDTA, 15% w/v PEG-8000, 510 mM NaCl) was added and mixed with the equal volume of pooled slats in a 2 mL round-bottom tube. The mixture was spun at 16k RCF for 25 min, the supernatant was gently extracted using a pipette, and the pellet was resuspended in 50 µL 1x TE buffer with 20 mM $MgCl_2$. The second round of PEG precipitation was performed using an equal volume of 2x PEG-purification buffer and the final slat pellet resuspended in a small volume of 1x TE buffer with 10 mM $MgCl_2$ so that the total concentration of slats was ~2 µM. However, the volume was adjusted as needed so that the pool was sufficiently concentrated to achieve the desired final per slat concentration in Method 9 or Method 10. The final slat pool was placed on a shaking incubator set to 1000 rpm at 33°C for about one hour. Finally, the concentration of DNA was measured on a Nanodrop 2000c Spectrophotometer (Thermo Scientific™) so as to estimate the final nM concentration of slats.

**Method 8 Agarose-gel extraction to purify pools of 12HB slats:** The 12HB slats in raw folded samples were low yielding and insufficient to prepare as described exactly in Method 7. See Supplementary Text 1, Fig. S3 and Fig. S4 where the yield and resulting challenges are discussed in detail. The 12HB slats were pooled as described in Method 7 and then separated on an agarose gel as described in Method 6. The gel was examined on a UV transilluminator to identify the monomer band which was then excised from the gel using a razor blade. The gel-band pieces were transferred to a 1.5 mL eppendorf tube, crushed using a plastic pestle, and purified using Freeze N' Squeeze spin columns (Bio-Rad, 732-6166) as published previously[48]. The gel-purified samples was typically too dilute, such that it was necessary to concentrate them into a smaller volume of 1x TE buffer with 10 mM $MgCl_2$ using one round of PEG precipitation as explained in Method 7.

**Method 9 Preparation of megastructure assembly reactions with fewer than 200 unique slats:** The raw gridiron seed (folded with 40 nM scaffold), purified slats (generally ~1 µM total slats per pool depending on the extent to which it was concentrated during purification), and 4x Megastructure buffer (5 mM Tris pH 8.0, 1 mM EDTA, 30 mM $MgCl_2$, 0.04% Tween-20) were mixed together in 5–20 µL at room temperature. We generally prepared the reactions in 0.1 mL Eppendorf PCR tubes (E0030124812) because of their tightly fitted lids which guarded against evaporative loss during extended growth periods. The final reaction generally contained ~0.5–2 nM seed, ~5–20 nM of each unique slat, 1x Megastructure buffer, with any excess volume filled with 1x TE buffer with 10 mM $MgCl_2$. We assumed that the seed and slats would be in 1x TE buffer with 10 mM $MgCl_2$, such that the final megastructure assembly reaction would contain 15 mM $MgCl_2$. Next, the reactions were placed on a thermocycler and incubated for 4 hours at a high temperature where slats could only bind the seed (i.e. 45°C or 55°C for the 6/7-nt or 8-nt designs, respectively), before the temperature was lowered for slat growth for 1–72 hours. We used 34°C for typical growth of the v16 periodic and finite megastructures using the 7-nt handle library, versus other designs which used growth temperatures as summarized with cyan markers in the Fig. S39 plot.

**Method 10 Preparation of megastructure assembly reactions with more than 200 unique slats:** As explained in Supplementary Text 3, megastructures composed of a large number of unique slats (i.e. > 100 slats) generally grew slowly if all the slats were mixed simultaneously. Hence, designs with more than 100 unique slats were either incubated for a longer growth time, or grown in stages with several additions of slats. The first growth stage with ~200 of the slats most proximal to the seed were prepared as in Method 9. We prepared a 4x MultiMegastructure buffer (5 mM Tris pH 8.0, 1 mM EDTA, 22 mM $MgCl_2$, 0.024% Tween-20). In each stage thereafter, 5 parts of 4x MultiMegastructure buffer were added to 8 parts of an aliquot of the reaction from the prior stage with ~200 of the next slats in 20 parts total. The purpose of the 4x MultiMegastructure buffer was to maintain the buffer with 5 mM Tris pH 8.0, 1 mM EDTA, 15 mM $MgCl_2$, and 0.01% Tween-20 as more slats were added.

**Method 11 Binding of the DNA nanocube to 6HB-slat canvases:** An aliquot of the periodic sheets or 1022-slat finite sheet was diluted 10–50 fold into agarose-gel purified nanocubes from Method 5. The sample was incubated at room temperature overnight to bind the nanocube to the megastructures. The subsequent day, the sample was purified from excess slats and nanocubes using two rounds of centrifugation, as outlined in Method 12. The sample was resuspended in 1x TE buffer with 10 mM $MgCl_2$ and 10% trehalose (w/v) and negatively stained with 1% uranyl formate, as described in Method 13. We qualitatively noted that trehalose and the lower percentage (i.e. versus 2%) uranyl formate provided better contrast for the nanocubes on the megastructure canvas.

**Method 12 Purification of periodic-ribbon and sheet megastructures by centrifugation:** The largest periodic ribbons and sheets were purified from excess free monomers by centrifuging the samples at low speed. See Fig. S26 for results using this method. Ribbons or sheets were prepared as in Method 9, diluted

~10 fold in 1x TE buffer with 10 mM $MgCl_2$, and then spun in a 1.5 mL eppendorf tube at 2.5k RCF for 15 minutes. The supernatant was gently extracted with a pipette and the pellet resuspended in 1x TE buffer with 10 mM $MgCl_2$ with the final volume as desired for the particular application. This procedure of centrifugation and resuspension could be repeated multiple times to further deplete excess free slats.

**Method 13 Transmission electron microscopy (TEM):** Raw assembled megastructure reactions were diluted 1:250–1:1000 in 1x TE buffer with either 10 mM or 15 mM $MgCl_2$. TEM grids (Electron Microscopy Sciences; FCF400-CU, FCF200-CU-TA, or FCF100-CU-TA) were negatively glow discharged at 15 mA for 25 s in a PELCO easiGlow. The diluted sample (4 µL) was applied to the glow-discharged grid, incubated for 2 min, and wicked off gently into Whatman paper (Fisher Scientific, 09-874-16B). We used one of two possible approaches to stain the sample with 2% uranyl formate: (1), When closer-up structural detail of individual slats was desired, we used a negative-stain approach where 4 µL of 2% aqueous filtered uranyl formate was applied, incubated for 1–2 seconds, and gently wicked from its side into Whatman paper so as to leave a thicker deposit of uranyl formate around the sample. Or (2), when clarity of the overall forms and length of the megastructure was desired in low-magnification images, we used a positive stain approach where 4 µL of 2% aqueous filtered uranyl formate was applied, incubated for 1–2 seconds, and wicked off completely into Whatman paper so as to leave a thinner layer of uranyl formate, which darkened each particle with respect to the grid substrate. All imaging was performed at 80 kV on a JEOL JEM 1400 Plus microscope. All images presented in this work were imported into FIJI ImageJ (v1.53c)[47], corrected for background noise using a pseudo flat-field image, and then contrast and brightness adjusted for clarity in publication.

**Method 14 Quantification of slat incorporation in TEM images:** We used the finite 64-slat square as a model system to assess the relative incorporation of a given slat into a megastructure (see Fig. 2). Each sample was assembled for a given period of time, diluted 750-fold so that single squares could be differentiated from excess free slats on the TEM substrate, negatively stained with uranyl formate, and then imaged by TEM. Images of ~50 well stained squares were collected at a magnification where the single slats could be differentiated. The slats in each layer of the square were quantified using the Cell Counter plugin in FIJI ImageJ (v1.53c)[47].

**Method 15 Quantification of completion of large finite megastructures in TEM images:** Aliquots of megastructures with ~1000 unique were concentrated ~10-fold and applied to TEM grids using a positive stain approach (see Method 12 and Method 13, respectively). Sufficient numbers of low magnification TEM images were collected so that ~200–300 individual megastructures could be assessed for their relative completion (i.e. determining if the larger morphological features of the design were present with no observable missing segments). We alternatively considered doing Method 14 for the largest finite structures and periodic ribbons and sheets; however, the large number of slats in the megastructures, variable sizes of the periodic designs, and manual approach to identifying the slats in images made this approach untenable.

**Method 16 Quantification of relative ribbon growth in TEM images:** We used periodic ribbons to determine how parameters including temperature, concentration of $Mg^{2+}$, concentration of the slats, time, strength of each binding site, the number of crisscross binding sites, growth pattern, and sequence assignment influences growth (see Fig. 5D–E, Fig. S37–Fig. S38, Fig. S19, Fig. S42–Fig. S43). We assembled ribbons using 0.5 nM seed as explained in Method 9, diluted the initial sample 250-fold, and applied it to a grid where the structures were positively stained with uranyl formate. Each sample was then imaged at low magnification with TEM. We collected enough images so that at least 150 ribbons could be measured. The lengths of the ribbons were calculated using the NeuronJ plugin in FIJI ImageJ (v1.53c)[47,49].

**Method 17 Quantification of stoichiometric control of megastructure formation from the seed in TEM images:** We counted the number of megastructures in low-magnification TEM images of reactions where different concentrations of seed were added (see results in Fig. 5C). The raw samples were diluted so as to be able to see single particles in the condition where the highest amount of seed was added. We positively stained the sample with uranyl formate, and collected ten TEM images at a suitable magnification where the structure in question could be identified, but still allowed us to observe a reasonably large number of particles. The number of megastructures in each image was counted using the Cell Counter plugin in FIJI ImageJ (v1.53c)[47]. We normalized the number of structures counted for a given seed concentration with respect to one of the concentrations tested and plotted the normalized count versus the concentration of seed (or scaffold) added.

**Method 18 Quantification of spontaneous nucleation of ribbons in TEM images:** We counted the formation of periodic ribbons in control reactions with no seed to quantify the amount of spontaneous nucleation. We considered the ribbons useful as a model for spontaneous nucleation of megastructures because they were easy to identify as micrometer-sized objects that could be readily counted in low-magnification TEM images (see results in Fig. 5D and Fig. S38). Samples were incubated at the reaction conditions that we desired to test, diluted ~250 fold, and added to the TEM substrate, positively stained with uranyl formate, and then ten low-magnification TEM images were collected. The number of ribbons in each image was counted using the Cell Counter plugin in FIJI ImageJ (v1.53c)[47]. The ribbons counted in each image area were normalized to the dilution and magnification, and then converted into a molar amount by comparing it to a standard curve of ribbons (see Fig. S35). The recorded ribbon detection limit was ~0.3 pM, which corresponded to observing a single ribbon in the ten images.

**Method 19 Preparation of megastructures for DNA-PAINT imaging:** The outside faces of slats on ribbons and sheets were decorated on one side with 3′ complementary handle sequences for PAINT imager strands and on the other side with 5′ biotin sites. Slats were folded, purified, assembled into ribbons and sheets, and purified from excess slat monomers and resuspended in buffer B (5 mM Tris-HCl pH 8.0, 10 mM MgCl2, 1 mM EDTA, 0.05% v/v Tween-20) at a final concentration of ~0.1 nM per megastructure as described in Method 4, Method 7, Method 9, and Method 12, respectively. The megastructure sample was further prepared as described previously[50], where an imaging chamber with an inner volume of ∼20 µl was created by adhering one coverslip (#1.5, 18 × 18 mm2, ∼0.17 mm thick) to a glass slide (75 × 26 mm2, 1 mm thick) with double-sided tape. Surfaces of the chamber were prepared by the following steps: (1) flowing 20 µl of 1 mg/ml biotin-labeled bovine albumin (Sigma-Aldrich, catalog number A8549) dissolved in buffer A (10 mM Tris-HCl (pH 8.0), 100 mM NaCl, 0.05% (v/v) Tween-20) and incubating it for 2 minutes; (2) washing with 40 µl of buffer A; (3) flowing 20 µl of 0.5 mg/ml streptavidin (Invitrogen, catalog number S-888) dissolved in buffer A and incubating it for 2 minutes; (4) washing with 40 µl of buffer A; (5) equilibrating with 40 µl of buffer B. Twenty microliter of the purified biotin-labeled megastructure sample was then flown into the chamber and incubated for 2 minutes followed by washing with 40 µl of buffer B. For the purpose of drift correction, fiducial markers (40 nm gold nanoparticles; Sigma-Aldrich, catalog number 753637) were diluted to 1:10 in buffer B, flown into the chamber, and incubated for 10 minutes followed by washing with 40 µl buffer B. Finally, imaging buffer (buffer B containing 2 U/ml PCD (OYC Americas, sold as rPCO), 2.5 mM PCA (Sigma-Aldrich, catalog number 37580), 1 mM Trolox (Sigma-Aldrich, catalog number 238813)) was flown into the chamber. The imaging chamber was sealed with nail polish before imaging.

**Method 20 DNA-PAINT super-resolution imaging:** DNA-PAINT imaging was performed on a Nikon Ti Eclipse inverted microscope with a Perfect Focus System and a custom-build TIRF illuminator. Laser excitation with a 532-nm laser (MPB Communications Inc., 1 W, DPSS-system) was used for excitation with a 100 mW input at an effective power density of ~2 kW/cm2. The excitation laser was passed through a quarter-wave plate (Thorlabs, WPQ05M-532), placed at 45° to the polarization axis, and directed to the objective through an excitation filter (Chroma ZET532/10x) via a long-pass dichroic mirror (Chroma ZT532RDC_UF2). The laser beam was then expanded using a commercial variable beam expander (Edmund Optics, Broadband VIS 2X-8X) and a custom-built Galilean telescope followed by coupling into the microscope objective using a motorized mirror to generate a total internal reflection illumination. Emission light was spectrally filtered (Chroma ET542LP and Chroma ET550 LP), directed into a 4f adaptive optics system containing a deformable mirror (Imagine Optic, MicAO 3DSR) to correct optical aberration and optimize point spread functions (PSFs), and imaged on scientific complementary metal-oxide–semiconductor sCMOS camera (Andor Technologies, Zyla 4.2+) using rolling shutter readout at a bandwidth of 200 MHz at 16 bit and a 150 ms exposure time with 6.5-μm pixels, resulting in an effective pixel size of 65 nm. At each imaging session, 10,000 frames with an exposure time of 150 ms per frame were captured. DNA-PAINT image data were processed and rendered using Picasso[51]. The lateral positions of single-molecule localization events were determined using Picasso Localize followed by drift correction using imaged gold nanoparticles as fiducial markers.

### Supplementary Text 1: Design notes regarding the 12HB slat

The following section describes how the 12HB slats were derived from the 6HB design and explains the consequences that this choice had for folding and experimental usage.

##### 1.1 Derivation of 12HB slats from 6HB slats

The crossover pattern and staple set for the 12HB slat was derived from the design for the 6HB (Fig. S3). The reason for this was so that we would not have to purchase additional handle strands for the 12HB slats. Rather, we were able to apply the 2048-strand library from the 6HBs to also make the shorter 12HB slats. For the 12HB, a subset of staple strands in the middlemost segment of the 6HB was left out or modified to act as a hinge, so that the 6HB could fold back upon itself to make a half-length 12HB.

##### 1.2 Yield differences and the consequence on downstream usage of the slats

One shortcoming of the approach in Supplementary Text 1.1 was that the yield of 12HBs was much lower compared to the equivalent 6HB (~10% versus ~80%, see Fig. S4). We realized that the process of folding the 6HB back upon itself was inefficient because there was a propensity for the slats to dimerize or else make other higher molecular-weight byproducts. The gel in Fig. S4B indicates that roughly similar amounts of desired monomer and byproduct dimer formed even in the best folding condition. We hypothesized that the hinge did not allow complete freedom for the 6HBs to flex into the desired half-length 12HBs and that this process was perhaps energetically unfavorable. We expect that we could make formation of the half-length 12HB higher yielding by changing the scaffold routing to eliminate this flexing back in the folding pathway. However, that would require purchasing additional handle library strands to accommodate the different scaffold routing.

We note that high yield in the folding of the slats is critical for making downstream preparation of the megastructures easy and straightforward. The ~80% yield of the 6HBs gave us the ability to pool and concentrate the slats into a smaller volume using only PEG precipitation. By contrast, the low ~10% yield of the 12HBs required us to pool the slats, separate them from the undesired dimers by excision from an agarose gel, and then subsequently concentrate the 12HBs into a smaller volume with PEG precipitation. We generally observed ~80% recovery of a pool of raw 6HB slats versus about an order of magnitude decrease for the 12HB slats. We would suggest further optimization of the 12HB design with a different library of staple strands with the 32 possible handles should a user desire to make larger quantities of 12HB megastructures than as tested here. We also advise any user who conceives of alternative slat designs to optimize design and folding of the slat to make it easier to prepare at high yield for use in megastructures.

### Supplementary Text 2: Algorithm for selection of the handle sequences

We generated 6-nt, 7-nt, and 8-nt handle sequences so that each sequence of a given length had roughly similar binding energies and were as orthogonal from one another as reasonably possible. The following section describes the sequence-selection algorithm that we used to choose sequences for the 32 7-nts in the 2048 handle strand library, as well for the 256 6-nts, 7-nts, or 8-nts in the various 512-handle strand libraries. The former 2048 -strand library was used to create the diversity of finite and periodic megastructures, versus the latter 512-strand libraries which were used to make only v16/v8 staggered ribbons to characterize growth and nucleation of different strength binding sites (as detailed in Supplementary Text 4).

#### 2.1 Initial sequence generation

We created all possible 6-nts, 7-nts, and 8-nts using a Python script. We removed the reverse complements for each sequence and sequences with either extremely high or low GC content, and computed the pairwise mean free energy of the sequence with its exact complement in NUPACK 3.0[52] (using parameters DNA with 25 mM $Mg^{2+}$ and 2 mM $Na^{+}$). We considered this value as the binding energy of the handle. To make the sequences roughly isoenergetic, we removed sequences where the binding energy was some arbitrary number of standard deviations above or below the mean energy of the total set.

#### 2.2 Optimizing for sequence orthogonality

We computed the matrices of pairwise mean free energies between each handle with the other handles, reverse complements of the other handles, and reverse complement of the handle with the reverse complements of the other handles. As a measure of orthogonality of one handle versus another, we normalized the computed free energies in each matrix to the binding energy of the handle in question. We assumed this ratio for a given pair of sequences would lessen if the handles being compared were more orthogonal. As such, we set an arbitrary maximum for this ratio (i.e. a minimum threshold for orthogonality) that we allowed for binding-handle sequences in the library. We removed subsets of sequences that failed to meet this criteria most frequently with other sequences in the library and repeated this process until every handle satisfied the threshold.

##### 2.3 Discussion about the final sequences and pertinence to megastructures

The computed sequence energies of the handles as selected for this study are shown in Fig. S7. We note that only the 6-, 7-, and 8-nt handles were subjected to the selection criteria as above. The 9- and 10-nt handles used initially to test viability of different handle lengths (Fig. S6) were chosen randomly without the careful selection criteria as described above. By linear fitting of the computed energies, we found that each increment of one base pair decreased the mean free energy per handle by -1.26 kcal $mol^{-1}$, where the spontaneous nucleation was negligible with 7-nt handles versus prolific with 8-nt handles.

It is unclear to what extent the handle-design process in the prior sections is necessary for the successful assembly of megastructures. In our implementation, we hypothesized that orthogonality of the individual handles would be important so as to limit misbinding of the slats where base pairing is made between a handle other than its appropriate complementary handle. We further hypothesized that by making the relative binding energies of a given length handle similar to one another would make the boundaries clearer between where spontaneous nucleation becomes prevalent versus the threshold length of binding handle. However, we did not seek to accept or refute either of these hypotheses and rather used them as best practices to choose the handle sequences that we thought would be most successful to create megastructures. We also note that our analysis is subject to the shortcomings of the *in silico* tools which we used to measure binding energies. For instance, most of our decisions about whether to use a sequence or not were derived using the pairwise mean free energy in NUPACK 3.0[52] (with parameters of DNA with 25 mM $Mg^{2+}$ and 2 mM $Na^{+}$). We are unclear as to how well these calculations would align to experimental measurements. It might be the case that an experimentalist would be able to create megastructures with less care in the sequence selection of the binding handles, though this topic must be studied further to make any conclusions.

### Supplementary Text 3: Relationship between the number of unique slats in a design versus the relative rate of growth

We sought to determine the relationship for the growth of megastructures versus the concentration of the slats and number of unique slats in the design. Our intuition was that a higher concentration of each slat would increase the growth rate of a megastructure, as shown in the middlemost panel of Fig. 5D where there were 16 unique slats in the design. However, we note that the apparent rate of growth seemed noticeably slower as the total concentration of slats was increased past 1 μM (Fig. S22). We speculated that at these concentrations, liquid crystalline behavior of the slats may be occurring, thereby slowing down the rotational diffusion of slats, which may retard the rate of slat addition. This section will describe how this effect was even more dramatic for when more unique slats were added to the design. Using these observations, we describe general best practices for selecting the slat concentration and multi-pot slat addition steps to optimize growth.

##### 3.1 Growth slows when large numbers of unique slats are mixed simultaneously

We realized that the fastest growth is a balance between the concentration of each free slat versus the total concentration of all the slats in the reaction. In Fig. S21, we studied the growth of v16 staggered ribbons versus designs with increasing numbers of unique slats (i.e. decreasing sequence symmetry as shown in Fig. S20). We note that Hamming distances between the different slats are also an important parameter influencing growth, as described in Supplementary Text 6. All the designs with different numbers of unique slats generally had similar Hamming distances (as shown in Fig. S45G–J) so as to not make this a confounding variable.

As expected, the 2x symmetry design with 32 unique slats grew longer per unit of time when 15 nM per slat was added (i.e. 480 nM total slats) versus when 10 nM per slat was added (i.e. 320 nM total slats). Similarly, this also occurred with the 1x symmetry design with 64 unique slats where the 15 nM per slat (i.e. 960 nM total slats) condition grew faster compared to when 5 nM per slat (i.e. 320 nM total slats) was added. However, this relationship of faster growth with higher per slat concentrations was not maintained with the 0.5x symmetry design with 128 unique slats. After 12 hours of growth, there was no measurable assembly with 15 nM per slat (i.e. 1920 nM total slats) versus some measurable assembly when only 2.5 nM or 5 nM per slat (i.e. 320 or 640 nM total slats) was added. While after 38 hours of growth there was some measurable assembly for the 15 nM per slat (i.e. 1920 nM total slats) condition, the ribbons were of comparable length for the 2.5 nM per slat (i.e. 320 nM total slats) condition or even longer for when 5 nM per slat (i.e. 640 nM total slats) was added.

We concluded that although higher concentrations of each slat generally allow for faster growth, there are limits to which trend applies. There is some threshold between 1 and 2 μM where the total concentration of slats seems to cause an impediment to growth. This was further substantiated in Fig. S22, where the relative rate of growth of ribbons (using small 4x symmetry designs with 16 unique slats) was diminished as the total concentration of slats was increased towards 2 μM.

##### 3.2 Growth of finite megastructures with large numbers of unique slats can be accelerated by adding the slats in multiple steps

We predicted that growth of finite megastructures with large numbers of unique slats (e.g. >100) would be slow if all the slats were added simultaneously, because the total concentration of slats would exceed the 1 μM threshold causing growth impediment (see Fig. S21 and explanation in Supplementary Text 3.1). For instance, addition of all slats for the finite 1022-slat sheet at a low concentration of 2 nM per slat would make the total concentration surpass 2 μM. Hence, we considered an alternative approach where we sequentially added the slats in multiple stages to maintain a relatively high per-slat concentration, but without the total slat concentration ever exceeding 1 μM. We note that the initially added seed (and therefore the growing megastructure) is progressively diluted as more slat stages are added, though this was not problematic in this work. Future approaches could attempt to grow megastructures attached to a solid support, enabling circumvention of megastructure dilution.

We initially tested this approach for the elongated 320-slat plus symbol (see Fig. S14). In one approach, we incubated all the slats simultaneously during the initial preparation of the reaction, versus the other approach where half of the total slats (i.e. 160 slats) were added during the initial preparation and incubated for ~38 hours, at which point the remaining 160 slats were added and incubated again for a comparable amount of time. We note that the concentration per slat in the first approach was ~3 nM (i.e. 1000 nM total slats), versus ~5 nM per slat (~800 nM total slats at any given time) in the second approach. We observed that growth using the second approach with multiple slat additions resulted in completed megastructures at an earlier time versus when all the slats were added at once. One explanation of this faster growth with the second approach could be the ~66% higher concentration of a given slat during each stage. Users are advised to keep in mind the observations as explained in Fig. S21 and Supplementary Text 3.1, which suggest that increasing the total slat concentration towards 2 μM would likely impede the growth.

Given all of the above observations, we concluded that growth of the finite megastructures with the largest numbers of unique components should be conducted with the slats being added in multiple stages (see Fig. 2Biii–vi for the final structures, with the different stages shown in Fig. S9C–D and Fig. S10). The megastructures with the initial set of slats were incubated with 5 nM seed for three days, at which point an aliquot was taken and diluted 2.5x fold into the next series of slats for three more days, with the latter process being repeated five more times until all the slats were added (seed concentration estimated at 50 pM after final dilution). In our implementation, we maintained a concentration per slat for a given stage of ~4–5 nM and ensured that the total concentration of slats did not exceed 1 μM.

### Supplementary Text 4: Detailed characterization of binding handles

To fully characterize how the base-pairing strength of binding handles informs growth and spontaneous nucleation, we studied v16 and v8 ribbons using one of either 6-, 7-, or 8-nt handles at various temperatures. We used the ribbons as a model system for megastructure growth and nucleation because they were easy to identify, had lengths that could be readily measured in low-magnification TEM images, and could be applied to the TEM-grid substrate with high density so as to count many ribbons per image but where the lengths of single particles could still be resolved. All ribbon reactions for assessment of growth versus temperature had 20 nM per slat, 0.5 nM of seed, and 15 mM $Mg^{2+}$. Spontaneous nucleation was quantified by counting ribbons in a control reaction where no seed was added. The following section elaborates and discusses the results from the main text more deeply.

#### 4.1 How the number of binding sites (i.e. v16 or v8) versus temperature influences spontaneous nucleation

To understand how the number and strength of binding sites on the slats inform nucleation and growth, we also tested v8 6-, 7-, and 8-nt ribbons. With fewer binding sites, the v8 designs formed the longest ribbons at lower temperatures where higher spontaneous nucleation was observed. This is evident with the 7-nt design where growth of v16 could be maintained at 31ºC, 34ºC, 36ºC, and 38ºC with no observable spontaneous nucleation, versus the v8 design where this could only be maintained at 30.5ºC and 32.5ºC (Fig. S38, Fig. S39). Similarly, low amounts of spontaneous nucleation (i.e. one order of magnitude greater than the 0.3 pM detection limit) could be maintained at 43.7ºC, 47.3ºC, and 50.2ºC with the v16 8-nt design, versus only over a narrower range (40ºC and 42.5ºC) with the v8 8-nt design (Fig. S38, Fig. S39). We were not able to observe these differences using the 6-nt binding sites because no spontaneous nucleation was observed at any of the temperatures tested (Fig. S38, Fig. S39). We conclude that the v16 design where more binding sites are used allows growth using a broader window of temperatures where spontaneous nucleation is not observed, which is consistent with prior findings with ssDNA slats[33].

#### 4.2 How the base-pairing strength of the binding site handles versus temperature influences growth rate and spontaneous nucleation

We grew v16 6-, 7-, and 8-nt ribbons with and without the seed for ~16 hours at various isothermal temperatures to determine if fast growth and controlled nucleation could be attained simultaneously (Fig. S38). Additionally, we determined the temperature where grown ribbons melted apart (Fig. S37). We sought to find the breadth of growth temperatures that were below the melting point where no measurable spontaneous nucleation occurred, because the span of this window represents the utility of a particular length binding site for controlling nucleation of the slats. We can see that the v16 8-nt design is subject to spontaneous nucleation at all temperatures tested (see rightmost Fig. S38A and top panel of Fig. S39). However, no spontaneous nucleation for the v16 6- and 7-nt designs was observed using a large window of temperatures up to ~11ºC below the melting temperatures (see left two panels in Fig. S38A and top section of Fig. S39). With the 7-nt design, only minimal spontaneous nucleation that was about an order of magnitude greater than the 0.3 pM detection limit was observed at the lowest 28ºC condition tested (middle panel of Fig. S38A). Taken together, we can see that either the 6- or 7-nt binding sites provide leeway with choosing fast assembly conditions that are far below the melting temperature where nucleation is strictly seed dependent. In practice, this window gives the experimentalist freedom to set up origami-megastructure growth reactions at room temperature without the slats becoming aggregated into undesirable byproducts, to be able to accommodate variations in the ion and slat concentrations, and be able to use a breadth of growth temperatures where fast growth and seeded nucleation predominate without significant formation of unseeded byproducts.

In the growth-temperature trials, we also sought to find the "optimal" growth-temperature range where the fastest growth occurred in the window of nucleation control. Temperatures in this window serve as a guideline for reaction conditions that an experimentalist might select for fast, seed-dependent growth. The results are summarized as dark blue bars in the top portion of Fig. S39. We find that the optimal growth temperature ranges were 22.5–25ºC, 31–34ºC, and 43.7–47.3ºC for the v16 6-, 7-, and 8-nt ribbons, respectively. Thus, each increment of one base pair in the v16 ribbon binding sites increases the optimal temperature range by ~10ºC. We note that defining exact bounds for an "optimal" temperature range is arbitrary, which would also depend on parameters such as slat and cation concentration.

### Supplementary Text 5: Kinetics of ribbon assembly

We determined the kinetics of slat addition by measuring the length of ribbons at different timepoints (see Fig. S40), then converting the length measurements into an estimated number of slat additions, and deducing an assumed pseudo-first order rate constant ($k_{on}$) from that estimated number of slat additions. The following section explains how these measurements were determined and how the base-pairing strength of the binding site influences these rates.

#### 5.1 Calculation of the number of slats added to the ribbon

We assume that each 6HB slat on the v16 staggered ribbons extends the ribbon roughly by ~10.1 nm, as explained in Fig. S19C. Hence, the number of slats in a ribbon is given by equation 5.1.

$$number\ of\ slats\ =\ \frac{(nm\ length)}{10.1\ nm\ slat^{-1}} \qquad (5.1)$$

#### 5.2 Approximation of growth using pseudo-first order kinetics

In ribbon reactions, the slats were in huge excess with respect to the seed. In a typical reaction such as where growth was measured versus time in Fig. S40, we used a 40-fold excess of each slat with respect to the seed (i.e. ~20 nM of each slat versus 0.5 nM seed). The ribbon designs in Fig. S40 used 4x sequence symmetry (see Fig. S20 for explanation of symmetry) and were thus composed of 16 unique periodic slats. If we imagine that every one of the seeds added triggers growth of a ribbon, then complete depletion of the slats would net ribbons that are each ~6.5 µm long. We may thus assume that the concentration of slats did not change appreciably over the duration of growth, especially during early time points where the mean length is well below this ~6.5 µm threshold. With this, we proceeded to compute second-order $k_{on}$ constants from the length assuming pseudo-first order kinetics, with equation 5.2. The length was divided by two to account for how the ribbons grow bidirectionally from the seed.

$$k_{on} = \frac{(nm\ length)}{2}\ \frac{1\ slat}{10.1\ nm}\ \frac{1}{[concentration\ per\ slat\ in\ M]\ (time)} \qquad (5.2)$$

It should also be noted that lack of consideration for slat depletion effects at the longest time points in Fig. S40, where mean lengths of ribbons approached or exceeded the ~6.5 µm threshold, would contribute to the gradual decline of the apparent second-order $k_{on}$ values versus time.

#### 5.3 How the base-pairing strength of the binding-site handles influences growth rate

The rate of ribbon growth was increasingly faster as the strength of the binding sites were increased from 6, to 7, and to 8 nt. This is shown in Fig. S40 where the length of v16 ribbons was compared at different time intervals using a growth temperature where little to no spontaneous nucleation was observed. The mean lengths for the ribbons after 1 hour of growth were ~1, ~1.3, and ~3.1 μm for the v16 6-, 7-, and 8-nt designs, respectively, as shown in Fig. S40Ai, Bi, and Ci. After 16 hours of growth, these increased to ~4.2, ~4.9, and ~8.1 μm for the v16 6-, 7-, and 8-nt designs, respectively. This suggests surprisingly fast kinetics for addition of the 6HB slats—converting the mean length of the ribbons (as explained in Supplementary Text 5.1) after 1 hour of growth gives the observed second-order rate constants for growth of ~$0.66*10^6$, ~$0.86*10^6$, and ~$2.13*10^6$ $M^{-1}s^{-1}$ for the 6-, 7-, and 8-nt ribbons respectfully, as shown in Fig. S40Aii, Bii, and Cii. After 16 hours of growth, the observed second-order rate constants for growth were ~$0.18*10^6$, ~$0.21*10^6$, and ~$0.35*10^6$ $M^{-1}s^{-1}$ for the 6-, 7-, and 8-nt ribbons. We attribute this decrease in the apparent rate constants over time to accumulation of errors on the slats on a growing ribbon end that might slow further growth, and lack of consideration of depletion of slats in the calculation. As with any folded DNA origami, there is only some probability (e.g. perhaps ~80–90% per handle) that a 3′ handle sequence is available on a 6HB or 12HB slat. A multitude of errors on a ribbon end might temporarily stall or entirely stop the addition of more slats.

### Supplementary Text 6: Sequence assignment to megastructures from the 2048-handle sequence library, optimization of the Hamming distances of slats within megastructure designs

The megastructure designs were implemented from the 2048-sequence library by randomly assigning one of the 32 possible handle sequences to each of the intersections in the interface between the perpendicular layers of slats, as explained in Method 2. In the initial explorations with periodic v16 7-nt ribbons as shown in Fig. 3Ai, we discovered that particular random assignments of sequences sometimes resulted in drastic growth differences. We realized that certain designs that grew less prolifically had particular pairs of perpendicular slats where a multitude of complementary handles were matching and aligned to one another. We hypothesized this unintentional alignment of complementary handles caused the formation of kinetic traps with pairs of slats undesirably bound to one another (see Fig. S41). We designated the maximum number of matches between some pair of slats in a megastructure design as having a kinetic trap of *k* strength. For example, a design with a maximum kinetic trap *k6* has one or more pairs of slats where the two slats may be aligned with six complementary binding handles engaged together. We quantified this design property of the random sequence assignments for megastructures by measuring Hamming distances between each of the slats in one layer with each of the slats in the other perpendicular layer. This section explains the initial findings with periodic ribbons that suggest the need to maximize Hamming distances (also see Fig. S42), results showing a mechanism for how single kinetic traps in a pair of slats influences ribbon growth (also see Fig. S43), and the Hamming distances of the megastructure designs as implemented in this paper (also see Fig. S44–Fig. S45).

#### 6.1 Increased pairwise complementarity between slats (i.e. minimized Hamming distances) causes drastic slowdown of 1D ribbon growth

We generated random permutations from the strand library of v16 ribbons and selected designs where the minimum Hamming was marginally increased (see Fig. S42A). The designs tested had distributions of slat pairs where there were maximally 6 and 8 complementary binding sites aligning between them (i.e. *k6* and *k8* strength kinetic traps, respectively). The length of ribbons as measured by TEM after overnight growth of the *k6* and *k8* designs at various temperatures is plotted in Fig. S42B. There was no appreciable growth of the *k8* ribbons at any of the temperatures tested, versus the *k6* ribbons which grew prolifically at all the temperatures. Examples of the differences between growth versus no growth for the *k6* and *k8* designs are shown in low-magnification TEM images in Fig. S42C–Di. Closer inspection revealed infrequent short, stubby *k8* ribbons in select higher-magnification images, but their relative number was too few to quantitatively compare to the *k6* counterpart (see Fig. S42Dii). To rationalize these extreme growth differences, we considered the Hamming distances in the *k8* design more closely. We realized that of the total eight top and eight bottom slats in the design, five of the slats from either layer had eight complementary binding sites that could align with one another as a *k8* strength kinetic trap. We hypothesized that these unintentional "strong" *k8* interactions between over half of the total slats among themselves prevented the slats from readily being recruited to the ribbons. Regardless of the cause of the varied growth, these results suggested that avoiding low Hamming distances between the slats is important to achieve the best possible growth of a megastructure.

#### 6.2 Possible mechanism for how complementarity between the slats causes growth slowdown

To test our hypothesis that alignment of complementary handles between slat pairs causes the formation of kinetic traps that prevent the slats from freely being added to megastructures, we tested growth of several ribbon designs where we incremented the strength of a kinetic trap between a single pair of slats (see Fig. S43A). The mean length of the ribbons at different temperatures versus the single *k8*, *k10*, and *k12* traps are shown in Fig. S43B–C, versus a *k6* control. The ribbons were progressively shorter at all temperatures versus the *k6* control as the strength of a single kinetic trap *k* was incremented from *k8* to *k10* to *k12*. We also found periodic defects in the *k10* and *k12* ribbons with every eighth slat in a particular layer that was frequently missing in close-up images of ribbon segments (see Fig. S43Diii–iv). We note that the designs tested here use 4x sequence symmetry with eight repeating top-slats and eight repeating bottom-slats, such that a particular missing slat in each layer would appear at such intervals (for details regarding symmetry see Fig. S20). The observation of both slower overall growth and missing single slats with the strongest kinetic traps support the mechanism that low Hamming distances cause kinetic trapping of slats that impede their addition to a megastructure. Different segments of other ribbons with the *k10* and *k12* traps were observed to have the full set of periodic slats without the missing slat, suggesting that the trapped slat pairs could eventually be added. We suspect ribbon growth where a slat pair is strongly trapped becomes temporarily stalled because it is often missing from the growing ribbon edge. The missing trapped slats are added at a slower rate compared to other free slats and are perhaps skipped and added to the completed segments of ribbon later in time. Henceforth, we concluded that the Hamming distances between slats should be maximized to the greatest extent possible to limit kinetic trapping to best allow free slats to be added to megastructures.

#### 6.3 Discussion of the Hamming distances used in the finite and periodic megastructure designs

For the optimized finite and periodic designs, the multiplicity for each possible number of matching, complementary handles for the slats from one layer with respect to the other layer of slats are shown in Fig. S44–Fig. S45. The largest finite designs in Fig. S44D–F maximally had *k8* kinetic traps, with the smaller finite designs having weaker traps as the number of slats in the design was lessened. In general, the maximal traps were larger for designs with more unique slats because there are more possible interactions between the various slats. For instance, the 64-slat square had maximally five undesired complementary binding sites between slats (i.e. a *k5*-strength kinetic trap) versus the 1022-slat sheet which had maximally eight undesired complementary binding sites (i.e. a *k8*-strength kinetic trap), as shown in Fig. S44A versus Fig. S44F.

We note that the results in Fig. S42 showed that ribbon designs with *k8* traps either struggled or were incapable of appreciably growing into ribbons, which suggest that the designs in Fig. S44D–F should not be viable for growth. However, we hypothesized that the multi-step addition of slats for the finite designs with the largest numbers of slats (as per Method 10) would have resulted in particular slats being at a different concentrations compared to other diluted slats from prior stages. That is, the growth hindrance resulting from particular kinetic traps could have been lessened because of the differing stoichiometric amounts of particular slats. We also hypothesize that a few isolated "strong" *k8* kinetic traps in the largest finite designs would be less deleterious to growth compared to a periodic design with a similar-strength trap. With the finite design of unique slats, a single pair of trapped slats would only be skipped once (and perhaps slowly added at some later time), versus periodic designs where skipping of the trapped slat would be encountered repeatedly.

The periodic ribbon designs used to study the nucleation and growth behavior of the origami megastructures were generated from permutations from 256 different 6-, 7-, or 8-nt handles using a 512-strand library, and had maximally *k4*-strength kinetic traps (see Fig. S45A–B). Other periodic designs that were generated from the 32 different 7-nt handles using the 2048-strand library all had maximally *k6*-strength kinetic traps (see Fig. S45C–J). We attribute these differences between the maximum-strength kinetic traps to the differences in the number of different handle sequences with each of the different strand libraries.

### Supplementary Figures Fig. S1–Fig. S45

#### Fig. S1–Fig. S8: Slat and seed design and folding, qualitative energetics of crisscross assembly, selection of initial binding-site strength, energies of various binding-handle sets, and need for T-linkers

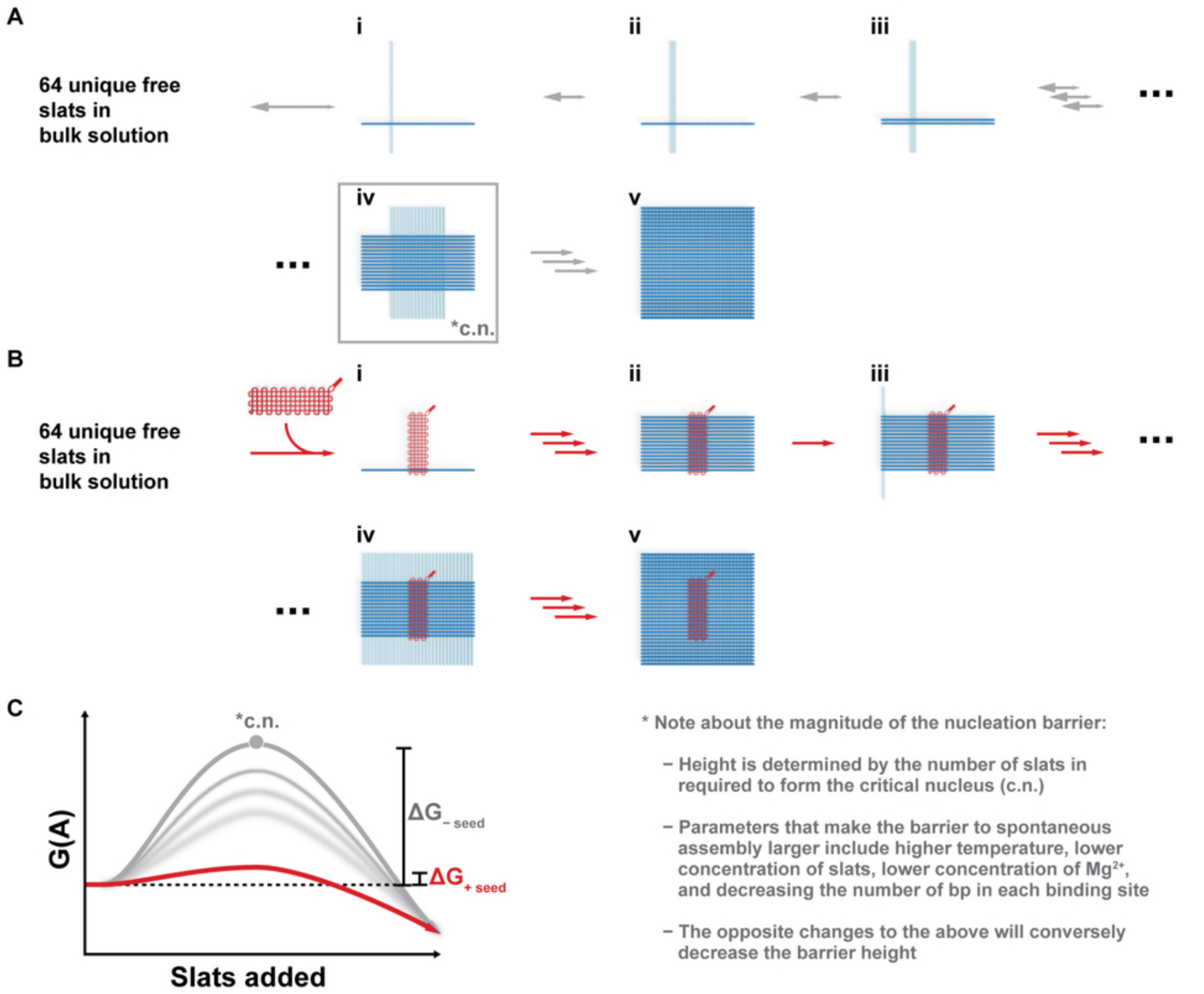

**Figure S1** Qualitative energy landscape of the nucleation barrier in crisscross assembly for the 64 6HB slat finite megastructure. This figure elaborates on the energetic pathways introduced in Fig. 1C. **A** shows unseeded assembly of the slats when no seed is added to the reaction. Single pairwise interactions between slats are weak and transient, with the reverse reaction of slat dissociation favored over forward assembly, as shown in *Ai–iii*. Stable addition of slats does not occur until the formation of the metastable critical nucleus (c.n.) in *Aiv*. Once the critical nucleus forms, the remaining slats may readily bind to the critical nucleus to complete the megastructure as in *Av*. **B** shows assembly when the seed is added to the reaction. Each of 16 cells in the seed have binding sites that strongly engage one 6HB slat, as shown in *Bi–ii*. Steps *Biii–v* with the remaining slats (e.g. 48 slats as shown here) bypasses the critical-nucleation pathway, because the parallel 16 slats in *Bii* stably localize columns of weak binding in close proximity. In the experimental assembly, we incubate the reactions initially for 4 hours at a higher temperature that does not let the reaction proceed past *Bii*. Next, the reaction is incubated at a lower temperature that favors binding of the remaining slats with weak binding sites, as in *Biii–v*. These two possible assembly pathways are qualitatively plotted, as shown in **C**. The various experimental and design parameters that influence the magnitude of the energy barrier to spontaneous assembly are described in the notes on the right.

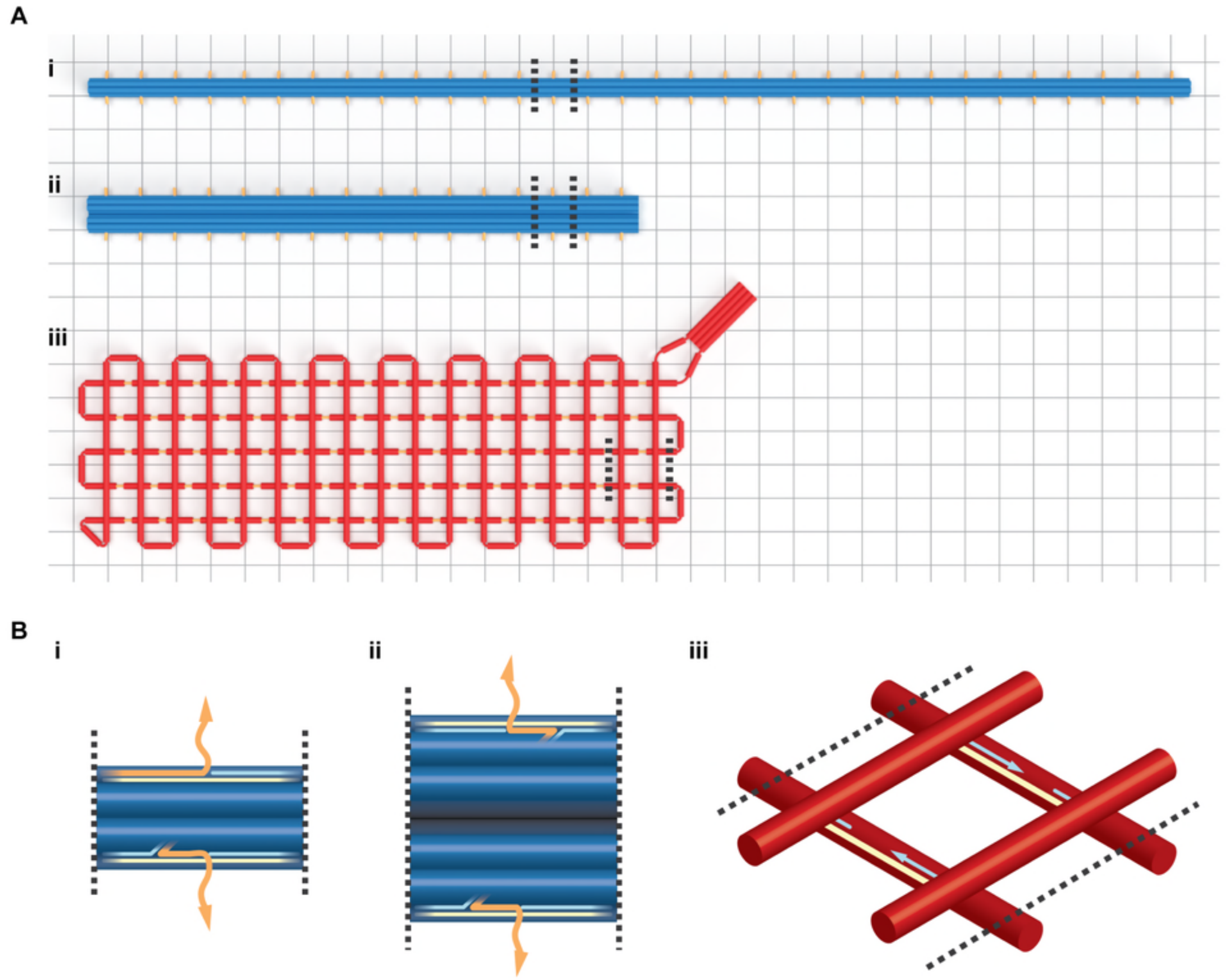

**Figure S2** Large renderings of designs for the 6HB slat, 12HB slat, and gridiron seed. Panel **A** shows the overall structure of each origami component, with the background grid representing 42 bp. The orange ssDNA sites along the top and bottom of each slat are the nodes which may be arbitrarily addressed with cargos as modifications to a staple will allow. In particular, we used these nodes for weak handles to bind the slats to other slats, for strong 10-nt handles to bind the slats to the sockets on the seed, for strong 16-nt handles to bind DNA nanocubes (see Fig. S28), or as 3′ biotin sites to bind DNA megastructures to substrates for DNA-PAINT imaging. The orange notches in the seed in *Aiii* are exposed 10-nt regions of scaffold which serve as “sockets” to strongly bind slats via complementary “plug” handles. The seed has 16 columns as oriented, where five horizontal helices (i.e. five sockets) in each column may cooperatively engage a single slat with five strong 10-nt handles. The black lines indicate the region shown more closely in panel **B**, where the orange line represents the staple strands for addressable nodes on a slat, light yellow lines represent the scaffold strand, and cyan lines as other core staple stands.

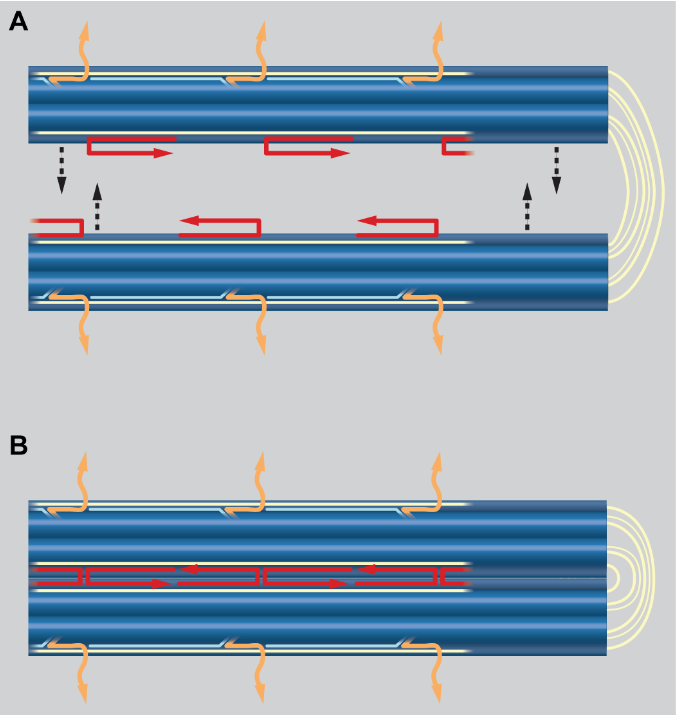

**Figure S3** Strategy in which the 6HB slat folds back upon itself to make a half-length 12HB slat. Extreme right in panel **A** shows a region of light yellow scaffold that was left as ssDNA without any staple strands. This region acts as a hinge to allow the 6HB to fold back into a 12HB, using the red staple strands to bridge opposite ends of the top helix together. Panel **B** shows a segment of completed 12HB that was properly folded back. This scaffold routing allows us to use the same base staple sequences (i.e. the orange staple strands) for the 12HB as is used with the 6HB, so that the same 2048 7-nt strand library can be applied to either slat. The consequence of this design on the final slat yield is explained in Supplementary Text 1 and shown in Fig. S4.

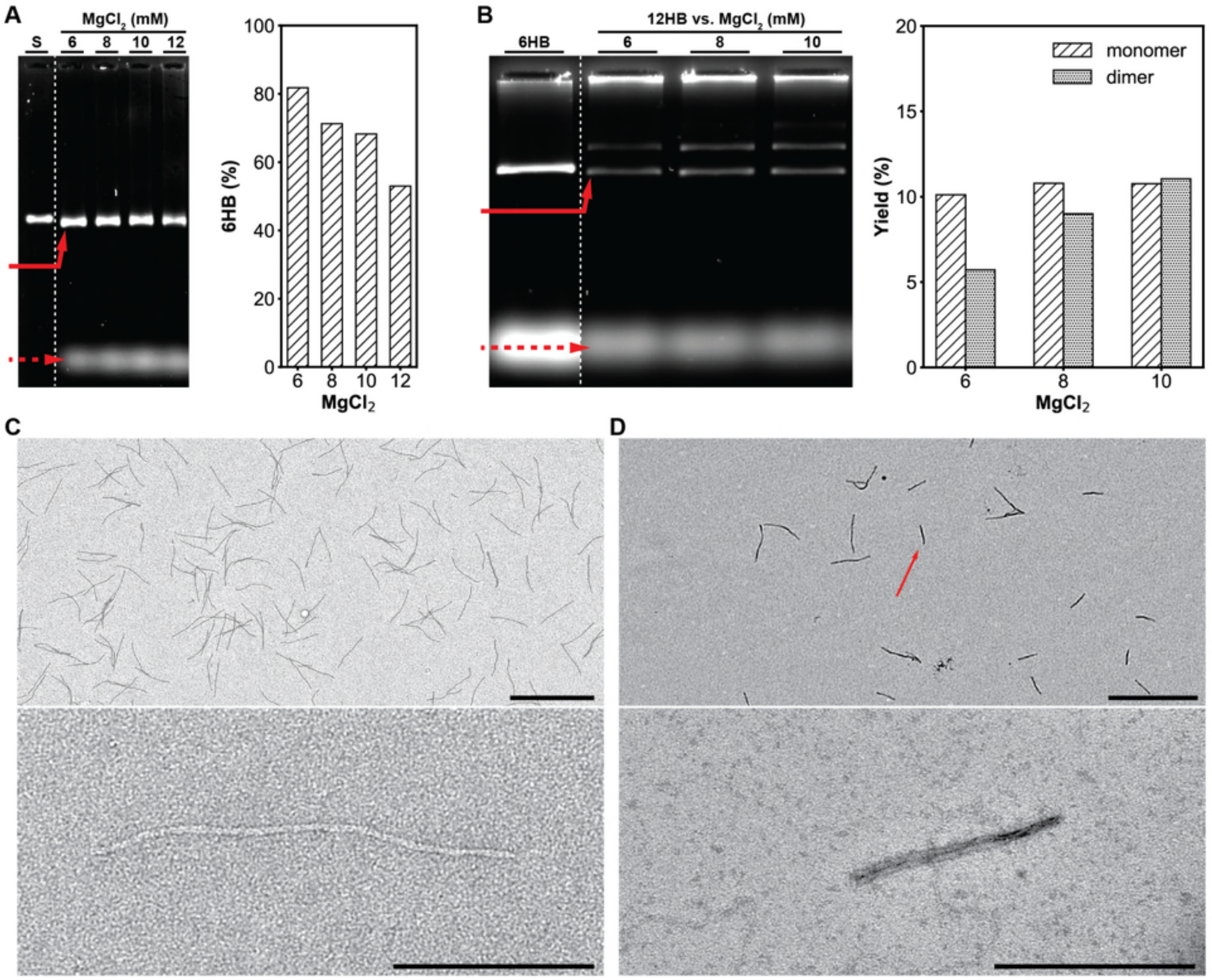

**Figure S4** Folding of the DNA-origami 6HB and 12HB slats. Leftmost **A** and **B** show agarose gels to optimize $MgCl_2$ for folding of the 6HB and 12HB slats respectively, with the solid red arrow pointing to the slats and the dashed arrow pointing to excess staple strands. Control lanes (S) is the scaffold only for the 6HB in *A*, versus a 6HB reference for the 12HB in *B*. The folding yields were determined in the rightmost of *A* and *B* using densitometry of the agarose gel, where the intensity of the desired slat band was compared to the overall intensity of all the elements that migrated slower than the staple band. Roughly 80% yield for 6HB slats was obtained in the 6 mM $MgCl_2$ condition, versus 10% yield for monomeric 12HB slats in the 8 mM $MgCl_2$ condition. See Supplementary Text 1 for further explanation of the differences between these slats. TEM images of the raw, folded 6HB and 12HB slats are shown in **C** and **D**, respectively. Only the monomeric 6HB is observed, versus the 12HB where there is a mixture of the desired 12HB monomer (as shown with the red arrow) and an undesired double length 12HB dimer. Scale bars are 1 μm and 200 nm in the top and lower images of *C* and *D*, respectively.

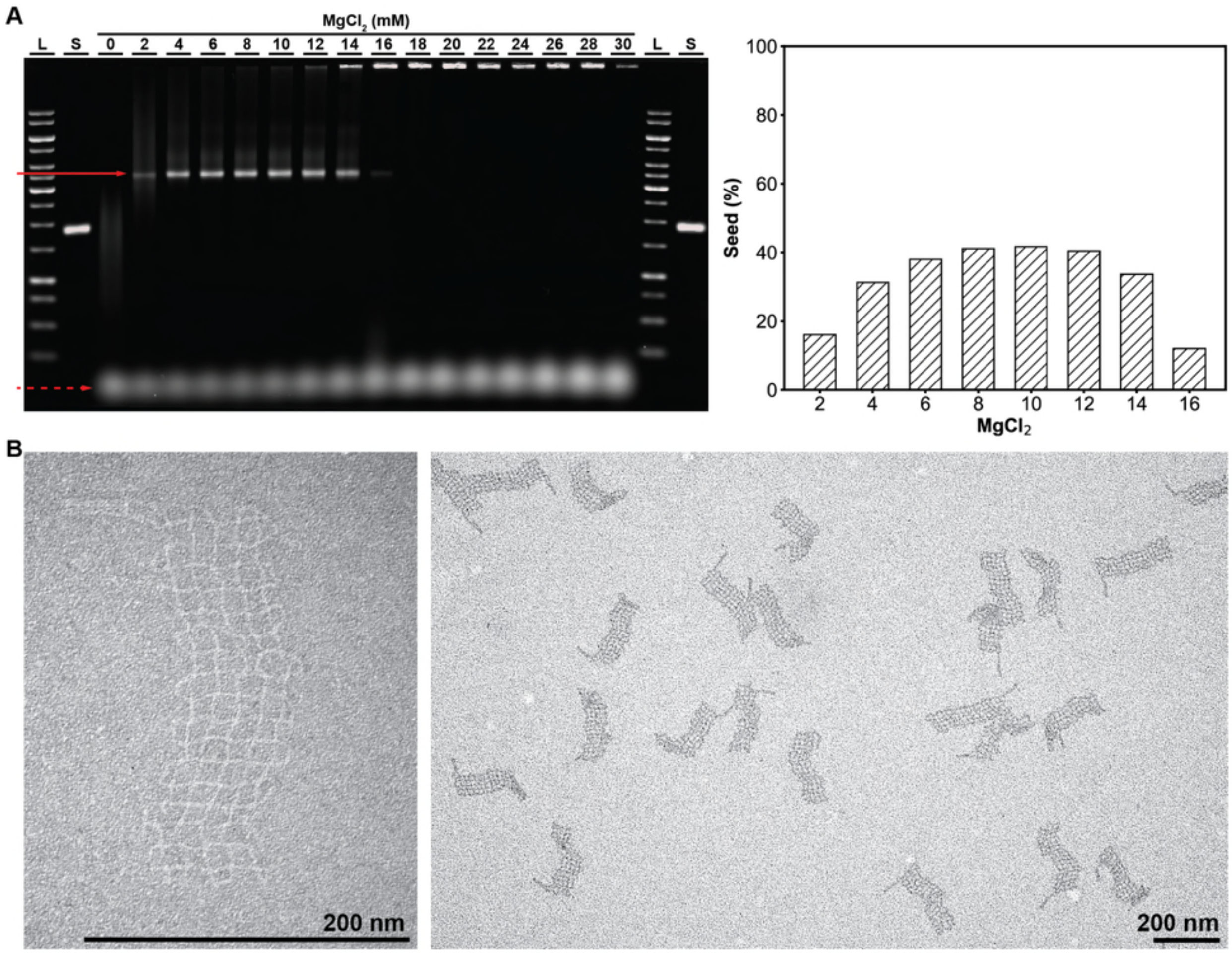

**Figure S5** Folding of the gridiron DNA-origami seed. Leftmost **A** shows an agarose gel to optimize $MgCl_2$ for folding of the seed, with the solid red arrow pointing to the seed band and the dashed arrow pointing to the staple strands. Lanes (L) indicate ladder and (S) indicate scaffold only. The folding yields were determined in rightmost **A** using densitometry of the agarose gel, where the intensity of the seed band was compared to the overall intensity of the elements that migrated slower than the staple band. Roughly 40% of seed monomer was obtained for the 8 mM and 10 mM $MgCl_2$ conditions, which we deemed to be the folding conditions with the highest yield. **B** shows TEM images of negatively (left) and positively (right) uranyl-formate-stained gridiron seeds. We note that only a single design seed with the same scaffold register was used for all the megastructures in this work, such that the socket sequences needed for each structure were universal. Scale bars are 200 nm.

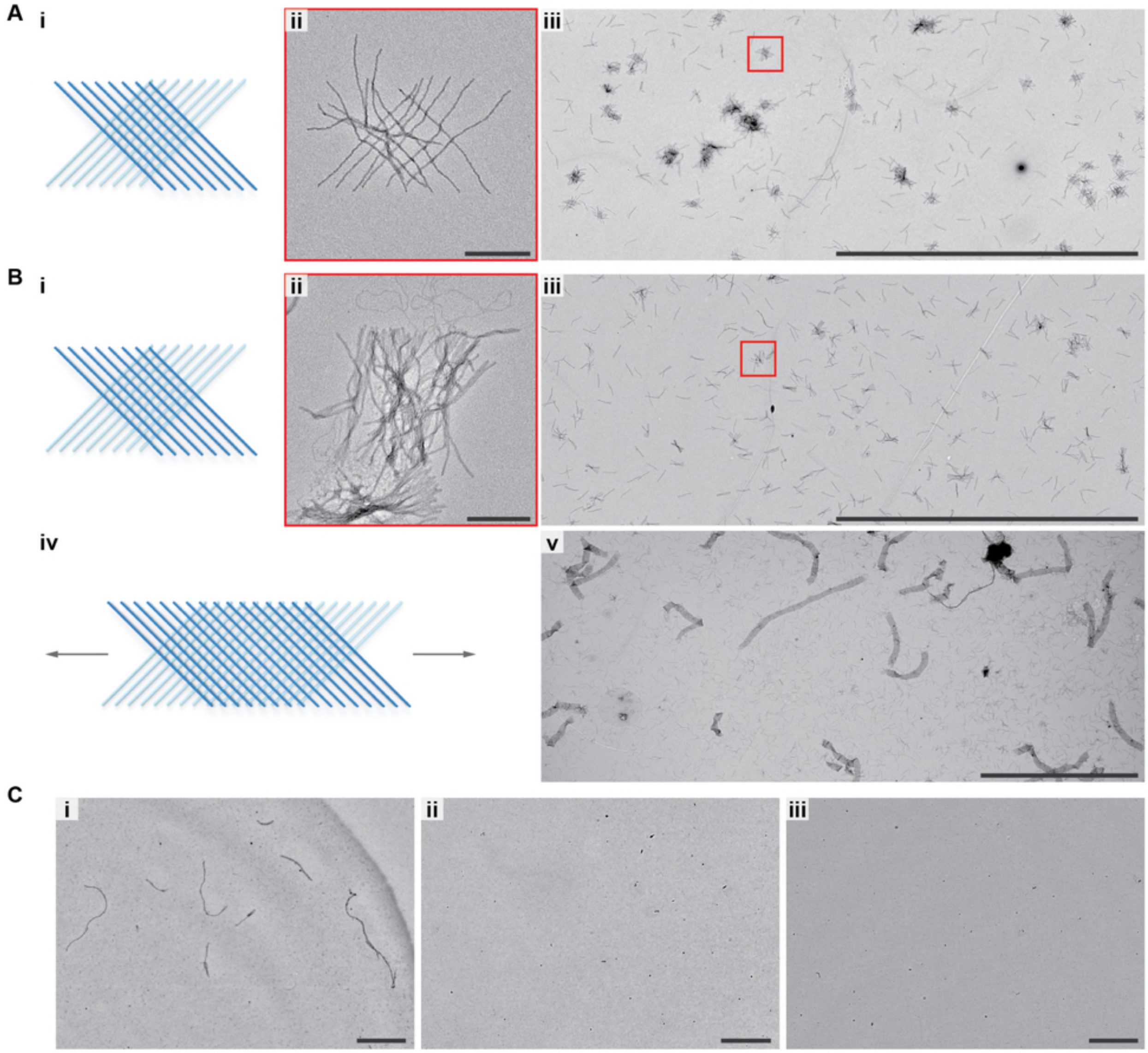

**Figure S6** Qualitative assessment of unseeded spontaneous nucleation of 10-, 9-, 8-, 7-, and 6-nt binding sites for crisscross assembly. Binding-site designs where nucleation of assemblies was observed using typical conditions that might be encountered in routine experimental procedures were determined not suitable for seeded nucleation control. **A** shows that 10-nt binding sites had prolific error-prone assembly of the finite 16-slat assembly (*Ai*), as shown by TEM (*Aii–iii*). Similarly, spontaneous assembly of something assumed to be some portion of the 16-slat test structure was also observed with 9-nt binding sites, in **Bi–iii**. Structures similar to the closeup from panel *ii* are highlighted in red in panel *iii*. In **Biv–v**, the propensity of spontaneous nucleation with the 9-nt design was further shown using a v8 ribbon design. As such, we further tested v8 ribbons using 8-, 7-, and 6-nt binding sites, in **Ci–iii** respectively. Prolific ribbon growth occurred using the 8-nt design. However, it was not observed for either the 7- or 6-nt designs such that we concluded either these strength binding sites would be suitable for seeded nucleation control of the origami slats. Assembly of the 16-slat test structure was for 16 hours at 25ºC in 10 mM $MgCl_2$ using 2 nM of each slat, and assembly of the v8 ribbons was for 16 hours at 32.5ºC in 15 mM $MgCl_2$ using either ~6.6 nM or 20 nM (in *Biv–v* and *C*, respectively) of each slat. Scale bars are 200 nm in *Aii* and *Bii*, and 10 µm in all other images.

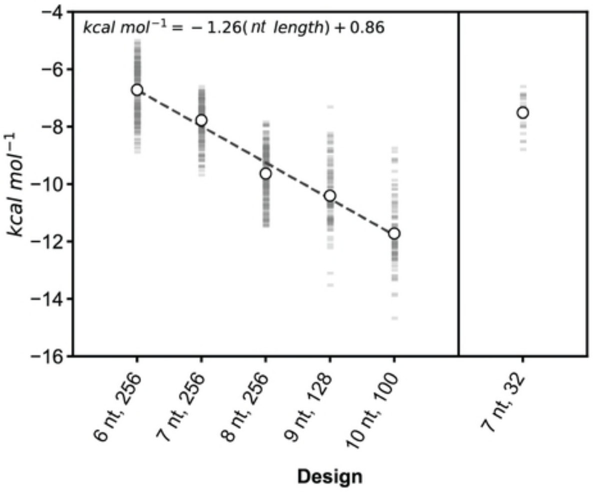

**Figure S7** Calculated mean free energy of the sequence handles versus their base-pair length, as determined in NUPACK 3.0[52]. Each faint gray bar plotted is the energy of a single handle, with the circular data points indicating the mean energy of the library tested. The 6-, 7-, and 8-nt sequences were selected as described in Supplementary Text 2. The 256-sequence set was used to characterize growth and nucleation of origami megastructures as described in Supplementary Text 4, and the rightward 7-nt 32-sequence set was used for the 2048-strand library to create a diversity of megastructures. The strongest 9-nt 128-sequence and 10-nt 100-sequence sets were used only for initial testing of sequence-handle length (see Fig. S6). For the leftward sequences, a linear fit suggests that each single-nt increment of the handle increases the binding energy by -1.26 kcal $mol^{-1}$.

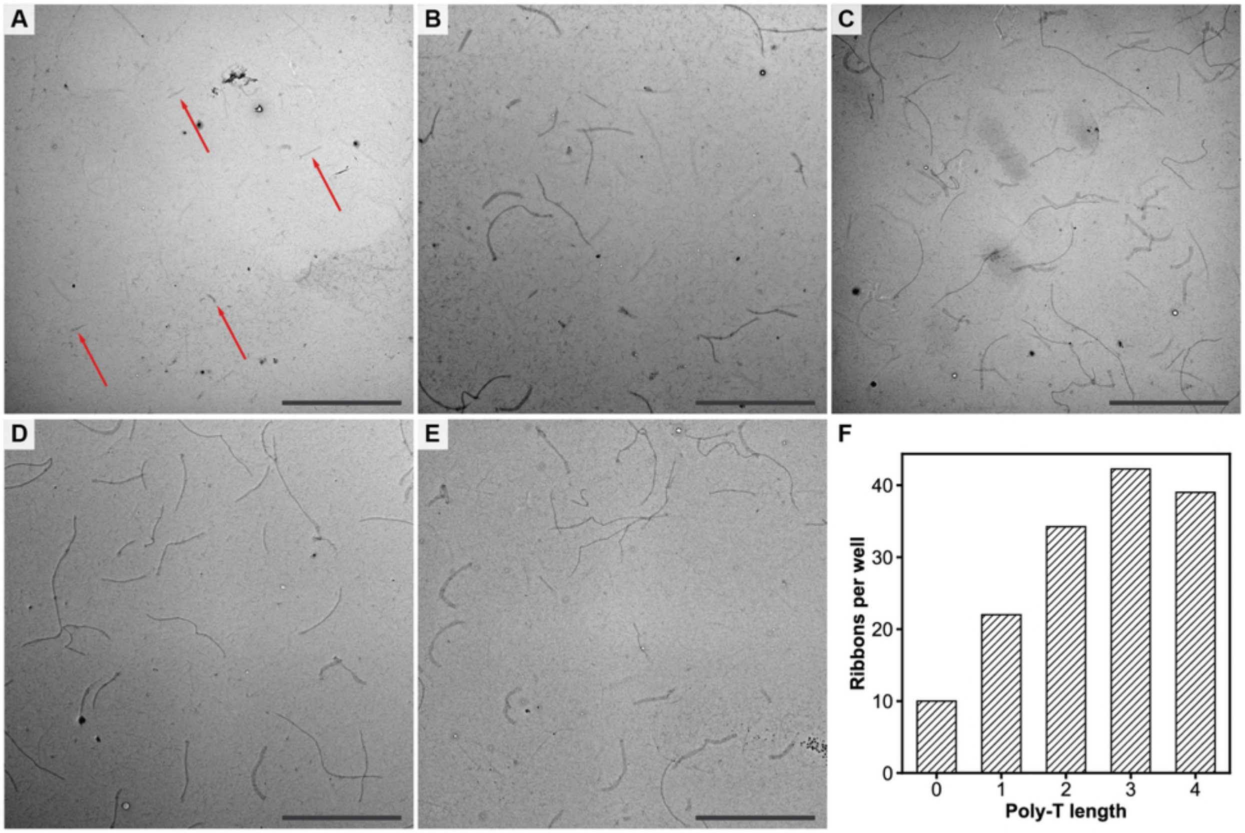

**Figure S8** Addition of a poly-T-linker before the binding sites is necessary for allowing assembly of megastructures from DNA-origami slats. The relative number of seeded v8 8-nt ribbons as counted on TEM grids was increased if a 1T, 2T, 3T, or 4T linker was used to separate the handle from the core staple sequence versus if no linker was used. Representative TEM images for the 0–4T linkers are in panels **A–E** respectively, with the average number of ribbons counted per defined imaging area plotted in **F**. The red arrows in panel A point to the stubby ribbons that were much shorter compared to the longer ribbons that formed when a linker was used. This observation may be due to relaxation of electrostatic repulsion of the slats that would otherwise impede megastructure assembly. The relative difference in the number of ribbons counted was similar for 2T, 3T, and 4T linkers. We therefore decided to use 2T linkers for the 8-, 7-, and 6-nt handle designs, as shown in Fig. 1Di. We further note that the 9- and 10-nt binding sites as tested in Fig. S6A–B did not use any T-linker, though were still able to nucleate prolifically into unseeded crisscross test structures. We conclude that T-linkers are necessary to allow the assembly of DNA-origami megastructures using weak binding sites that are less than or equal to 8 nt. This observation does not seem to hold true for when stronger handles are used, though we did not study this phenomenon with such designs. Scale bars are 10 µm. N = 3 images were counted for the 0T design, and N = 4 images were counted for all the other designs.

**Fig. S9–Fig. S15: Models of finite megastructures, preparation time for each design, and additional TEM results of finite megastructures**

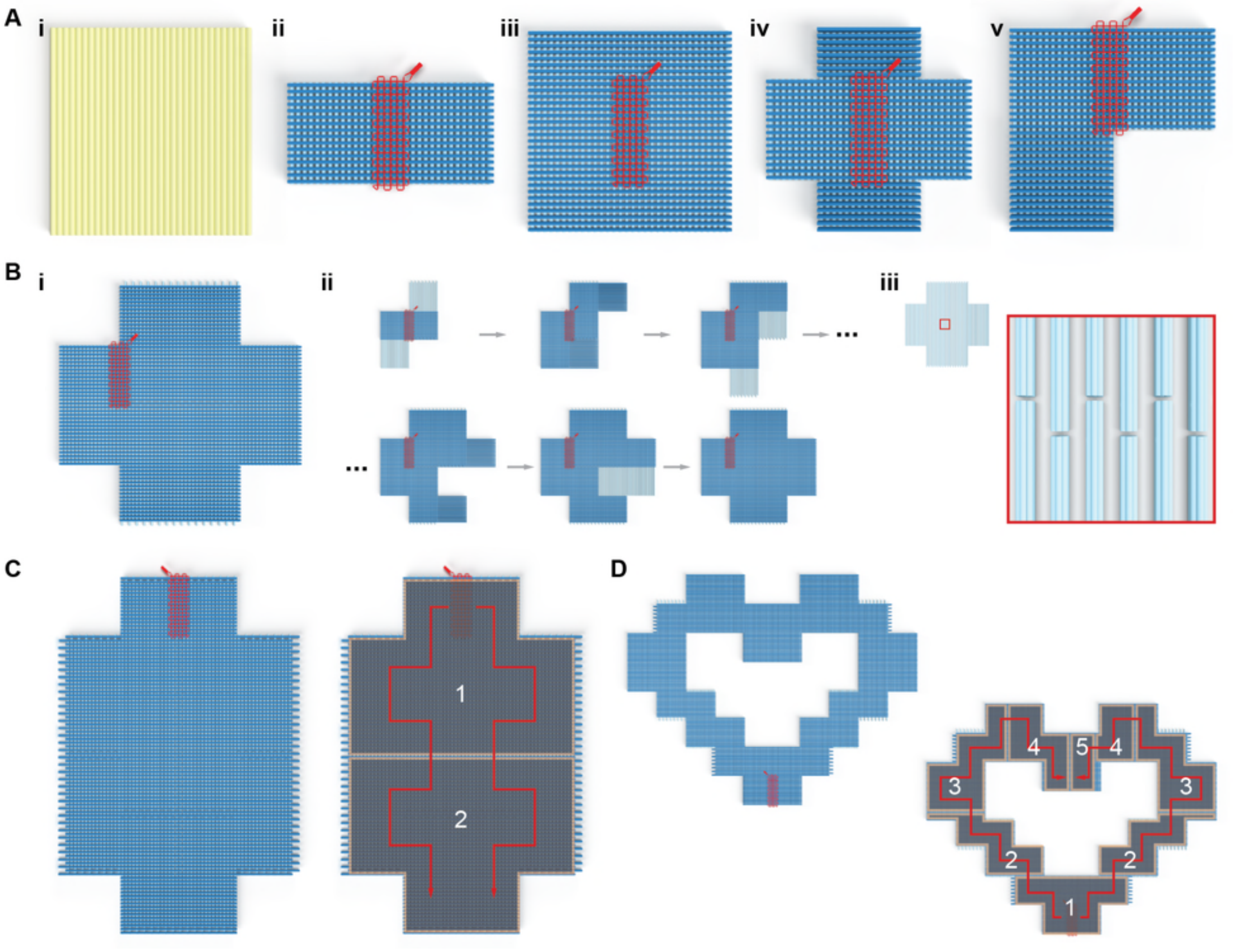

**Figure S9** Enlarged design renders of the various finite origamis and megastructures from Fig. 1C. **Ai** is a single-layer flat-sheet DNA origami that is folded from the same scaffold that is used to make the slats, **Aii** is a 48-slat rectangle composed of 16 6HBs and 32 12HBs, **Aiii** is a 64-slat square composed of entirely 6HBs, **Aiv** is a 64-slat plus symbol composed of 32 6HBs and 32 12HBs, and **Av** is a 64-slat L symbol composed of 32 6HBs and 32 12HBs. Panel **Bi** is a larger 191-slat plus symbol composed entirely of 6HBs, with various steps in its raster-fill growth in **Bii**, and **Biii** showing how the bottom-most vertical cyan slats were staggered to seal the middlemost horizontal seam. Such seams exist between the abutting edges of each ribbon-like section of the megastructure. The slats in one or both of the two layers can be staggered, such as done with the finite megastructures in *C–D* and Fig. S10, and the ribbon in Fig. 3iii. **C** leftward is the 320-slat elongated plus symbol. **D** leftward is the 568-slat heart. Rightward panels in C and D show the raster-fill growth as a red line, with the darkened overlays showing each of the distinct stages where up to 200 slats were added. The megastructures were incubated for about three days of isothermal growth before additional slats for the next stage were added. Images are not drawn to scale.

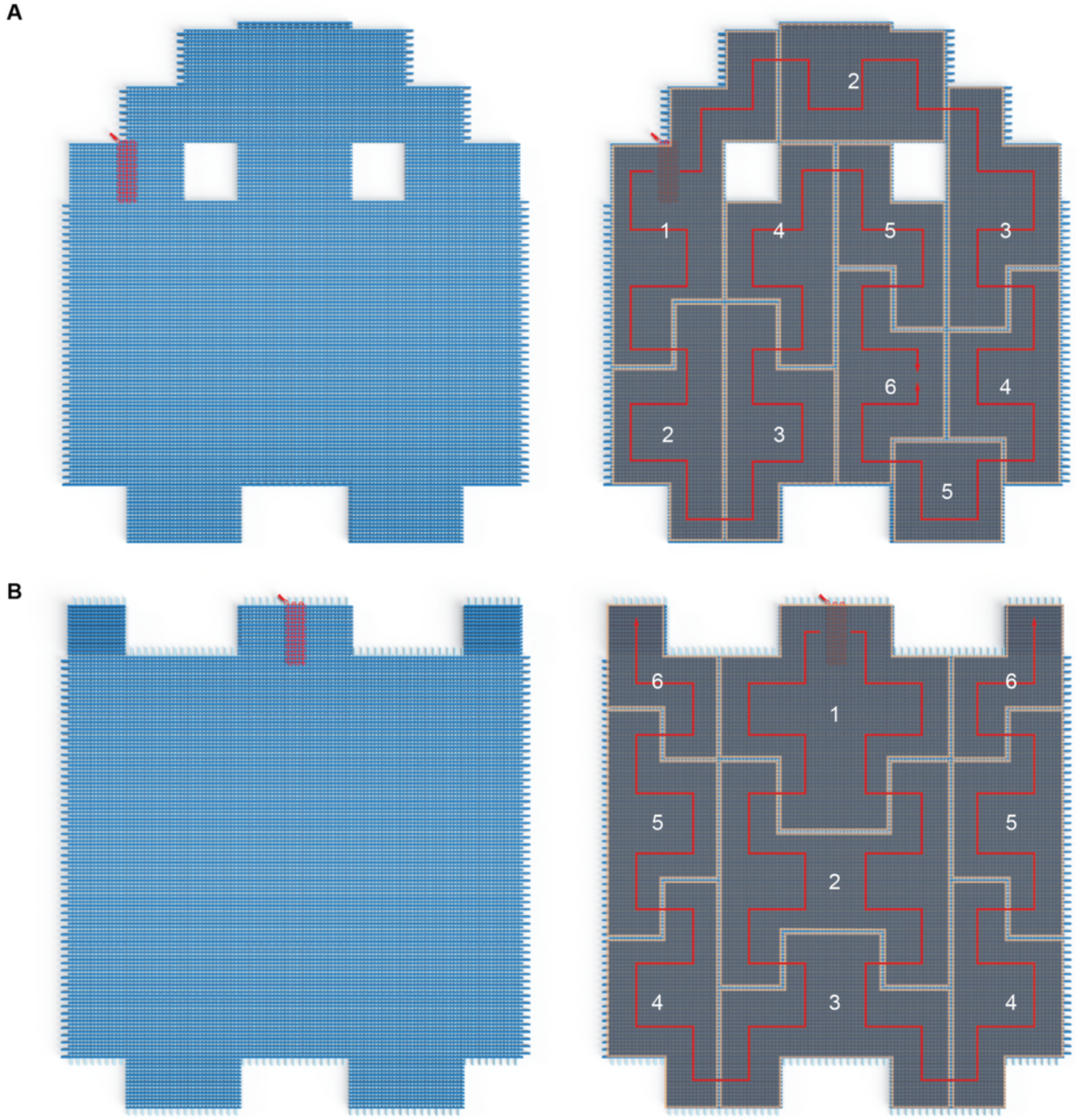

**Figure S10** Enlarged renders of the largest designs in Fig. 1C. **A** leftward is the 968-slat ghost and **B** leftward is the 1022-slat sheet. Rightward panels in *A* and *B* show the raster-fill growth as a red line, with the darkened overlays showing each of the distinct stages where up to 200 slats were added. The megastructures were incubated for about three days of isothermal growth before additional slats for the next stage were added. Images are not drawn to scale.

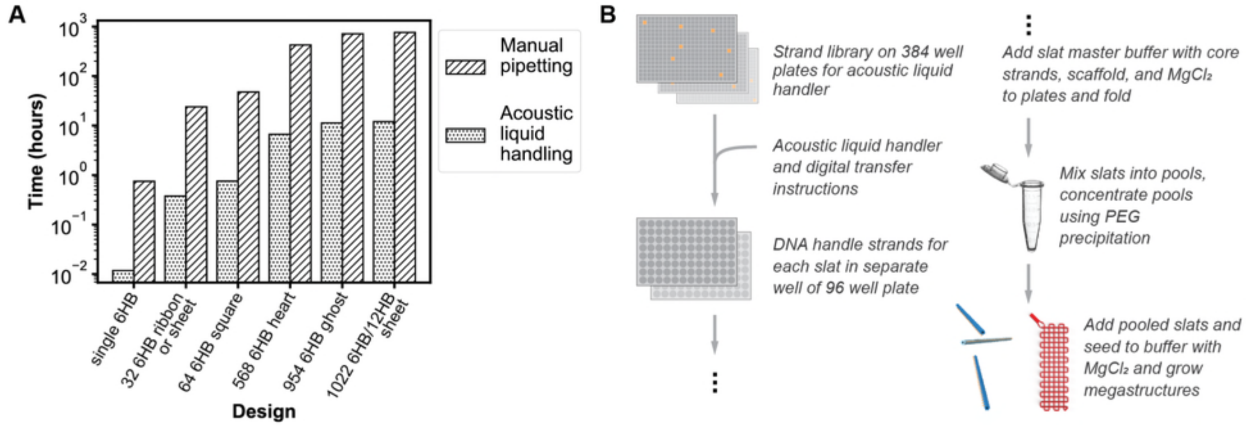

**Figure S11** General workflow to assemble DNA megastructures from slats. **A** shows approximate time to mix DNA handle strands for various crisscross megastructures from the 2048-strand library using an automated Labcyte Echo acoustic liquid handler versus a single-channel manual pipette. We determined an average transfer time of 42 seconds per strand using a manual pipette versus 0.66 seconds per strand using the liquid handler, as extrapolated using each method. These timings allow for preparations including manual loading of plates into the liquid handler, manual application of seals to the plates after automated strand transfer, and limited break times for the user. It would be untenable to make the largest megastructures from the strand library using a manual pipette. For instance, the 1022-slat finite sheet requires over 65,000 total strand transfers from the library and would require about one month of manual pipetting versus ~12 hours using the acoustic liquid handler. We note that the need for the acoustic liquid handler could be circumvented by purchasing the strands needed to make the slats for a given design arranged in an order where they could be manually combined using a multichannel pipette. However, this latter approach would not be amenable to having the strands readily rearranged into different crisscross megastructures. **B** shows the workflow for how megastructures are experimentally implemented from the strand library.

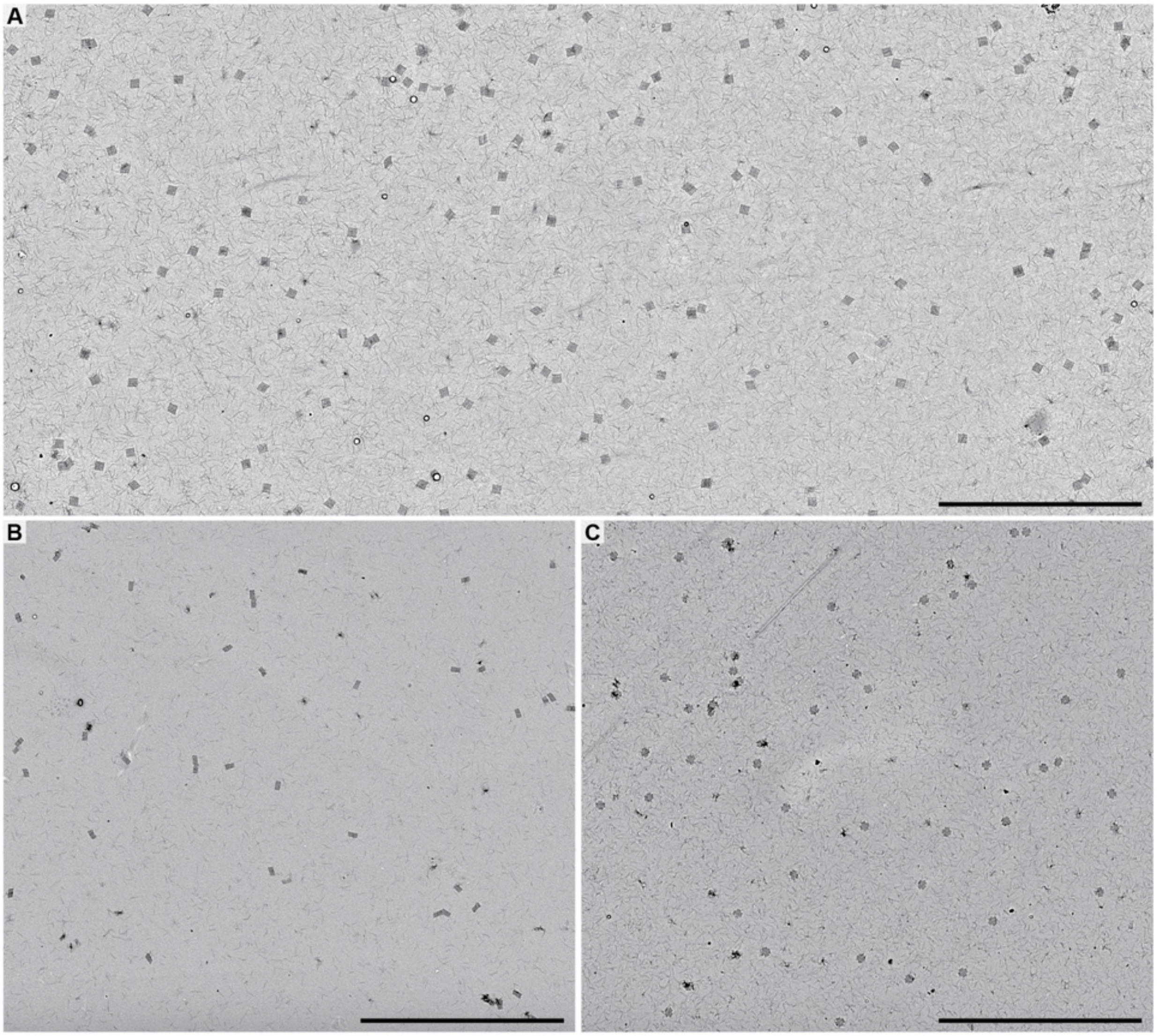

**Figure S12** Low-magnification TEM images of small finite shapes showing that they assemble as dispersed, single particles. **A** is the 64-slat square assembled with 6 nM seed (also see design in Fig. S9Aiii), **B** is the 48-slat square assembled with 1.5 nM seed (also see design in Fig. S9Aii), and **C** is the 64-slat plus symbol assembled with 6 nM seed (also see design in Fig. S9Aiv). Scale bars are 10 µm.

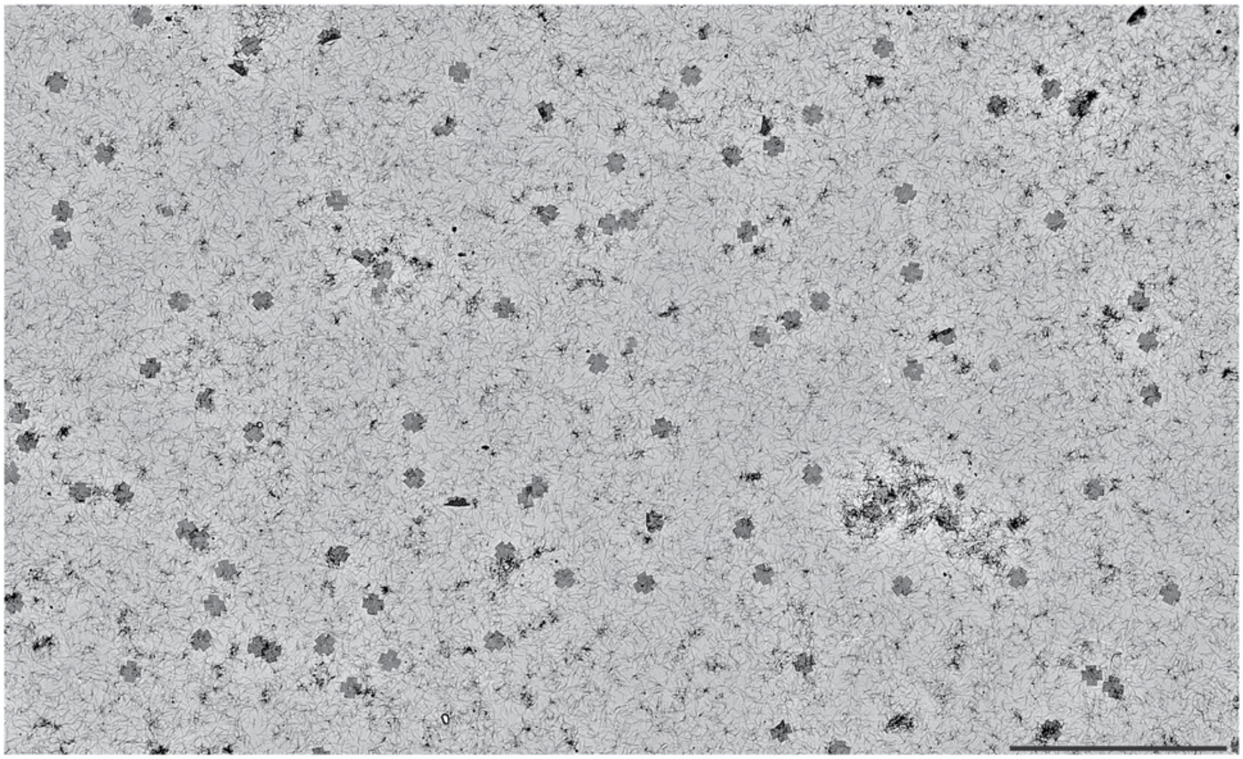

**Figure S13** Low-magnification TEM image of the rastering 191-slat plus symbol showing that it assembles as dispersed, single particles, with all slats added simultaneously to the reaction mixture. The raw assembly reaction used 1 nM seed with all the slats added simultaneously and a total isothermal incubation time of ~1.5 weeks. We note that ~3 days was a more typical incubation time for this number of slats, and the longer incubation time may explain why almost all the structures above have gone to completion. An aliquot of the final reaction was concentrated (and also to remove some of the excess slats) by centrifugation about 25-fold, so that it could be imaged with a higher density of particles as above. The scale bar is 10 μm.

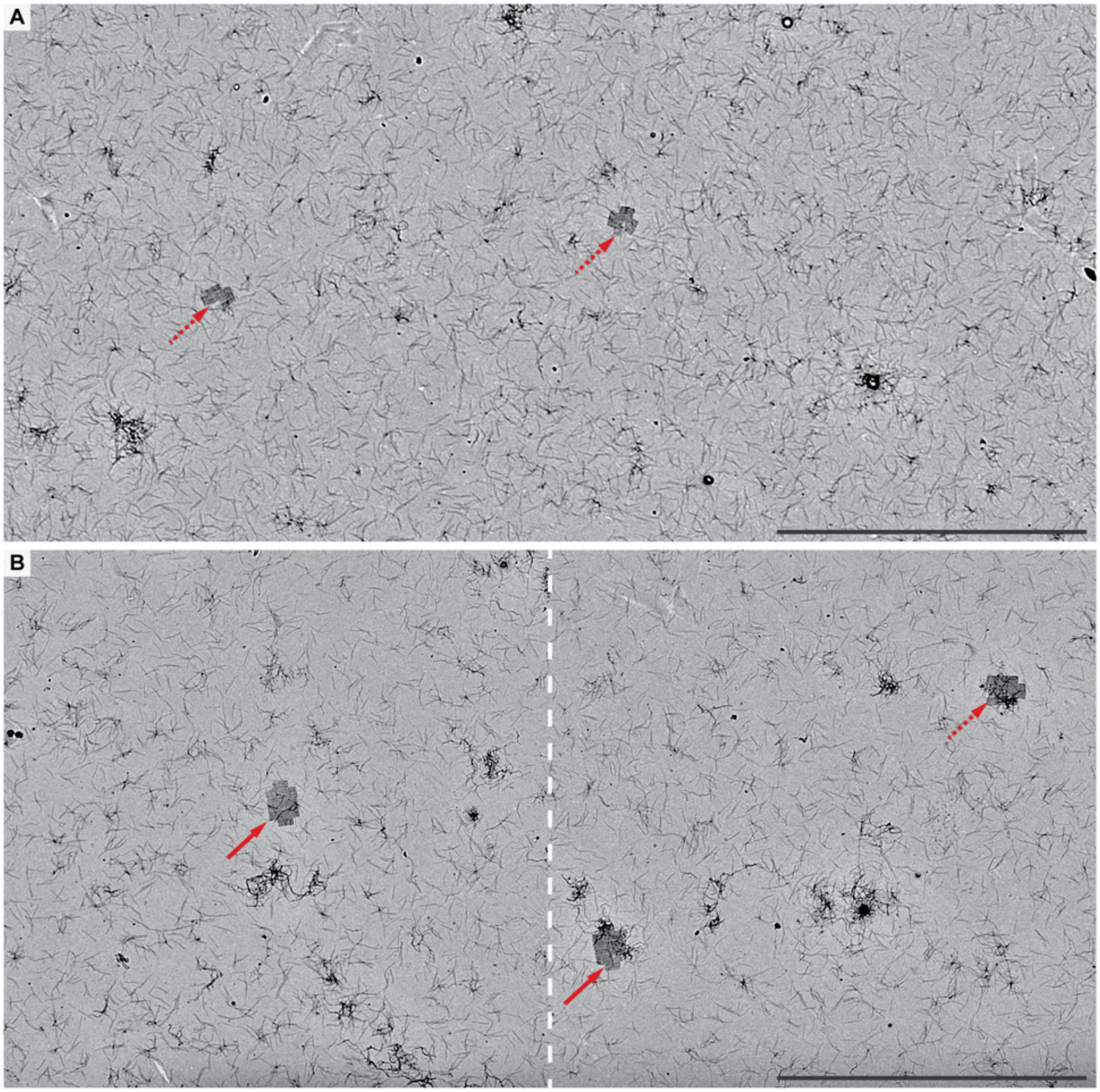

**Figure S14** Low-magnification TEM images of the finite elongated 320-slat plus symbol after 90 hours of isothermal growth. Solid-red versus dotted-red arrows show complete and incomplete assemblies respectively. In **A**, all 320 slats were simultaneously added during the initial preparation of the reaction, versus **B** where half of the total slats (i.e. 160 slats) were added during the initial preparation. After 38 hours of growth, the remaining 160 slats were added to the reaction in *B*. The completed shape was not observed after 90 hours using approach *A*, versus *B* where it was frequently observed. We note that the concentration per slat in *A* was ~3 nM, versus ~5 nM per slat in *B*. This was selected so as to ensure that the total slat concentration did not exceed ~1 μM, because extremely high concentrations of slats tended to impede growth as explained in Supplementary Text 3 and Fig. S21. Scale bars are 10 μm.

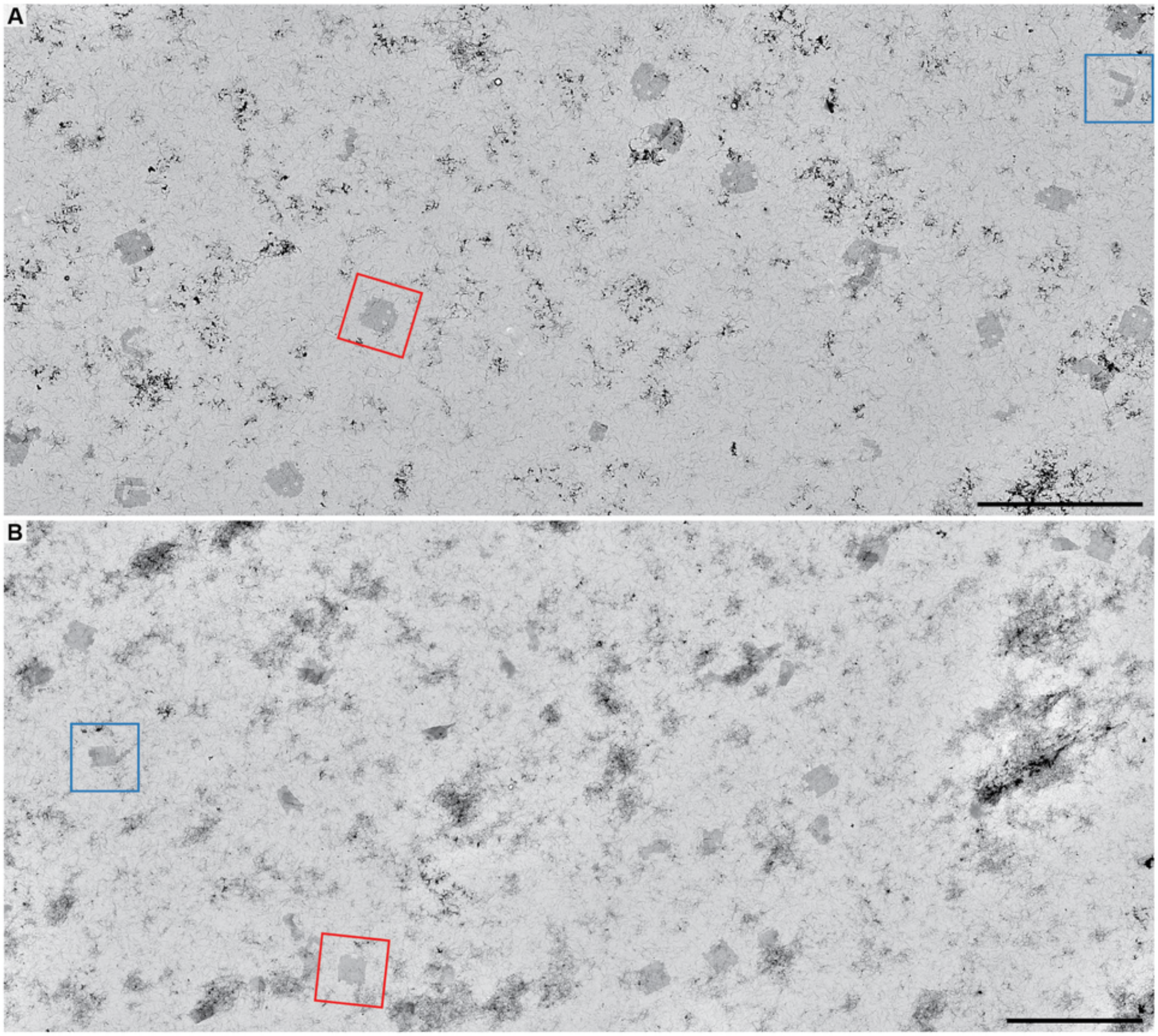

**Figure S15** Low-magnification TEM images of the largest finite 954-slat ghost (**A**) and 1022-slat sheet (**B**) to assess their relative completion. Completion of the megastructure was deemed all the corners and middle sections of the shape appropriately filled with slats. Examples in red boxes were deemed complete, versus examples in blue boxes were deemed incomplete. Roughly 22.5% of each of the above megastructures were complete by the end of the last stage of assembly. We made the assumption that incomplete structures at the end of a given assembly stage would not be able to continue growth once additional slats were added for the next stage of assembly. This assumes that partially formed structures at the end of a growth period are stalled/trapped because they have accumulated too many defective slats (e.g. those with missing or truncated handles). We considered that by adding more slats for the next growth stage, we would dilute out the remaining free slats needed to recover growth of a given stalled stage, to the extent that recovery would be exceedingly slow. We postulated that the probability for any given growth stage to go to completion could be treated as the probability of a series of independent events, suggesting that over 75% of the assemblies at each stage were complete and suitable for continuing growth (i.e. $0.225^{1/6}$, as per six distinct stages of assembly). The multi-stage addition of slats is described in Method 10. N = 225 seeded particles were counted for the 954-slat ghost, and N = 267 particles were counted for the 1022-slat sheet. Scale bars are 10 μm.

**Fig. S16–Fig. S27: Models of periodic megastructures, growth comparison of different ribbon designs, relationship between growth rate and number of unique slats, and additional TEM results of periodic megastructures**

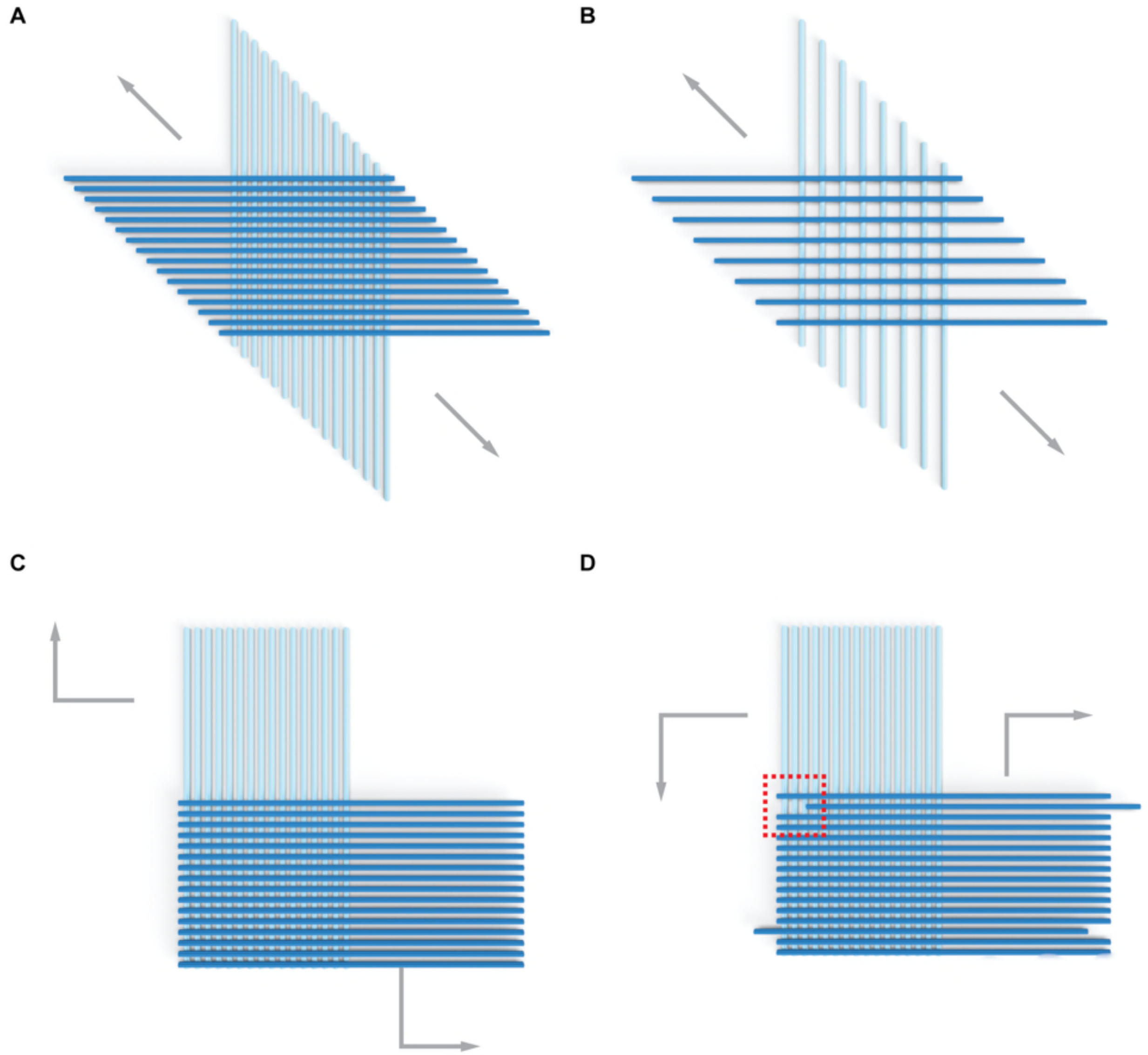

**Figure S16** Renders of the ribbon designs shown in Fig. 3. Slats are oriented parallel/perpendicular to the edges of the page to show differences in the staggering of the slats added to the ribbon. In **A** and **B**, each slat added staggers one and two 42-bp spacings respectively, compared to the parallel slat that preceded it. (note: such ribbons within the supplement are defined as the "staggered" designs). The v16 ribbon in *A* is differentiated from v8 ribbon in *B* by the maximum number of binding sites that an incoming slat may make with perpendicular slats on the growing ribbon front (i.e. 16 versus 8 binding sites). In the completed v16 ribbon, a given slat has 32 perpendicular slats bound to all of its 32 possible binding handles, versus v8 which only has 16 slats bound to every other of its 32 possible binding sites. Assuming the binding energy per binding site is the same, the greater number of binding sites allow the v16 designs to be grown under more stringent reaction conditions (e.g. higher temperatures) where the rate of spontaneous nucleation is lower. The differences in nucleation performance are experimentally explained in Supplementary Text 4.1 and Fig. S38, and the energetic models explaining these differences are as discussed previously[33]. **C** and **D** are v16 ribbons where each slat added does not stagger compared to either immediate parallel slat, creating "zig-zag" ribbons with jagged and flush edges. In *D*, two of the horizontal top blue slats were each staggered by two 42-bp units (as boxed in red) to seal the vertical seams (note: such ribbons within the supplement are defined as the "non-staggered" designs).

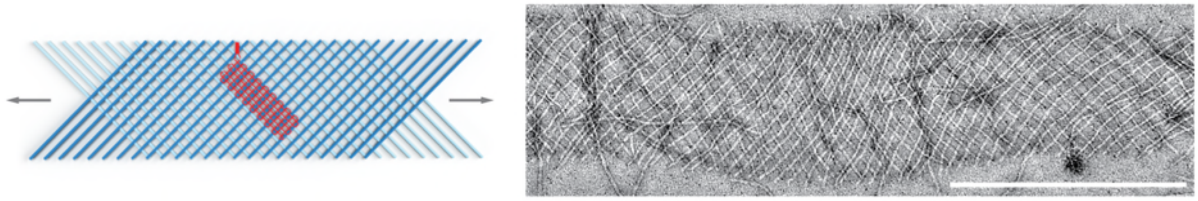

**Figure S17** Design and testing of the periodic v8 ribbon. The v8 design uses every other binding site on the 6HB slat, with 16 slats bound to every other of the 32 possible binding sites. There is half the density of slats compared to its v16 counterpart. As such, the v8 ribbons were flexible and had a propensity to stretch and elongate compared to v16 ribbons which remained straighter and flatter (also see Fig. S18B versus Fig. S18A). The close-up rightward negative-stain TEM image shows a flat segment of ribbon, with the inherent flexibility causing the curving, meandering ribbon edge. The scale bar is 500 nm.

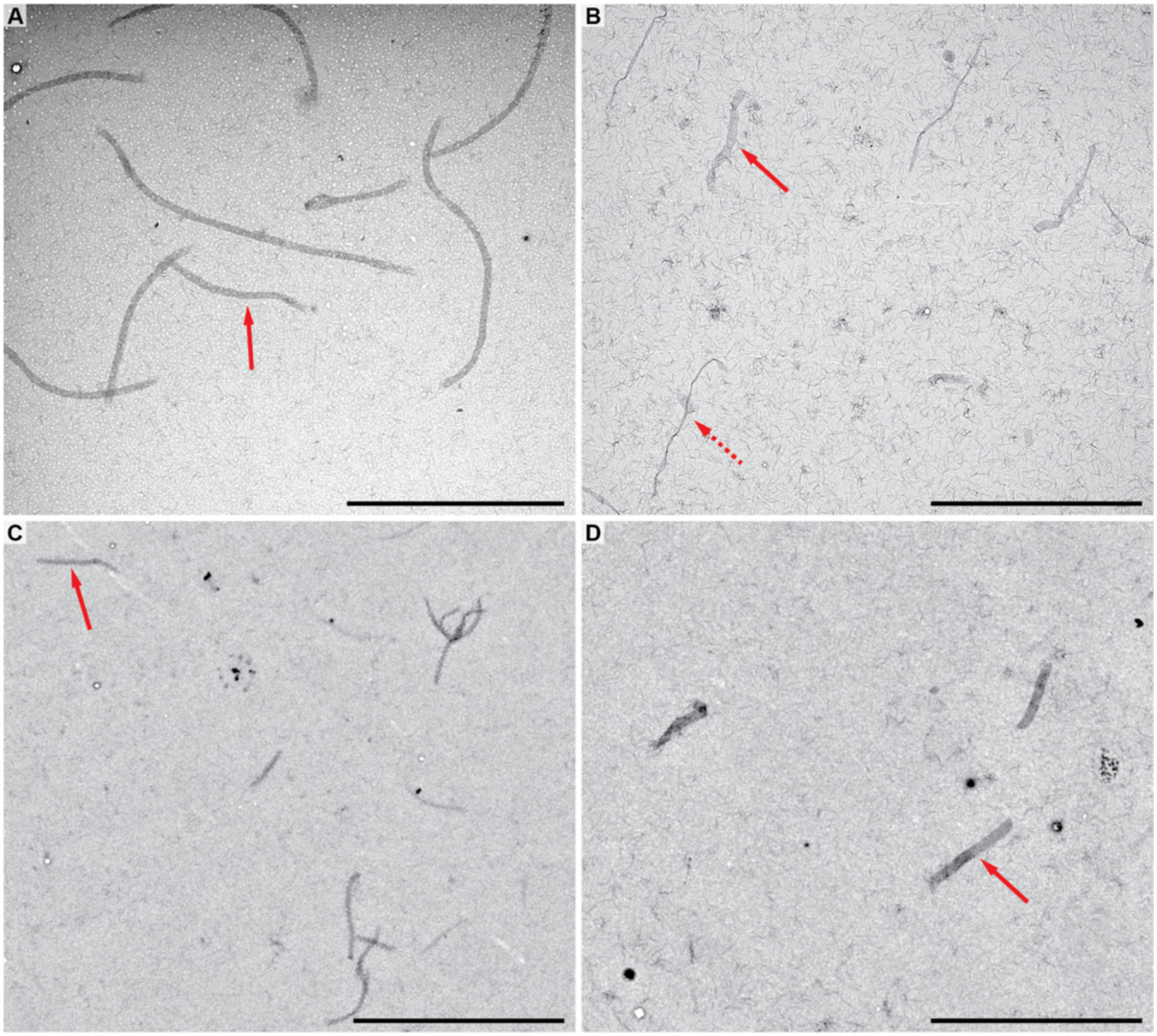

**Figure S18** Low-magnification TEM images of periodic 1D ribbons. Images in **A–D** correspond to the designs v16 staggered, v8 staggered, v16 non-staggered jagged, and v16 non-staggered flush ribbons respectively (also see Fig. 3i–iv, Fig. S11). Solid-red arrows point to flat-lying ribbons. The dotted-red arrow in *B* shows how the v8 ribbons had a propensity to adopt stretched and elongated morphologies on the TEM grids. Scale bars are 10 µm.

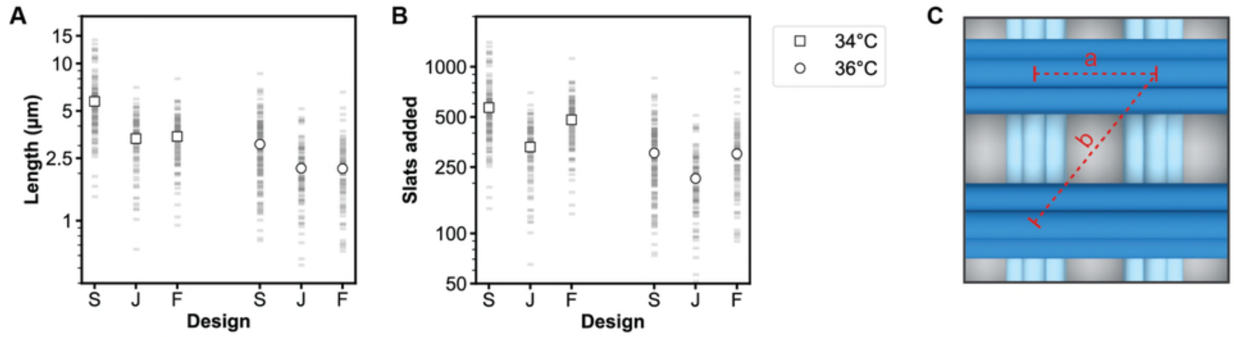

**Figure S19** Length comparison of the various designs of v16 ribbons after 16 hours of isothermal growth. **A** shows the lengths as measured in low-magnification TEM images, versus **B** which shows the number of slats per ribbon as extrapolated using the theoretical nm per slat as calculated in **C**. Each faint gray bar in the plots is the measurement of a single ribbon, with the mean lengths shown with either the square or circular data points. 'S', 'J', and 'F' indicate the v16 staggered, non-staggered jagged, and non-staggered flush ribbons. The ribbons for the various designs for a given temperature grew to roughly similar numbers of slats (e.g. 200–300 slats at 36°C), suggesting that the pattern in which the slats are added does not greatly influence the kinetics of assembly. Small differences in growth might be due to small variations in experimental parameters or differences in sequence symmetry of the designs. The v16 staggered ribbons used 4x sequence symmetry, versus 2x sequence symmetry of the v16 non-staggered jagged and flush ribbons (see Supplementary Text 3 for an explanation of the relationship between symmetry and growth). In *C*, dimension 'a' is 42 bp or ~14.2 nm, versus dimension 'b' which is ~20.2 nm. We can see by considering the pattern of slat as per the design renders in Fig. 3: each pair of slats for the v16 staggered ribbons and v16 non-staggered jagged ribbons add one unit of 'b'; conversely, each pair of slats for the v16 non-staggered flush ribbons add one unit of 'a'. $N_{S\_34°C}$ = 133, $N_{J\_34°C}$ = 104, $N_{F\_34°C}$ = 109, $N_{S\_36°C}$ = 170, $N_{J\_36°C}$ = 102, and $N_{F\_36°C}$ = 95 ribbons were measured.

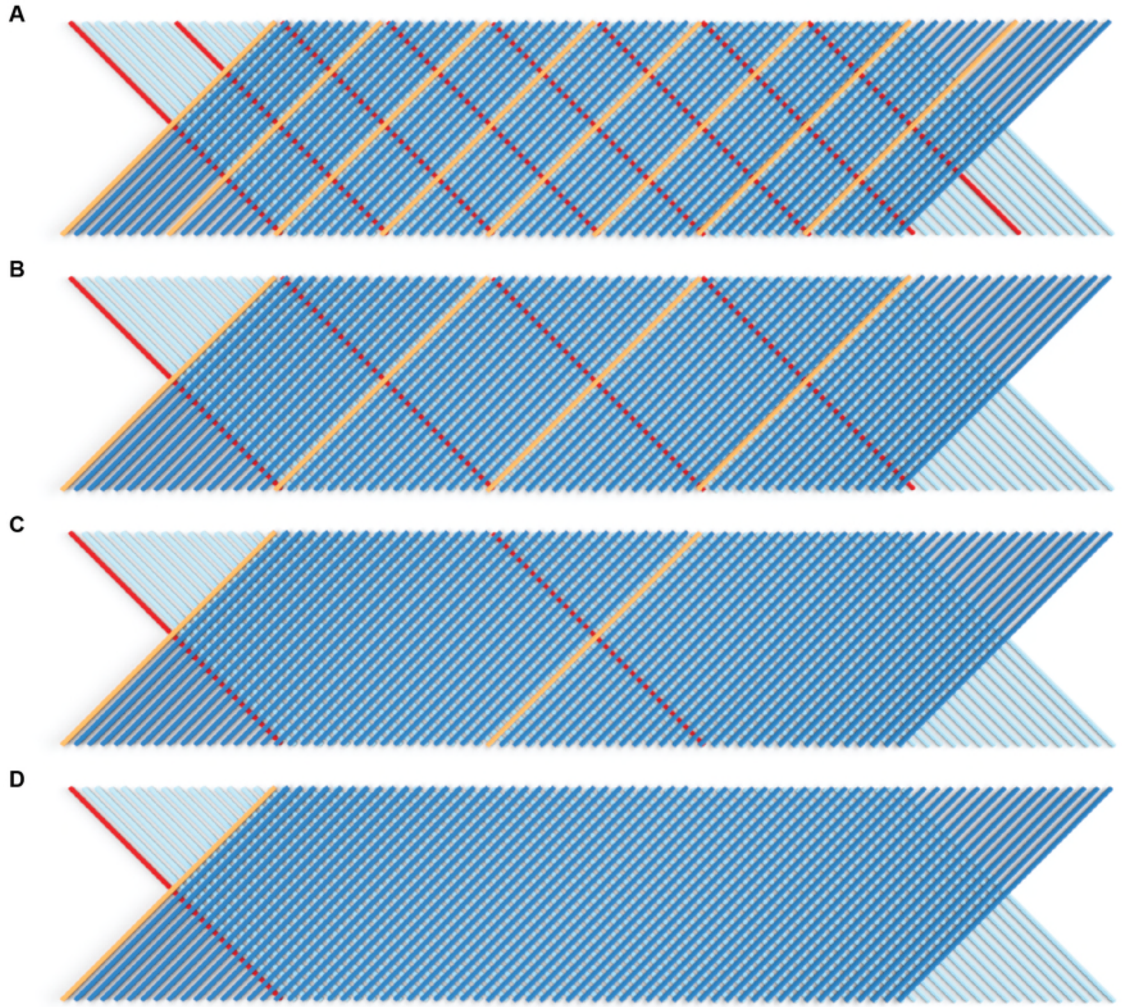

**Figure S20** Design renderings showing variable sequence symmetries of the v16 staggered ribbons. That is, the number of slats that repeat in the periodic unit is arbitrary and can be programmed with as many unique slats as desired. Alternatively, sequence symmetry of a given slat is the copy number of another particular slat from the perpendicular direction to which it is bound. This is observed in the above where a given slat from the top-layer is colored orange, and a given slat in the bottom-layer that is colored red. **A** is 4x symmetry with 16 unique slats (i.e. 8+8 slats in each respective perpendicular layer), **B** is 2x symmetry with 32 unique slats (i.e. 16+16), **C** is 1x symmetry with 64 unique slats (i.e. 32+32), and **D** is 0.5x symmetry with 128 unique slats (i.e. 64+64). The v16 staggered ribbons in Fig. 3i use the 4x symmetry, the v8 staggered ribbons in Fig. 3ii use the 2x symmetry, the non-staggered jagged and flush ribbons in Fig. 3iii–iv use the 2x symmetry, the v16 staggered sheets in Fig. 4 and Fig. 5 use the 2x sequence symmetry. All origami crisscross growth characterization with respect to reaction conditions was studied using v16 staggered ribbons with 4x symmetry, or v8 staggered ribbons with 2x symmetry. In Fig. S21, growth versus symmetry design as shown above are compared.

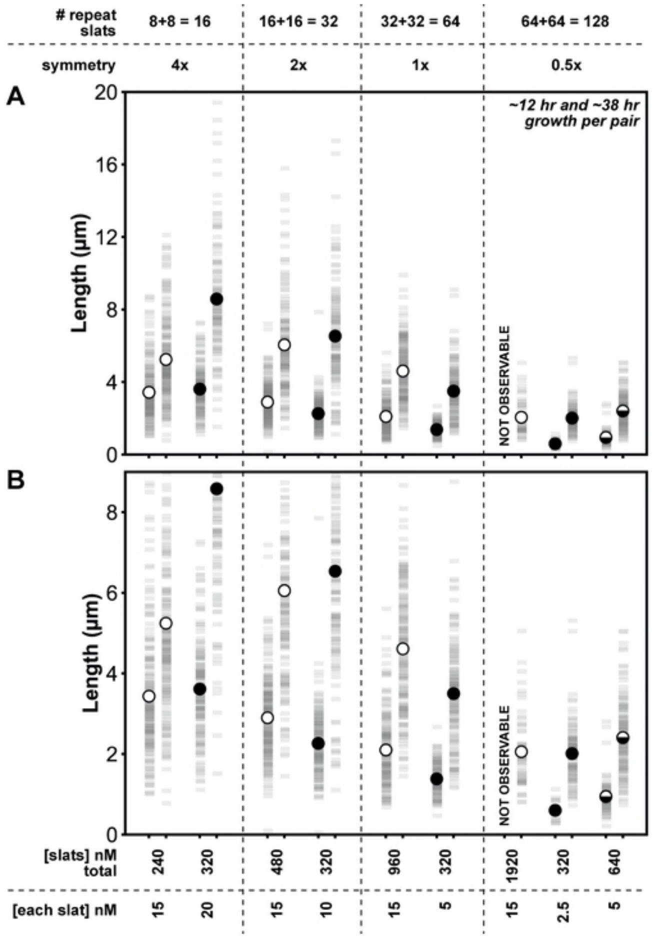

**Figure S21** Length of v16 staggered ribbons versus the number of unique slats in the unit repeat of the ribbon (i.e. symmetry as explained in Fig. S20), as obtained by measuring ribbons in TEM images after 12 and 38 hours of isothermal 34ºC growth. Panels **A** and **B** are the same data, with the y-axis in ***B*** over a smaller range to better show the length differences. Each faded rectangle is the measurement of a single ribbon, with circular data points showing the mean length. In general, designs with a larger number of unique slats (i.e. lower symmetry) grew more slowly. We observed that the concentration of the slats is an important determinant for the rate of growth of a given symmetry design. The white data points indicate where we maintained the per slat concentration at 15 nM, versus the black data points where we maintained the total concentration of slats at 320 nM. The following number of ribbons were measured for each condition: $N_{8+8\text{ slats; 15 nM [each slat], 12 hr}} = 177$, $N_{8+8\text{ slats; 15 nM [each slat], 38 hr}} = 192$, $N_{8+8\text{ slats; 20 nM [each slat], 12 hr}} = 147$, $N_{8+8\text{ slats; 20 nM [each slat], 38 hr}} = 117$, $N_{16+16\text{ slats; 15 nM [each slat], 12 hr}} = 217$, $N_{16+16\text{ slats; 15 nM [each slat], 38 hr}} = 147$, $N_{16+16\text{ slats; 10 nM [each slat], 12 hr}} = 146$, $N_{16+16\text{ slats; 10 nM [each slat], 38 hr}} = 129$, $N_{32+32\text{ slats; 15 nM [each slat], 12 hr}} = 169$, $N_{32+32\text{ slats; 15 nM [each slat], 38 hr}} = 179$, $N_{32+32\text{ slats; 5 nM [each slat], 12 hr}} = 117$, $N_{32+32\text{ slats; 5 nM [each slat], 38 hr}} = 143$, $N_{64+64\text{ slats; 15 nM [each slat], 12 hr}} = 0$, $N_{64+64\text{ slats; 15 nM [each slat], 38 hr}} = 58$, $N_{64+64\text{ slats; 2.5 nM [each slat], 12 hr}} = 15$, $N_{64+64\text{ slats; 2.5 nM [each slat], 38 hr}} = 115$, $N_{64+64\text{ slats; 5 nM [each slat], 12 hr}} = 78$, $N_{64+64\text{ slats; 5 nM [each slat], 38 hr}} = 132$.

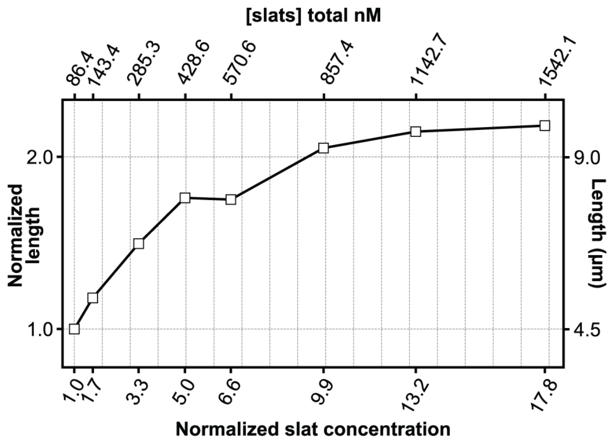

**Figure S22** Normalized mean lengths of the ribbons are shown versus the concentration of the slats for the 4x sequence symmetry design (i.e. 16 total unique slats) using a constant concentration of seed. There is roughly a linear change in the mean length versus the slat concentration when the total slat concentration is below 500 nM, versus a lesser increase in length for when higher concentrations of slats were used. This experiment is replotted on a linear axis with the data from Fig. 5D. The dotted grid lines are spaced to represent an increase in either the length or total slat concentration of one unit compared to the condition to which the data was normalized.

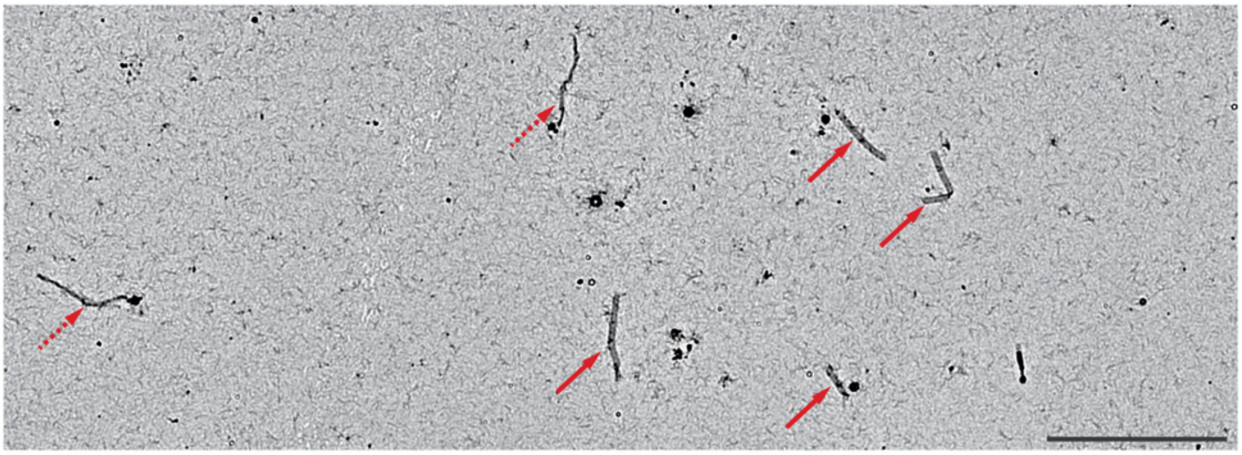

**Figure S23** Tri-layer arrangement of slats on a v8 staggered ribbon for 2D growth, as shown in an overview image of the ribbon sample from Fig. 3B. The initial v8 ribbon with two layers of slats was grown isothermally overnight, at which point the additional third layer of slats was added and incubated for two further days. The ribbons with solid red arrows were those where a third layer was successfully added, versus the ribbons with dashed arrows where the third layer did not bind. We note that binding of the third layer appeared to be an all-or-nothing event for a given ribbon. It is unclear why the third layer slats did not bind to certain ribbons and will require further study and optimization to improve its efficiency. The scale bar is 10 µm.

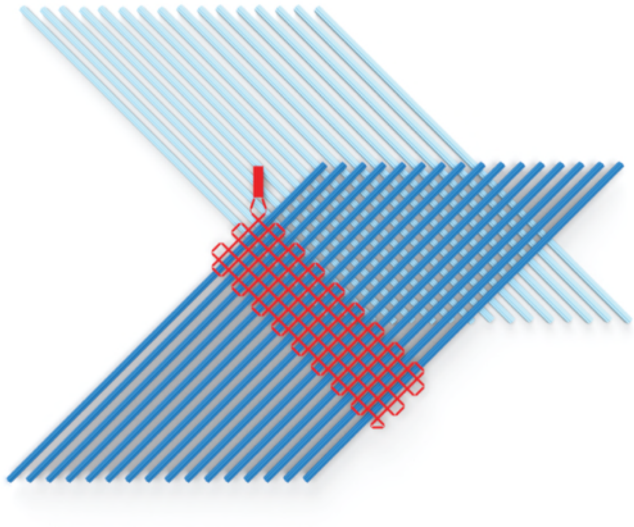

**Figure S24** Render of the repeating slat unit for periodic growth of 2D sheets. The pattern of growth is similar to the v16 staggered ribbons (see Fig. S16A), except that here the top- and bottom-layers of slats are shifted with respect to one another. The particular render and designs as tested in this paper used 2x sequence symmetry and were composed of 32 unique slats total.

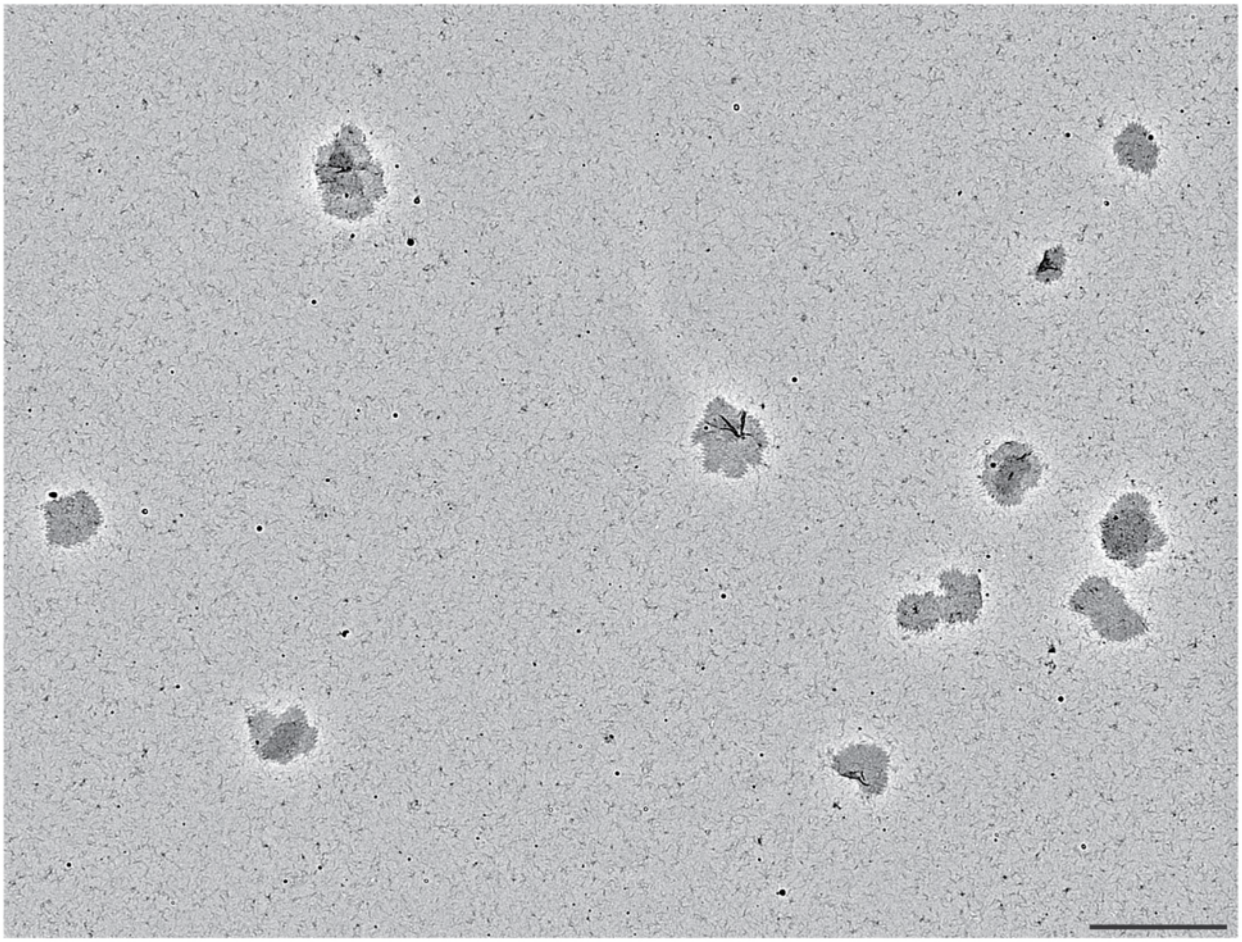

**Figure S25** Low-magnification TEM images of the periodic sheets after three days of isothermal growth.

The sheets were grown with 0.2 nM seed and generally appeared as well distributed single particles. The scale bar is 10 µm.

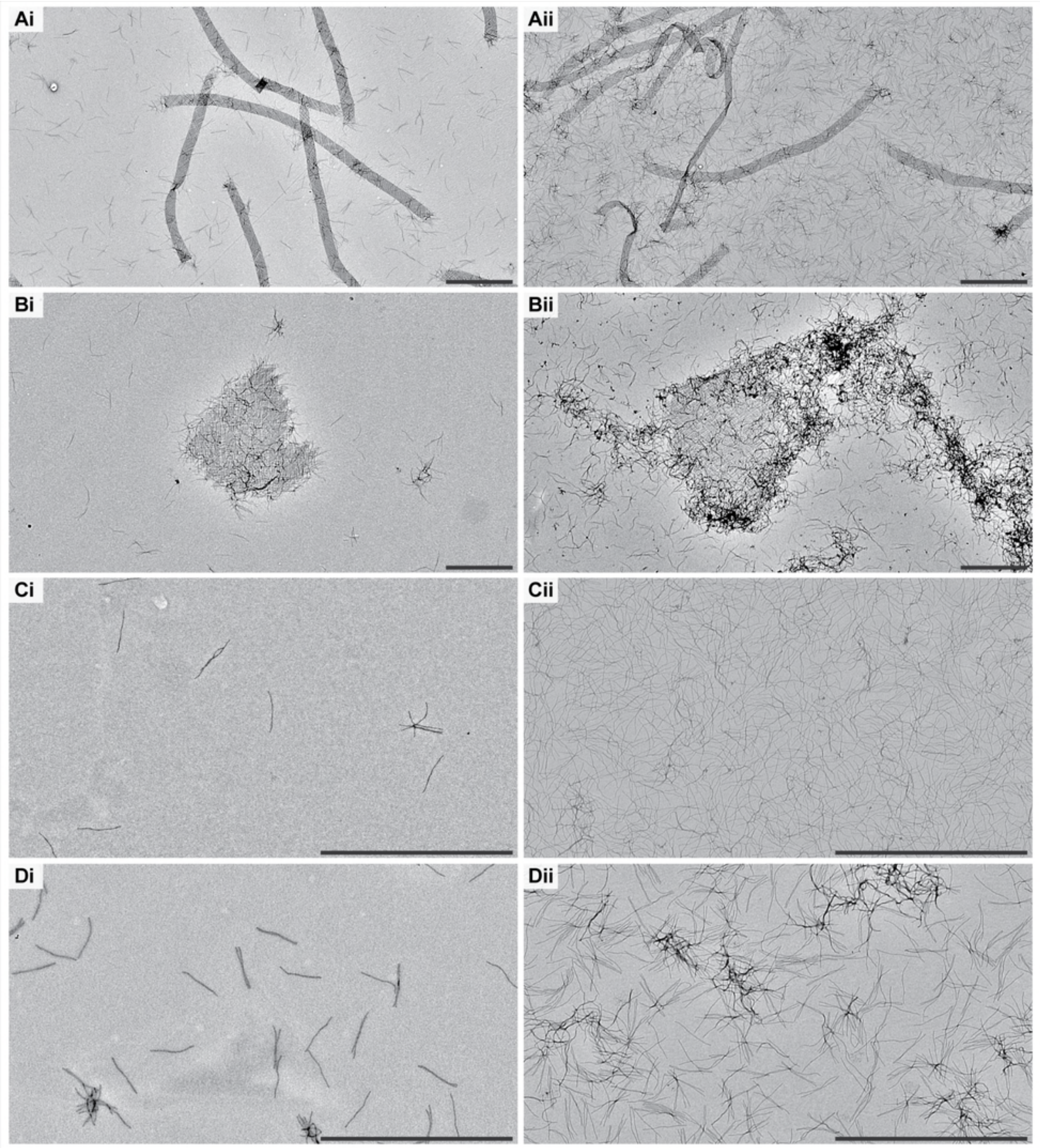

**Figure S26** Large periodic megastructures can be purified from excess free slats to a limited extent by low speed centrifugation. Rows **A–D** are ribbons, sheets, one layer of unassembled ribbon slats with no seed as one control, and both layers of unassembled ribbon slats with no seed as another control, respectively. In column **i**, the samples were purified as described in Method 12, versus column **ii** which is the raw sample with no centrifugation. There are qualitatively fewer background free slats in the resuspended pellets in comparison to the raw samples. We note that the background free slats could be further lessened by repeating the centrifugation process two or more times. However, we were unable to extract all the background slats by centrifugation alone. There was a higher background of free slats in sample *Di* where the control slats had both handles and complementary handles from the other perpendicular layer, versus *Ci* where the control slats only had handles from one layer. We assume that transient interactions between free slats with complementary handles makes them prone to pelleting during centrifugation and that is the reason we could not completely remove the slat background in the megastructures samples in *A* and *B* by centrifugation alone. Scale bars are 2 µm.

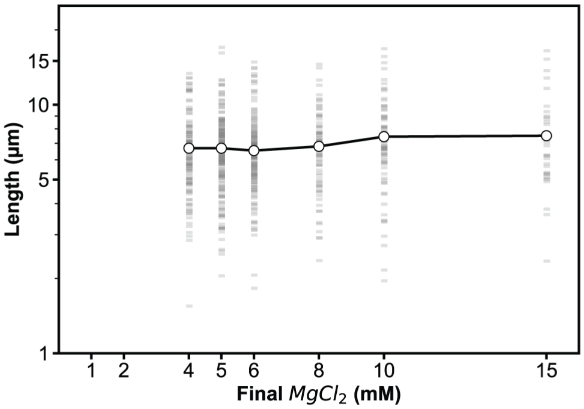

**Figure S27** Megastructures (i.e. periodic v16 ribbons) using the 7-nt binding sites were stable in lower-magnesium conditions from which they were initially grown. The ribbons were initially grown in 15 mM $MgCl_2$, then diluted 15-fold into the concentration of $MgCl_2$ as listed on the x-axis, incubated for ~48 hours at room temperature. There was negligible difference in the length of the ribbons that were incubated in 15 mM $MgCl_2$ versus those that were incubated in 4 mM $MgCl_2$. No ribbons could be observed in either 1 mM or 2 mM $MgCl_2$ conditions, where they were presumed to have fallen apart. Each faint gray bar is the length measurement of a single ribbon, with the mean length indicated by the circular, white data point. N ribbons were measured for the conditions tested: $N_{4\,mM}$ = 121, $N_{5\,mM}$ = 188, $N_{6\,mM}$ = 171, $N_{8\,mM}$ = 78, $N_{10\,mM}$ = 79, $N_{15\,mM}$ = 38.

**Fig. S28–Fig. S31: Model of the DNA nanocube, additional TEM of nanocube patterns, and DNA-PAINT results of 1D ribbons and 2D sheets**

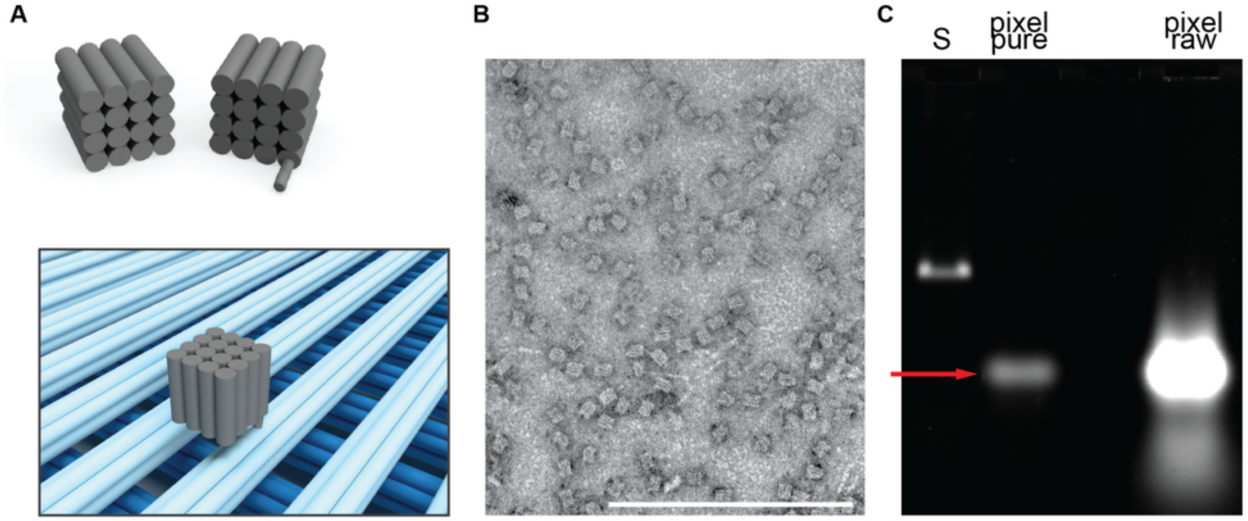

**Figure S28** Strategy and folding of the DNA nanocube contrast agent to visualize arbitrary patterns on 6HB megastructure canvases. Uppermost **A** shows a rendering of the 10 × 10 × 10 $nm^3$ DNA nanocube, where we added a 16-nt handle with a 4T linker to the 3′ end of one of the strands in the nanocube as published previously[37]. The lowermost rendering shows the nanocube bound to the top layer of slats in a megastructure. The 3′ end of one strand in the top helix of a cyan 6HB was extended with the 16-nt complementary handle at one of the 32 possible addressable sites. The negative-stain TEM image in **B** shows the nanocube after purification by excising them from an agarose gel. **C** shows an agarose gel of the nanocubes after and before purification, with the desired band shown with the solid red arrow. Densitometry of the raw folded nanocube suggests that ~64% of the total material was assembled into the desired structure. The scale bar in the TEM image is 200 nm.

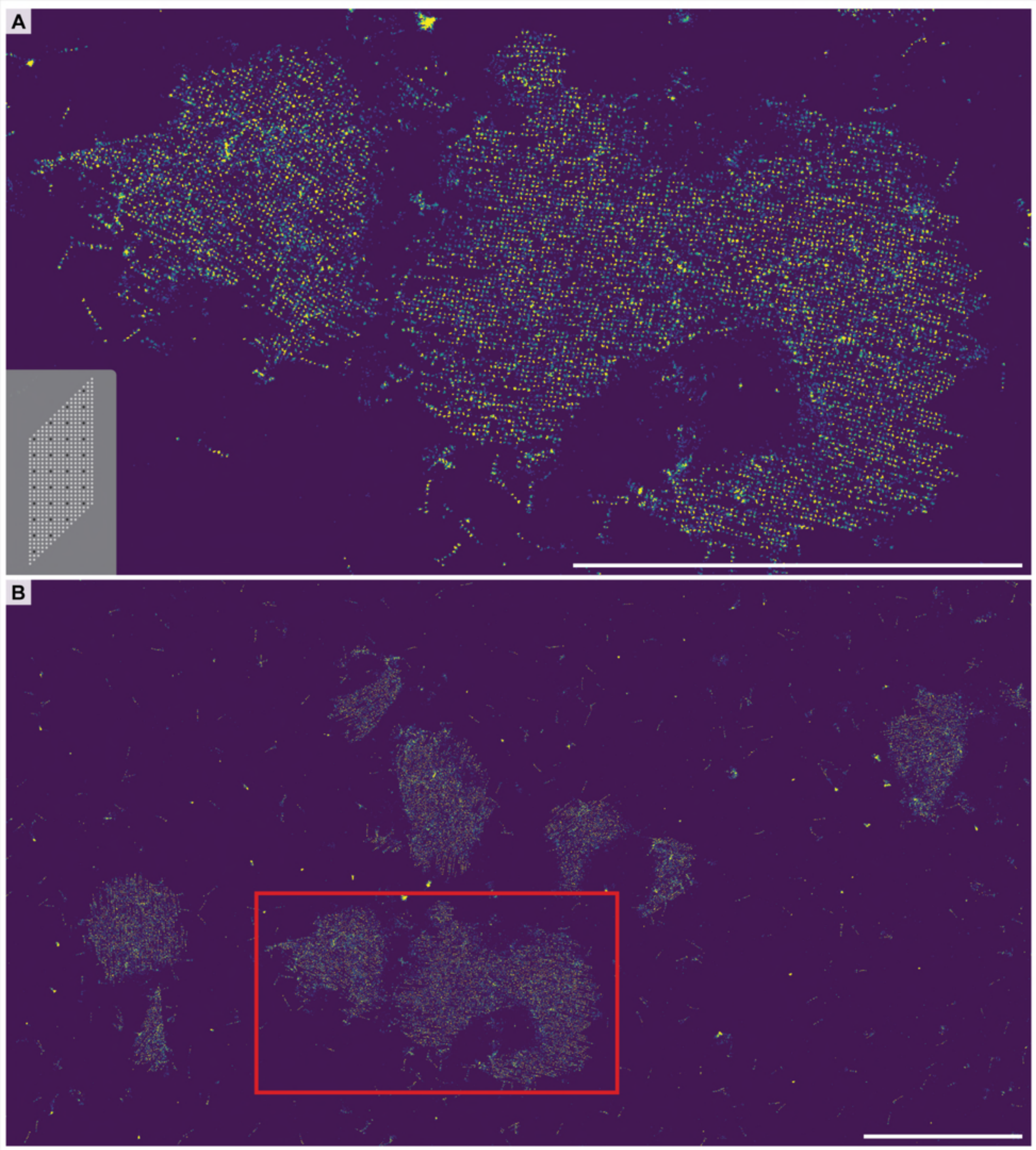

**Figure S29** DNA-PAINT overview images of 2D sheets (also note, this is the same sample as shown in Fig. 4C) where the top layer of slats is tagged with complementary handles to the PAINT imager strands. The close-up region shown in **A** was selected from the boxed region in **B**. How the top slats are patterned is shown with the dark dots in the lower left of *A*, where every fourth node (~56.3 nm or 168 bp spacings) was decorated. We achieved single handle resolution, with each dot in the above images corresponding to a single node on the canvas. The base of the sheets were intermittently decorated with biotinylated strands for docking to the imaging substrate. Scale bars are 5 µm.

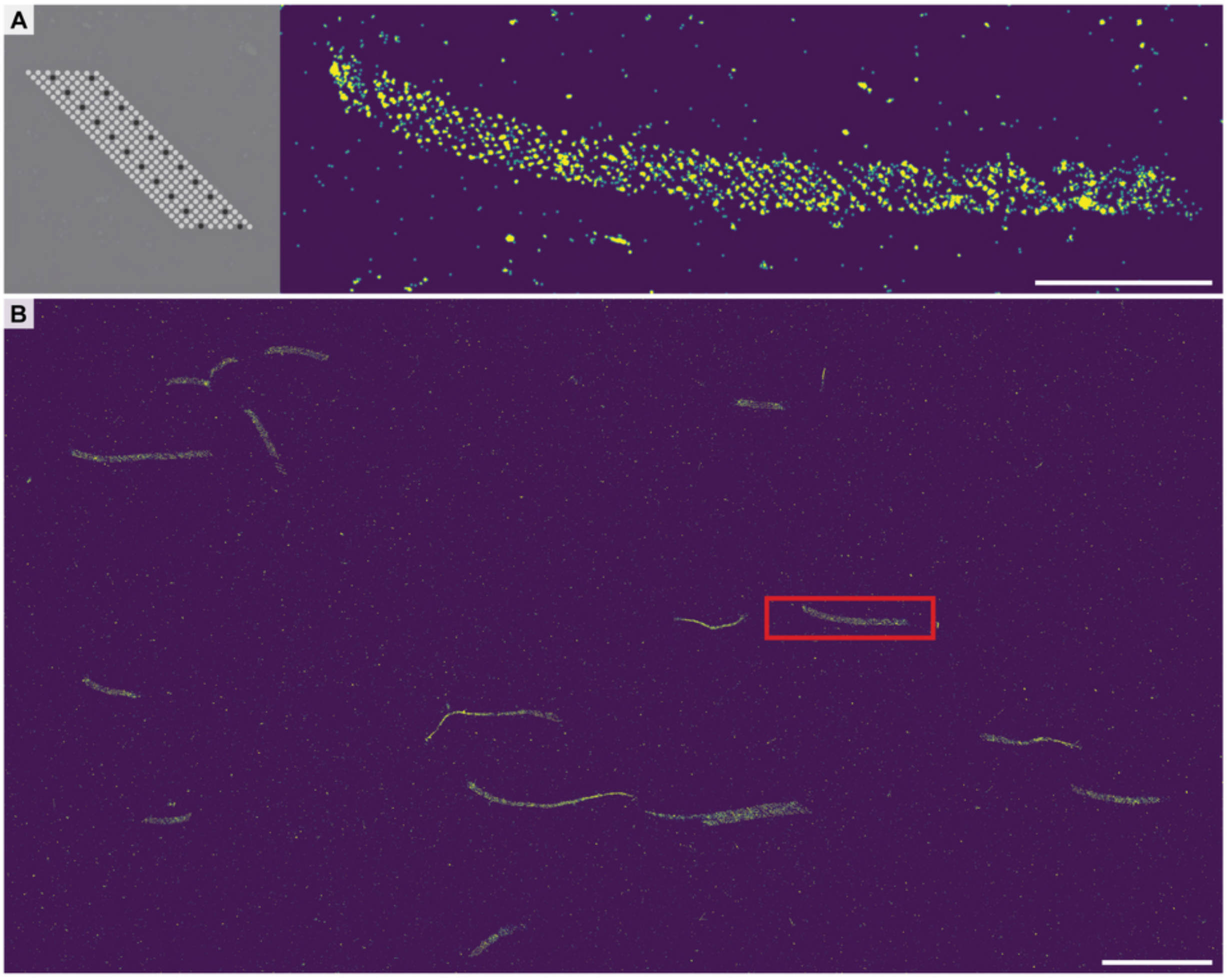

**Figure S30** DNA-PAINT overview images of periodic v16 staggered ribbons where the top layer of slats is tagged with complementary handles to the PAINT imager strands. The close-up region shown in **A** was selected from the boxed region in **B**. How the top slats are patterned is shown with the dark dots in the left of *A*, where minimally every third node (~42.2 nm or 126 bp spacings) was decorated. We achieved single handle resolution on some examples of slats in *A*, though the resolution of the single handles was not as clear as in Fig. S29. We also note that the ribbons had a propensity to not bind completely flat to the imaging substrate and frequently appeared stretched and elongated. The base of the sheets were intermittently decorated with biotinylated strands for docking to the imaging substrate. Scale bars are 1 µm in *A* and 5 µm in *B*.

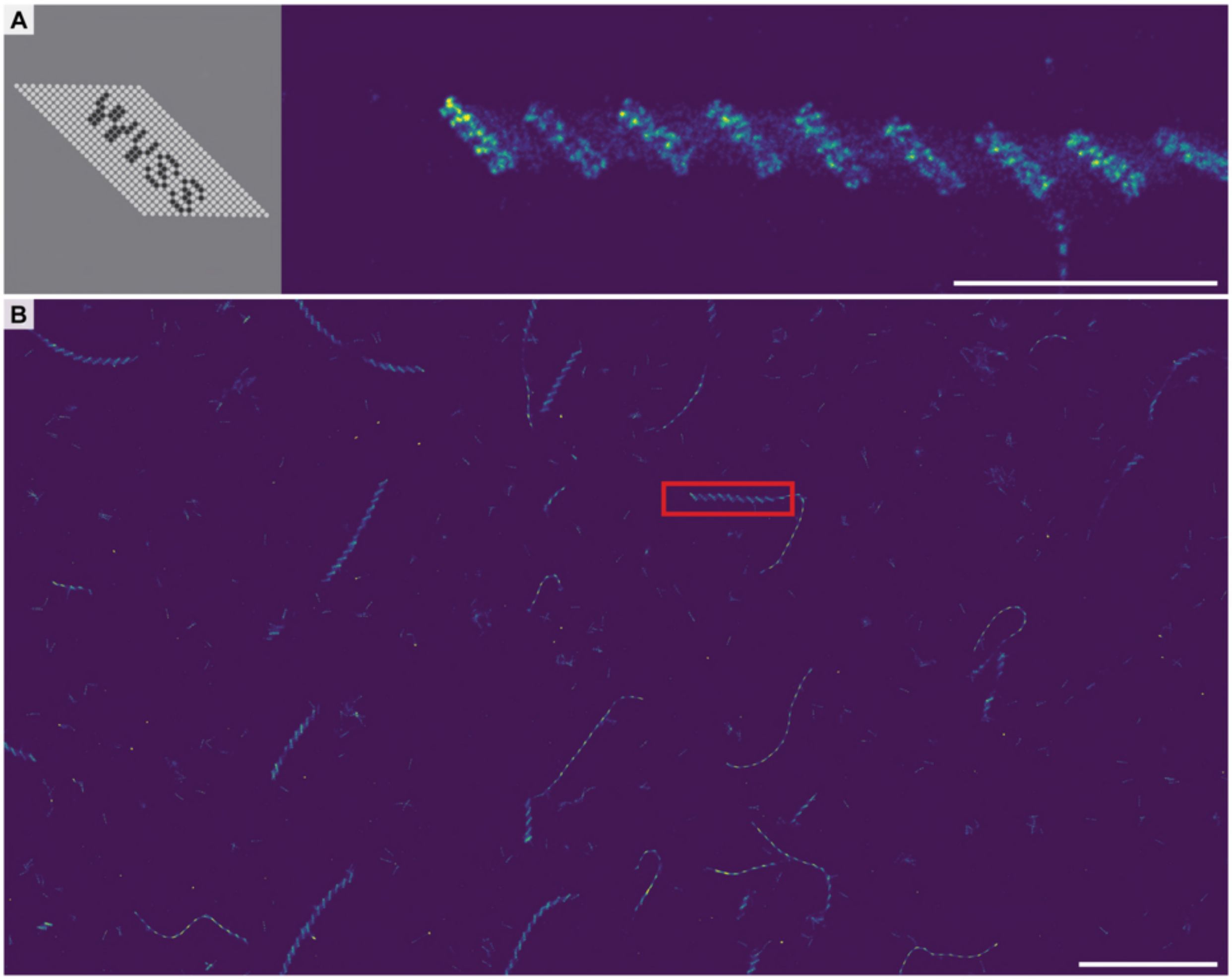

**Figure S31** DNA-PAINT overview images of 1D v16 staggered ribbons where the top layer of slats is tagged with complementary handles to the PAINT imager strands, where the pattern resembles the lettering "WYSS." The close-up region shown in **A** was selected from the boxed region in **B**. How the top slats are patterned is shown with the dark dots in the left of *A*, where minimally every node (~14.1 nm or 42 bp spacings) was decorated. The resolution of single handles could not be distinguished, though there is resemblance to the designed pattern. We also note that the ribbons had a propensity to not bind completely flat to the imaging substrate and frequently appeared stretched and elongated. The base of the sheets was intermittently decorated with biotinylated strands for docking to the imaging substrate. Scale bars are 1 μm in *A* and 5 μm in *B*.

**Fig. S32–Fig. S34: TEM results when no seed added, AGE results of single DNA origami square versus scaffold, representative TEM of megastructures versus concentration of seed**

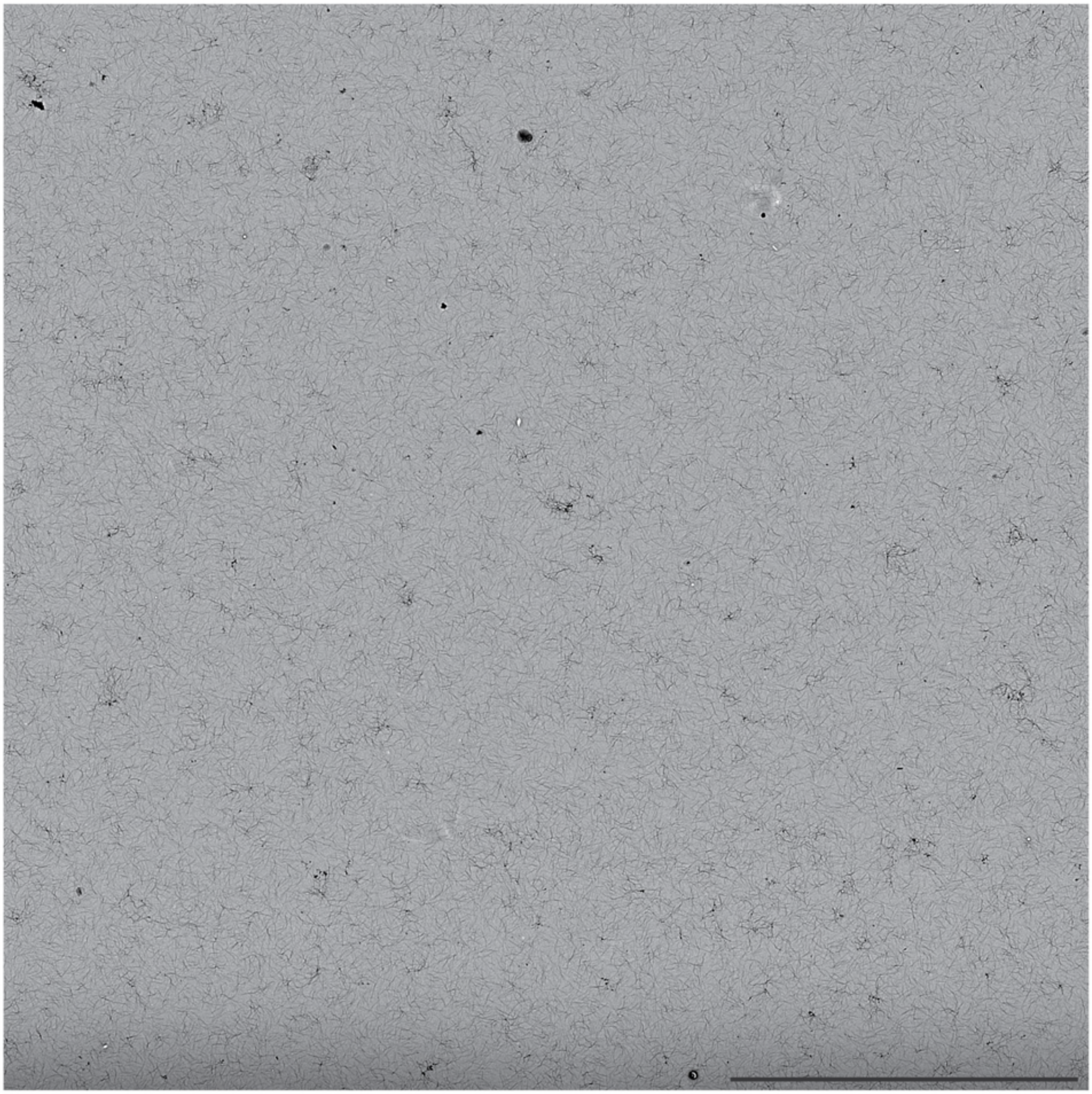

**Figure S32** Low-magnification TEM image of a typical megastructure assembly reaction when no seed is added. No assembly was observed in the control for the finite 1022-slat sheet, as above. This was similarly observed in all control reactions for other finite and periodic megastructures from Fig. 2 –Fig. 4, with typical reaction conditions where fast, seeded growth is favored. For brevity, we show only the image of a single design here. The particular reaction as above is perhaps an extreme challenge of the nucleation control attainable with origami slats; slats were added in multiple three-day stages for the largest structures as explained in Fig. S15. As such, the slats in the above 1022-slat sheet control were incubated at 34°C for over 18 days. In this particular experiment, we also had the thermocycler where we were growing the reaction fail. The reaction sat at room temperature for about one day before we noticed the breakdown, and despite this extended incubation at both these temperatures, no spontaneous growth could be observed. We argue that this leeway with experimental conditions where useful nucleation control can still be attained is a large factor of what makes crisscross assembly of origami a powerful advance to make it easier to create DNA megastructures. The scale bar is 10 μm.

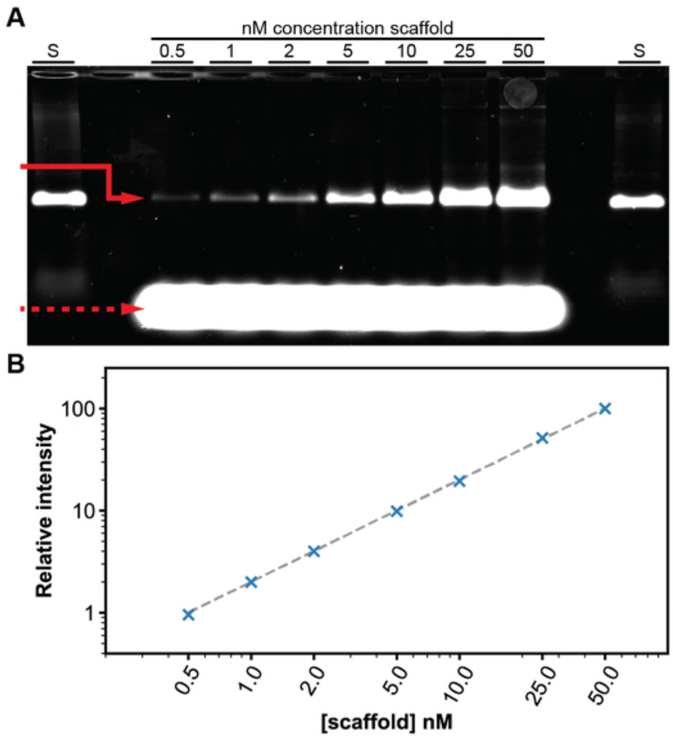

**Figure S33** Agarose gel showing the amount of single DNA-origami reference square folded varies precisely with the amount of DNA scaffold added, to serve as a benchmark for nucleation control that we wished to match with assembly of origami-slat megastructures. **A** shows the agarose gel with the solid red arrow indicating the gel band for the square, the dashed red arrow indicating the excess staple strands, and lanes 'S' containing just the scaffold. In **B**, the gel density of the solid red gel band with respect to the 50 nM condition is plotted. The dotted line is a linear fitting of the data, indicating an almost perfect stoichiometric relationship between the amount of reference square folded versus scaffold added. The concentration of scaffold was varied from 0.5–50 nM with the staple strands maintained at 400 nM per strand. This reference square folded as shown above and was also quantified by direct counting TEM images (see Fig. 5Ci).

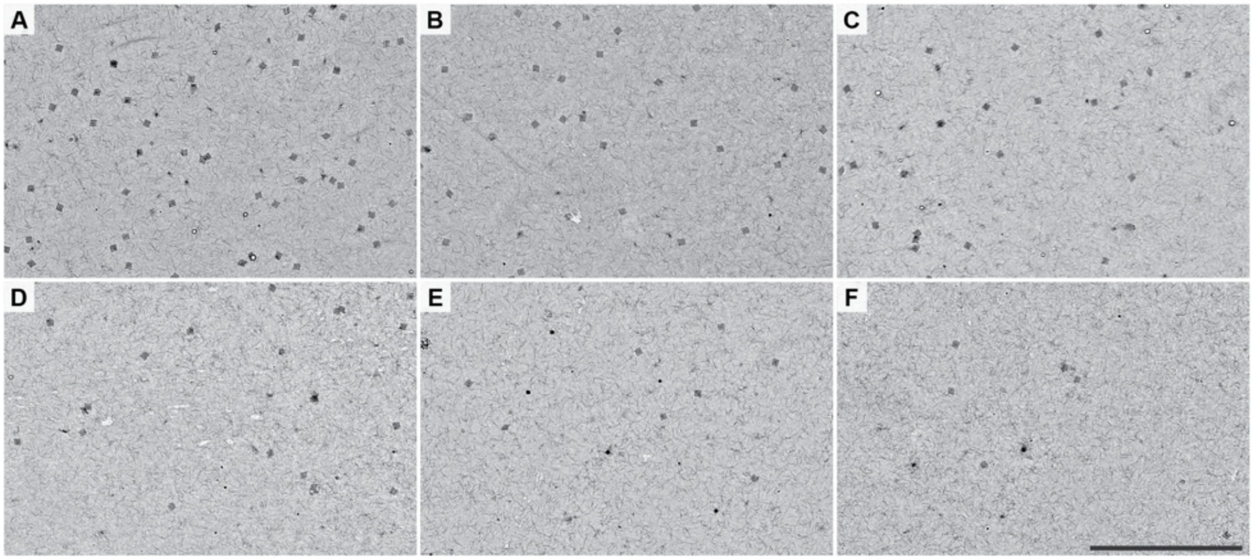

**Figure S34** Representative low-magnification TEM images of the 64-slat square versus different concentrations of seed. The concentrations of seed in **A–F** were 6, 4, 2, 1, 0.5, and 0.25 nM, respectively. There is a roughly linear relationship between the mean number of squares counted versus the amount of seed added, as plotted in the leftward panel of Fig. 5Cii. This approach of direct counting from ten TEM images to show control of assembly was also used for the reference origami square and other finite and periodic megastructures, as shown in Fig. 5C. The scale bar (which applies to all images *A–F*) is 10 μm.

**Fig. S35–Fig. S40: Standard curve of ribbons, melt temperature of v16 7-nt ribbons, temperature characterization and melt temperature of v8 7-nt, v16/v8 6-nt, and v16/v8 8-nt ribbons, growth versus time for 6-nt and 8-nt v16 ribbons**

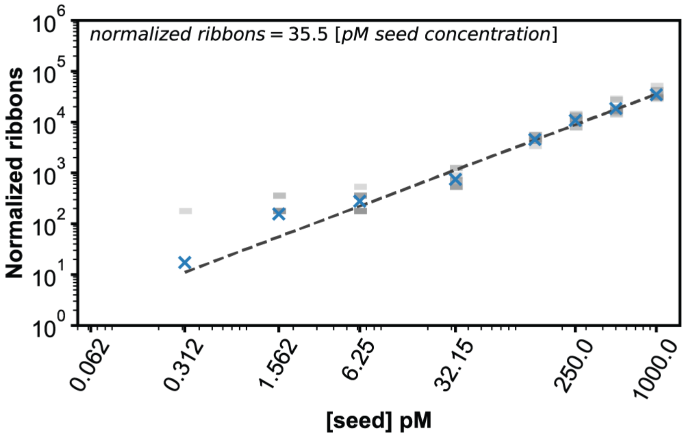

**Figure S35** Standard curve of the ribbons as derived from the experiment shown in the leftward panel of Fig. 5Ci. Ribbons were assembled in reactions where the seed was added in different amounts to a constant concentration of slats. The number of ribbons was counted in ten low-magnification (i.e. 400x) TEM images and the number of ribbons was normalized with respect to the factor that the sample was diluted (i.e. multiply by 250) and to a 100 µm × 100 µm area (i.e. divide by ~1.44, which is the number of 100 µm × 100 µm regions covered in a 400x image area with our microscope setup). This standard curve shows a ~0.3 pM limit-of-detection, where a single ribbon was observed in the ten images. We also note that we saw no ribbons in either the sample with no seed or the sample where ~0.062 pM seed was added. The blue 'x' markers indicate the mean number of ribbons counted per seed concentration. The dotted line and upper left formula is the linear fitting of the data, where the y-intercept was forced through zero to account for how no ribbons were observed when no seed was added. This equation was rearranged to compute the pM amount of spontaneously formed ribbons in Fig. 5D and Fig. S38, with the meaning of the white versus red data points versus the standard curve explained in Fig. S36.

| Marker style | x (pM spontaneous nucleation) | Relative unseeded background (vs. 0.5 nM seed) |
|---|---|---|
| □ | $x < 0.3$ | < 0.06% |
| ▲ | $0.3 \leq x < 5$ | < 0.99% |
| ● | $5 \leq x < 50$ | < 9.09% |
| ■ | $50 \leq x < 500$ | < 50% |

**Figure S36** Table of markers used in Fig. 5D and Fig. S38 to denote spontaneous nucleation of ribbons as counted in the control reactions with no seed. There were fewer pM spontaneously nucleated ribbons counted in the average of ten TEM images of unseeded control reactions, as compared to the standard curve of ribbons in the leftmost panel in Fig. 5Ciii. Each seeded reaction was initiated with 500 pM seed, with the assumed relative proportion ribbons that would have been unseeded background for each temperature condition as listed for each marker. We posit that the relative background of spontaneously nucleated ribbons is especially low for up to and including the red triangle data point, and would be of limited concern for routine nanofabrication of DNA megastructures, though the requirement of nucleation stringency might vary depending on the end application of the crisscross assembly. For instance, one could envision using crisscross growth to amplify some biomarker signal that might require lower background nucleation yet, which might be satisfied using reaction conditions with the white square marker.

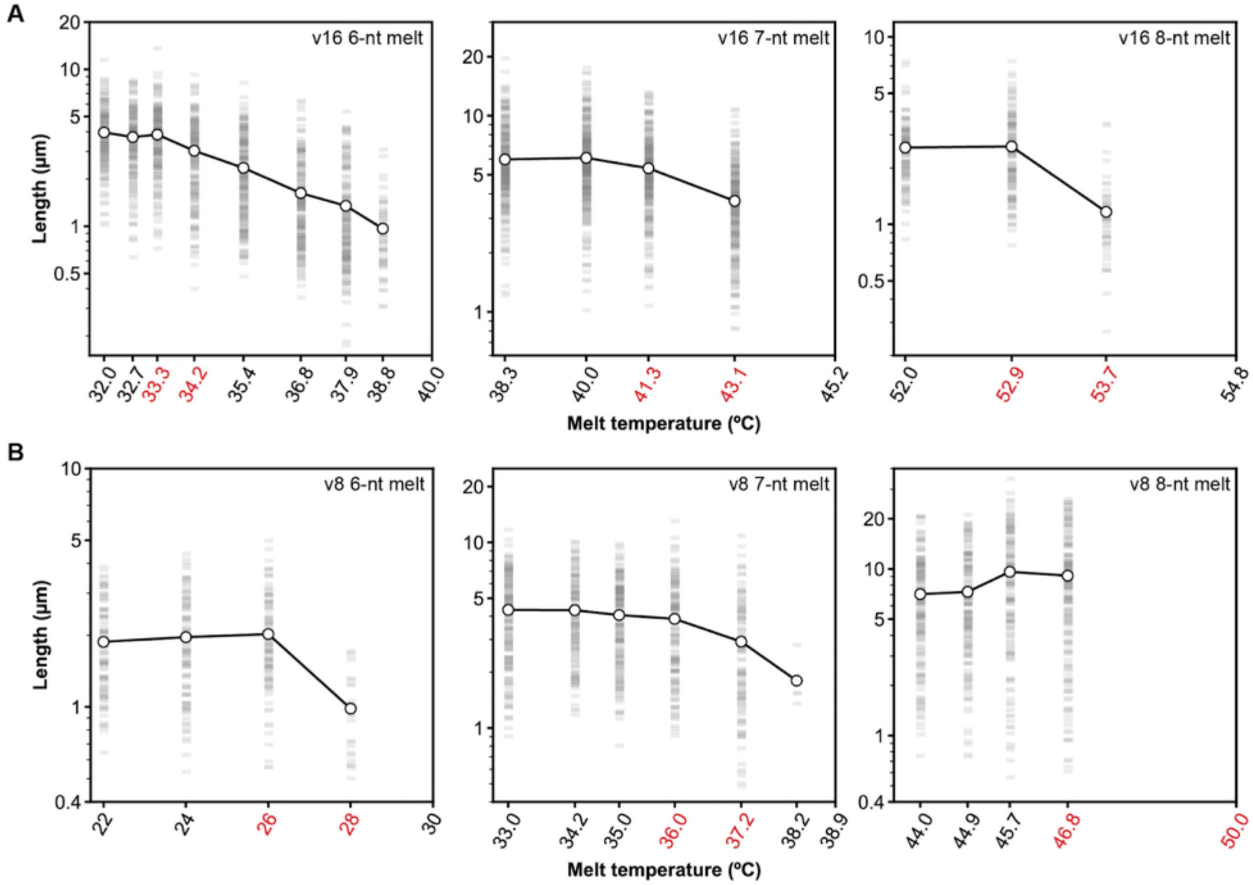

**Figure S37** Approximate melting temperatures of the v16 (**A**) and v8 (**B**) staggered ribbons using 6-, 7-, and 8-nt binding sites are within the red temperature bounds in the above plots. Ribbons were grown isothermally for ~12–16 hours at a temperature favoring seeded growth of long ribbons, at which point the temperature was increased across a gradient of possible melt temperatures for 3–4 hours. Samples were collected on TEM grids and the lengths of ribbons were recorded at the putative melt temperatures. The highest temperature tick on the plots is where no ribbons remained, and is hence above the melting temperature. Some range of temperatures below this upper bound was where the ribbons started to fall apart slowly. The shrinkage of the ribbons was typically most noticeable as observed at the second highest temperature tested, to the extent where sometimes only a small number of ribbons could be observed in the sample. The lower and upper temperature bounds containing this melt transition temperature are highlighted in red as above. N ribbons were measured for the conditions tested:

- v16 6-nt: $N_{32°C}$ = 172, $N_{32.7°C}$ = 189, $N_{33.3°C}$ = 205, $N_{34.2°C}$ = 184, $N_{35.4°C}$ = 216, $N_{36.8°C}$ = 204, $N_{37.9°C}$ = 180, $N_{38.8°C}$ = 56

- v16 7-nt: $N_{38.3°C}$ = 297, $N_{40°C}$ = 337, $N_{41.3°C}$ = 305, $N_{43.1°C}$ = 283
- v16 8-nt: $N_{52°C}$ = 124, $N_{52.9°C}$ = 141, $N_{53.7°C}$ = 51
- v8 6-nt: $N_{22°C}$ = 97, $N_{24°C}$ = 101, $N_{26°C}$ = 102, $N_{28°C}$ = 28
- v8 7-nt: $N_{33°C}$ = 174, $N_{34.2°C}$ = 148, $N_{35°C}$ = 183, $N_{36°C}$ = 146, $N_{37.2°C}$ = 114, $N_{38.2°C}$ = 4
- v8 8-nt: $N_{44°C}$ = 183, $N_{44.9°C}$ = 166, $N_{45.7°C}$ = 151, $N_{46.8°C}$ = 177

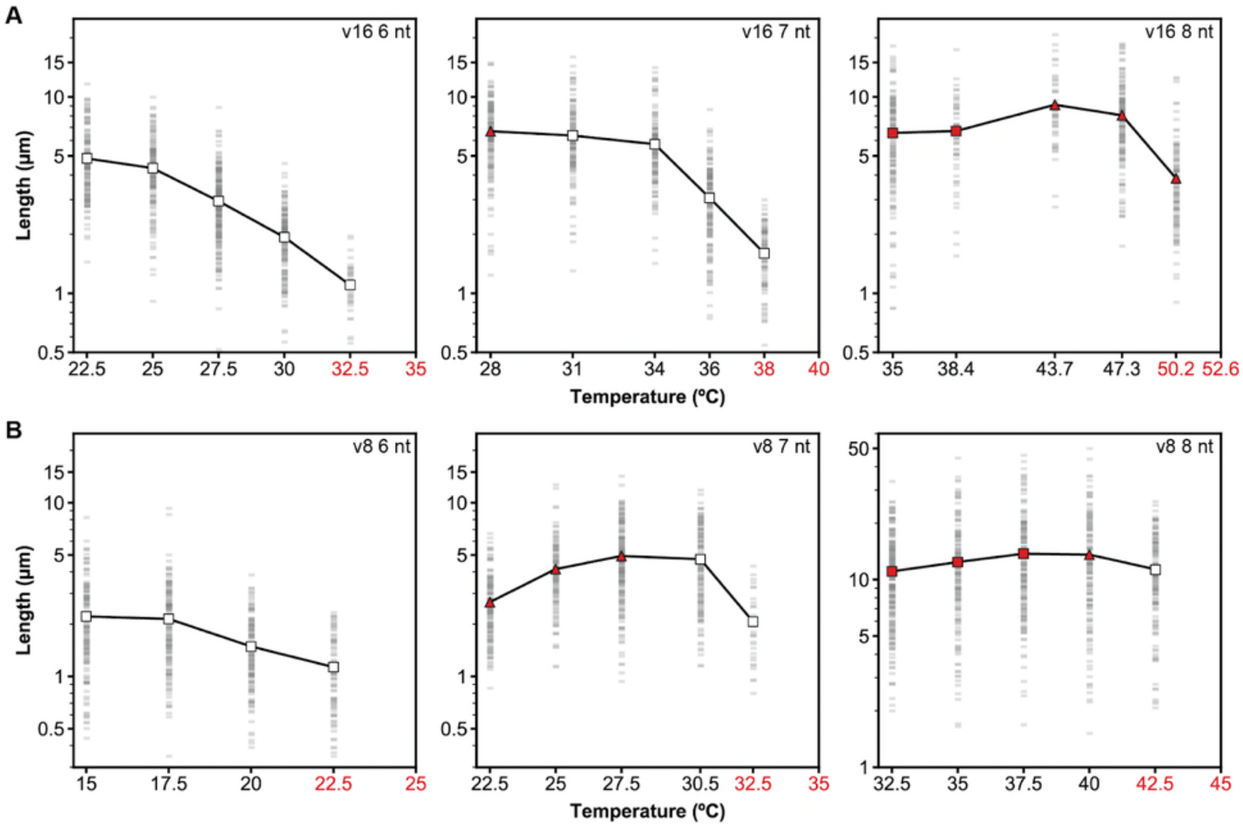

**Figure S38** Length of the ribbons versus growth temperature for seeded v16 (**A**) or v8 (**B**) ribbons using either 6-, 7-, or 8-nt binding sites. The v16 7-nt middle plot is the same data as from the leftmost plot in Fig. 5D and is replicated for easy comparison to the other designs. The x-axis ticks shown in red are the growth temperatures flanking the reversible temperature for slat binding to a ribbon end, where the seeded ribbons were first observed to be short and then entirely not observed. The longest seeded ribbons with the fastest growth were generally observed at lower growth temperatures several degrees below reversibility. The shape and color of the mean-length data points represent the number of spontaneously nucleated ribbons as counted in control reactions where no seed was added. White square points are where no measurable ribbons were observed and were below the 0.3 pM ribbon detection limit, and red points represent some amount of measurable unseeded growth. Details regarding the degradations of these red data points are explained in Fig. S36. Further discussion and consideration of the above results are in Supplementary Text 4 and Fig. S39. Each seeded reaction was mixed together with 20 nM per slat, 0.5 nM of seed, and 15 mM $Mg^{2+}$ and incubated for 4 hours at a temperature above the ribbon melt temperature where binding of slats to the seed was favored. Subsequently, the reaction was incubated isothermally for ~16 hours at the temperature as indicated on the x-axis, at which point was diluted and with images of the ribbons collected by TEM. Each faint gray box is the length measurement of a single ribbon, with the large data point representing the mean length. N ribbons were measured for the conditions tested:

- v16 6-nt: $N_{22.5°C}$ = 156, $N_{25°C}$ = 154, $N_{27.5°C}$ = 234, $N_{30°C}$ = 162, $N_{32.5°C}$ = 38
- v16 7-nt: $N_{28°C}$ = 149, $N_{31°C}$ = 139, $N_{34°C}$ = 133, $N_{36°C}$ = 170, $N_{38°C}$ = 91
- v16 8-nt: $N_{35°C}$ = 144, $N_{38.4°C}$ = 51, $N_{43.7°C}$ = 46, $N_{47.3°C}$ = 154, $N_{50.2°C}$ = 120
- v8 6-nt: $N_{15°C}$ = 145, $N_{17.5°C}$ = 166, $N_{20°C}$ = 137, $N_{22.5°C}$ = 81
- v8 7-nt: $N_{22.5°C}$ = 141, $N_{25°C}$ = 128, $N_{27.5°C}$ = 201, $N_{30°C}$ = 140, $N_{32.5°C}$ = 34
- v8 8-nt: $N_{32.5°C}$ = 206, $N_{35°C}$ = 150, $N_{37.5°C}$ = 171, $N_{40°C}$ = 141, $N_{42.5°C}$ = 150

A

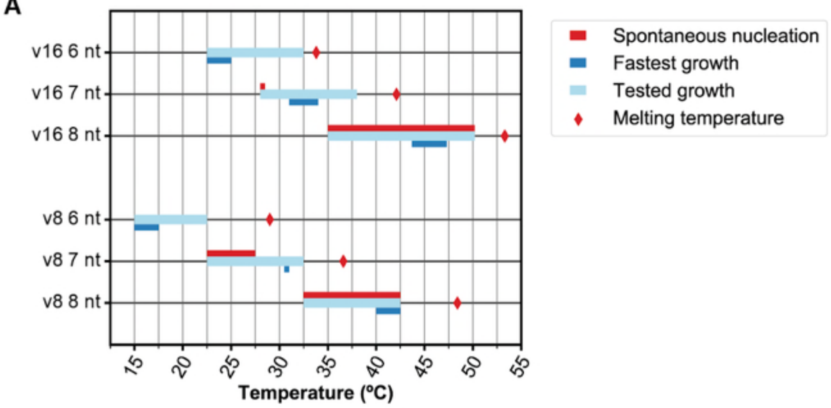

B

| design | "optimal" °C | reversible °C | melting °C |
|---|---|---|---|
| v16 6 nt | ~22.5–25 | >32.5 | ~33.8 |
| v8 6 nt | ~15–17.5 | >22.5 | ~29 |
| **v16 7 nt** | **~31–34** | **>38** | **~42.1** |
| v8 7 nt | ~30.5 | >32.5 | ~36.6 |
| v16 8 nt | ~43.7–47.3 | >50.2 | ~53.3 |
| v8 8 nt | ~40–42.5 | >42.5 | ~48.4 |

**Figure S39** Growth and melt temperatures as determined for various ribbon designs are plotted in **A**, with the plotted data numerically shown in table **B**. The light-blue lines indicate the different temperatures tested in Fig. S38, where growth of ribbons was observed when the seed was added. The highest light-blue points are the temperatures where ribbon growth drastically slowed, and approached the reversible temperature for slats to bind either 16 (for v16) or 8 (for v8) binding handles on a ribbon end. The red diamonds are the average temperatures where previously assembled ribbons were observed to fall apart (i.e. the approximate melt temperature), as determined in Fig. S37. The lowest light-blue points are the lowest growth temperature as tested in Fig. S38, though it would likely be possible to carry out seeded growth at lower temperatures yet. The dark-blue points were designated as the "optimal" growth temperatures favoring fast, seed-driven growth of the ribbons as determined from Fig. S38. We discuss the selection of these temperature optimums in more depth in Supplementary Text 4.1. The red points were where measurable spontaneous nucleation was observed in Fig. S38 that was some amount greater than the 0.3 pM ribbon detection limit.

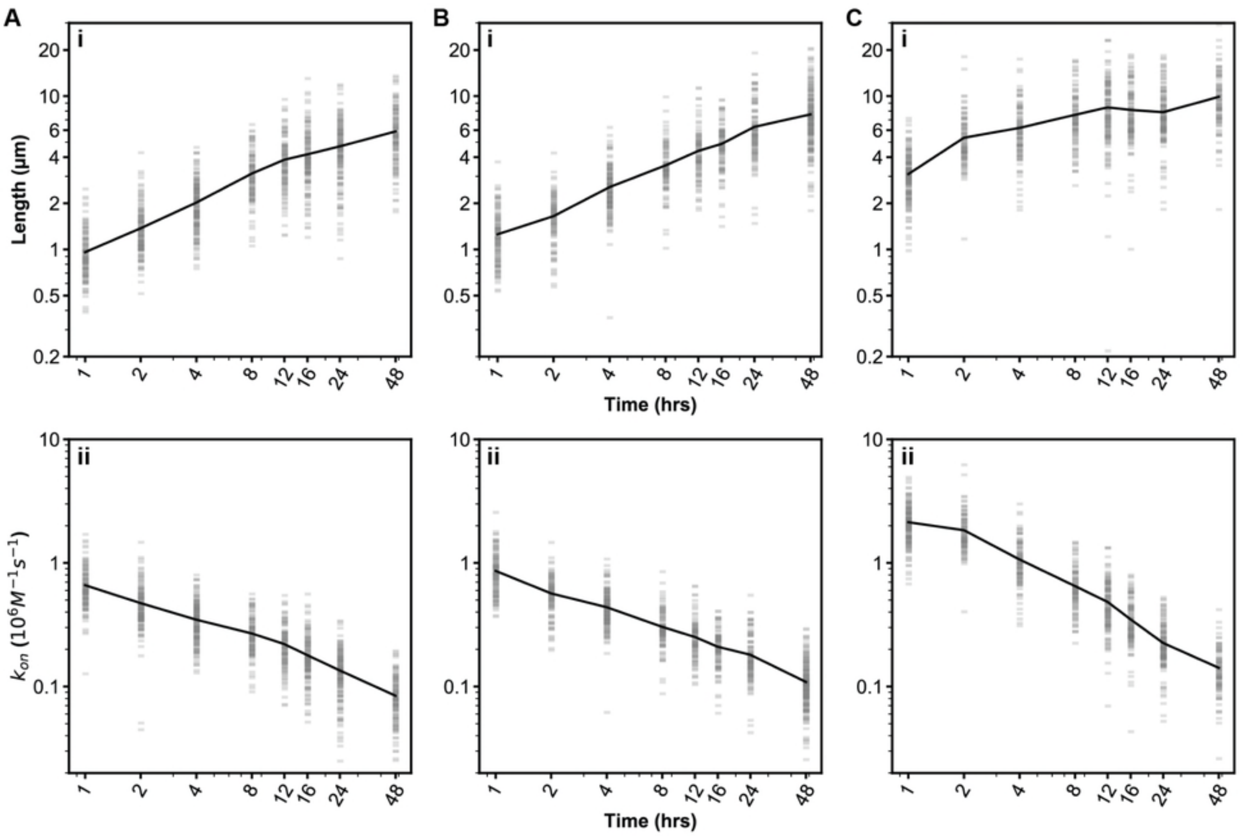

**Figure S40** Length of the ribbons (see **i**) and extrapolated kinetics (see **ii**) for v16 6-, 7-, and 8-nt ribbons versus time in **A**, **B**, and **C** respectively. Panel Bi is replicated from Fig. 5E for clearer comparison of the three designs. Each faint gray bar represents the length measurement of a single ribbon, with the mean lengths indicated by the line. The mean lengths for the ribbons after 1 hour of growth were ~1, ~1.3, and ~3.1 µm, and after 16 hours of growth were ~4.2, ~4.9, and ~8.1 µm for the 6-, 7-, and 8-nt designs, respectively. Explanation for how the mean lengths were extrapolated into $k_{on}$ rates are discussed in Supplementary Text 5. The rates of growth were increasingly faster for the 6-, 7-, and 8-nt designs. The observed average forward rate of assembly after one hour of growth was ~0.66*$10^6$, ~0.86*$10^6$, and ~2.13*$10^6$ $M^{-1}s^{-1}$ for the 6-, 7-, and 8-nt designs. The average rates were observed to gradually decline over time, such as ~0.18*$10^6$, ~0.21*$10^6$, and ~0.35*$10^6$ $M^{-1}s^{-1}$ for the 6-, 7-, and 8-nt designs after 16 hours of growth. We hypothesize that this decline was because of temporary stalling and permanent halting of growth due to accumulation of errors with missing handles on the growing fronts of ribbons, or else due to depletion of the free slats which may have caused inaccuracy of the pseudo-first order kinetic assumptions as explained in Supplementary Text 5. All assembly reactions had 20 nM per slat, 0.5 nM of seed, and 15 mM $Mg^{2+}$ and were conducted at growth temperatures where spontaneous nucleation was limited. N ribbons were measured for the conditions tested:

- v16 6-nt: $N_{1\ hr}$ = 130, $N_{2\ hr}$ = 141, $N_{4\ hr}$ = 179, $N_{8\ hr}$ = 114, $N_{12\ hr}$ = 119, $N_{16\ hr}$ = 146, $N_{24\ hr}$ = 177, $N_{48\ hr}$ = 128
- v16 7-nt: $N_{1\ hr}$ = 131, $N_{2\ hr}$ = 107, $N_{4\ hr}$ = 158, $N_{8\ hr}$ = 107, $N_{12\ hr}$ = 106, $N_{16\ hr}$ = 106, $N_{24\ hr}$ = 118, $N_{48\ hr}$ = 204
- v16 8-nt: $N_{1\ hr}$ = 188, $N_{2\ hr}$ = 108, $N_{4\ hr}$ = 112, $N_{8\ hr}$ = 108, $N_{12\ hr}$ = 143, $N_{16\ hr}$ = 125, $N_{24\ hr}$ = 148, $N_{48\ hr}$ = 100

**Fig. S41–Fig. S45: Hamming-distance analysis of 1D ribbon growth, mechanistic testing for growth changes to 1D ribbons vs. kinetic trapping of slats resulting from Hamming distance, and optimized Hamming distances of all the megastructure designs tested in this work**

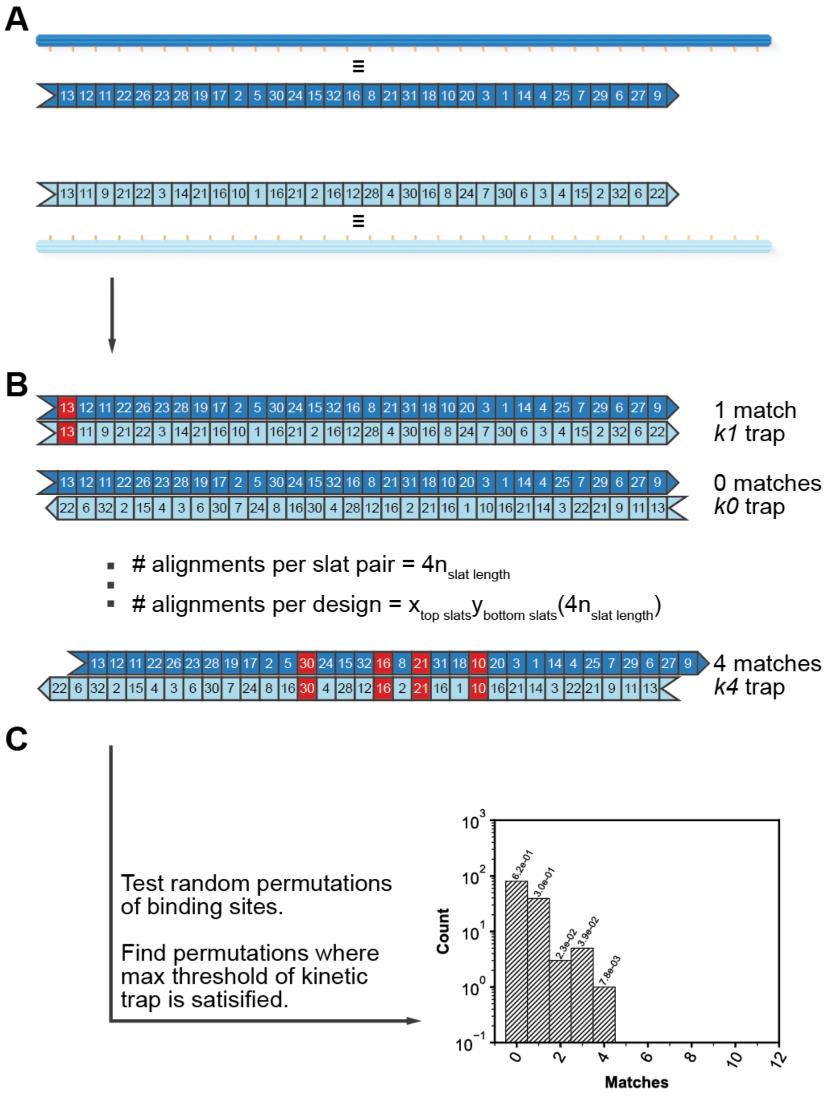

**Figure S41** Schematic showing analysis of Hamming distances between a single pair of slats. In **A**, a pair of example slats are extracted from some random permutation of binding sites for a megastructure. Rendered images of 6HB slats are drawn proximally to the numeric representation, where each slat is abstracted as a one-dimensional list which is $n_{slat\ length}$ long (i.e. $n = 32$ for the 6HB slat). In **B**, the Hamming distance between the various forward and reverse alignments is measured to determine the number of matching, complementary handles between the pair of slats (where matches in the sample alignments are in red). There are $4n_{slat\ length}$ alignments per slat pair, and $x*y$ slat pairs in a megastructure composed of $x$ top-slats and $y$ bottom-slats. For the example slat pair, the maximum number of matches is four and is designated as a kinetic trap of strength *k4*. In **C**, the strength of the total 128 alignments for the example slat pair are shown in a histogram, with the max-strength kinetic trap shown in the rightmost bin.

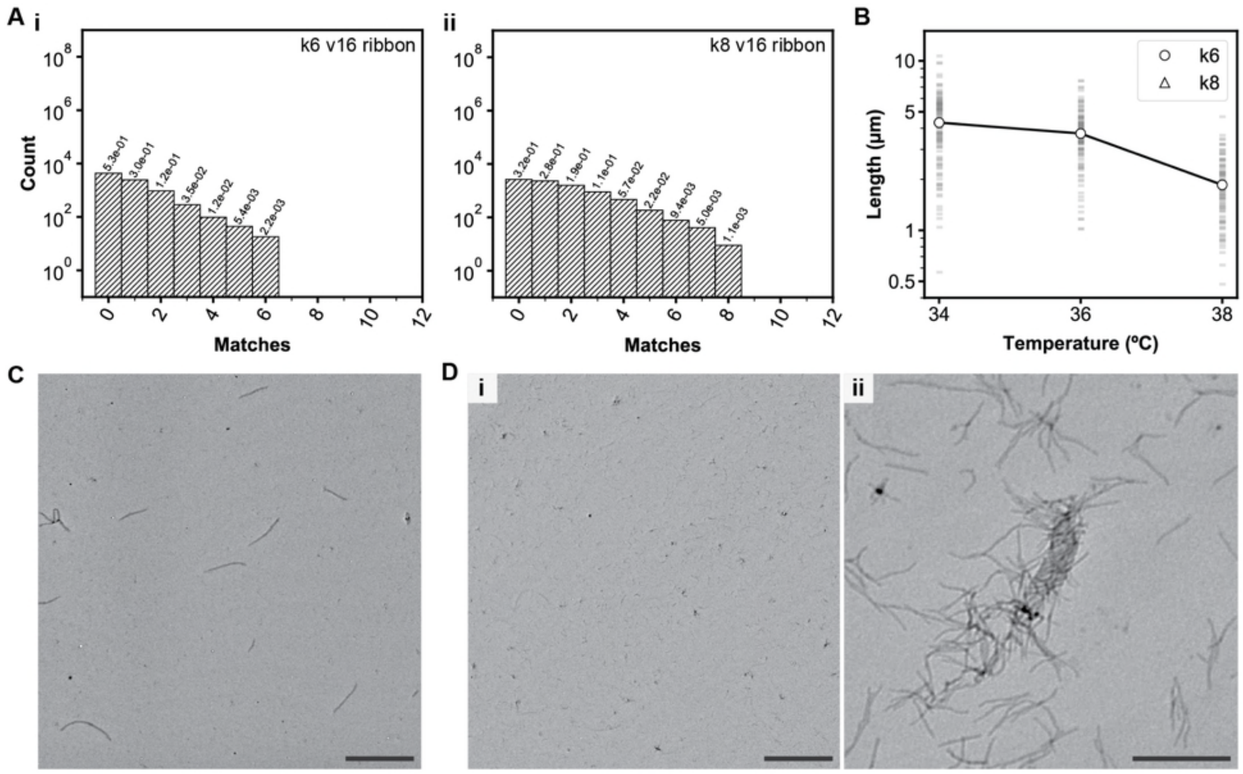

**Figure S42** Increasing the *k* kinetic-trap strength (i.e. the number of matching complementary binding sites between pairs of slats resulting from smaller Hamming distances) nearly stops the growth of v16 7-nt ribbons. The number of kinetic traps between one layer of slats and the other perpendicular layer is shown for two ribbon designs in **Ai–ii**. There are maximally six or eight matching binding sites between the slats (i.e. *k6* and *k8* strength kinetic traps) in each of the designs, respectively. The relative fraction of each strength trap of the total interactions measured is shown above each histogram bar. In **B**, the length of ribbons as measured by TEM after overnight growth of the *k6* and *k8* designs at various temperatures is plotted. We note that no appreciable growth of the *k8* ribbons was measured at any of the temperatures tested. Each faint gray box represents the length measurement of a single ribbon, with the circular and triangular white points showing the mean. Low-magnification TEM image of the *k6* growth is in **C**, versus the unobservable growth of the *k8* design in **Di** and one of the infrequently observed short *k8* ribbons in **Dii**. Isothermal growth was conducted for 16 hours using 15 mM $Mg^{2+}$, with either ribbon design using 4x sequence symmetry with binding sites as selected from the 2048-strand library. Scale bars are 10 µm in *C* and *Di*, and 1 µm in *Dii*. N ribbons were measured for the conditions tested: $N_{k6\ 34°C}$ = 118, $N_{k6\ 36°C}$ = 108, $N_{k6\ 38°C}$ = 94.

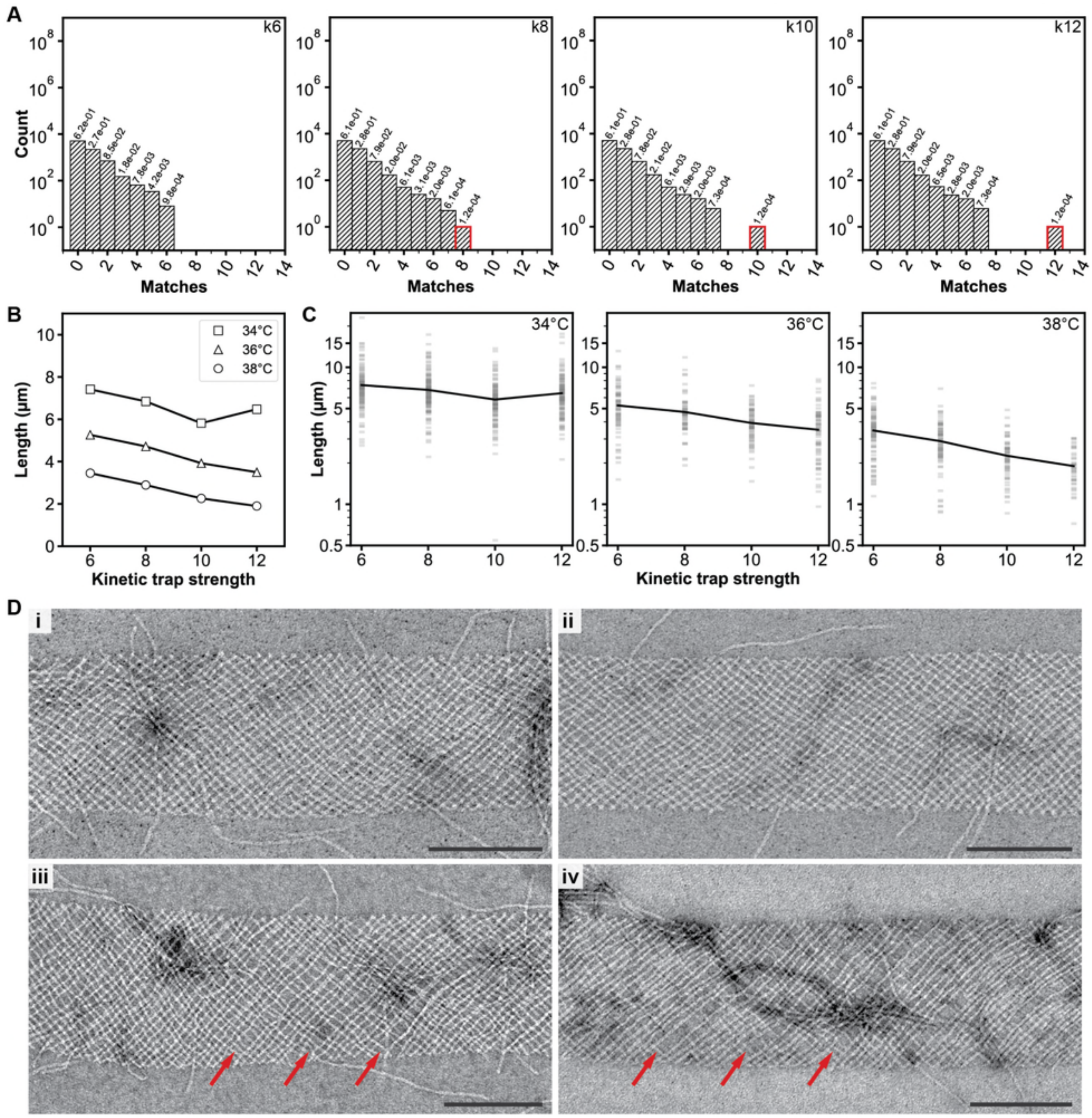

**Figure S43** Introducing a single kinetic trap between a single pair of slats of max strength *k* progressively slows the growth of v16 7-nt ribbons as *k* is increased. Histograms in **A** show the number of kinetic traps between one layer of slats and the other perpendicular layer in the different ribbon designs tested. The *k6* sequence permutations is a control with the maximal Hamming distance that we attained for a 4x symmetry ribbon using the 2048-strand library. The *k8* permutation is distinct from the *k6* control, with the *k10* and *k12* that were generated from small manipulations of the *k8* design. Each of these latter three designs have only a single kinetic trap of either *k8*, *k10*, or *k12* strength. The relative fraction that each kinetic trap comprises of the total interactions measured is written above each bar. **B** shows the mean length of the ribbons at different temperatures versus the different max-strength kinetic traps. In general, the ribbons were shorter at a given temperature when the kinetic trap was stronger with a larger *k* trap where more matching binding sites entrapped one pair of slats. **C** shows the raw measured-length data of each design for each temperature tested. In **D**, close-up negative-stain TEM images show segments of a ribbon for *k6*, *k8*, *k10*, and *k12* traps in **i–iv**, respectively. Red arrows point to periodic missing slats in one layer of the ribbon which appeared every eighth slat. Scale bars are 200 nm. The length of N ribbons were for each of the conditions tested above:

- At 34°C: $N_{k6}$ = 124, $N_{k8}$ = 138, $N_{k10}$ = 122, $N_{k12}$ = 129.
- At 36°C: $N_{k6}$ = 77, $N_{k8}$ = 61, $N_{k10}$ = 78, $N_{k12}$ = 72.
- At 38°C: $N_{k6}$ = 91, $N_{k8}$ = 81, $N_{k10}$ = 61, $N_{k12}$ = 41.

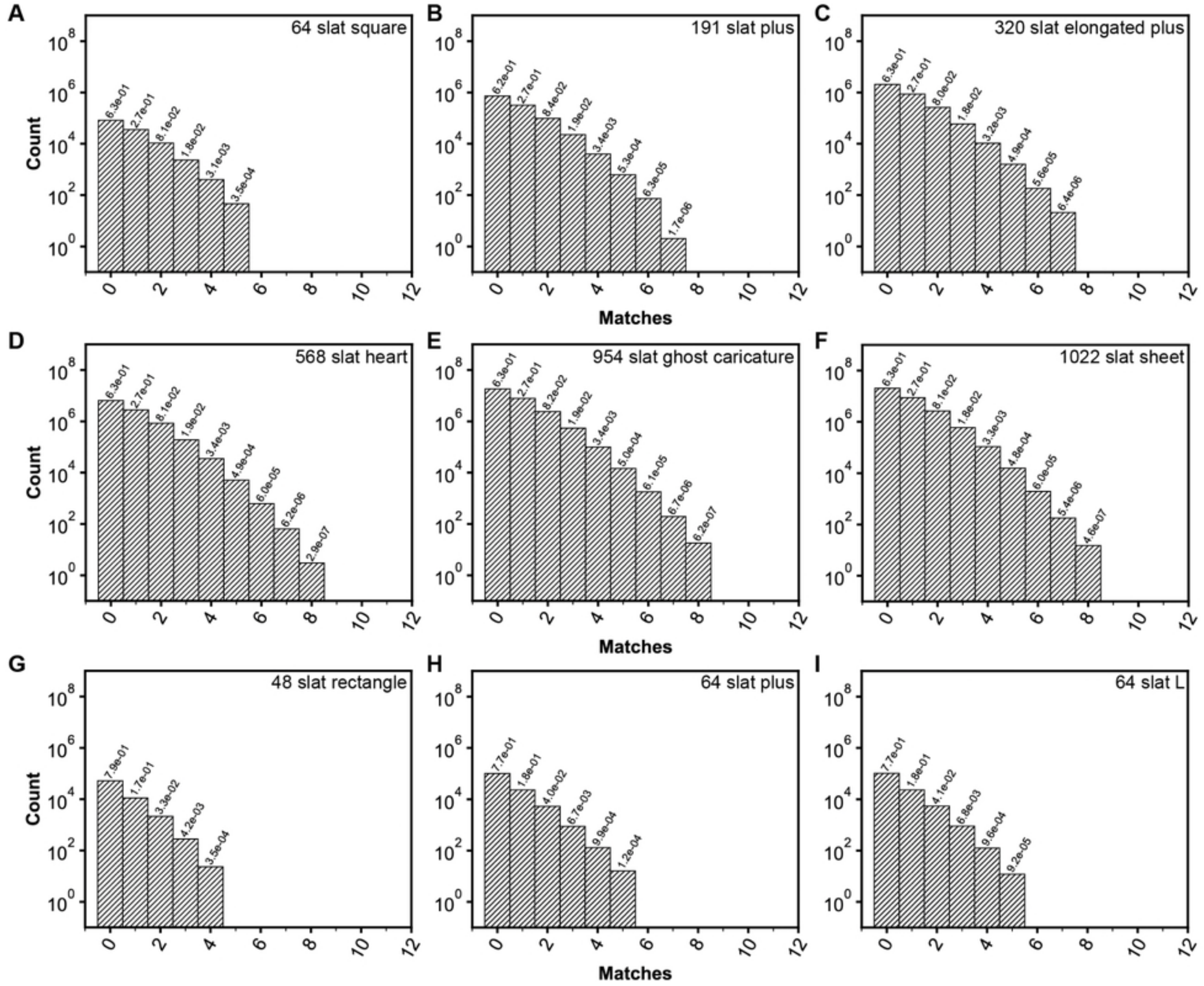

**Figure S44** The multiplicity for each possible number of matching, complementary binding sites for each of the slats from one layer with respect to each of the slats in the other layer are shown as histograms, for the optimized finite megastructure designs as tested in Fig. 3. The relative fraction that each match comprises of the total interactions is written above each bin. We minimized the maximum number of matches for each design as discussed in Method 2 and Supplementary Text 6.4.

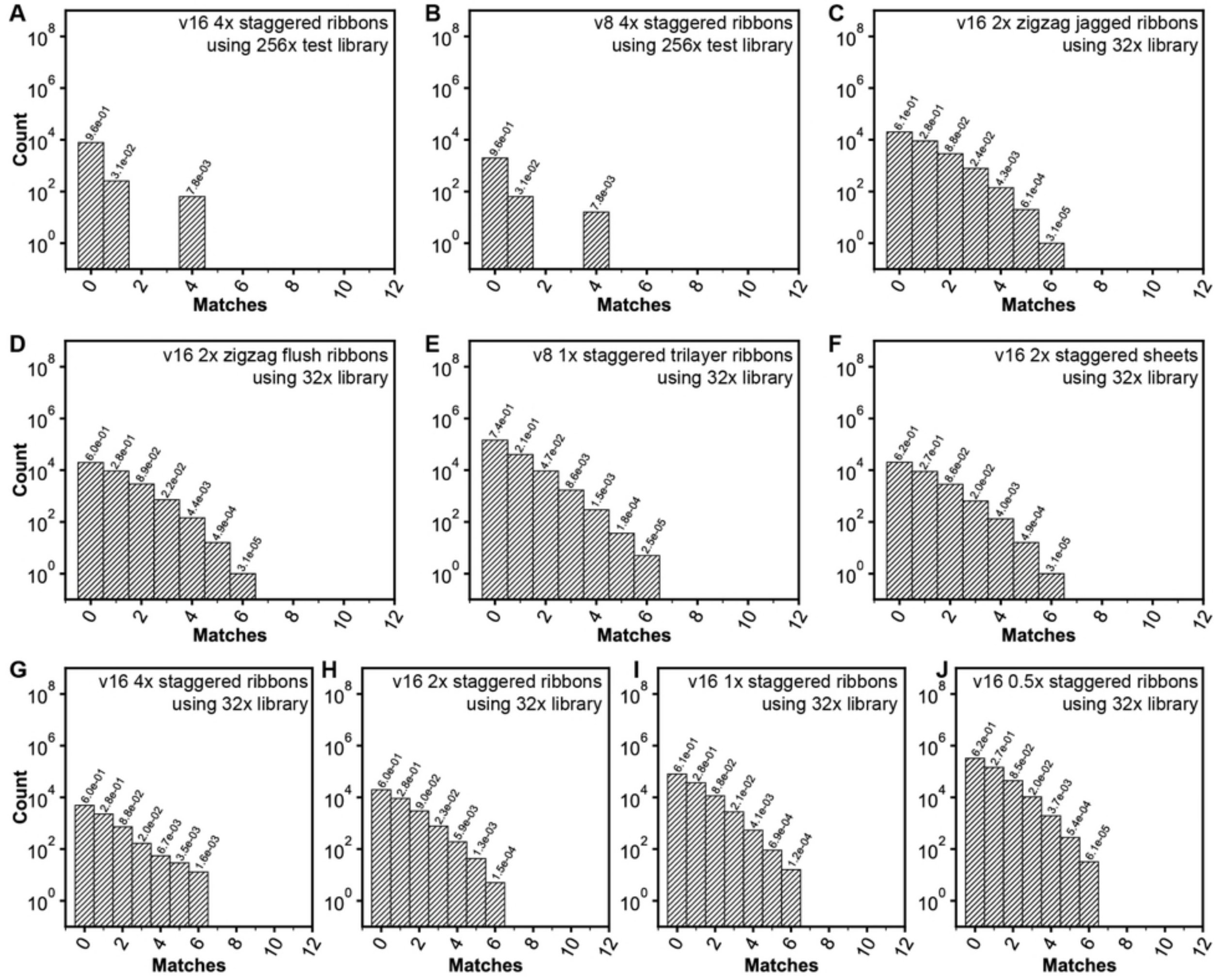

**Figure S45** The multiplicity for each possible number of matching, complementary binding sites for each of the slats from one layer with respect to each of the slats in the other layer are shown as histograms, for the optimized periodic megastructures in this paper. The relative fraction that each match comprises of the total interactions is written above each bin. We minimized the maximum number of matches for each design as discussed in Method 2 and Supplementary Text 6.3. The ribbon designs in **A–B** have maximally *k4* traps, with permutations generated from 256 different 6-, 7-, or 8-nt handles from a 512-strand library. All other periodic designs in **C–J** have maximally *k6* traps, with permutations generated from 32 different

7-nt handles in the 2048-strand library. Design *A* is for the 4x symmetry v16 staggered ribbon as used in Fig. 3Ai, Fig. S18A, Fig. S19A–B design 'S', Fig. S27, Fig. S35, Fig. 5Ciii ribbons, and v16 ribbon characterization in Fig. 5D–E and Fig. S37–Fig. S40; design *B* is for the 4x symmetry v8 staggered ribbon as used in Fig. S17Aii, Fig. S18B, and v8 ribbon characterization in Fig. S37–Fig. S38; designs *C*–*D* are for the 2x symmetry v16 non-staggered ribbons as used in Fig. 3Aii–iii, Fig. S18C–D, and Fig. S19A–B designs 'J' and 'F'; design *E* is for all tri-layer designs; design *F* was used for any staggered sheet; designs *G*–*J* are for the ribbons in Fig. S21.

### Table S1–Table S8: Core-strand sequences for 6HB, 12HB, seed, and single origami reference square; Sequences of the 6-, 7-, 8-, 9-, and 10-nt handles; Nanocube strand sequences

| 6HB universal core staples |
| --- |
| ACCATCACCCAAACTATTAAAGAACGTGGAGAGAGAGTTGCA<br>GCAAGCGGTCCACGGTTTTTCTTTTCACCATAAAGCCTGGGG<br>TGCCTAATGAGTGGTGTGAAATTGTTATCCCTGCGGCCAGAA<br>TGCGGCGGGCCGTTGCCCCCTGCATCAGACGCGCGGTTGCGG<br>TATGAGCCGGGTCGCTGGAGGTGTCCAGCACGTCGCTGGCAG<br>CCTCCGGCCAGAGAGGGTAAAGTTAAACGAGGTGAAGGGATA<br>GCTCTCACGGAAAGAAACGTACAGCGCCATCCAGCTGGCGAA<br>AGGGGGATGTGCTCCATTCAGGCTGCGCAAGGGATAGGTCAC<br>GTTGGTGTAGATGGTGAGCGAGTAACAACCTATAAGCAAATA<br>TTTAAATTGTAAACATGTCAATCATATGTAAATCACCATCAA<br>TATGATATTCAACAATGCCTGAGTAATGTGAAGAATTAGCAA<br>AATTAAGCAATAAGGCATCAATTCTACTAACTAAAGTACGGT<br>GTCTGGAAGTTTCTGGCTTAGAGCTTAATTCATCAAAAAGAT<br>TAAGAGGAAGCCCCCATAAATCAAAAATCACAAAATAGCGAG<br>AGGCTTTTGCAAAACGAGGCATAGTAAGAGGCTCATTATACC<br>AGTCAGGACGTTGGAGATGGTTTAATTTCAAGACCAGGCGCA<br>TAGGCTGGCTGACGGTCAATCATAAGGGAAGCAAAAGAATAC<br>ACTAAAACACTCACCATTAAACGGGTAAAAATAACCGATATA<br>TTCGGTCGCTGAGTTCTTAAACAGCTTGATCAACTTTCAACA<br>GTTTCAGCGGAGTAAGTTTTGTCGTCTTTCGAACCGCCACCC<br>TCAGAGCCACCACTAGGTGTATCACCGTACGCCCCCTGCCTA<br>TTTCGGAACCTATGGTAATAAGTTTTAACGGCATTGACAGGA<br>GGTTGAGGCAGGTCACCCTCAGAGCCACCATTGCCTTTAGCG<br>TCAGACTGTAGCGAAACGTCACCAATGAAAAAAGGGCGACAT<br>TCAACCGATTGAGATAAGTTTATTTTGTCATACCAGAAGGAA<br>ACCGAGGAAACGCGCAATAGCTATCTTACCGAGAATAACATA<br>AAAACAGGGAAGCTCCAAATAAGAAACGATACTTGCGGGAGG<br>TTTTGAAGCCTTACAGATATAGAAGGCTTATTACGAGCATGT<br>AGAAACCAATCAAGCGCCTGTTTATCAACAAATTGAGAATCG<br>CCATATTTAACAATATCATATGCGTTATACCAAAGAACGCGA<br>GAAAACTTTTTCACCGGCTTAGGTTGGGTTAGTGAATAACCTTGCT<br>AACAAGAGTCCATCAAGTTTTTTGGCCGTAAAG<br>CACTAAATCGGAAGATAGGGTTGAGTGTTGCGAAAATCCTGT<br>TTGATGGTGGTTAACGCGCGGGGAGAGGCGCGTTGCGCTCAC<br>TGCCCGCTTTCCTACCGAGCTCGAATTCGGGTTTCTGCCAGC<br>ACGCGTGCCTGTACGGCATCAGATGCCGGGTAATGGGTAAAG<br>GTTTCTTTGCTCGCAGCACCGTCGGTGGTCGTGCCGGACTTG<br>TAGAACGTCAGCAATCCCGTAAAAAAAGCGGATCAAACTTAA<br>ATTTCTGCTCATAGCCGCCACGGGAACGGGTAACGCCAGGGT<br>TTTCCCAGTCACGCACCGCTTCTGGTGCCATCTGCCAGTTTG<br>AGGGGACGACGAGCGTCTGGCCTTCCTGTATTCGCATTAAAT<br>TTTTGTTAAATCGAGCAAACAAGAGAATCAATGCCGGAGAGG<br>GTAGCTATTTTTCAAGGATAAAAATTTTTTAAATCGGTTGTA<br>CCAAAAACATTATGTTTAGCTATATTTTCTTCCCAATTCTGC<br>GAACGAGTAGATACCTTTAATTGCTCCTTCGTTTTAATTCGA<br>GCTTCAAAGCGATCCCCCTCAAATGCTTTGGTAATAGTAAAA<br>TGTTTAGACTGGCCACATTCAACTAATGCAATAAAACGAACT<br>AACGGAACAACAGCCCTGACGAGAAACACTGACAAGAACCGG<br>ATATTCATTACCGCGACCTGCTCCATGTTTATACCAAGCGCG<br>AAACAAAGTACACAGAGGCTTTGAGGACTGCCGCTTTTGCGG<br>GATCGTCACCCTCTTTAATTGTATCGGTTCTAAAGGAATTGC<br>GAATAATAATTTAGCATTCCACAGACAGCCAAGCCCAATAGG<br>AACCCATGTACCATAAGTGCCGTCGAGAGTATTAAGAGGCTG<br>AGACTCCTCAAGGTAAGCGTCATACATGGATTCACAAACAAA<br>TAAATCCTCATTACCGGAACCGCCTCCCTCGGTCATAGCCCC<br>CTTATTAGCGTTAGCAAAATCACCAGTAGGACGGAAATTATT<br>CATTAAAGGTGAAAAGGTGGCAACATATAAAAAGAACTGGCA<br>TGATTAAGACTCTTGAGTTAAGCCCAATAAACTGAACACCCT<br>GAACAAAGTCAGTTTGCCAGTTACAAAATTTTTGCACCCAGC<br>TACAATTTTATCTTCATCGTAGGAATCATTATCATTCCAAGA<br>ACGGGTATTAAACAGACGACGACAATAAAGCAGAGGCATTTT<br>CGAGCCAGTAATATAAACACCGGAATCATTCATCTTCTGACC<br>TAAATTTAATGGTTTATCAAAATCATAGGTTCCCTTAGAATC<br>GGAGCCCCCTTTTTTTTTTGATTTAGAGAAAACCGTCTATCA<br>GCAAAATCCCTTATGAGAAAGGAAGGGAATTGCCCTTCACCG<br>ACCTGTCGTGCCAGGCTGGCAAGTGTAGATACGAGCCGGAAG<br>CCGTGAGCCTCCTCCGCCGCGCTTAATGGCGCGCCTGTGCAC<br>ATCCCTTACACTGGCTTTGACGAGCACGCGCTTTCGCACTCA<br>GTCTGGTCAGCAGCAGCGGGAGCTAAACAAACGCGGTCCGTT<br>GCAGTTGGGCGGTTGGAACGGTACGCCATGTGAGAGATAGAC<br>AAACGACGGCCAGTTGAGGCCACCGAGTGTGCGGGCCTCTTC<br>CCTCAGGAAGATCGCCGTTGTAGCAATAACAAACGGCGGATT<br>TTTAACCAATAGGACCTGAGTAGAAGAAAAGCCCCAAAAACA<br>ACAAAGGCTATCAGCCAGAACAATATTAGTGAGAAAGGCCGG<br>AATACTTTTGCGGGTCATGGAAATACCTCAATAAATCATACA<br>CCATTAGATACATTGATTATTTACATTGCTCAACATGTTTTA<br>GAAGCAAACTCCAATAAAAGGGACATTCATAGTCAGAAGCAA<br>AATACTGCGGAATCCTGAAAGCGTAAGAGTTTACCAGACGAC<br>GTAGAAAGATTCATGGCTATTAGTCTTTACCTTATGCGATTT<br>GTAACAAAGCTGCTCGCCATTAAAAATAAAGAGGACAGATGA<br>TGTATCATCGCCTGCAGAGGTGAGGCGGGGCACCAACCTAAA<br>AAGACAGCATCGGACACGCTGAGAGCCAATGACAACAACCAT<br>GAAAATCTCCAAAACTTGCTGAACCTCATTTCTGTATGGGAT |

| 6HB top staples (h5) |
| --- |
| 1; CCTGGCCCTCTCCAACGTCAAAGGGCGACTTGACGGGGAAAG<br>2; CATAAAGTGGTGAGACGGGCAACAGCTGAGAAAGCGAAAGGA<br>3; TCTGTGGTGGCTCACAATTCCACACAACCGGTCACGCTGCGC<br>4; ATCCGCCGGGATCCAGCGCAGTGTCACTCGCCGCTACAGGGC<br>5; TTTTCGTCTTCAGCGGGGTCATTGCAGGTATAACGTGCTTTC<br>6; TTTCTCCGTTGCTGATTGCCGTTCCGGCAGGAGGCCGATTAA<br>7; GCTATTACGGTTTACCAGTCCCGGAATTGAATCCTGAGAAGT<br>8; GACCGTAATCTGTTGGGAAGGGCGATCGAAAAGAGTCTGTCC<br>9; GGAAGATTGCGTCGGATTCTCCGTGGGACTTCTTTGATTAGT<br>10; AGACAGTCACCCCGGTTGATAATCAGAACTCAAACTATCGGC<br>11; GGCAAGGCATAGGTAAAGATTCAAAAGGCCGCCAGCCATTGC<br>12; AATATGCAATAGTAGTAGCATTAACATCACATTTTGACGCTC<br>13; AGCGGATTGGCTGAATATAATGCTGTAGGCAGATTCACCAGT<br>14; GATAAAAACGGTCTTTACCCTGACTATTTGGCCAACAGAGAT<br>15; TAAGAACTGCAACACTATCATAACCCTCATACGTGGCACAGA<br>16; ACGGTGTACACTTTAATCATTGTGAATTAATGCGCGAACTGA<br>17; ACGAAAGAGCCGAACTGACCAACTTTGACCGAACGAACCACC<br>18; CGCCCACGCTACGTAATGCCACTACGAATCAGTATTAACACC<br>19; TTTGCTAAAACCGATAGTTGCGCCGACAGCAGCAAATGAAAA<br>20; CCACCCTCACAGACGTTAGTAAATGAATAATATCAAACCCTC<br>21; ACAGTTAATTCAGGAGGTTTAGTACCGCACAGTTGAAAGGAA<br>22; CCGCCGCCAGGGTCAGTGCCTTGAGTAAGGAGCACTAACAAC<br>23; GAATCAAGTCCCTCAGAGCCGCCACCAGACATTTGAGGATTT<br>24; CCAAAGACACCATCGATAGCAGCACCGTACAACTCGTATTAA<br>25; GAACAAAGTCAATCAATAGAAAATTCATAAAGTTTGAGTAAC<br>26; CTTTACAGAGAAGCCCTTTTTAAGAAAACCAGAAGGAGCGGA<br>27; AACCTCCCGTTTTTGTTTAACGTCAAAAGATGGCAATTCATC<br>28; CATCCTAATTCCGGTATTCTAAGAACGCTTCTGAATAATGGA<br>29; GTAGGGCTTATAGATAAGTCCTGAACAATTTGCACGTAAAAC<br>30; ATCGCAAGAAAATTCTTACCAGTATAAAGGTTTAACGTCAGA<br>31; ATATATGTGATATAACTATATGTAAATGTCGGGAGAAACAAT<br>32; TCTGTAAATTAACAATTTCATTTTTTTAATGGAAACAAGTTACAAAATC |

| 6HB bottom staples (h2) |
| --- |
| 1; GGGCGCCAGGGTGCTGGTTTGCCCCAGCAGGTTCCAGTTTGG<br>2; TAGCTGTTTCCTAGCTAACTCACATTAATTGGTTTGCGTATT<br>3; CAGCGGTGCCGGTTTCACGGTCATACCGGGTAATCATGGTCA<br>4; AACCAGCTTACGACTGTTGCCCTGCGGCTGGTTACCTGCAGC<br>5; CTTTAGTGATGACACATCCTCATAACGGAAGCCATCCCACGC<br>6; GAAACAATCGGCAAGAGACGCAGAAACAGCCGCACAGGCGGC<br>7; AAGCGCCATTCGGCAAGGCGATTAAGTTGGATAACCTCACCG<br>8; TCAACATTAAATGGCGCATCGTAACCGTGCGGAAACCAGGCA<br>9; TCGTAAAACTAGCGTTAATATTTTGTTAAAAGCCAGCTTTCA<br>10; TATTTTAAATGCCGTTCTAGCTGATAAATTGATGAACGGTAA<br>11; AGCTGAAAAGGTAGCCTCAGAGCATAAAGCAGAACCCTCATA<br>12; CATTTTTGCGGAATTCCATATAACAGTTGAATTTGGGGCGCG<br>13; AAACGAGAATGAGAAAGACTTCAAATATCGTTGATAAGAGGT<br>14; CCAAAAGGAATTAGAAGTTTTGCCAGAGGGAAACAGTTCAGA<br>15; TAAATTGGGCTTGGAAGAAAAATCTACGTTAGATACATAACG<br>16; CGAGGCGCAGACCTTCATCAAGAGTAATCTCAGAACGAGTAG<br>17; ATGAGGAAGTTTTCTTTGACCCCCAGCGATACTTAGCCGGAA<br>18; TTCGAGGTGAATGCTTGCAGGGAGTTAAAGAAAGACTTTTTC<br>19; CGTAACGATCTAGAGAATAGAAAGGAACAATATCAGCTTGCT<br>20; TATAGCCCGGAACCTCATTTTCAGGGATAGCCTCATAGTTAG<br>21; CAGGAGTGTACTTATTCTGAAACATGAAAGGGTTGATATAAG<br>22; CCTCAGAACCGCCAGACGATTGGCCTTGATCTTTTGATGATA<br>23; TAGCAAGGCCGGCGTTTTCATCGGCATTTTCAGAGCCGCCAC<br>24; AGACACCACGGAGGAGGGAAGGTAAATATTCACCATTACCAT<br>25; AACAATGAAATAAATAATAACGGAATACCCAAAGAAACGCAA<br>26; TATTTATCCCAAGCATTAGACGGGAGAATTATAAGAGCAAGA<br>27; TAGCAAGCAAATAATCAAGATTAGTTGCTAAAACAGCCATAT<br>28; CTAATGCAGAACTAATCGGCTGTCTTTCCTTACCGCGCCCAA<br>29; AAGCCTGTTTAGCGCCAACATGTAATTTAGCAACATGTTCAG<br>30; CCTTTTTAACCTAATATATTTTAGTTAATTAATTACTAGAAA<br>31; ATTAATTACATTCGTCGCTATTAATTAATTTCTGAGAGACTA<br>32; CTTGAAAACATAGATGATGAAACAAAACAAA |

| |
|---|
| AGTTTCGTCACCAGCAGTTGGCAAATCACACCCTCAGAACCG |
| AGGATTAGCGGGGTCTAAAATATCTTTACAGTGCCCGTATAA |
| ATGGAAAGCGCAGTCGTCAATAGATAATAACCACCACCAGAG |
| TTCATAATCAAAATTTACAAACAATTCGAATCAGTAGCGACA |
| GTCACCGACTTGAGCGTTATTAATTTTAATGGTTTACCAGCG |
| CAGTATGTTAGCAAGAACAAAGAAACCAGTAAGCAGATAGCC |
| GAGCGCTAATATCACCTGATTATCAGATATGAAAATAGCAGC |
| ACCAACGCTAACGATGTTTGGATTATACGAGGCGTTTTAGCG |
| GCACTCATCGAGAACCATATCAAAATTAGAAAAATAATATCC |
| TAAAGTACCGACAATGCGTAGATTTTCAGCCAACGCTCAACA |
| CCGACCGTGTGATACAGTACCTTTTACACTGATGCAAATCCA |
| TAGATTAAGACGCTATTGCTTTGAATACCAGTACATAAATCA |
| CCGGCGAACGTGGCAAATCAAAAGAATAGCCCGACCCTAAAG |
| GCGGGCGCTAGGGCCTGCATTAATGAATCGGCCCCGAAATCG |
| GTAACCACCACACCACAGTTGAGGATCCCCGGGAGTCGGGAA |
| GCGTACTATGGTTGTGTGTTCAGCAAATCGTTATCTTCGCGT |
| CTCGTTAGAATCAGAACCGCAAGAATGCCAACGGTCATAAAC |
| AGGGATTTTAGACAGTGTACATCGACATAAAAAGTGGTGCTG |
| GTTTTTATAATCAGGCCAAGCTTTCAGAGGTGGTTGCCGCCA |
| ATCACGCAAATTAACACTCCAGCCAGCTTTCCGGACGTTGTA |
| AATAACATCACTTGACGCCATCAAAAATAATTCCAGTATCGG |
| CTTGCTGGTAATATGTCATTGCCTGAGAGTCTGAGCTCATTT |
| AACAGGAAAAACGCAGAAGCCTTTATTTCAACGGAGAGATCT |
| AATCGTCTGAAATGTCGCAAATGGTCAATAACCTGACCCTGT |
| CACACGACCAGTAACAGGTCAGGATTAGAGAGTTTAGTTTGA |
| AGAACCCTTCTGACGTCATAAATATTCATTGAAACCAGACCG |
| CAATATTTTTGAATCAGTTGAGATTTAGGAATAATAGCGTCC |
| TAGCCCTAAAACATCATTCAGTGAATAAGGCTTTTATTACAG |
| AGCAGAAGATAAAAATAAATTGTGTCGAAATCCCAAATCAAC |
| GCCTGCAACAGTGCACGAGGGTAGCAACGGCTAACGGAGATT |
| ATCTAAAGCATCACAAAAGGCTCCAAAAGGAGCCAGCAGCGA |
| AATCAATATCTGGTTACAAACTACAACGCCTGTTTTCACGTT |
| TTGAGGAAGGTTATTTTGCTCAGTACCAGGCGGGTAACACTG |
| TAATAGATTAGAGCCTCTGAATTTACCGTTCCAAGAAGGATT |
| AGAAGTATTAGACTCACCGGAACCAGAGCCACCAAAGCCAGA |
| ATCCTTTGCCCGAACCATTTGGGAATTAGAGCCTGCCATCTT |
| ATTATCATTTTGCGACGTAGAAAATACATACATATTATCACC |
| ATTATCATCATATTGAGAGATAACCCACAAGAACTTATTACG |
| AATATAATCCTGATGCGTCTTTCCAGAGCCTAAAGGGTAATT |
| AGGGTTAGAACCTACAAGCAAGCCGTTTTTATTCTGAATCTT |
| AGAAATAAAGAAATAAGGTAAAGTAATTCTGTCCCAAGTACC |
| TGAATATACAGTAAAATAAGGCGTTAAATAAGAAAGAGAATA |
| AACGGATTCGCCTGGAGAAGAGTCAATAGTGAATTTGAAATA |
| GCGCAGAGGCGAATAATTACCTGAGCAAAAGAAGCGATAGCT |

**Table S1** Oligonucleotide staple strand sequences for the 6HB slat. The 127 universal core staple strands are unchanging across all the 6HB slats folded, versus the 32 top- and 32 bottom-staple strands which are modified on their 3′ end to encode functionality in the slats. The latter top- and bottom-staple strands are arranged in the order in which they are displayed along the length of the slat, where the nodal position is indicated with the leading number. Given top- and bottom-staple strand pairs with the same number overlay one another. In general, we reserved the top-staple strands to append handle or complementary handle sequences to bind the slats to one another. The bottom-staple strands were alternatively appended with strong handle "plug" sequences to bind slats to the gridiron seed, the complementary 16-nt handle sequence to bind the nanocube contrast agent, the complementary handle sequence to bind DNA-PAINT imager strands, with biotin to affix the megastructures to streptavidin coated flow chambers for DNA-PAINT, or with complementary handle sequences to bind other slats in case of the tri-layer ribbon.

| **12HB universal core staples** | **12HB top staples** |
|---|---|
| ACCATCACCCAAACTATTAAAGAACGTGGAGAGAGAGTTGCA | 1; CCTGGCCCTCTCCAACGTCAAAGGGCGACTTGACGGGGAAAG |
| GCAAGCGGTCCACGGTTTTTCTTTTCACCATAAAGCCTGGGG | 2; CATAAAGTGGTGAGACGGGCAACAGCTGAGAAAGCGAAAGGA |
| TGCCTAATGAGTGGTGTGAAATTGTTATCCCTGCGGCCAGAA | 3; TCTGTGGTGGCTCACAATTCCACACAACCGGTCACGCTGCGC |
| TGCGGCGGGCCGTTGCCCCCTGCATCAGACGCGCGGTTGCGG | 4; ATCCGCCGGGATCCAGCGCAGTGTCACTCGCCGCTACAGGGC |
| TATGAGCCGGGTCGCTGGAGGTGTCCAGCACGTCGCTGGCAG | 5; TTTTCGTCTTCAGCGGGGTCATTGCAGGTATAACGTGCTTTC |
| CCTCCGGCCAGAGAGGGTAAAGTTAAACGAGGTGAAGGGATA | 6; TTTCTCCGTTGCTGATTGCCGTTCCGGCAGGAGGCCGATTAA |
| GCTCTCACGGAAAGAAACGTACAGCGCCATCCAGCTGGCGAA | 7; GCTATTACGGTTTACCAGTCCCGGAATTGAATCCTGAGAAGT |
| AGGGGGATGTGCTCCATTCAGGCTGCGCAAGGGATAGGTCAC | 8; GACCGTAATCTGTTGGGAAGGGCGATCGAAAAGAGTCTGTCC |
| GTTGGTGTAGATGGTGAGCGAGTAACAACCTATAAGCAAATA | 9; GGAAGATTGCGTCGGATTCTCCGTGGGACTTCTTTGATTAGT |
| TTTAAATTGTAAACATGTCAATCATATGTAAATCACCATCAA | 10; AGACAGTCACCCCGGTTGATAATCAGAACTCAAACTATCGGC |
| TATGATATTCAACAATGCCTGAGTAATGTGAAGAATTAGCAA | 11; GGCAAGGCATAGGTAAAGATTCAAAAGGCCGCCAGCCATTGC |

AATTAAGCAATAAGGCATCAATTCTACTAACTAAAGTACGGT
GTCTGGAAGTTTCTGGCTTAGAGCTTAATTCATCAAAAAGAT
TAAGAGGAAGCCCCCATAAATCAAAAATCACAAAATAGCGAG
AGGCTTTTGCAAAACGAGGCATAGTAAGAGGCTCATTATACC
ACTAAAACACTCACCATTAAACGGGTAAAAATAACCGATATA
TTCGGTCGCTGAGTTCTTAAACAGCTTGATCAACTTTCAACA
GTTTCAGCGGAGTAAGTTTTGTCGTCTTTCGAACCGCCACCC
TCAGAGCCACCACTAGGTGTATCACCGTACGCCCCCTGCCTA
TTTCGGAACCTATGGTAATAAGTTTTAACGGCATTGACAGGA
GGTTGAGGCAGGTCACCCTCAGAGCCACCATTGCCTTTAGCG
TCAGACTGTAGCGAAACGTCACCAATGAAAAAAGGGCGACAT
TCAACCGATTGAGATAAGTTTATTTTGTCATACCAGAAGGAA
ACCGAGGAAACGCGCAATAGCTATCTTACCGAGAATAACATA
AAAACAGGGAAGCTCCAAATAAGAAACGATACTTGCGGGAGG
TTTTGAAGCCTTACAGATATAGAAGGCTTATTACGAGCATGT
AGAAACCAATCAAGCGCCTGTTTATCAACAAATTGAGAATCG
CCATATTTAACAATATCATATGCGTTATACCAAAGAACGCGA
GAAAACTTTTTCACCGGCTTAGGTTGGGTTAGTGAATAACCTTGCT
CACTAAATCGGAAGATAGGGTTGAGTGTTGCGAAAATCCTGT
TTGATGGTGGTTAACGCGCGGGGAGAGGCGCGTTGCGCTCAC
TGCCCGCTTTCCTACCGAGCTCGAATTCGGGTTTCTGCCAGC
ACGCGTGCCTGTACGGCATCAGATGCCGGGTAATGGGTAAAG
GTTTCTTTGCTCGCAGCACCGTCGGTGGTCGTGCCGGACTTG
TAGAACGTCAGCAATCCCGTAAAAAAAGCGGATCAAACTTAA
ATTTCTGCTCATAGCCGCCACGGGAACGGGTAACGCCAGGGT
TTTCCCAGTCACGCACCGCTTCTGGTGCCATCTGCCAGTTTG
AGGGGACGACGAGCGTCTGGCCTTCCTGTATTCGCATTAAAT
TTTTGTTAAATCGAGCAAACAAGAGAATCAATGCCGGAGAGG
GTAGCTATTTTTCAAGGATAAAAATTTTTTAAATCGGTTGTA
CCAAAAACATTATGTTTAGCTATATTTTCTTCCCAATTCTGC
GAACGAGTAGATACCTTTAATTGCTCCTTCGTTTTAATTCGA
GCTTCAAAGCGATCCCCCTCAAATGCTTTGGTAATAGTAAAA
TGTTTAGACTGGCCACATTCAACTAATGCAATAAAACGAACT
AAACAAAGTACACAGAGGCTTTGAGGACTGCCGCTTTTGCGG
GATCGTCACCCTCTTTAATTGTATCGGTTCTAAAGGAATTGC
GAATAATAATTTAGCATTCCACAGACAGCCAAGCCCAATAGG
AACCCATGTACCATAAGTGCCGTCGAGAGTATTAAGAGGCTG
AGACTCCTCAAGGTAAGCGTCATACATGGATTCACAAACAAA
TAAATCCTCATTACCGGAACCGCCTCCCTCGGTCATAGCCCC
CTTATTAGCGTTAGCAAAATCACCAGTAGGACGGAAATTATT
CATTAAAGGTGAAAAGGTGGCAACATATAAAAAGAACTGGCA
TGATTAAGACTCTTGAGTTAAGCCCAATAAACTGAACACCCT
GAACAAAGTCAGTTTGCCAGTTACAAAATTTTTGCACCCAGC
TACAATTTTATCTTCATCGTAGGAATCATTATCATTCCAAGA
ACGGGTATTAAACAGACGACGACAATAAAGCAGAGGCATTTT
CGAGCCAGTAATATAAACACCGGAATCATTCATCTTCTGACC
TAAATTTAATGGTTTATCAAAATCATAGGTTCCCTTAGAATC
GGAGCCCCCTTTTTTTTTTGATTTAGAGAAAACCGTCTATCA
GCAAAATCCCTTATGAGAAAGGAAGGGAATTGCCCTTCACCG
ACCTGTCGTGCCAGGCTGGCAAGTGTAGATACGAGCCGGAAG
CCGTGAGCCTCCTCCGCCGCGCTTAATGGCGCGCCTGTGCAC
ATCCCTTACACTGGCTTTGACGAGCACGCGCTTTCGCACTCA
GTCTGGTCAGCAGCAGCGGGAGCTAAACAAACGCGGTCCGTT
GCAGTTGGGCGGTTGGAACGGTACGCCATGTGAGAGATAGAC
AAACGACGGCCAGTTGAGGCCACCGAGTGTGCGGGCCTCTTC
CCTCAGGAAGATCGCCGTTGTAGCAATAACAAACGGCGGATT
TTTAACCAATAGGACCTGAGTAGAAGAAAAGCCCCAAAAACA
ACAAAGGCTATCAGCCAGAACAATATTAGTGAGAAAGGCCGG
AATACTTTTGCGGGTCATGGAAATACCTCAATAAATCATACA
CCATTAGATACATTGATTATTTACATTGCTCAACATGTTTTA
GAAGCAAACTCCAATAAAAGGGACATTCATAGTCAGAAGCAA
AATACTGCGGAATCCTGAAAGCGTAAGAGTTTACCAGACGAC
GTAGAAAGATTCATGGCTATTAGTCTTTACCTTATGCGATTT
GTAACAAAGCTGCTCGCCATTAAAAATAAAGAGGACAGATGA
TGTATCATCGCCTGCAGAGGTGAGGCGGGGCACCAACCTAAA
AAGACAGCATCGGACACGCTGAGAGCCAATGACAACAACCAT
GAAAATCTCCAAAACTTGCTGAACCTCATTTCTGTATGGGAT
AGTTTCGTCACCAGCAGTTGGCAAATCACACCCTCAGAACCG
AGGATTAGCGGGGTCTAAAATATCTTTACAGTGCCCGTATAA
ATGGAAAGCGCAGTCGTCAATAGATAATAACCACCACCAGAG
TTCATAATCAAAATTTACAAACAATTCGAATCAGTAGCGACA
GTCACCGACTTGAGCGTTATTAATTTTAATGGTTTACCAGCG
CAGTATGTTAGCAAGAACAAAGAAACCAGTAAGCAGATAGCC
GAGCGCTAATATCACCTGATTATCAGATATGAAAATAGCAGC
ACCAACGCTAACGATGTTTGGATTATACGAGGCGTTTTAGCG
GCACTCATCGAGAACCATATCAAAATTAGAAAAATAATATCC
TAAAGTACCGACAATGCGTAGATTTTCAGCCAACGCTCAACA
CCGACCGTGTGATACAGTACCTTTTACACTGATGCAAATCCA
TAGATTAAGACGCTATTGCTTTGAATACCAGTACATAAATCA
CCGGCGAACGTGGCAAATCAAAAGAATAGCCCGACCCTAAAG
GCGGGCGCTAGGGCCTGCATTAATGAATCGGCCCCGAAATCG
GTAACCACCACACCACAGTTGAGGATCCCCGGGAGTCGGGAA
GCGTACTATGGTTGTGTGTTCAGCAAATCGTTATCTTCGCGT
CTCGTTAGAATCAGAACCGCAAGAATGCCAACGGTCATAAAC
AGGGATTTTAGACAGTGTACATCGACATAAAAAGTGGTGCTG
GTTTTTATAATCAGGCCAAGCTTTCAGAGGTGGTTGCCGCCA
ATCACGCAAATTAACACTCCAGCCAGCTTTCCGGACGTTGTA
AATAACATCACTTGACGCCATCAAAAATAATTCCAGTATCGG
CTTGCTGGTAATATGTCATTGCCTGAGAGTCTGAGCTCATTT
AACAGGAAAAACGCAGAAGCCTTTATTTCAACGGAGAGATCT
AATCGTCTGAAATGTCGCAAATGGTCAATAACCTGACCCTGT
CACACGACCAGTAACAGGTCAGGATTAGAGAGTTTAGTTTGA
AGAACCCTTCTGACGTCATAAATATTCATTGAAACCAGACCG
CAATATTTTTGAATCAGTTGAGATTTAGGAATAATAGCGTCC
TAGCCCTAAAACATCATTCAGTGAATAAGGCTTTTATTACAG
GCCTGCAACAGTGCACGAGGGTAGCAACGGCTAACGGAGATT
ATCTAAAGCATCACAAAAGGCTCCAAAAGGAGCCAGCAGCGA
AATCAATATCTGGTTACAAACTACAACGCCTGTTTTCACGTT
TTGAGGAAGGTTATTTTGCTCAGTACCAGGCGGGTAACACTG
TAATAGATTAGAGCCTCTGAATTTACCGTTCCAAGAAGGATT
AGAAGTATTAGACTCACCGGAACCAGAGCCACCAAAGCCAGA
ATCCTTTGCCCGAACCATTTGGGAATTAGAGCCTGCCATCTT

12; AATATGCAATAGTAGTAGCATTAACATCACATTTTGACGCTC
13; AGCGGATTGGCTGAATATAATGCTGTAGGCAGATTCACCAGT
14; GATAAAAACGGTCTTTACCCTGACTATTTGGCCAACAGAGAT
15; TAAGAACTGCAACACTATCATAACCCTCATACGTGGCACAGA
16; ACGGTGTACACTTTAATCATTGTGAATTAATGCGCGAACTGA

#### 12HB bottom staples

1; TCTGTAAATTAACAATTTCATTTTTTTAATGGAAACAAGTTACAAAATC
2; ATATATGTGATATAACTATATGTAAATGTCGGGAGAAACAAT
3; ATCGCAAGAAAATTCTTACCAGTATAAAGGTTTAACGTCAGA
4; GTAGGGCTTATAGATAAGTCCTGAACAATTTGCACGTAAAAC
5; CATCCTAATTCCGGTATTCTAAGAACGCTTCTGAATAATGGA
6; AACCTCCCGTTTTTGTTTAACGTCAAAAGATGGCAATTCATC
7; CTTTACAGAGAAGCCCTTTTTAAGAAAACCAGAAGGAGCGGA
8; GAACAAAGTCAATCAATAGAAAATTCATAAAGTTTGAGTAAC
9; CCAAAGACACCATCGATAGCAGCACCGTACAACTCGTATTAA
10; GAATCAAGTCCCTCAGAGCCGCCACCAGACATTTGAGGATTT
11; CCGCCGCCAGGGTCAGTGCCTTGAGTAAGGAGCACTAACAAC
12; ACAGTTAATTCAGGAGGTTTAGTACCGCACAGTTGAAAGGAA
13; CCACCCTCACAGACGTTAGTAAATGAATAATATCAAACCCTC
14; TTTGCTAAAACCGATAGTTGCGCCGACAGCAGCAAATGAAAA
15; CGCCCACGCTACGTAATGCCACTACGAATCAGTATTAACACC
16; ACGAAAGAGCCGAACTGACCAACTTTGACCGAACGAACCACC

| |
|---|
| ATTATCATTTTGCGACGTAGAAAATACATACATATTATCACC |
| ATTATCATCATATTGAGAGATAACCCACAAGAACTTATTACG |
| AATATAATCCTGATGCGTCTTTCCAGAGCCTAAAGGGTAATT |
| AGGGTTAGAACCTACAAGCAAGCCGTTTTTATTCTGAATCTT |
| AGAAATAAAGAAATAAGGTAAAGTAATTCTGTCCCAAGTACC |
| TGAATATACAGTAAAATAAGGCGTTAAATAAGAAAGAGAATA |
| AACGGATTCGCCTGGAGAAGAGTCAATAGTGAATTTGAAATA |
| GCGCAGAGGCGAATAATTACCTGAGCAAAAGAAGCGATAGCT |
| AGTCAGGACGTTGGAGATGGTTTAATTTCAAGACCAGGCtttttttt |
| AGCAGAAGATAAAAATAAATTGTGTCGAAATCCCAAATCAACtttttttt |
| ACATTAATTGGTTTGCGTATTCCTTTTTAACCTAATATATTT |
| CATACCGGGTAATCATGGTCAAAGCCTGTTTAGCGCCAACAT |
| CTGCGGCTGGTTACCTGCAGCCTAATGCAGAACTAATCGGCT |
| ATAACGGAAGCCATCCCACGCTAGCAAGCAAATAATCAAGAT |
| AGAAACAGCCGCACAGGCGGCTATTTATCCCAAGCATTAGAC |
| TTAAGTTGGATAACCTCACCGAACAATGAAATAAATAATAAC |
| TAACCGTGCGGAAACCAGGCAAGACACCACGGAGGAGGGAAG |
| TTTGTTAAAAGCCAGCTTTCATAGCAAGGCCGGCGTTTTCAT |
| TGATAAATTGATGAACGGTAACCTCAGAACCGCCAGACGATT |
| GCATAAAGCAGAACCCTCATACAGGAGTGTACTTATTCTGAA |
| AACAGTTGAATTTGGGGCGCGTATAGCCCGGAACCTCATTTT |
| CAAATATCGTTGATAAGAGGTCGTAACGATCTAGAGAATAGA |
| GCCAGAGGGAAACAGTTCAGATTCGAGGTGAATGCTTGCAGG |
| ATCTACGTTAGATACATAACGATGAGGAAGTTTTCTTTGACC |
| CTCAGAACGAGTAGCGAGGCGCAGACCT |
| CCCCAGCAGGTTCCAGTTTGGATTAATTACATTCGTCGCTAT |
| AACGGAACAACAGCCCTGACGAGAAACACTGtttttttt |
| GAGTTAAAGAAAGACTTTTTCCCAAAAGGAATTAGAAGTTTT |
| AAGGAACAATATCAGCTTGCTAAACGAGAATGAGAAAGACTT |
| CAGGGATAGCCTCATAGTTAGCATTTTTGCGGAATTCCATAT |
| ACATGAAAGGGTTGATATAAGAGCTGAAAAGGTAGCCTCAGA |
| GGCCTTGATCTTTTGATGATATATTTTAAATGCCGTTCTAGC |
| CGGCATTTTCAGAGCCGCCACTCGTAAAACTAGCGTTAATAT |
| GTAAATATTCACCATTACCATTCAACATTAAATGGCGCATCG |
| GGAATACCCAAAGAAACGCAAAAGCGCCATTCGGCAAGGCGA |
| GGGAGAATTATAAGAGCAAGAGAAACAATCGGCAAGAGACGC |
| TAGTTGCTAAAACAGCCATATCTTTAGTGATGACACATCCTC |
| GTCTTTCCTTACCGCGCCCAAAACCAGCTTACGACTGTTGCC |
| GTAATTTAGCAACATGTTCAGCAGCGGTGCCGGTTTCACGGT |
| TAGTTAATTAATTACTAGAAATAGCTGTTTCCTAGCTAACTC |
| TAATTAATTTCTGAGAGACTAGGGCGCCAGGGTGCTGGTTTG |
| CCCAGCGATACTTAGCCGGAATAAATTGGGCTTGGAAGAAAA |
| ttttttttTTCATTACCGCGACCTGCTCCATGTTTATACCAAGCGCG |
| CTTGAAAACATAGATGATGAAACAAAACAAAAACAAGAGTCCATCAAGTTTTTTGGCCGTAAAG |
| ttttttttACGGTCAATCATAAGGGAAGCAAAAGAATAC |

**Table S2** Oligonucleotide staple strand sequences for the 12HB slat. The 158 universal core staple strands are unchanging across all the 12HB slats folded, versus the 16 top- and 16 bottom-staple strands which are modified on their 3′ end to encode functionality in the slats. The latter top- and bottom-staple strands are arranged in the order in which they are displayed along the length of the slat, where the nodal position is indicated with the leading number. Given top- and bottom-staple strand pairs with the same number overlay one another. In general, we reserved the top-staple strands to append handle or complementary handle sequences to bind the slats to one another.

| seed universal core staples | seed non-socketed staples |
|---|---|
| 1; TACCGCCACCCTCAGAACCGCCGTCGAGAGGGTTGATATAAG | 65c1; GGGAATTAGAGCCAGCAAAATGTTTATGTAGATGAAGGTATA |
| 2; TCATTAAAGGTGAATTATCACTTCTGCAATGTGCGAGAAATG | 66c1; ATTCATATGGTTTACCAGCGCTATCACGAGTACGGTGGAAAC |
| 3; TAGCAAGCAAATCAGATATAGCAGGGATAGCAAGCCCAATAG | 67c1; CCTTGAAAACATAGCGATAGCGAGTTAGAGTCTGAGCAAAAA |
| 4; GCTTCTGTAAATCGTCGCTATAAACATATAGATGATTAAACC | 68c1; ATTAATTACATTTAACAATTTGCACTCGCGGGGATTTATTTT |
| 5; AGTATTAACACCGCCTGCAACACAGACAGCCCTCATAGTTAG | 69c1; TGACGGGGAAAGCCGGCGAACCTTACTGTTTCTTTACATAAA |
| 6; GCACTAAATCGGAACCCTAAATTTTGTTTTATGGAGATGATA | 70c2; TAGCAAGGCCGGAAACGTCACCGAACAAGACCCGTTAGTAAC |
| 7; CAAAGGGCGAAAAACCAACAGCTGATTGCCCTGCGCCAGG | 71c2; AGACACCACGGAATAAGTTTATGCAGATCCGGTGTCTTGTCT |
| 8; AGTCCACTATTAAAGAAGAGAGTTGCAGCAAGCAACGCGC | 72c2; AGTCAATAGTGAATTTATCAAGTATCTGCATATGATGTCTGA |
| 9; TAGGGTTGAGTGTTGTGCCCCAGCAGGCGAAAACCTGTCG | 73c2; CTGAGCAAAAGAAGATGATGAGAAACGACATACATTGCAAGG |
| 10; TTTTTTTTTTAATCCCTTATAAATCAGTTCCGAAATCGGCAATTTTTTTTTT | 74c2; AAAGCGAAAGGAGCGGGCGCTCTGAATTTCGCGTCGTCTTCA |
| 11; TTTTTTTTTTGTGAGACGGGCGTCTATCA | 75c3; CACCGTAATCAGTAGCGACAGGTTTCTTGTTGTTCGCCATCC |
| 12; GTGGTTTTTGTTTCCTGTGTGAAA | 76c3; AAAATACATACATAAAGGTGGCTATTACGGGGTTGGAGGTCA |
| 13; CGTATTGGTCACCGCCTGGCCCTGACGTGGACTCCAACGT | 77c3; CCTTTTTAACCTCCGGCTTAGTGAGTATTACGAAGGTGTTAT |
| 14; GGGGAGAGATTCCACACAACATAC | 78c3; AGTTACAAAATCGCGCAGAGGAGAGTGAGATCGGTTTTGTAA |
| 15; GAATCGGCCGGTCCACGCTGGTTTTCCAGTTTGGAACAAG | 79c3; GTCACGCTGCGCGTAACCACCCCAGGAGAACGAGGATATTGC |
| 16; TGCCAGCTGTGTAAAGCCTGGGGT | 80c4; CAGACTGTAGCGCGTTTTCATAACGAAGACGCCTGGTCGTTC |
| 17; GTCGGGAAATCCTGTTTGATGGTGAAAGAATAGCCCGAGA | 81c4; ATGATTAAGACTCCTTATTACTGCTAAACTGGAAAGCAACGA |
| 18; TTTTTTTTTTGTTGCGCTCACTGCCCAACTCACATTAATTGCTTTTTTTTTT | 82c4; TAAATGCTGATGCAAATCCAACGAAGTGAGCGAAATTAACTC |
| 19; GTCATAGCTCTTTTCACCATTTTTTTTTT | 83c4; GGGAGAAACAATAACGGATTCTGTTGAGCTTGAAACAGCAAA |
| 20; TTGTTATCCGCTCACAGCGGTTTG | 84c4; CCGCTACAGGGCGCGTACTATTTTCCATGAATTGGTAACACC |
| 21; GAGCCGGAAGCATAAAGCATTAAT | 85c5; CCTTATTAGCGTTTGCCATCTGCAACACAGCAATAAAAATGC |
| 22; GCCTAATGAGTGAGCTGCTTTCCA | 86c5; CCGAGGAAACGCAATAATAACGTTGCCAGGAGGATCTGGAAC |

23; AAACATCGGGTTGAGTATTATGTGGCGAGAAAGGAAGGGAAG
24; GCAATACATCAAACGCCGCGAACACCCGCCGCGCTTAATGCG
25; GGGGGATGTGCTGCAAGGCGAATCAGAGCGGGAGCTAAACAG
26; TCGTAAAACTAGCATGTCAATATCAGTGAGGCCACCGAGTAA
27; CGAACGAGTAGATTTAGTTTGACTTGCCTGAGTAGAAGAACT
28; ACCCTCGTTTACCAGACGACGAACGCTCATGGAAATACCTAC
29; TGTGTCGAAATCCGCGACCTGAGTAATAAAAGGGACATTCTG
30; TTTCAGCGGAGTGAGAATAGATGAATGGCTATTAGTCTTTAA
31; AGAGTTTCTGCGGCAGTTAATCAATGAAACCATCGATAGCAG
32; AGTGTGGCGATCCGATAGATGCGGCATTTTCGGTCATAGCCC
33; GTGGGAACAAACGGCGGATTGCGCCTCCCTCAGAGCCGCCAC
34; CTTTATTTCAACGCAAGGATACGCCGCCAGCATTGACAGGAG
35; AAGAGGAAGCCCGAAAGACTTAATGGAAAGCGCAGTCTCTGA
36; TAAATTGGGCTTGAGATGGTTTTTTAACGGGGTCAGTGCCTT
37; GGATCGTCACCCTCAGCAGCGACATGAAAGTATTAAGAGGCT
38; TCTAATGAAGACAAATCCCCACGTCACCGACTTGAGCCATTT
39; GCACTTCCGAGTCACAGGAGAATGGATC
40; ATAGAGTCGGCATACAAATATTCCATGAAGGTTTAT
41; CGATGCCAGAGTCTGTAGTGTCAGATGATGACCGTA
42; GAAGACTCCTGTTATCAAGCACTGCACTGGTGACCT
43; GATTGTGCTGGAACTGCTGGATGAACGGGAAAGAAA
44; GCCTGAACCACCAGGCTATATCTGCCACTCATTGTT
45; TCGGTGCGGGCCTCTTCGCTATTACGCCAGCTGGCG
46; ACATTAAATGTGAGCGAGTAACAACCCGTCGGATTC
47; TGAGAGTCTGGAGCAAACAAGAGAATCGATGAACGG
48; AAAAACATTATGACCCTGTAATACTTTTGCGGGAGA
49; GGAAGTTTCATTCCATATAACAGTTGATTCCCAATT
50; ATTATAGTCAGAAGCAAAGCGGATTGCATCAAAAAG
51; AAAGGAATTACGAGGCATAGTAAGAGCAACACTATC
52; AATAAGGCTTGCCCTGACGAGAAACACCAGAACGAG
53; ACAAAGTACAACGGAGATTTGTATCATCGCCTGATA
54; GTCGCTGAGGCTTGCAGGGAGTTAAAGGCCGCTTTT
55; AATTTTCTGTATGGGATTTTGCTAAACAACTTTCAA
56; CACCGTACTCAGGAGGTT
57; AGCCCGGAATAGGTGTAT
58; CCGATTGAGGGAGGGAAGGTAAATATTGACGGAAAT
59; CGTTTTTATTTTCATCGTAGGAATCATTACCGCGCC
60; AAACAGTACATAAATCAATATATGTGAGTGAATAAC
61; AATCAAGTTTTTTGGGGTCGAGGTGCCGT
62; CGAACCACCAGCAGAAGATAAAACAGAGGTGAGGCG
63; CGATGGCCCACTACGTGAACCATCACCCA
64; CCCGGGTACCGAGCTCGAATTCGTAATC
145; ACTGATACCGTGCAAAATTATCAAAGACAAAAGGGCGACATT
146; CCATGCAGACATCACGAAGGTCACCAGTAGCACCATTACCAT
147; TATCGACATCATTACGCATCGCAACATATAAAAGAAACGCAA
148; AGTCCGTGAAGACGGAAACCAAATCAAGTTTGCCTTTAGCGT
149; AGGGTTGTCGGACTTGTGCAAGGAATACCCAAAAGAACTGGC
150; CAGAAATAGAAGAATTACAGCTTTCATAATCAAAATCACCGG
151; TTGGTGTAGATGGGCGCATCGATCTTACCGAAGCCCTTTTTA
152; TCAAAAATAATTCGCGTCTGGAGCCACCACCCTCAGAGCCGC
153; TATTTTAAATGCAATGCCTGATGAGCGCTAATATCAGAGAGA
154; ATTAAGCAATAAAGCCTCAGAGGCCTTGATATTCACAAACAA
155; AGCTTCAAAGCGAACCAGACCTTTACAGAGAGAATAACATAA
156; AAACGAGAATGACCATAAATCCATACATGGCTTTTGATGATA
157; TGAATTACCTTATGCGATTTTTTACAAAATAAACAGCCATAT
158; GATATTCATTACCCAAATCAACAGTTAATGCCCCCTGCCTAT
159; GTAGCAACGGCTACAGAGGCTTAGTTGCTATTTTGCACCCAG
160; CCGACAATGACAACAACCATCTAGGATTAGCGGGGTTTTGCT
161; CAGAGCCACCACCCTCATTTTAAGGCTTATCCGGTATTCTAA
162; GATACTTGCCCTCTCTGTACATAATTAATTTTCCCTTAGAAT
163; GATTGGGCGTTATCAATGTTGTTTTGTCACAATCAATAGAAA
164; ATGGGTTCAGGATGCAGGTGAAATCATAGGTCTGAGAGACTA
165; CTCGGATGGGAGTAAGCGTATGCAGTATGTTAGCAAACGTAG
166; TTATCAGTAAACAGAGAGGTTTCGCAAGACAAAGAACGCGAG
167; TCAGGGATTAATGAAAGATGGAACAAAGTTACCAGAAGGAAA
168; GAGGGGACGACGACAGTATCGACCGACCGTGTGATAAATAAG
169; TTTTTGTTAAATCAGCTCATTAGCCCAATAATAAGAGCAAGA
170; AAAAGGGTGAGAAAGGCCGGACGTTATACAAATTCTTACCAG
171; AACATCCAATAAATCATACAGGGGAGAATTAACTGAACACCC
172; AGGATTAGAGAGTACCTTTAAGTAATTTAGGCAGAGGCATTT
173; AATATTCATTGAATCCCCCTCGAAACGATTTTTTGTTTAACG
174; GTCAGGACGTTGGGAAGAAAAGACAATAAACAACATGTTCAG
175; AGGCTGGCTGACCTTCATCAATACCAACGCTAACGAGCGTCT
176; ATGAGGAAGTTTCCATTAAACATCCTAATTTACGAGCATGTA
177; TTCGAGGTGAATTTCTTAAACACCTCCCGACTTGCGGGAGGT
178; GAACCCATGTACCGTAACACTCGCACTCATCGAGAACAAGCA
179; CAATATTACATAACAATCCTCCATTTGAATTACCTTTTTTAA
180; AGTTTATAAATGAGTATCAATTTAGATTAAGACGCTGAGAAG
181; CGCTGGCATTCGCATCAAAGGCGAATTATTCATTTCAATTAC
182; ATCAAAACTCAACGAGCAGCGGTTGGGTTATATAACTATATG
183; TCAGGCACTGCGTGAAGCGGCAGTAACAGTACCTTTTACATC
184; CTTTATTATTCGCATTCACCCTAGTTAATTTCATCTTCTGAC
185; AGCCAGCTTTCCGGCACCGCTACCTACCATATCAAAATTATT
186; TTAAATTGTAAACGTTAATATCGGAATCATAATTACTAGAAA
187; ATGATATTCAACCGTTCTAGCATATTCCTGATTATCAGATGA
188; AGCTGAAAAGGTGGCATCAATTAGGGCTTAATTGAGAATCGC
189; CATTTTTGCGGATGGCTTAGACCGAACGTTATTAATTTTAAA
190; ATGTTTAGACTGGATAGCGTCATAAAGTACCGACAAAAGGTA
191; TAACGGAACAACATTATTACAAGAGCCGTCAATAGATAATAC
192; CTTTGAAAGAGGACAGATGAATATCAACAATAGATAAGTCCT
193; TACGAAGGCACCAACCTAAAACTGGTCAGTTGGCAAATCAAC
194; CTCCAAAAGGAGCCTTTAATTGTCTTTCCTTATCATTCCAAG
195; CTACAACGCCTGTAGCATTCCAGTGCCACGCTGAGAGCCAGC
196; ATCTGAACTCGCTACGGCGGGGGGAGCCCCCGATTTAGAGCT
197; CATTGCTGATACCGTTTAGCTAACAAACATCAAGAAAACAAA
198; AAGATAACGCTTGTGAAAATGAGGGCGCTGGCAAGTGTAGCG
199; GCTAACAGTAGGGAAACTGCGGCCTGATTGCTTTGAATACCA
200; CTGGGGATTTGACGCAGACCTGGTTGCTTTGACGAGCACGTA
201; TTTTCCCAGTCACGACGTTGTGAAATTGCGTAGATTTTCAGG
202; AAGCGCCATTCGCCATTCAGGAGACAGGAACGGTACGCCAGA
203; TCAGAAAAGCCCCAAAAACAGCTGATTGTTTGGATTATACTT
204; GGTAGCTATTTTTGAGAGATCATTAACCGTTGTAGCAATACT
87c5; AAAACTTTTTCAAATATATTTTCATGCGTATTAACCAACAGT
88c5; TTTAACGTCAGATGAATATACAGAGCAGGCAATGCATGACGA
89c5; TAACGTGCTTTCCTCGTTAGATTAAGTTGGGTAACGCCAGGG
90c6; AACCAGAGCCACCACCGGAACACCGTAATGGGATAGGTCACG
91c6; AGAAAAGTAAGCAGATAGCCGCAGACATCATTGATTCAGCAT
92c6; CTAAATTTAATGGTTTGAAATGCCTCAGGAAGATCGCACTCC
93c6; TGCACGTAAAACAGAAATAAAAAAACGACGGCCAGTGCCAAG
94c6; GAGGCCGATTAAAGGGATTTTCTGCGCAACTGTTGGGAAGGG
95c7; CCTCAGAACCGCCACCCTCAGCCTTCCTGTAGCCAGCTTTCA
96c7; AACAATGAAATAGCAATAGCTTAACCGTGCATCTGCCAGTTT
97c7; GCGTTAAATAAGAATAAACACTTTGTTAAAATTCGCATTAAA
98c7; CTGAATAATGGAAGGGTTAGATCTGGTGCCGGAAACCAGGCA
99c7; ATCCTGAGAAGTGTTTTTATACATATGTACCCCGGTTGATAA
100c8; CACCAGAACCACCACCAGAGCAAAATTTTTAGAACCCTCATA
101c8; TAACCCACAAGAATTGAGTTATTTTAACCAATAGGAACGCCA
102c8; AAGCCTGTTTAGTATCATATGGACAGTCAAATCACCATCAAT
103c8; TGGCAATTCATCAATATAATCGAAGATTGTATAAGCAAATAT
104c8; AAGAGTCTGTCCATCACGCAATACAAAGGCTATCAGGTCATT
105c9; GTTGAGGCAGGTCAGACGATTGCATAAAGCTAAATCGGTTGT
106c9; TGAACAAAGTCAGAGGGTAATGTAATGTGTAGGTAAAGATTC
107c9; TATAAAGCCAACGCTCAACAGTCTACTAATAGTAGTAGCATT
108c9; AGAAGGAGCGGAATTATCATCTGATAAATTAATGCCGGAGAG
109c9; TCTTTGATTAGTAATAACATCACCATTAGATACATTTCGCAA
110c10; ATAAATCCTCATTAAAGCCAGCAAATATCGCGTTTTAATTCG
111c10; AAACAGGGAAGCGCATTAGACGCAAGGCAAAGAATTAGCAAA
112c10; CATATTTAACAACGCCAACATTTGCTCCTTTTGATAAGAGGT
113c10; AGTTTGAGTAACATTATCATTTATATTTTCATTTGGGGCGCG
114c10; CAAACTATCGGCCTTGCTGGTATATGCAACTAAAGTACGGTG
115c11; ATTTACCGTTCCAGTAAGCGTAAAAATCAGGTCTTTACCCTG
116c11; TCAAAAATGAAAATAGCAGCCGGAAGCAAACTCCAACAGGTC
117c11; TCGAGCCAGTAATAAGAGAATCAATACTGCGGAATCGTCATA
118c11; AACTCGTATTAAATCCTTTGCGCTTAATTGCTGAATATAATG
119c11; GCCAGCCATTGCAACAGGAAAATAAAAACCAAAATAGCGAGA
120c12; CAGGAGTGTACTGGTAATAAGTAATTTCAACTTTAATCATTG
121c12; TATTTATCCCAATCCAAATAAAAATGCTTTAAACAGTTCAGA
122c12; AAGTAATTCTGTCCAGACGACATCTACGTTAATAAAACGAAC
123c12; ATTTGAGGATTTAGAAGTATTGCCAGAGGGGGTAATAGTAAA
124c12; ATTTTGACGCTCAATCGTCTGAACTAATGCAGATACATAACG
125c13; GAGTAACAGTGCCCGTATAAACGTAACAAAGCTGCTCATTCA
126c13; TTCCAGAGCCTAATTTGCCAGAAGAACTGGCTCATTATACCA
127c13; CTAATGCAGAACGCGCCTGTTCGGTGTACAGACCAGGCGCAT
128c13; AGCACTAACAACTAATAGATTGGTAGAAAGATTCATCAGTTG
129c13; AGATTCACCAGTCACACGACCCTCCATGTTACTTAGCCGGAA
130c14; TTCGGAACCTATTATTCTGAAAAAGACAGCATCGGAACGAGG
131c14; CTACAATTTTATCCTGAATCTGAGTAATCTTGACAAGAACCG
132c14; GAACAAGAAAAATAATATCCCGGGTAAAATACGTAATGCCAC
133c14; AGTTGAAAGGAATTGAGGAAGTAAGGGAACCGAACTGACCAA
134c14; GCCAACAGAGATAGAACCCTTCCCAGCGATTATACCAAGCGC
135c15; GAGACTCCTCAAGAGAAGGATGCCCACGCATAACCGATATAT
136c15; TTTGAAGCCTTAAATCAAGATTTGAGGACTAAAGACTTTTTC
137c15; GAAACCAATCAATAATCGGCTGTATCGGTTTATCAGCTTGCT
138c15; TATCAAACCCTCAATCAATATCGAAAGAGGCAAAAGAATACA
139c15; ACGTGGCACAGACAATATTTTAAGGAACAACTAAAGGAATTG
140c16; CAGTACCAGGCGGATAAGTGCCACCCTCAGAACCGCCACCCT
141c16; GAACGCGAGGCGTTTTAGCGAAGCTTGATACCGATAGTTGCG
142c16; AACGGGTATTAAACCAAGTACGAGTTTCGTCACCAGTACAAA
143c16; AGCAAATGAAAAATCTAAAGCTGAAAATCTCCAAAAAAAAGG
144c16; TGCGCGAACTGATAGCCCTAACGTCTTTCCAGACGTTAGTAA

#### seed socketed staples

65c1; GCCAGCAAAATGTTTATGTAGATGAAGGTATA
66c1; TTTACCAGCGCTATCACGAGTACGGTGGAAAC
67c1; ATAGCGATAGCGAGTTAGAGTCTGAGCAAAAA
68c1; TTTAACAATTTGCACTCGCGGGGATTTATTTT
69c1; AGCCGGCGAACCTTACTGTTTCTTTACATAAA
70c2; GGAAACGTCACCGAACAAGACCCGTTAGTAAC
71c2; GAATAAGTTTATGCAGATCCGGTGTCTTGTCT
72c2; GAATTTATCAAGTATCTGCATATGATGTCTGA
73c2; GAAGATGATGAGAAACGACATACATTGCAAGG
74c2; GAGCGGGCGCTCTGAATTTCGCGTCGTCTTCA
75c3; AGTAGCGACAGGTTTCTTGTTGTTCGCCATCC
76c3; CATAAAGGTGGCTATTACGGGGTTGGAGGTCA
77c3; CTCCGGCTTAGTGAGTATTACGAAGGTGTTAT
78c3; TCGCGCAGAGGAGAGTGAGATCGGTTTTGTAA
79c3; GCGTAACCACCCCAGGAGAACGAGGATATTGC
80c4; CGCGTTTTCATAACGAAGACGCCTGGTCGTTC
81c4; CTCCTTATTACTGCTAAACTGGAAAGCAACGA
82c4; TGCAAATCCAACGAAGTGAGCGAAATTAACTC
83c4; ATAACGGATTCTGTTGAGCTTGAAACAGCAAA
84c4; GCGCGTACTATTTTCCATGAATTGGTAACACC
85c5; GTTTGCCATCTGCAACACAGCAATAAAAATGC
86c5; GCAATAATAACGTTGCCAGGAGGATCTGGAAC
87c5; CAAATATATTTTCATGCGTATTAACCAACAGT
88c5; GATGAATATACAGAGCAGGCAATGCATGACGA
89c5; TCCTCGTTAGATTAAGTTGGGTAACGCCAGGG
90c6; ACCACCGGAACACCGTAATGGGATAGGTCACG
91c6; GCAGATAGCCGCAGACATCATTGATTCAGCAT
92c6; TGGTTTGAAATGCCTCAGGAAGATCGCACTCC
93c6; ACAGAAATAAAAAAACGACGGCCAGTGCCAAG
94c6; AAAGGGATTTTCTGCGCAACTGTTGGGAAGGG
95c7; GCCACCCTCAGCCTTCCTGTAGCCAGCTTTCA
96c7; TAGCAATAGCTTAACCGTGCATCTGCCAGTTT
97c7; AGAATAAACACTTTGTTAAAATTCGCATTAAA
98c7; GAAGGGTTAGATCTGGTGCCGGAAACCAGGCA
99c7; GTGTTTTTATACATATGTACCCCGGTTGATAA
100c8; ACCACCAGAGCAAAATTTTTAGAACCCTCATA
101c8; GAATTGAGTTATTTTAACCAATAGGAACGCCA

205; ATGGTCAATAACCTGTTTAGCTTGCGGAACAAAGAAACCACC
206; CTGTAGCTCAACATGTTTTAAAATATCCAGAACAATATTACC
207; GGCTTTTGCAAAAGAAGTTTTAGACTTTACAAACAATTCGAC
208; AGATTTAGGAATACCACATTCAAATGGATTATTTACATTGGC
209; CGAGGCGCAGACGGTCAATCAGTTATCTAAAATATCTTTAGG
210; CTAAAACACTCATCTTTGACCCTGACCTGAAAGCGTAAGAAT
211; CGAATAATAATTTTTTCACGTATCACCTTGCTGAACCTCAAA
212; CGTAACGATCTAAAGTTTTGTAACATCGCCATTAAAAATACC

102c8; AGTATCATATGGACAGTCAAATCACCATCAAT
103c8; TCAATATAATCGAAGATTGTATAAGCAAATAT
104c8; CCATCACGCAATACAAAGGCTATCAGGTCATT
105c9; GTCAGACGATTGCATAAAGCTAAATCGGTTGT
106c9; CAGAGGGTAATGTAATGTGTAGGTAAAGATTC
107c9; ACGCTCAACAGTCTACTAATAGTAGTAGCATT
108c9; GAATTATCATCTGATAAATTAATGCCGGAGAG
109c9; GTAATAACATCACCATTAGATACATTTCGCAA
110c10; ATTAAAGCCAGCAAATATCGCGTTTTAATTCG
111c10; GCGCATTAGACGCAAGGCAAAGAATTAGCAAA
112c10; AACGCCAACATTTGCTCCTTTTGATAAGAGGT
113c10; ACATTATCATTTATATTTTCATTTGGGGCGCG
114c10; GCCTTGCTGGTATATGCAACTAAAGTACGGTG
115c11; CCAGTAAGCGTAAAAATCAGGTCTTTACCCTG
116c11; AAATAGCAGCCGGAAGCAAACTCCAACAGGTC
117c11; AATAAGAGAATCAATACTGCGGAATCGTCATA
118c11; AAATCCTTTGCGCTTAATTGCTGAATATAATG
119c11; GCAACAGGAAAATAAAAACCAAAATAGCGAGA
120c12; CTGGTAATAAGTAATTTCAACTTTAATCATTG
121c12; AATCCAAATAAAAATGCTTTAAACAGTTCAGA
122c12; GTCCAGACGACATCTACGTTAATAAAACGAAC
123c12; TTAGAAGTATTGCCAGAGGGGGTAATAGTAAA
124c12; TCAATCGTCTGAACTAATGCAGATACATAACG
125c13; GCCCGTATAAACGTAACAAAGCTGCTCATTCA
126c13; TAATTTGCCAGAAGAACTGGCTCATTATACCA
127c13; ACGCGCCTGTTCGGTGTACAGACCAGGCGCAT
128c13; ACTAATAGATTGGTAGAAAGATTCATCAGTTG
129c13; GTCACACGACCCTCCATGTTACTTAGCCGGAA
130c14; ATTATTCTGAAAAAGACAGCATCGGAACGAGG
131c14; ATCCTGAATCTGAGTAATCTTGACAAGAACCG
132c14; AATAATATCCCGGGTAAAATACGTAATGCCAC
133c14; AATTGAGGAAGTAAGGGAACCGAACTGACCAA
134c14; ATAGAACCCTTCCCAGCGATTATACCAAGCGC
135c15; AAGAGAAGGATGCCCACGCATAACCGATATAT
136c15; TAAATCAAGATTTGAGGACTAAAGACTTTTTC
137c15; AATAATCGGCTGTATCGGTTTATCAGCTTGCT
138c15; TCAATCAATATCGAAAGAGGCAAAAGAATACA
139c15; GACAATATTTTAAGGAACAACTAAAGGAATTG
140c16; CGGATAAGTGCCACCCTCAGAACCGCCACCCT
141c16; CGTTTTAGCGAAGCTTGATACCGATAGTTGCG
142c16; AAACCAAGTACGAGTTTCGTCACCAGTACAAA
143c16; AAATCTAAAGCTGAAAATCTCCAAAAAAAAGG
144c16; GATAGCCCTAACGTCTTTCCAGACGTTAGTAA

#### plug sequences for the seed

65c1; GGGAATTAGA
66c1; ATTCATATGG
67c1; CCTTGAAAAC
68c1; ATTAATTACA
69c1; TGACGGGGAA
70c2; TAGCAAGGCC
71c2; AGACACCACG
72c2; AGTCAATAGT
73c2; CTGAGCAAAA
74c2; AAAGCGAAAG
75c3; CACCGTAATC
76c3; AAAATACATA
77c3; CCTTTTTAAC
78c3; AGTTACAAAA
79c3; GTCACGCTGC
80c4; CAGACTGTAG
81c4; ATGATTAAGA
82c4; TAAATGCTGA
83c4; GGGAGAAACA
84c4; CCGCTACAGG
85c5; CCTTATTAGC
86c5; CCGAGGAAAC
87c5; AAAACTTTTT
88c5; TTTAACGTCA
89c5; TAACGTGCTT
90c6; AACCAGAGCC
91c6; AGAAAAGTAA
92c6; CTAAATTTAA
93c6; TGCACGTAAA
94c6; GAGGCCGATT
95c7; CCTCAGAACC
96c7; AACAATGAAA
97c7; GCGTTAAATA
98c7; CTGAATAATG
99c7; ATCCTGAGAA
100c8; CACCAGAACC
101c8; TAACCCACAA
102c8; AAGCCTGTTT
103c8; TGGCAATTCA
104c8; AAGAGTCTGT
105c9; GTTGAGGCAG
106c9; TGAACAAAGT
107c9; TATAAAGCCA
108c9; AGAAGGAGCG
109c9; TCTTTGATTA
110c10; ATAAATCCTC
111c10; AAACAGGGAA
112c10; CATATTTAAC
113c10; AGTTTGAGTA
114c10; CAAACTATCG
115c11; ATTTACCGTT
116c11; TCAAAAATGA

| | |
|---|---|
| | 117c11; TCGAGCCAGT<br>118c11; AACTCGTATT<br>119c11; GCCAGCCATT<br>120c12; CAGGAGTGTA<br>121c12; TATTTATCCC<br>122c12; AAGTAATTCT<br>123c12; ATTTGAGGAT<br>124c12; ATTTTGACGC<br>125c13; GAGTAACAGT<br>126c13; TTCCAGAGCC<br>127c13; CTAATGCAGA<br>128c13; AGCACTAACA<br>129c13; AGATTCACCA<br>130c14; TTCGGAACCT<br>131c14; CTACAATTTT<br>132c14; GAACAAGAAA<br>133c14; AGTTGAAAGG<br>134c14; GCCAACAGAG<br>135c15; GAGACTCCTC<br>136c15; TTTGAAGCCT<br>137c15; GAAACCAATC<br>138c15; TATCAAACCC<br>139c15; ACGTGGCACA<br>140c16; CAGTACCAGG<br>141c16; GAACGCGAGG<br>142c16; AACGGGTATT<br>143c16; AGCAAATGAA<br>144c16; TGCGCGAACT |

**Table S3** Oligonucleotide staple strand sequences for the gridiron seed. There are 212 strands in the overall gridiron origami seed, with the leading numbers in front of each strand denoting its identity. Strands 65–144 are necessary for making the 16 seed columns that each bind a single slat using five sockets (see Fig. S2). The non-socketed staples are full length 42-nt strands which will not allow binding of plug handles on slats, versus the socketed staples which have 10-nt removed from their 5′ end to form a socket from the exposed seed scaffold. The column location for each of these strand types is denoted with c1–c16, where a matching set of five 10-nt plug sequences must be added to each corresponding slat in the order as written. The universal core staples are not implicated in binding of slats to the seed.

| universal reference square staples | universal reference square staples |
|---|---|
| ttttttttAAATCGGCAAAATCCCTTTGATGGTGGTTCCGtttttttt | TCATCAACGCATCGTAACCGTGCAGGGCGATC |
| ttttttttGAAAGGAGCGGGCGCTAGGAAGGGAAGAAAGCtttttttt | CCGTCGGAAATGGGATAGGTCACCCAGCTG |
| ttttttttAGTGAGCTAACTCACACCTGGGGTGCCTAATGtttttttt | CTGACCTAAAATAAGGCGTTAAATATATAAAG |
| ttttttttAGTAAAAGAGTCTGTCATCAGTGAGGCCACCGtttttttt | TTTTCAAAAATCATAATTACTAGAGCAGAGGC |
| ttttttttGTCACTGCGCGCCTGTGACGATCCAGCGCAGTtttttttt | CCAATCGTCATATGCGTTATACCCATATTT |
| ttttttttAAAAGGGACATTCTGGTCACACGACCAGTAATtttttttt | TTATATAAAAAGCCAACGCATATGATATTCA |
| ttttttttTCCCACGCAACCAGCTACCGTCGGTGGTGCCAtttttttt | TTTTGTTAAGCTGATAAATTAATGGGGTGAGA |
| ttttttttAAGCATCACCTTGCTGGCAAATGAAAAATCTAtttttttt | TAAGCAATTTTTGAGAGATCTAGCCTGAGT |
| ttttttttTTGGGCGGTTGTGTACCATTTGCCGCCAGCAGtttttttt | ATCAGAAAATTGCCTGAGAGTCTGTTTTTAGA |
| ttttttttGTAACATTATCATTTTAATTTTAAAAGTTTGAtttttttt | AACACCGGTATATTTTAGTTAATGCTTAGA |
| ttttttttCGATTAAGTTGGGTAAGGGATGTGCTGCAAGGtttttttt | GTTTAGTACAAGACAAAGAACGCGGTGAATTT |
| ttttttttATCGGGAGAAACAATATAACAGTACCTTTTACtttttttt | ACCAGTATCTATATGTAAATGCTGCTACCTTT |
| ttttttttACGGCGGATTGACCGTTTCTCCGTGGGAACAAtttttttt | ACCGTTCTAAATTCGCATTAAATTTTTGGGG |
| ttttttttTTTCCCTTAGAATCCTGTCGCTATTAATTAATtttttttt | GGTAGCTAATATTTAAATTGTAAATCAAAAAT |
| ttttttttGTAAAACTAGCATGTCTCGATGAACGGTAATCtttttttt | ATCAGGTCAGCCCCAAAAACAGGAGCCAGCTT |
| ttttttttATACCGACCGTGTGATAATTTAATGGTTTGAAtttttttt | CAAGAGAAAATCATATGTACCCCTAACAAC |
| ttttttttAGAAGCCTTTATTTCAGTAATACTTTTGCGGGtttttttt | TACCGACAATAAACAACATGTTCATATTTTCA |
| ttttttttTGTCCAGACGACGACAAAAGGTAAAGTAATTCtttttttt | ATTTTCGAGCCTGTTTATCAACAACCGCACTC |
| ttttttttGATTCCCAATTCTGCGATTCCATATAACAGTTtttttttt | AACAACGAAGAAAAATAATATCATCATTCC |
| ttttttttATAGCAAGCAAATCAGATCATTACCGCGCCCAtttttttt | ACAGTAGGGCATGTAGAAATAGTAGCATTAA |
| ttttttttAGCGGATTGCATCAAATTATAGTCAGAAGCAAtttttttt | AAGGCCGGAAATCATACAGGCAAGCGCGAGCT |
| ttttttttTTTGTTTAACGTCAAAAAATAAGAAACGATTTtttttttt | AATGTGTATTAAGCAATAAAGCATAACCTG |
| ttttttttACGAGGCATAGTAAGAAACGCCAAAAGGAATTtttttttt | ACCCTCATAAATCGGTTGTACCAATTTGACCA |
| ttttttttAACAAAGTTACCAGAAAGTAAGCAGATAGCCGtttttttt | CAGAACGCGCCAGTAATAAGAGAAAGAATA |
| ttttttttATAAGGCTTGCCCTGAGCTGCTCATTCAGTGAtttttttt | TCCTGAACCCAACATGTAATTTAGAAAAGCCT |
| ttttttttGGGAGGGAAGGTAAATACATTCAACCGATTGAtttttttt | ATTTACGAGCTTAATTGAGAATCGAAATTCTT |
| ttttttttGCGCGAAACAAAGTACCCAGCGATTATACCAAtttttttt | CATCCAATAGACAGTCAAATCACCATCATCA |
| ttttttttCTTATTAGCGTTTGCCTTTCGGTCATAGCCCCtttttttt | TTAGCAAAAGGTAAAGATTCAAAACCGGAGAG |
| ttttttttTAACCGATATATTCGGAACCATCGCCCACGCAtttttttt | ATAAAGCTATATTTTAAATGCAATCAAAGGCT |
| ttttttttAGTCTCTGAATTTACCCCAGAATGGAAAGCGCtttttttt | ATGACCCTACGCAAGGATAAAAAGAGCAAA |
| ttttttttGTCTTTCCAGACGTTAGATCTAAAGTTTTGTCtttttttt | TCGTAGGAATATAGAAGGCTTATCTTATTTAT |
| ttttttttTCGAGAGGGTTGATATAGGCGGATAAGTGCCGtttttttt | ATCGAGAAGCGAGGCGTTTTAGCGTAATTTGC |
| GGCGAGAAAGGGCGCTGGCAAGTGGAGAAGTG | AAGAACGGGAGGTTTTGAAGCCTACCAACG |
| TTAGAGCTGCGTAACCACCACACCATTTTAGA | ATCAATAATGCTATTTTGCCTCCTTTTGATA |
| CTAAATCCCGCTACAGGGCGCGGCGGGAGC | GAAAAGGTTTTTTGCGGATGGCTTCTCCAACA |
| CAAGTTTTACGAGCACGTAGGTTTTTCTTTT | TTTAGCTATATAATGCTGTAGCGAGCTTCA |

| | |
|---|---|
| ACGTCAAAAGACGGGCAACAGCTGGCGGGGAG | TTAGATACATGCAACTAAAGTACGCCCGAAAG |
| AAGAGTCTGGCCCTGAGAGAGTGTGCCAGC | CTAAGAACCAAGCAAGCCGTTTTGCTAATG |
| AGATAGGGCGCTGGTTTGCCCCAGTCACTGCC | GACTTGCGGGTATTAAACCAAGTATAGATAAG |
| CACGCTGCTGACGGGGAAAGCCGGCGAACGT | AGATTAGTTCGGCTGTCTTTCCTTCCATCCTA |
| TTAATGCGGGAACCCTAAAGGGAGCCCCCGAT | AGAGGTCAGGCATCAATTCTACTAATAGCCA |
| TTGCTTTGTTGGGGTCGAGGTGCCGTAAAGCA | ATTGCTGAATATTTTCATTTGGGGGCAAAGAA |
| CACCAGTGGGGCGAAAAACCGTCTATCAAAT | TTTTAAATATTTCGCAAATGGTCACTCAGAGC |
| TCACCGCCCACTATTAAAGAACGTGGACTCCA | GAAGTTTCAACGAGTAGATTTAGAAACATT |
| GCGGTCCATTGAGTGTTGTTCCAGTTTGGAAC | CCCAATCCAATGAAAATAGCAGCCAGCCCTTT |
| AAATCCTGTTATAAATCAAAAGAATAGCCCG | CAGTTACAACATAAAAACAGGGAAAACAATGA |
| TTTTTATACATCACGCAAATTAACTTGGCAGA | CTAACGAAATTAACTGAACACCAATTGAGT |
| CAGGAACGTCTTTGATTAGTAATAACGCTCAA | CAGCTACAGGTAATTGAGCGCGTCCAATACT |
| TAAACAGTAGAAGAACTCAAACGAAAAACG | GGTCAGGAGTCATAAATATTCATTCCAGAGGG |
| CGTGCTTTAATATCCAGAAATCCCCGGGTAC | AAGCGAACTTTAAACAGTTCAGATAGCGAG |
| AGGCGGTTAATTCGTAATCATGGTTGTTCTTC | ACTTCAAATAAATCAAAAATCAGGCCCTCGTT |
| TGCATTAGTGAAATTGTTATCCCATACCGG | AGAGAATAAAATAAACAGCCATACGGTATT |
| CGCTTTCCAACATACGAGCCGGAAGCGGCCAG | GACGGGAGGCGTCTTTCCAGAGCCAACCTCCC |
| GCAATACTGTACGCCAGAATCCTTAGCGGT | AGTCAGAGATTTTATCCTGAATCTTTAAATCA |
| TGCCTGAGGAGGCCGATTAAAGGGCGCCGCGC | GCGGAATCTTAGAGAGTACCTTTAATTGACC |
| TTGCTGGTCCTCGTTAGAATCAGATACTATGG | CTCAAATGCCAGACCGGAAGCAAAAGAGCTTA |
| CGAGCTCGTGCGTATTGGGCGCCAGGGTTAA | AATGACCATATCGCGTTTTAATTCTCAACATG |
| TTTCCTGTATGAATCGGCCAACGCATTGCCCT | CCTGACTAAAGATTAAGAGGAAGGTGTCTG |
| TTCCACACAGTCGGGAAACCTGTCTGCAGCAA | TTAAGAAAGGAAACCGAGGAAACGCAAAGACA |
| GTGTAAAGTTAATTGCGTTGCGCCAGGCGA | AATAGCAAACCCAAAAGAACTGGCTCAATAGA |
| TTCACCAGCCAACAGAGATAGAACACGCTGAG | TAAGCCCATTACGCAGTATGTTGACACCAC |
| TCGTCTGAGCGTAAGAATACGTGGGGTCAGTA | AATATCAGACATACATAAAGTTGGGAAGAAA |
| CTCATGGGAATGGCTATTAGTCCCACCAGC | GGTAATAGTTAATAAAACGAACTACGATTTTA |
| TATTACCGATAGCCCTAAATAAAGGTTTCTT | AGGCTTTACAGGTAGAAAGATTTTCAACTT |
| GCGTCCGTATAAACATCCCTTACAGAGCCGGG | TACCAGACGGAATACCACATTCAACGAGTAGT |
| GGGTTTCATCGTTAACGGCATCGCACTCAA | AACGGAATTAGCTATCTTACCGATTTACAG |
| AATGCGGCGCAGCCAGCGGTGCCGCCAGCATC | GACTCCTTAATAATAAGAGCAAGAGCGCATTA |
| ACCTGAAAAATGGATTATTTACACGTTGTA | TAGAAAATAGAGATAACCCACAAGCTGAACAA |
| ATATTTTTAAATACCTACATTTTGACATCACT | AATCTACGTAAAATGTTTAGACTGGATAGCT |
| GCGAACTGCCAGCCATTGCAACAGTATCGGCC | ACATTATTTGCAAAAGAAGTTTTGGAATCCCC |
| TGCTCGTCGAGCCTCCTCACAGTTGAGGCAA | GAGATTTAGACGATAAAAACCAAAAAAACGAG |
| TTCAGCAATGCCAGCACGCGTGCCCATAGCTG | AGATACATGCAACACTATCATAATCTTTAC |
| GGTTACCTGGGCCGTTTTCACGGTGCTCACAA | AAAGGGCGATTGACGGAAATTATTGCGTTTTC |
| CTGCATCAGCACTCTGTGGTGCTGCATAAA | AAATTCATATCACCGTCACCGACTATCAAGTT |
| AGCCAGCAAACCTCAAATATCAAATTGCCCGA | GGAATAATAGAGCCAGCAAAATCGATAGCA |
| TTAACACCTGGTCAGTTGGCAAATACAAACAA | GGCAACATACCATTAGCAACTTTGAAAGAGG |
| AGAAGATTTGAGGAAGGTTATCTACATTTG | AGAACTGGACGGTGTACAGACCAGGCAGACGG |
| TCGCCATTGCACTAACAACGTCGCTGGCAGC | TAATCATACCTTCATCAAGAGTCCTGCTCC |
| TCACTGTTAGAGCACATCCTCATATGCCGTTC | AAATTGGGGGATATTCATTACCCACATCGCCT |
| TCCGCCGTTGTAGAACGTCAGCGTGATGAA | GGTGAATTATGGTTTACCAGCGCCAATAAT |
| AGCGGGGTCAGCAGCAACCGCAAGTCCCGTAA | TTGGGAATGTTTATTTTGTCACAAATGATTAA |
| TCAATATCGCCTGCAACAGTGCCCCTTCTG | GCACCATTATAAAAGAAACGCAAAAGCAAACG |
| GAAAGGAAAAAACAGAGGTGAGGCCACAGACA | ACAGATGACTCATTATACCAGTCAGGACGGT |
| CTTTAGGAAAAAATACCGAACGAATTTAATGC | GCTGGCTGTGTGAATTACCTTATGACGGAACA |
| CTCCGGCCGCCCTGCGGCTGGTAATGGGACA | CAAGAACCCTTGAGATGGTTTAATCATCAGTT |
| TGCCGGACGGCGCGGTTGCGGTATCTGGTGTG | GTAACAAACGAGAAACACCAGAACTAATGC |
| GGTCTGGTCATTGCAGGCGCTTTCAGATGCCG | ATCGGCATATCTTTTCATAATCAATAAATCCT |
| ACGGCAGCTACGGCTGGAGGTGTGTGCCCC | TGCCTTTAAGCCACCACCGGAACCCAGACGAT |
| ACGTTATTGCGGAACAAAGAAACCTCAGATGA | GCACCGTCCACCCTCAGAACCGCCGCCAGC |
| TTCGACAAATTATCATCATATTCCAAAGAAAT | CGGAAACGCACCCTCAGAGACTTTTTCATGA |
| AGGATTTGCAATTCATCAATATTATCAAAA | TCAATCATCCATTAAACGGGTAAAGGGTAGCA |
| TAGATTAGATTATACTTCTGCGCCATGTTTA | ATGTTACCGAAGGCACCAACCTCCCTCAGC |
| CGGCAAACGGAATTTGTGAGAGATGGATAACC | GATAAATTAAGAATACACTAAAACGGGAGTTA |
| GGGTAAAGAAGGGATAGCTCTCAAGCTTTC | GGAACCAGGCGTCAGACTGTAGCCATTAAA |
| AAAAAGCCAGAAACAGCGGATCAAAGTCACGA | CAGAGCCGAATCAGTAGCGACAGATGAGCCAT |
| GGAGCGGACTCGTATTAAATCCTCCCTCAA | AGAGCCACTCACCAATGAAACCATCACCAGTA |
| AGATGATGAGAAGTATTAGACTTTCAACAGTT | GGAAGTTTAAGGGAACCGAACTGACCAAGGC |
| TTGTTTGGAGCCGTCAATAGATAATAAAATAT | TGCCACTATTAGCCGGAACGAGGCGCGCATAG |
| CCAGTCCCGCGGTCCGTTTTTTCGTCTCTAA | AGAGGCAAGTGTCGAAATCCGCGAAATCTTGA |
| TCCGTGGTGTTAAACGATGCTGATACGGAACG | TTTGACCCAACGGAGATTTGTATAATCAAC |
| AGAGACGCGCACAGGCGGCCTTTAGTGGTGCT | CATTAAAGGTTCCAGTAAGCGTCAGGTTTTGC |
| TTTCTGCTATCGACATAAAAAAAAATGCCA | TGGCCTTGGATACAGGAGTGTACTGACTCCTC |
| ATATACAGACGGATTCGCCTGATTCTTGCTTC | ATTGACAGGGGTCAGTGCCTTGATTCTGAA |
| TGCGTAGATTACAAAATCGCGCAGGTACATAA | CCACCAGATAAACAGTTAAATCTCCAAAAAA |
| TTATTTGTCAATTACCTGAGCATTCATTTG | ACGGCTACAAAAGGAGCCTTTAATAAAGGAAT |
| TAATGGAACAAACATCAAGCACCGCTTCTGG | AGCGAAACTTGCTTTCGAGGTGTTCAGCGG |
| TCACCGGAACCAGGCAAAGCGCCAGGCCTCAG | AAGGCCGCTGATACCGATAGTTGCGTATGGGA |
| AGAGGTGCGCAACTGTTGGGAATCTGCCAG | CTTTTGATATATTCACAAACAAAAATCACC |
| CGTTGTAACCTCTTCGCTATTACGGTTGGTGT | GTTTTAACGGAGGTTGAGGCAGGTGCCTCCCT |
| ATACCAAGTTTTCAGGTTTAACGACCAGAA | TGCCCGTAACCACCACCAGAGCCGCCACCCTC |
| TATTCATTCACGTAAAACAGAAATTGATTATC | AAGGCTCCAGAGGCTTTGAGGACTAAAGCCG |
| TGATGAAAGGGTTAGAACCTACCAAATCCTGA | TTTATCAGGACAGCATCGGAACGAATACGTAA |
| TGCCGGAAAACAATCGGCGAAACGTACAGAA | AAACAGCTTTTTGCGGGATCGTCAAAAACGAA |
| TCAGGCTGGAGCCGCCACGGGAACAGACTTTC | ATGACAACTCGCTGAGGCTTGCAACTCATC |
| GGTGCGGGAACGACGGCCAGTGCCACGGAAAA | TCAGTACCAAGTATAGCCCGGAATAGGTGTA |
| GCGAAAGGCGCCAGGGTTTTCCCACTTAAA | AAGAGAAGCTCAGGAGGTTTAGTACCGCCACC |
| TGTAAATCTGAAAACATAGCGATATTCATCTT | ACATGAACGCCACCCTCAGAACCGCCACCC |
| ATCAATATCTGAGAAGAGTCAATAAGAAAACT | CCCCTGCCACCACCCTCATTTTCAGGGATAG |
| AATTACCCATAGGTCTGAGAGAATGCAAAT | TGCGAATAATAGGAACCCATGTACCGTAACAC |
| ACAAAATTCGGCTTAGGTTTTAAATCAGCTC | AGTGAGAGTCACCAGTACAAACTACAACGC |
| GAAGATCGACCAATAGGAACGCCACGTTAATA | TTTTGCTATTCCACAGACAGCCCTCATAGTT |
| TTTGAGGTCTGGCCTTCCTGTAAGATTGTA | TCACCGTAGATTAGGATTAGCGGTACATGG |
| AGATGGGCATTAAATGTGAGCGAGGGTTGATA | CTCAGAACAGTATTAAGAGGCTGAGGTAATAA |
| TTAAGACGATGTGAGTGAATAACGCTTTGA | TCAGAGCCTATTTCGGAACCTATTAGTAACAG |
| ATCAAAATTTTTTTAATGGAAACAAGGCGAAT | CAAGCCCAATAATTTTTTCACGTTGAAATGC |
| TTAACCTCAATTACATTTAACAATAAAGAAGA | TGAGTTTCATAGAAAGGAACAACTTGTATCGG |
| ATTTTTTACACTCCAGCCAGCTTTCCGGAAA | CTGTAGCAAACAACTTTCAACAGTAATTTCTT |
| AATTCGCGGGACGACGACAGTATCTTCGCCAT | AGCGTAACGTAAATGAATTTTCTGCCGACA |

**Table S4** Oligonucleotide staple strand sequences for the single DNA origami reference square.

| | |
|---|---|
| **p8064** | GAGTCCACGTTCTTTAATAGTGGACTCTTGTTCCAAACTGGAACAACACTCAACCCTATCTCGGGCTATTCTTTTGATTTATAAGGGATTTTGCCGATTTCGGAACCACCATCAAACAGGATTTT<br>CGCCTGCTGGGGCAAACCAGCGTGGACCGCTTGCTGCAACTCTCTCAGGGCCAGGCGGTGAAGGGCAATCAGCTGTTGCCCGTCTCACTGGTGAAAAGAAAAACCACCCTGGCGCCCAATACGCA<br>AACCGCCTCTCCCCGCGCGTTGGCCGATTCATTAATGCAGCTGGCACGACAGGTTTCCCGACTGGAAAGCGGGCAGTGAGCGCAACGCAATTAATGTGAGTTAGCTCACTCATTAGGCACCCCAG<br>GCTTTACACTTTATGCTTCCGGCTCGTATGTTGTGTGGAATTGTGAGCGGATAACAATTTCACACAGGAAACAGCTATGACCATGATTACGAATTCGAGCTCGGTACCCGGGGATCCTCAACTGT<br>GAGGAGGCTCACGGACGCGAAGAACAGGCACGCGTGCTGGCAGAAACCCCCGGTATGACCGTGAAAACGGCCCGCCGCATTCTGGCCGCAGCACCACAGAGTGCACAGGCGCGCAGTGACACTGC<br>GCTGGATCGTCTGATGCAGGGGGCACCGGCACCGCTGGCTGCAGGTAACCCGGCATCTGATGCCGTTAACGATTTGCTGAACACACCAGTGTAAGGGATGTTTATGACGAGCAAAGAAACCTTTA<br>CCCATTACCAGCCGCAGGGCAACAGTGACCCGGCTCATACCGCAACCGCGCCCGGCGGATTGAGTGCGAAAGCGCCTGCAATGACCCCGCTGATGCTGGACACCTCCAGCCGTAAGCTGGTTGCG<br>TGGGATGGCACCACCGACGGTGCTGCCGTTGGCATTCTTGCGGTTGCTGCTGACCAGACCAGCACCACGCTGACGTTCTACAAGTCCGGCACGTTCCGTTATGAGGATGTGCTCTGGCCGGAGGC<br>TGCCAGCGACGAGACGAAAAAACGGACCGCGTTTGCCGGAACGGCAATCAGCATCGTTTAACTTTACCCTTCATCACTAAAGGCCGCCTGTGCGGCTTTTTTTACGGGATTTTTTTATGTCGATG<br>TACACAACCGCCCAACTGCTGGCGGCAAATGAGCAGAAATTTAAGTTTGATCCGCTGTTTCTGCGTCTCTTTTTCCGTGAGAGCTATCCCTTCACCACGGAGAAAGTCTATCTCTCACAAATTCC<br>GGGACTGGTAAACATGGCGCTGTACGTTTCGCCGATTGTTTCCGGTGAGGTTATCCGTTCCCGTGGCGGCTCCACCTCTGAAAGCTTGGCACTGGCCGTCGTTTTACAACGTCGTGACTGGGAAA<br>ACCCTGGCGTTACCCAACTTAATCGCCTTGCAGCACATCCCCCTTTCGCCAGCTGGCGTAATAGCGAAGAGGCCCGCACCGATCGCCCTTCCCAACAGTTGCGCAGCCTGAATGGCGAATGGCGC<br>TTTGCCTGGTTTCCGGCACCAGAAGCGGTGCCGGAAAGCTGGCTGGAGTGCGATCTTCCTGAGGCCGATACTGTCGTCGTCCCCTCAAACTGGCAGATGCACGGTTACGATGCGCCCATCTACAC<br>CAACGTGACCTATCCCATTACGGTCAATCCGCCGTTTGTTCCCACGGAGAATCCGACGGGTTGTTACTCGCTCACATTTAATGTTGATGAAAGCTGGCTACAGGAAGGCCAGACGCGAATTATTT<br>TTGATGGCGTTCCTATTGGTTAAAAAATGAGCTGATTTAACAAAAATTTAATGCGAATTTTAACAAAATATTAACGTTTACAATTTAAATATTTGCTTATACAATCTTCCTGTTTTTGGGGCTTT<br>TCTGATTATCAACCGGGGTACATATGATTGACATGCTAGTTTTACGATTACCGTTCATCGATTCTCTTGTTTGCTCCAGACTCTCAGGCAATGACCTGATAGCCTTTGTAGATCTCTCAAAAATA<br>GCTACCCTCTCCGGCATTAATTTATCAGCTAGAACGGTTGAATATCATATTGATGGTGATTTGACTGTCTCCGGCCTTTCTCACCCTTTTGAATCTTTACCTACACATTACTCAGGCATTGCATT<br>TAAAATATATGAGGGTTCTAAAAATTTTTATCCTTGCGTTGAAATAAAGGCTTCTCCCGCAAAAGTATTACAGGGTCATAATGTTTTTGGTACAACCGATTTAGCTTTATGCTCTGAGGCTTTAT<br>TGCTTAATTTTGCTAATTCTTTGCCTTGCCTGTATGATTTATTGGATGTTAATGCTACTACTATTAGTAGAATTGATGCCACCTTTTCAGCTCGCGCCCCAAATGAAAATATAGCTAAACAGGTT<br>ATTGACCATTTGCGAAATGTATCTAATGGTCAAACTAAATCTACTCGTTCGCAGAATTGGGAATCAACTGTTATATGGAATGAAACTTCCAGACACCGTACTTTAGTTGCATATTTAAAACATGT<br>TGAGCTACAGCATTATATTCAGCAATTAAGCTCTAAGCCATCCGCAAAAATGACCTCTTATCAAAAGGAGCAATTAAAGGTACTCTCTAATCCTGACCTGTTGGAGTTTGCTTCCGGTCTGGTTC<br>GCTTTGAAGCTCGAATTAAAACGCGATATTTGAAGTCTTTCGGGCTTCCTCTTAATCTTTTTGATGCAATCCGCTTTGCTTCTGACTATAATAGTCAGGGTAAAGACCTGATTTTTGATTTATGG<br>TCATTCTCGTTTTCTGAACTGTTTAAAGCATTTGAGGGGGATTCAATGAATATTTATGACGATTCCGCAGTATTGGACGCTATCCAGTCTAAACATTTTACTATTACCCCCTCTGGCAAAACTTC<br>TTTTGCAAAAGCCTCTCGCTATTTTGGTTTTTATCGTCGTCTGGTAAACGAGGGTTATGATAGTGTTGCTCTTACTATGCCTCGTAATTCCTTTTGGCGTTATGTATCTGCATTAGTTGAATGTG<br>GTATTCCTAAATCTCAACTGATGAATCTTTCTACCTGTAATAATGTTGTTCCGTTAGTTCGTTTTATTAACGTAGATTTTTCTTCCCAACGTCCTGACTGGTATAATGAGCCAGTTCTTAAAATC<br>GCATAAGGTAATTCACAATGATTAAAGTTGAAATTAAACCATCTCAAGCCCAATTTACTACTCGTTCTGGTGTTTCTCGTCAGGGCAAGCCTTATTCACTGAATGAGCAGCTTTGTTACGTTGAT<br>TTGGGTAATGAATATCCGGTTCTTGTCAAGATTACTCTTGATGAAGGTCAGCCAGCCTATGCGCCTGGTCTGTACACCGTTCATCTGTCCTCTTTCAAAGTTGGTCAGTTCGGTTCCCTTATGAT<br>TGACCGTCTGCGCCTCGTTCCGGCTAAGTAACATGGAGCAGGTCGCGGATTTCGACACAATTTATCAGGCGATGATACAAATCTCCGTTGTACTTTGTTTCGCGCTTGGTATAATCGCTGGGGGT<br>CAAAGATGAGTGTTTTAGTGTATTCTTTTGCCTCTTTCGTTTTAGGTTGGTGCCTTCGTAGTGGCATTACGTATTTTACCCGTTTAATGGAAACTTCCTCATGAAAAAGTCTTTAGTCCTCAAAG<br>CCTCTGTAGCCGTTGCTACCCTCGTTCCGATGCTGTCTTTCGCTGCTGAGGGTGACGATCCCGCAAAAGCGGCCTTTAACTCCCTGCAAGCCTCAGCGACCGAATATATCGGTTATGCGTGGGCG<br>ATGGTTGTTGTCATTGTCGGCGCAACTATCGGTATCAAGCTGTTTAAGAAATTCACCTCGAAAGCAAGCTGATAAACCGATACAATTAAAGGCTCCTTTTGGAGCCTTTTTTTTGGAGATTTTCA<br>ACGTGAAAAAATTATTATTCGCAATTCCTTTAGTTGTTCCTTTCTATTCTCACTCCGCTGAAACTGTTGAAAGTTGTTTAGCAAAATCCCATACAGAAAATTCATTTACTAACGTCTGGAAAGAC<br>GACAAAACTTTAGATCGTTACGCTAACTATGAGGGCTGTCTGTGGAATGCTACAGGCGTTGTAGTTTGTACTGGTGACGAAACTCAGTGTTACGGTACATGGGTTCCTATTGGGCTTGCTATCCC<br>TGAAAATGAGGGTGGTGGCTCTGAGGGTGGCGGTTCTGAGGGTGGCGGTTCTGAGGGTGGCGGTACTAAACCTCCTGAGTACGGTGATACACCTATTCCGGGCTATACTTATATCAACCCTCTCG<br>ACGGCACTTATCCGCCTGGTACTGAGCAAAACCCCGCTAATCCTAATCCTTCTCTTGAGGAGTCTCAGCCTCTTAATACTTTCATGTTTCAGAATAATAGGTTCCGAAATAGGCAGGGGGCATTA<br>ACTGTTTATACGGGCACTGTTACTCAAGGCACTGACCCCGTTAAAACTTATTACCAGTACACTCCTGTATCATCAAAAGCCATGTATGACGCTTACTGGAACGGTAAATTCAGAGACTGCGCTTT<br>CCATTCTGGCTTTAATGAGGATTTATTTGTTTGTGAATATCAAGGCCAATCGTCTGACCTGCCTCAACCTCCTGTCAATGCTGGCGGCGGCTCTGGTGGTGGTTCTGGTGGCGGCTCTGAGGGTG<br>GTGGCTCTGAGGGTGGCGGTTCTGAGGGTGGCGGCTCTGAGGGAGGCGGTTCCGGTGGTGGCTCTGGTTCCGGTGATTTTGATTATGAAAAGATGGCAAACGCTAATAAGGGGGCTATGACCGAA<br>AATGCCGATGAAAACGCGCTACAGTCTGACGCTAAAGGCAAACTTGATTCTGTCGCTACTGATTACGGTGCTGCTATCGATGGTTTCATTGGTGACGTTTCCGGCCTTGCTAATGGTAATGGTGC<br>TACTGGTGATTTTGCTGGCTCTAATTCCCAAATGGCTCAAGTCGGTGACGGTGATAATTCACCTTTAATGAATAATTTCCGTCAATATTTACCTTCCCTCCCTCAATCGGTTGAATGTCGCCCTT<br>TTGTCTTTGGCGCTGGTAAACCATATGAATTTTCTATTGATTGTGACAAAATAAACTTATTCCGTGGTGTCTTTGCGTTTCTTTTATATGTTGCCACCTTTATGTATGTATTTTCTACGTTTGCT<br>AACATACTGCGTAATAAGGAGTCTTAATCATGCCAGTTCTTTTGGGTATTCCGTTATTATTGCGTTTCCTCGGTTTCCTTCTGGTAACTTTGTTCGGCTATCTGCTTACTTTTCTTAAAAAGGGC<br>TTCGGTAAGATAGCTATTGCTATTTCATTGTTTCTTGCTCTTATTATTGGGCTTAACTCAATTCTTGTGGGTTATCTCTCTGATATTAGCGCTCAATTACCCTCTGACTTTGTTCAGGGTGTTCA<br>GTTAATTCTCCCGTCTAATGCGCTTCCCTGTTTTTATGTTATTCTCTCTGTAAAGGCTGCTATTTTCATTTTTGACGTTAAACAAAAAATCGTTTCTTATTTGGATTGGGATAAATAATATGGCT<br>GTTTATTTTGTAACTGGCAAATTAGGCTCTGGAAAGACGCTCGTTAGCGTTGGTAAGATTCAGGATAAAATTGTAGCTGGGTGCAAAATAGCAACTAATCTTGATTTAAGGCTTCAAAACCTCCC<br>GCAAGTCGGGAGGTTCGCTAAAACGCCTCGCGTTCTTAGAATACCGGATAAGCCTTCTATATCTGATTTGCTTGCTATTGGGCGCGGTAATGATTCCTACGATGAAAATAAAAACGGCTTGCTTG<br>TTCTCGATGAGTGCGGTACTTGGTTTAATACCCGTTCTTGGAATGATAAGGAAAGACAGCCGATTATTGATTGGTTTCTACATGCTCGTAAATTAGGATGGGATATTATTTTTCTTGTTCAGGAC<br>TTATCTATTGTTGATAAACAGGCGCGTTCTGCATTAGCTGAACATGTTGTTTATTGTCGTCGTCTGGACAGAATTACTTTACCTTTTGTCGGTACTTTATATTCTCTTATTACTGGCTCGAAAAT<br>GCCTCTGCCTAAATTACATGTTGGCGTTGTTAAATATGGCGATTCTCAATTAAGCCCTACTGTTGAGCGTTGGCTTTATACTGGTAAGAATTTGTATAACGCATATGATACTAAACAGGCTTTTT<br>CTAGTAATTATGATTCCGGTGTTTATTCTTATTTAACGCCTTATTTATCACACGGTCGGTATTTCAAACCATTAAATTTAGGTCAGAAGATGAAATTAACTAAAATATATTTGAAAAAGTTTTCT<br>CGCGTTCTTTGTCTTGCGATTGGATTTGCATCAGCATTTACATATAGTTATATAACCCAACCTAAGCCGGAGGTTAAAAAGGTAGTCTCTCAGACCTATGATTTTGATAAATTCACTATTGACTC<br>TTCTCAGCGTCTTAATCTAAGCTATCGCTATGTTTTCAAGGATTCTAAGGGAAAATTAATTAATAGCGACGATTTACAGAAGCAAGGTTATTCACTCACATATATTGATTTATGTACTGTTTCCA<br>TTAAAAAAGGTAATTCAAATGAAATTGTTAAATGTAATTAATTTTGTTTTCTTGATGTTTGTTTCATCATCTTCTTTTGCTCAGGTAATTGAAATGAATAATTCGCCTCTGCGCGATTTTGTAAC<br>TTGGTATTCAAAGCAATCAGGCGAATCCGTTATTGTTTCTCCCGATGTAAAAGGTACTGTTACTGTATATTCATCTGACGTTAAACCTGAAAATCTACGCAATTTCTTTATTTCTGTTTTACGTG<br>CAAATAATTTTGATATGGTAGGTTCTAACCCTTCCATTATTCAGAAGTATAATCCAAACAATCAGGATTATATTGATGAATTGCCATCATCTGATAATCAGGAATATGATGATAATTCCGCTCCT<br>TCTGGTGGTTTCTTTGTTCCGCAAAATGATAATGTTACTCAAACTTTTAAAATTAATAACGTTCGGGCAAAGGATTTAATACGAGTTGTCGAATTGTTTGTAAAGTCTAATACTTCTAAATCCTC<br>AAATGTATTATCTATTGACGGCTCTAATCTATTAGTTGTTAGTGCTCCTAAAGATATTTTAGATAACCTTCCTCAATTCCTTTCAACTGTTGATTTGCCAACTGACCAGATATTGATTGAGGGTT<br>TGATATTTGAGGTTCAGCAAGGTGATGCTTTAGATTTTTCATTTGCTGCTGGCTCTCAGCGTGGCACTGTTGCAGGCGGTGTTAATACTGACCGCCTCACCTCTGTTTTATCTTCTGCTGGTGGT<br>TCGTTCGGTATTTTTAATGGCGATGTTTTAGGGCTATCAGTTCGCGCATTAAAGACTAATAGCCATTCAAAAATATTGTCTGTGCCACGTATTCTTACGCTTTCAGGTCAGAAGGGTTCTATCTC<br>TGTTGGCCAGAATGTCCCTTTTATTACTGGTCGTGTGACTGGTGAATCTGCCAATGTAAATAATCCATTTCAGACGATTGAGCGTCAAAATGTAGGTATTTCCATGAGCGTTTTTCCTGTTGCAA<br>TGGCTGGCGGTAATATTGTTCTGGATATTACCAGCAAGGCCGATAGTTTGAGTTCTTCTACTCAGGCAAGTGATGTTATTACTAATCAAAGAAGTATTGCTACAACGGTTAATTTGCGTGATGGA<br>CAGACTCTTTTACTCGGTGGCCTCACTGATTATAAAAACACTTCTCAGGATTCTGGCGTACCGTTCCTGTCTAAAATCCCTTTAATCGGCCTCCTGTTTAGCTCCCGCTCTGATTCTAACGAGGA<br>AAGCACGTTATACGTGCTCGTCAAAGCAACCATAGTACGCGCCCTGTAGCGGCGCATTAAGCGCGGCGGGTGTGGTGGTTACGCGCAGCGTGACCGCTACACTTGCCAGCGCCCTAGCGCCCGCT<br>CCTTTCGCTTTCTTCCCTTCCTTTCTCGCCACGTTCGCCGGCTTTCCCCGTCAAGCTCTAAATCGGGGGCTCCCTTTAGGGTTCCGATTTAGTGCTTTACGGCACCTCGACCCCAAAAAACTTGA<br>TTTGGGTGATGGTTCACGTAGTGGGCCATCGCCCTGATAGACGGTTTTTCGCCCTTTGACGTTG |
| **p8634** | GAGTCCACGTTCTTTAATAGTGGACTCTTGTTCCAAACTGGAACAACACTCAACCCTATCTCGGGCTATTCTTTTGATTTATAAGGGATTTTGCCGATTTCGGAACCACCATCAAACAGGATTTT<br>CGCCTGCTGGGGCAAACCAGCGTGGACCGCTTGCTGCAACTCTCTCAGGGCCAGGCGGTGAAGGGCAATCAGCTGTTGCCCGTCTCACTGGTGAAAAGAAAAACCACCCTGGCGCCCAATACGCA<br>AACCGCCTCTCCCCGCGCGTTGGCCGATTCATTAATGCAGCTGGCACGACAGGTTTCCCGACTGGAAAGCGGGCAGTGAGCGCAACGCAATTAATGTGAGTTAGCTCACTCATTAGGCACCCCAG<br>GCTTTACACTTTATGCTTCCGGCTCGTATGTTGTGTGGAATTGTGAGCGGATAACAATTTCACACAGGAAACAGCTATGACCATGATTACGAATTCGAGCTCGGTACCCGGGGATCCATTCTCCT<br>GTGACTCGGAAGTGCATTTATCATCTCCATAAAACAAAACCCGCCGTAGCGAGTTCAGATAAAATAAATCCCCGCGAGTGCGAGGATTGTTATGTAATATTGGGTTTAATCATCTATATGTTTTG<br>TACAGAGAGGGCAAGTATCGTTTCCACCGTACTCGTGATAATAATTTTGCACGGTATCAGTCATTTCTCGCACATTGCAGAATGGGGATTTGTCTTCATTAGACTTATAAACCTTCATGGAATAT<br>TTGTATGCCGACTCTATATCTATACCTTCATCTACATAAACACCTTCGTGATGTCTGCATGGAGACAAGACACCGGATCTGCACAACATTGATAACGCCCAATCTTTTTGCTCAGACTCTAACTC<br>ATTGATACTCATTTATAAACTCCTTGCAATGTATGTCGTTTCAGCTAAACGGTATCAGCAATGTTTATGTAAAGAAACAGTAAGATAATACTCAACCCGATGTTTGAGTACGGTCATCATCTGAC<br>ACTACAGACTCTGGCATCGCTGTGAAGACGACGCGAAATTCAGCATTTTCACAAGCGTTATCTTTTACAAAACCGATCTCACTCTCCTTTGATGCGAATGCCAGCGTCAGACATCATATGCAGAT<br>ACTCACCTGCATCCTGAACCCATTGACCTCCAACCCCGTAATAGCGATGCGTAATGATGTCGATAGTTACTAACGGGTCTTGTTCGATTAACTGCCGCAGAAACTCTTCCAGGTCACCAGTGCAG<br>TGCTTGATAACAGGAGTCTTCCCAGGATGGCGAACAACAAGAAACTGGTTTCCGTCTTCACGGACTTCGTTGCTTTCCAGTTTAGCAATACGCTTACTCCCATCCGAGATAACACCTTCGTAATA<br>CTCACGCTGCTCGTTGAGTTTTGATTTTGCTGTTTCAAGCTCAACACGCAGTTTCCCTACTGTTAGCGCAATATCCTCGTTCTCCTGGTCGCGGCGTTTGATGTATTGCTGGTTTCTTTCCCGTT<br>CATCCAGCAGTTCCAGCACAATCGATGGTGTTACCAATTCATGGAAAAGGTCTGCGTCAAATCCCCAGTCGTCATGCATTGCCTGCTCTGCCGCTTCACGCAGTGCCTGAGAGTTAATTTCGCTC<br>ACTTCGAACCTCTCTGTTTACTGATAAGTTCCAGATCCTCCTGGCAACTTGCACAAGTCCGACAACCCTGAACGACCAGGCGTCTTCGTTCATCTATCGGATCGCCACACTCACAACAATGAGTG<br>GCAGATATAGCCTGGTGGTTCAGGCGGCGCATTTTTATTGCTGTGTTGCGCTGTAATTCTTCTATTTCTGATGCTGAATCAATGATGTCTGCCATCTTTCATTAATCCCTGAACTGTTGGTTAAT<br>ACGCATGAGGGTGAATGCGAATAATAAAGCTTGGCACTGGCCGTCGTTTTACAACGTCGTGACTGGGAAAACCCTGGCGTTACCCAACTTAATCGCCTTGCAGCACATCCCCCTTTCGCCAGCTG<br>GCGTAATAGCGAAGAGGCCCGCACCGATCGCCCTTCCCAACAGTTGCGCAGCCTGAATGGCGAATGGCGCTTTGCCTGGTTTCCGGCACCAGAAGCGGTGCCGGAAAGCTGGCTGGAGTGCGATC<br>TTCCTGAGGCCGATACTGTCGTCGTCCCCTCAAACTGGCAGATGCACGGTTACGATGCGCCCATCTACACCAACGTGACCTATCCCATTACGGTCAATCCGCCGTTTGTTCCCACGGAGAATCCG<br>ACGGGTTGTTACTCGCTCACATTTAATGTTGATGAAAGCTGGCTACAGGAAGGCCAGACGCGAATTATTTTTGATGGCGTTCCTATTGGTTAAAAAATGAGCTGATTTAACAAAAATTTAATGCG<br>AATTTTAACAAAATATTAACGTTTACAATTTAAATATTTGCTTATACAATCTTCCTGTTTTTGGGGCTTTTCTGATTATCAACCGGGGTACATATGATTGACATGCTAGTTTTACGATTACCGTT<br>CATCGATTCTCTTGTTTGCTCCAGACTCTCAGGCAATGACCTGATAGCCTTTGTAGATCTCTCAAAAATAGCTACCCTCTCCGGCATTAATTTATCAGCTAGAACGGTTGAATATCATATTGATG<br>GTGATTTGACTGTCTCCGGCCTTTCTCACCCTTTTGAATCTTTACCTACACATTACTCAGGCATTGCATTTAAAATATATGAGGGTTCTAAAAATTTTTATCCTTGCGTTGAAATAAAGGCTTCT<br>CCCGCAAAAGTATTACAGGGTCATAATGTTTTTGGTACAACCGATTTAGCTTTATGCTCTGAGGCTTTATTGCTTAATTTTGCTAATTCTTTGCCTTGCCTGTATGATTTATTGGATGTTAATGC<br>TACTACTATTAGTAGAATTGATGCCACCTTTTCAGCTCGCGCCCCAAATGAAAATATAGCTAAACAGGTTATTGACCATTTGCGAAATGTATCTAATGGTCAAACTAAATCTACTCGTTCGCAGA<br>ATTGGGAATCAACTGTTATATGGAATGAAACTTCCAGACACCGTACTTTAGTTGCATATTTAAAACATGTTGAGCTACAGCATTATATTCAGCAATTAAGCTCTAAGCCATCCGCAAAAATGACC<br>TCTTATCAAAAGGAGCAATTAAAGGTACTCTCTAATCCTGACCTGTTGGAGTTTGCTTCCGGTCTGGTTCGCTTTGAAGCTCGAATTAAAACGCGATATTTGAAGTCTTTCGGGCTTCCTCTTAA<br>TCTTTTTGATGCAATCCGCTTTGCTTCTGACTATAATAGTCAGGGTAAAGACCTGATTTTTGATTTATGGTCATTCTCGTTTTCTGAACTGTTTAAAGCATTTGAGGGGGATTCAATGAATATTT<br>ATGACGATTCCGCAGTATTGGACGCTATCCAGTCTAAACATTTTACTATTACCCCCTCTGGCAAAACTTCTTTTGCAAAAGCCTCTCGCTATTTTGGTTTTTATCGTCGTCTGGTAAACGAGGGT<br>TATGATAGTGTTGCTCTTACTATGCCTCGTAATTCCTTTTGGCGTTATGTATCTGCATTAGTTGAATGTGGTATTCCTAAATCTCAACTGATGAATCTTTCTACCTGTAATAATGTTGTTCCGTT<br>AGTTCGTTTTATTAACGTAGATTTTTCTTCCCAACGTCCTGACTGGTATAATGAGCCAGTTCTTAAAATCGCATAAGGTAATTCACAATGATTAAAGTTGAAATTAAACCATCTCAAGCCCAATT<br>TACTACTCGTTCTGGTGTTTCTCGTCAGGGCAAGCCTTATTCACTGAATGAGCAGCTTTGTTACGTTGATTTGGGTAATGAATATCCGGTTCTTGTCAAGATTACTCTTGATGAAGGTCAGCCAG<br>CCTATGCGCCTGGTCTGTACACCGTTCATCTGTCCTCTTTCAAAGTTGGTCAGTTCGGTTCCCTTATGATTGACCGTCTGCGCCTCGTTCCGGCTAAGTAACATGGAGCAGGTCGCGGATTTCGA<br>CACAATTTATCAGGCGATGATACAAATCTCCGTTGTACTTTGTTTCGCGCTTGGTATAATCGCTGGGGGTCAAAGATGAGTGTTTTAGTGTATTCTTTTGCCTCTTTCGTTTTAGGTTGGTGCCT<br>TCGTAGTGGCATTACGTATTTTACCCGTTTAATGGAAACTTCCTCATGAAAAAGTCTTTAGTCCTCAAAGCCTCTGTAGCCGTTGCTACCCTCGTTCCGATGCTGTCTTTCGCTGCTGAGGGTGA<br>CGATCCCGCAAAAGCGGCCTTTAACTCCCTGCAAGCCTCAGCGACCGAATATATCGGTTATGCGTGGGCGATGGTTGTTGTCATTGTCGGCGCAACTATCGGTATCAAGCTGTTTAAGAAATTCA |

| | |
|---|---|
| | CCTCGAAAGCAAGCTGATAAACCGATACAATTAAAGGCTCCTTTTGGAGCCTTTTTTTTGGAGATTTTCAACGTGAAAAAATTATTATTCGCAATTCCTTTAGTTGTTCCTTTCTATTCTCACTC<br>CGCTGAAACTGTTGAAAGTTGTTTAGCAAAATCCCATACAGAAAATTCATTTACTAACGTCTGGAAAGACGACAAAACTTTAGATCGTTACGCTAACTATGAGGGCTGTCTGTGGAATGCTACAG<br>GCGTTGTAGTTTGTACTGGTGACGAAACTCAGTGTTACGGTACATGGGTTCCTATTGGGCTTGCTATCCCTGAAAATGAGGGTGGTGGCTCTGAGGGTGGCGGTTCTGAGGGTGGCGGTTCTGAG<br>GGTGGCGGTACTAAACCTCCTGAGTACGGTGATACACCTATTCCGGGCTATACTTATATCAACCCTCTCGACGGCACTTATCCGCCTGGTACTGAGCAAAACCCCGCTAATCCTAATCCTTCTCT<br>TGAGGAGTCTCAGCCTCTTAATACTTTCATGTTTCAGAATAATAGGTTCCGAAATAGGCAGGGGGCATTAACTGTTTATACGGGCACTGTTACTCAAGGCACTGACCCCGTTAAAACTTATTACC<br>AGTACACTCCTGTATCATCAAAAGCCATGTATGACGCTTACTGGAACGGTAAATTCAGAGACTGCGCTTTCCATTCTGGCTTTAATGAGGATTTATTTGTTTGTGAATATCAAGGCCAATCGTCT<br>GACCTGCCTCAACCTCCTGTCAATGCTGGCGGCGGCTCTGGTGGTGGTTCTGGTGGCGGCTCTGAGGGTGGTGGCTCTGAGGGTGGCGGTTCTGAGGGTGGCGGCTCTGAGGGAGGCGGTTCCGG<br>TGGTGGCTCTGGTTCCGGTGATTTTGATTATGAAAAGATGGCAAACGCTAATAAGGGGGCTATGACCGAAAATGCCGATGAAAACGCGCTACAGTCTGACGCTAAAGGCAAACTTGATTCTGTCG<br>CTACTGATTACGGTGCTGCTATCGATGGTTTCATTGGTGACGTTTCCGGCCTTGCTAATGGTAATGGTGCTACTGGTGATTTTGCTGGCTCTAATTCCCAAATGGCTCAAGTCGGTGACGGTGAT<br>AATTCACCTTTAATGAATAATTTCCGTCAATATTTACCTTCCCTCCCTCAATCGGTTGAATGTCGCCCTTTTGTCTTTGGCGCTGGTAAACCATATGAATTTTCTATTGATTGTGACAAAATAAA<br>CTTATTCCGTGGTGTCTTTGCGTTTCTTTTATATGTTGCCACCTTTATGTATGTATTTTCTACGTTTGCTAACATACTGCGTAATAAGGAGTCTTAATCATGCCAGTTCTTTTGGGTATTCCGTT<br>ATTATTGCGTTTCCTCGGTTTCCTTCTGGTAACTTTGTTCGGCTATCTGCTTACTTTTCTTAAAAAGGGCTTCGGTAAGATAGCTATTGCTATTTCATTGTTTCTTGCTCTTATTATTGGGCTTA<br>ACTCAATTCTTGTGGGTTATCTCTCTGATATTAGCGCTCAATTACCCTCTGACTTTGTTCAGGGTGTTCAGTTAATTCTCCCGTCTAATGCGCTTCCCTGTTTTTATGTTATTCTCTCTGTAAAG<br>GCTGCTATTTTCATTTTTGACGTTAAACAAAAAATCGTTTCTTATTTGGATTGGGATAAATAATATGGCTGTTTATTTTGTAACTGGCAAATTAGGCTCTGGAAAGACGCTCGTTAGCGTTGGTA<br>AGATTCAGGATAAAATTGTAGCTGGGTGCAAAATAGCAACTAATCTTGATTTAAGGCTTCAAAACCTCCCGCAAGTCGGGAGGTTCGCTAAAACGCCTCGCGTTCTTAGAATACCGGATAAGCCT<br>TCTATATCTGATTTGCTTGCTATTGGGCGCGGTAATGATTCCTACGATGAAAATAAAAACGGCTTGCTTGTTCTCGATGAGTGCGGTACTTGGTTTAATACCCGTTCTTGGAATGATAAGGAAAG<br>ACAGCCGATTATTGATTGGTTTCTACATGCTCGTAAATTAGGATGGGATATTATTTTTCTTGTTCAGGACTTATCTATTGTTGATAAACAGGCGCGTTCTGCATTAGCTGAACATGTTGTTTATT<br>GTCGTCGTCTGGACAGAATTACTTTACCTTTTGTCGGTACTTTATATTCTCTTATTACTGGCTCGAAAATGCCTCTGCCTAAATTACATGTTGGCGTTGTTAAATATGGCGATTCTCAATTAAGC<br>CCTACTGTTGAGCGTTGGCTTTATACTGGTAAGAATTTGTATAACGCATATGATACTAAACAGGCTTTTTCTAGTAATTATGATTCCGGTGTTTATTCTTATTTAACGCCTTATTTATCACACGG<br>TCGGTATTTCAAACCATTAAATTTAGGTCAGAAGATGAAATTAACTAAAATATATTTGAAAAAGTTTTCTCGCGTTCTTTGTCTTGCGATTGGATTTGCATCAGCATTTACATATAGTTATATAA<br>CCCAACCTAAGCCGGAGGTTAAAAAGGTAGTCTCTCAGACCTATGATTTTGATAAATTCACTATTGACTCTTCTCAGCGTCTTAATCTAAGCTATCGCTATGTTTTCAAGGATTCTAAGGGAAAA<br>TTAATTAATAGCGACGATTTACAGAAGCAAGGTTATTCACTCACATATATTGATTTATGTACTGTTTCCATTAAAAAAGGTAATTCAAATGAAATTGTTAAATGTAATTAATTTTGTTTTCTTGA<br>TGTTTGTTTCATCATCTTCTTTTGCTCAGGTAATTGAAATGAATAATTCGCCTCTGCGCGATTTTGTAACTTGGTATTCAAAGCAATCAGGCGAATCCGTTATTGTTTCTCCCGATGTAAAAGGT<br>ACTGTTACTGTATATTCATCTGACGTTAAACCTGAAAATCTACGCAATTTCTTTATTTCTGTTTTACGTGCAAATAATTTTGATATGGTAGGTTCTAACCCTTCCATTATTCAGAAGTATAATCC<br>AAACAATCAGGATTATATTGATGAATTGCCATCATCTGATAATCAGGAATATGATGATAATTCCGCTCCTTCTGGTGGTTTCTTTGTTCCGCAAAATGATAATGTTACTCAAACTTTTAAAATTA<br>ATAACGTTCGGGCAAAGGATTTAATACGAGTTGTCGAATTGTTTGTAAAGTCTAATACTTCTAAATCCTCAAATGTATTATCTATTGACGGCTCTAATCTATTAGTTGTTAGTGCTCCTAAAGAT<br>ATTTTAGATAACCTTCCTCAATTCCTTTCAACTGTTGATTTGCCAACTGACCAGATATTGATTGAGGGTTTGATATTTGAGGTTCAGCAAGGTGATGCTTTAGATTTTTCATTTGCTGCTGGCTC<br>TCAGCGTGGCACTGTTGCAGGCGGTGTTAATACTGACCGCCTCACCTCTGTTTTATCTTCTGCTGGTGGTTCGTTCGGTATTTTTAATGGCGATGTTTTAGGGCTATCAGTTCGCGCATTAAAGA<br>CTAATAGCCATTCAAAAATATTGTCTGTGCCACGTATTCTTACGCTTTCAGGTCAGAAGGGTTCTATCTCTGTTGGCCAGAATGTCCCTTTTATTACTGGTCGTGTGACTGGTGAATCTGCCAAT<br>GTAAATAATCCATTTCAGACGATTGAGCGTCAAAATGTAGGTATTTCCATGAGCGTTTTTCCTGTTGCAATGGCTGGCGGTAATATTGTTCTGGATATTACCAGCAAGGCCGATAGTTTGAGTTC<br>TTCTACTCAGGCAAGTGATGTTATTACTAATCAAAGAAGTATTGCTACAACGGTTAATTTGCGTGATGGACAGACTCTTTTACTCGGTGGCCTCACTGATTATAAAAACACTTCTCAGGATTCTG<br>GCGTACCGTTCCTGTCTAAAATCCCTTTAATCGGCCTCCTGTTTAGCTCCCGCTCTGATTCTAACGAGGAAAGCACGTTATACGTGCTCGTCAAAGCAACCATAGTACGCGCCCTGTAGCGGCGC<br>ATTAAGCGCGGCGGGTGTGGTGGTTACGCGCAGCGTGACCGCTACACTTGCCAGCGCCCTAGCGCCCGCTCCTTTCGCTTTCTTCCCTTCCTTTCTCGCCACGTTCGCCGGCTTTCCCCGTCAAG<br>CTCTAAATCGGGGGCTCCCTTTAGGGTTCCGATTTAGTGCTTTACGGCACCTCGACCCCAAAAAACTTGATTTGGGTGATGGTTCACGTAGTGGGCCATCGCCCTGATAGACGGTTTTTCGCCCT<br>TTGACGTTG |

**Table S5** Sequences of the p8064 and p8634 scaffolds used to fold the 6HB and 12HB slats, single DNA origami reference square, and gridiron seed.

| 7-nt x32 | 6-nt, x256 | 7-nt, x256 | 8-nt, x256 |
|---|---|---|---|
| (GATTGTC, -7.365) | (CGAAGT, -7.209) | (GTTATCG, -7.365) | (AGCAGACG, -11.032) |
| (GTAAGAC, -7.035) | (CCAACC, -7.639) | (GATTGTC, -7.365) | (AAATGAGG, -8.622) |
| (TTACACC, -7.695) | (CCGAAA, -7.329) | (GTTCGAA, -8.155) | (AAATCTGG, -8.622) |
| (GATAACC, -7.035) | (CTACAT, -5.649) | (GTGTGTG, -8.665) | (AAACCGGG, -11.002) |
| (GTAGAAG, -6.875) | (GAAATG, -5.699) | (ACCCCAT, -9.185) | (AAGTGCAT, -9.562) |
| (GTTACAG, -7.185) | (CCAAGT, -7.029) | (GAAGATC, -7.055) | (AGGTTGTT, -9.272) |
| (CTTATGC, -7.425) | (CAAAGT, -6.189) | (GTTGGAT, -7.855) | (AGGGCTAG, -10.212) |
| (CTGAACT, -7.695) | (GGCGAT, -8.449) | (GAAACTC, -7.315) | (AGCGTAGG, -10.702) |
| (CGGAAAA, -8.255) | (CTTCCT, -6.719) | (CTGTTCT, -7.695) | (AACGGGAC, -10.902) |
| (CCTTACT, -7.365) | (CGAATT, -6.369) | (GTCGAAA, -8.155) | (AACTCCGC, -11.142) |
| (CATTTGC, -8.015) | (CTAGGT, -6.439) | (CAAGATG, -7.355) | (ACCTATCC, -9.032) |
| (GTTGCAA, -8.525) | (AGATAG, -5.339) | (TTAGTGC, -7.935) | (ATTGACGC, -10.352) |
| (GTTTGTC, -7.625) | (CATTAC, -5.419) | (GTTCGTT, -8.295) | (AGCATTTC, -9.022) |
| (GGAATTC, -7.315) | (ACCTCT, -7.109) | (CCTCAAA, -7.815) | (AAGAGACT, -8.702) |
| (GTGAATC, -7.365) | (TTCTGC, -7.289) | (GTTGCTT, -8.355) | (AAGGATCT, -8.702) |
| (TTTGACC, -7.975) | (AGAAAG, -5.879) | (CTTGCCT, -9.035) | (ATTGACCC, -9.622) |
| (GTTTCAC, -7.625) | (AGGAAC, -6.879) | (CTGAACT, -7.695) | (AAGGAATC, -8.472) |
| (CAAACTC, -7.465) | (AGTGGG, -7.869) | (GTAAGAG, -6.875) | (AACCAGCG, -11.292) |
| (CTAGATC, -6.615) | (ACCGAC, -8.209) | (GCGAAAC, -9.145) | (AGGTCAAA, -9.132) |
| (TTTAGCC, -7.885) | (GAAGAA, -5.899) | (GTCATCT, -7.595) | (AGATGTCT, -8.752) |
| (CTTGTGT, -8.005) | (TTCGGG, -8.169) | (CATCAAG, -7.355) | (AAAGCAGC, -10.362) |
| (CTTTTCC, -7.415) | (GATCAC, -6.439) | (ACGATCC, -8.875) | (AAATACCG, -8.782) |
| (GTATGAG, -6.925) | (CGGAAG, -7.659) | (CAACGAT, -8.185) | (AGGCAAGG, -10.802) |
| (GATGTCG, -8.535) | (CTTCTT, -5.879) | (CCTTTGG, -8.405) | (AGAGGTAT, -8.422) |
| (GTGATAG, -6.925) | (CGTATC, -6.439) | (CGTGAAA, -8.305) | (AGCACAAT, -9.562) |
| (CTTTGAG, -7.305) | (GAGAAT, -5.779) | (CAGAAAG, -7.305) | (AAGTCGGC, -11.142) |
| (CGCAAAA, -8.805) | (ATGCTC, -7.169) | (CTTTGCT, -8.195) | (GACGTAAA, -8.802) |
| (CTTCTGT, -7.695) | (ATGGTC, -6.929) | (GTAGATG, -6.925) | (AGCTTCAC, -9.862) |
| (GTATCAG, -6.925) | (CACAAG, -6.689) | (CCTCCAT, -8.535) | (AAACCCCT, -10.062) |
| (AAGAGAG, -7.385) | (GGGAAG, -7.329) | (CCCCAAT, -8.795) | (AGATCGAC, -9.542) |
| (GATTTCC, -7.315) | (CCAATT, -6.189) | (GTTAACC, -7.295) | (AAGTGCCG, -11.292) |
| (CCTAATC, -6.875) | (ATGGGT, -7.419) | (CTTCAAG, -7.305) | (ACGATTCC, -9.802) |
| | (AGGTAC, -6.599) | (CGAGAAA, -7.995) | (AAAATCCC, -8.732) |

| | | | |
|---|---|---|---|
| | (CAAAAG, -5.799)<br>(AAGGAT, -6.269)<br>(GACAAC, -6.699)<br>(CTTACT, -5.599)<br>(CGTGAG, -7.709)<br>(AGGACG, -8.049)<br>(GCTCAG, -7.619)<br>(ACTGAA, -6.439)<br>(TTTGCT, -6.939)<br>(GGCGTT, -8.709)<br>(GGTTTT, -6.299)<br>(CTCCAG, -7.219)<br>(GTAGGT, -6.599)<br>(GGTCAG, -7.379)<br>(TTAGCT, -6.349)<br>(CGTGAC, -7.869)<br>(AGATAC, -5.499)<br>(GCTAAT, -5.999)<br>(CGTAAT, -6.089)<br>(CAAAAC, -5.959)<br>(ATCGTC, -7.109)<br>(GTTAAG, -5.369)<br>(ACGAGC, -8.449)<br>(CCTCAG, -7.219)<br>(CCTCAT, -6.769)<br>(TTGTGC, -7.599)<br>(ATACCC, -6.599)<br>(AAGGAG, -6.719)<br>(GTACTT, -5.759)<br>(GAGACG, -7.559)<br>(AACTCT, -6.269)<br>(CTAGAT, -5.339)<br>(GAATCT, -5.779)<br>(AACTTG, -6.189)<br>(GCAAAC, -7.199)<br>(GTGATT, -6.089)<br>(AGGTCG, -8.049)<br>(TTCAGC, -7.289)<br>(GTTCAA, -6.209)<br>(CTATCC, -5.949)<br>(GATCTT, -5.779)<br>(AAGGAC, -6.879)<br>(AAGCAC, -7.429)<br>(GTTCAT, -6.089)<br>(CAACAT, -6.239)<br>(GCAAAA, -6.709)<br>(GTAGAC, -6.109)<br>(GTGCAG, -7.929)<br>(GTACAC, -6.419)<br>(CATAAC, -5.419)<br>(TTGTCG, -7.379)<br>(TTGGGG, -7.989)<br>(AGAGCT, -7.349)<br>(ATCGGG, -8.049)<br>(ATCGAC, -7.109)<br>(AAGGGT, -7.369)<br>(GTAACT, -5.759)<br>(GAGTTT, -6.039)<br>(AAGCAA, -6.939)<br>(GTACAA, -5.929)<br>(CTTGAC, -6.539)<br>(GCTAAC, -6.609)<br>(CTCGTG, -7.709)<br>(CAAACT, -6.189)<br>(AGTCAT, -6.319)<br>(TTTGGC, -7.549)<br>(GGATAA, -5.619)<br>(AACTGT, -6.579)<br>(CTATAG, -4.669)<br>(GAGAAC, -6.389) | (CGATCAA, -8.045)<br>(GTTGGAA, -7.975)<br>(CATCGAA, -8.045)<br>(CGTTTTC, -7.905)<br>(GTCAATC, -7.365)<br>(CCTAGAC, -7.715)<br>(CCTTAAG, -6.975)<br>(GATGTCG, -8.535)<br>(GCGTAAT, -8.255)<br>(ATTGCCT, -8.585)<br>(ATATCGC, -7.995)<br>(CTTTCGT, -8.135)<br>(CGCAAAA, -8.805)<br>(GTAACAG, -7.185)<br>(CACAATG, -7.665)<br>(ATGACTC, -7.595)<br>(GTGTAGC, -8.425)<br>(GTCATAG, -6.925)<br>(GTTATGC, -7.585)<br>(GTTCTTG, -7.465)<br>(ATCTCCG, -8.715)<br>(GCAACCT, -9.195)<br>(CTGTACT, -7.415)<br>(TTACTGC, -7.935)<br>(ACTTCCG, -8.975)<br>(CCTAATC, -6.875)<br>(GAAATCC, -7.315)<br>(CCACAAA, -8.125)<br>(CATTGAG, -7.355)<br>(CTCGCAA, -9.385)<br>(ATTTCGC, -8.535)<br>(GTTCGAT, -8.035)<br>(GGTGAAC, -8.465)<br>(CGCTTAA, -8.215)<br>(CCAAAAC, -7.725)<br>(CCTATGG, -7.865)<br>(TTCTTCC, -7.665)<br>(GCTGTAA, -7.935)<br>(GTTTCTG, -7.465)<br>(GAAGCAT, -8.095)<br>(GTTACAC, -7.345)<br>(CTGTAAG, -7.025)<br>(GTTGCAA, -8.525)<br>(CCTAAAC, -7.135)<br>(CTAATCG, -7.205)<br>(CTTAGAG, -6.715)<br>(GTACCAT, -7.575)<br>(GCTATTG, -7.425)<br>(CCATTAC, -7.185)<br>(CTATCCT, -7.105)<br>(CTCCCAA, -8.655)<br>(CAACCTT, -7.955)<br>(GAGTATC, -6.775)<br>(CTAACTC, -6.875)<br>(GTGAATC, -7.365)<br>(GCTGAAA, -8.215)<br>(GTTGTTG, -7.775)<br>(GCCTAAT, -7.765)<br>(CACCAAA, -8.125)<br>(TTGAGCG, -9.385)<br>(TTCGCAA, -9.055)<br>(CAGTCAA, -7.865)<br>(GTAGGAA, -7.385)<br>(CATATGC, -7.475)<br>(CTACATG, -7.075)<br>(GGTATTC, -7.035)<br>(GTGAGTT, -7.855)<br>(CCTTATG, -7.025)<br>(TTGGTCG, -9.145)<br>(ATTGCTC, -8.095) | (AACCTGGG, -10.562)<br>(AGAGTCCT, -9.542)<br>(AAACTTCG, -9.062)<br>(AACGCAAT, -10.002)<br>(AGGGTGTG, -10.612)<br>(AGATCGAT, -8.932)<br>(AAGAGTGC, -9.862)<br>(AAGAGCAC, -9.862)<br>(AGCTTCGC, -11.382)<br>(AAAGCTGT, -9.512)<br>(AAACTGGG, -9.722)<br>(AAGGGTCG, -10.742)<br>(AAACACCT, -9.272)<br>(ACTTACGC, -10.022)<br>(AAGGAAAG, -8.572)<br>(AGGTACGG, -10.462)<br>(GATAAAGC, -8.202)<br>(GGTATAGT, -7.912)<br>(AGCATCCT, -10.092)<br>(AAACCCAT, -9.272)<br>(GCATATAC, -7.972)<br>(AGCTCATT, -9.252)<br>(GACGTTAA, -8.802)<br>(AGGCTGAG, -10.542)<br>(AACGGTAT, -9.172)<br>(AGGCAGAG, -10.542)<br>(AAGGGAAT, -8.962)<br>(ATATGGCC, -9.582)<br>(AAGTCACT, -9.012)<br>(AAACTCGG, -9.902)<br>(AGTTCCGG, -10.742)<br>(AGGTGAAA, -9.132)<br>(AACGTCGG, -11.232)<br>(AAGGTAGC, -9.532)<br>(AAACCGAG, -9.902)<br>(AGGGGTAC, -10.132)<br>(AACAGGCG, -11.292)<br>(AAGTGACC, -9.622)<br>(AAATGGTC, -8.782)<br>(ACATCTCC, -9.362)<br>(ACGGTACT, -10.012)<br>(AAAGTCCC, -9.572)<br>(AGATGAGT, -8.752)<br>(AGACACAT, -9.062)<br>(AGGCCAAG, -10.802)<br>(ACGATCAA, -9.362)<br>(AAGGGGAG, -10.252)<br>(AGAGACAT, -8.752)<br>(AAAGCCAT, -9.512)<br>(AACTGCAC, -10.172)<br>(AGCAACAT, -9.562)<br>(GCGTATAA, -8.762)<br>(AGGGCTAC, -10.372)<br>(AGACCTAT, -8.422)<br>(AGGATCCT, -9.542)<br>(GACCTAAA, -8.312)<br>(AAGCACGT, -10.842)<br>(AACCCGAC, -10.902)<br>(AGCCTATT, -8.922)<br>(AAGCTCCT, -10.042)<br>(ACCTAAGC, -9.532)<br>(AAATCCAC, -8.782)<br>(AGACGAAC, -9.802)<br>(AGCCTTAT, -8.922)<br>(AAACAGCT, -9.512)<br>(AGACTGCT, -10.092)<br>(AGGTTGAT, -9.012)<br>(AAAGCGAG, -10.142)<br>(AGCTCGAG, -10.722)<br>(ATATGCGC, -10.312) |

| | | | |
|---|---|---|---|
| | (GCCAAG, -7.879)<br>(CTACGG, -7.379)<br>(TTGCCT, -7.779)<br>(ATGACC, -6.929)<br>(CTCAAT, -5.929)<br>(ATGTCC, -6.929)<br>(GTCAAT, -6.089)<br>(ATTTGG, -6.189)<br>(AGAGAC, -6.619)<br>(ATTGGC, -7.429)<br>(CAATGT, -6.239)<br>(AATCAG, -5.929)<br>(CAATAG, -5.259)<br>(AGACAC, -6.929)<br>(ATCAGC, -7.169)<br>(GTGACC, -7.539)<br>(CAATTG, -5.849)<br>(AGGTGG, -7.869)<br>(ACCGAG, -8.049)<br>(AAAGCT, -6.769)<br>(GCCTAG, -7.289)<br>(CTTCGT, -7.209)<br>(CGATCG, -7.889)<br>(GGATCG, -7.559)<br>(GATCAT, -5.829)<br>(CTTGAA, -6.049)<br>(CCTAAT, -5.599)<br>(GATAAG, -5.109)<br>(GGTAAA, -5.879)<br>(GCACAG, -7.929)<br>(GTATTG, -5.419)<br>(ATGTGC, -7.479)<br>(GTGAGC, -7.779)<br>(AGGGAG, -7.559)<br>(CATTAG, -5.259)<br>(CTGAAT, -5.929)<br>(GAATGT, -6.089)<br>(CGGAAT, -7.209)<br>(ATTTCG, -6.369)<br>(CGAAGG, -7.659)<br>(AAACAG, -6.189)<br>(ATGCCG, -8.599)<br>(CTCAAC, -6.539)<br>(GAAGCG, -8.059)<br>(ACGGAC, -8.209)<br>(GTAGAA, -5.619)<br>(AGGAGC, -7.959)<br>(AAGCAG, -7.269)<br>(TTGGAG, -6.889)<br>(TTCGCG, -8.899)<br>(CTTCAC, -6.539)<br>(GGAAAA, -6.159)<br>(CACAAC, -6.849)<br>(CTAGAG, -5.789)<br>(CTTGCT, -7.269)<br>(CTGTTC, -6.539)<br>(GTCGAC, -7.719)<br>(GATCAA, -5.949)<br>(GAGATT, -5.779)<br>(ACACAT, -6.629)<br>(CATCTT, -5.929)<br>(AGCAAC, -7.429)<br>(ACGGAG, -8.049)<br>(CCTATC, -5.949)<br>(GTGAGG, -7.379)<br>(ATGGCT, -7.659)<br>(CTGAAG, -6.379)<br>(CTACAA, -5.769)<br>(GAATAG, -5.109)<br>(GGAAAT, -6.039) | (GTTGACG, -8.795)<br>(CTGAGTT, -7.695)<br>(AAGACAG, -7.695)<br>(TTTACCC, -7.645)<br>(GTAGAAC, -7.035)<br>(CATGGAA, -7.865)<br>(GTAGGTT, -7.525)<br>(GTCATTC, -7.365)<br>(TTAGACC, -7.385)<br>(CAAAACC, -7.725)<br>(CAGAGAA, -7.555)<br>(ACGATCG, -9.205)<br>(CTGATCT, -7.435)<br>(GATTTCC, -7.315)<br>(GCTATAC, -6.995)<br>(TTGTGCG, -9.695)<br>(GTCTGAA, -7.715)<br>(CCTTTTC, -7.415)<br>(CTCACCT, -8.535)<br>(CCAGTAA, -7.535)<br>(TTGTGTC, -8.025)<br>(CCATTAG, -7.025)<br>(GATAACC, -7.035)<br>(CCTAGAA, -7.225)<br>(GCCTATT, -7.765)<br>(CTCAAGG, -8.145)<br>(GCTTTTG, -7.965)<br>(CTGTAGG, -7.865)<br>(GTGAGAT, -7.595)<br>(TTCTGCG, -9.385)<br>(CTATTGC, -7.425)<br>(CTGGAAT, -7.695)<br>(CCCATAA, -7.535)<br>(CGTCAAA, -8.305)<br>(TTGTCTC, -7.715)<br>(TTCAGTC, -7.715)<br>(GTCTCTT, -7.545)<br>(CTTAACC, -7.135)<br>(CATTTGC, -8.015)<br>(CGTCTAA, -7.715)<br>(GTAATCC, -7.035)<br>(CTAACTG, -7.025)<br>(CTAAAGC, -7.375)<br>(CTCATGG, -8.195)<br>(CCTTAAC, -7.135)<br>(CAAGATC, -7.205)<br>(ACTGTCG, -9.025)<br>(GTCTAAG, -6.875)<br>(GCTAAAC, -7.535)<br>(GAATGCG, -9.035)<br>(CCTGAAT, -7.695)<br>(CTGATGC, -8.595)<br>(GTTCATC, -7.365)<br>(TTGTACC, -7.695)<br>(GTATCTC, -6.775)<br>(CAAACAG, -7.615)<br>(CTCCAAT, -7.695)<br>(CCTAGAT, -7.105)<br>(CATAGAG, -6.765)<br>(GTTGTGT, -8.165)<br>(GAACATC, -7.365)<br>(GAAAGTC, -7.315)<br>(GTAACTC, -7.035)<br>(GCTAAAG, -7.375)<br>(CTACATC, -6.925)<br>(TTTAGCC, -7.885)<br>(CTTGTGT, -8.005)<br>(GTTCTCT, -7.545)<br>(GTCTGAT, -7.595)<br>(CGTTAAG, -7.465) | (ACTTACCC, -9.292)<br>(ACCATTGT, -9.322)<br>(AGAGGAAT, -8.702)<br>(ACGTTGAT, -9.502)<br>(AGCTGAAC, -9.862)<br>(AGATGTGT, -9.062)<br>(ACATAGCC, -9.582)<br>(AGCTCTAT, -8.662)<br>(AGTGAAGC, -9.862)<br>(AGCCTAGG, -10.212)<br>(AGGGTGAC, -10.462)<br>(AGTGACAT, -9.062)<br>(AGTTGGAT, -9.012)<br>(AAACGAGG, -9.902)<br>(AGATTGCT, -9.252)<br>(ACATACGC, -10.072)<br>(ATTGAGCC, -9.862)<br>(AGCATGGC, -11.252)<br>(AGGTACAA, -8.852)<br>(AGCAGAAA, -9.372)<br>(GACTTTAC, -7.962)<br>(AGATCGCG, -11.212)<br>(AGCCTAAT, -8.922)<br>(AGCTGGAG, -10.542)<br>(AAATAGCG, -9.022)<br>(AAATAGGC, -8.692)<br>(AACAGGTT, -9.272)<br>(ACATAGGC, -9.582)<br>(ACGTGAAT, -9.502)<br>(AGGCGTAG, -10.702)<br>(AGAGCAAC, -9.862)<br>(ACCTACAA, -8.852)<br>(AAGGGTGG, -10.562)<br>(AGGATTTC, -8.472)<br>(AGCTTCAT, -9.252)<br>(AAGTACGC, -10.022)<br>(AAATTGCC, -9.282)<br>(AGGCCTAG, -10.212)<br>(AACTCCCG, -10.742)<br>(AGGTACGC, -10.862)<br>(AACCCACG, -11.052)<br>(AACACCGC, -11.452)<br>(AGGTCGAG, -10.482)<br>(AGCACAGC, -11.252)<br>(ACGATTTC, -8.962)<br>(AAGGATAC, -8.192)<br>(AGGTAGGC, -10.372)<br>(ACCGTAAA, -9.292)<br>(ATTTCCCC, -9.572)<br>(ACGCTAAC, -10.022)<br>(AGTTCGAT, -9.192)<br>(AGCAAGGG, -10.802)<br>(AGCAATGT, -9.562)<br>(ACATTCCC, -9.622)<br>(AAAGCTCT, -9.202)<br>(AGCATCGC, -11.432)<br>(ACCATTTC, -8.782)<br>(AGTAGCGC, -11.102)<br>(ACCTAGAT, -8.422)<br>(AGGGCTTC, -10.652)<br>(ACTAACCT, -8.682)<br>(AGCGTTAT, -9.412)<br>(AAAAGCGT, -9.952)<br>(AAGAGTGG, -9.462)<br>(AGGCAAAT, -9.512)<br>(AAAGCCAC, -10.122)<br>(AAATACCC, -8.452)<br>(AGATCAGT, -8.752)<br>(AAGGTGTT, -9.272)<br>(AGTCCTAA, -8.542) |

| | | | |
|---|---|---|---|
| | (CTACTT, -5.599)<br>(CTAAGG, -6.049)<br>(GTGACG, -7.869)<br>(TTCCAG, -6.889)<br>(CTTTAG, -5.209)<br>(CTTTGT, -6.189)<br>(GTACAT, -5.809)<br>(AACCAA, -6.699)<br>(CAATCT, -5.929)<br>(GTAGAT, -5.499)<br>(CTAAGT, -5.599)<br>(CATGAT, -5.979)<br>(GCCATG, -7.929)<br>(CCAAAA, -6.309)<br>(TTCACG, -7.379)<br>(GAATTC, -5.549)<br>(GAAAAG, -5.649)<br>(AATGAG, -5.929)<br>(AGCGAG, -8.289)<br>(GTGAAA, -6.209)<br>(CATCAA, -6.099)<br>(CATAGT, -5.649)<br>(GAACCG, -7.819)<br>(CTGAAC, -6.539)<br>(GTTCGT, -7.369)<br>(ATGCAC, -7.479)<br>(GCAATT, -6.589)<br>(CGGTAT, -6.929)<br>(GTCGAG, -7.559)<br>(GTTTAG, -5.369)<br>(AATACG, -6.089)<br>(CGTTTT, -6.629)<br>(GCTGAG, -7.619)<br>(GTTGAA, -6.209)<br>(GACTTT, -6.039)<br>(GGTTCT, -6.879)<br>(GTGAAT, -6.089)<br>(ATGCGT, -8.149)<br>(TTCTCC, -6.739)<br>(CTTCAT, -5.929)<br>(TTGTCC, -7.049)<br>(CTTGAG, -6.379)<br>(GAGAGG, -7.069)<br>(GTACCG, -7.539)<br>(GCTAAA, -6.119)<br>(CATATG, -5.309)<br>(TTGCAC, -7.599)<br>(ATGGGG, -7.869)<br>(CAGTTT, -6.189)<br>(CATCAT, -5.979)<br>(GTAGCG, -7.779)<br>(CTTTAC, -5.369)<br>(ACGTCG, -8.539)<br>(ATCGCG, -8.779)<br>(GATGAT, -5.829)<br>(CTACCG, -7.379)<br>(ACTGCT, -7.659)<br>(CTAACT, -5.599)<br>(GTGGTG, -7.689)<br>(GGAACG, -7.819)<br>(GATATC, -5.009)<br>(GTAGTC, -6.109)<br>(CATTCG, -6.869)<br>(AGAAAC, -6.039)<br>(AACTAC, -5.759)<br>(ATACGC, -7.329)<br>(GTTGAT, -6.089)<br>(CGGAAA, -7.329)<br>(CATTCT, -5.929)<br>(ATGCGG, -8.599) | (TTCTGTC, -7.715)<br>(CCATATC, -6.925)<br>(GAAGTTC, -7.315)<br>(CCTAAGG, -7.815)<br>(GTCAGAA, -7.715)<br>(CCATTCT, -7.695)<br>(CCGAAAT, -8.135)<br>(GAGATCG, -8.225)<br>(CTAATGC, -7.425)<br>(GTTTGTG, -7.775)<br>(CATAACG, -7.515)<br>(GCCAAAT, -8.355)<br>(AACAGAG, -7.695)<br>(GTTTGAG, -7.465)<br>(GCAGAAC, -8.705)<br>(TTCCAAG, -7.815)<br>(CTGTCAG, -8.195)<br>(CCTGAAA, -7.815)<br>(GCTAATG, -7.425)<br>(CAGCTAT, -7.655)<br>(AGCCCAA, -9.545)<br>(GTTACTG, -7.185)<br>(AAGTCAG, -7.695)<br>(GTGTATC, -7.085)<br>(CCAGAAA, -7.815)<br>(CTTGTCT, -7.695)<br>(TTCCTCC, -8.505)<br>(GTATACC, -6.755)<br>(GTTAACG, -7.625)<br>(CGTAGAA, -7.715)<br>(CACATAG, -7.075)<br>(AAGTGAG, -7.695)<br>(CCTTACT, -7.365)<br>(CGAAAAG, -7.745)<br>(GTTCAAC, -7.625)<br>(GGATATC, -6.775)<br>(GGAATTC, -7.315)<br>(ACCGTAG, -8.695)<br>(GTTGATG, -7.515)<br>(CATGCAA, -8.415)<br>(GTTGATC, -7.365)<br>(GTTTCAC, -7.625)<br>(TTGAACC, -7.975)<br>(CTACAAG, -7.025)<br>(CTAGATC, -6.615)<br>(CCTATTC, -6.875)<br>(GTCAGAT, -7.595)<br>(GTTAGTC, -7.035)<br>(GTATTGC, -7.585)<br>(ACCTGCT, -9.425)<br>(GTGTCAT, -7.905)<br>(TTTGTGC, -8.525)<br>(CGGTAAA, -7.975)<br>(TTAGTCC, -7.385)<br>(GCTCAAA, -8.215)<br>(GAAGTCG, -8.485)<br>(CCAAGAA, -7.815)<br>(TTCGTCC, -8.995)<br>(CTCATCT, -7.435)<br>(GTTCCAT, -7.855)<br>(GTTCCCT, -8.645)<br>(GCTATAG, -6.835)<br>(CTAAGTG, -7.025)<br>(GCCTAAA, -7.885)<br>(CAAATCG, -7.795)<br>(CTGCTAT, -7.655)<br>(CAAGTAG, -7.025)<br>(GTGACTT, -7.855)<br>(GTTTTCG, -7.905)<br>(GTGAAAG, -7.465) | (ATTTGGGC, -10.122)<br>(AACTCGAC, -9.802)<br>(AGTAGAGC, -9.272)<br>(AGGTTGAC, -9.622)<br>(AAAAGGTG, -8.882)<br>(AGGGTAGG, -9.972)<br>(ACTAGACT, -8.422)<br>(AGCACTAT, -8.972)<br>(AGTACGCG, -11.192)<br>(AGGTGGAG, -10.302)<br>(AAACACGG, -10.212)<br>(GAGCTAAT, -8.432)<br>(AAAAGACG, -9.062)<br>(AGCCTAGC, -10.612)<br>(AAACTGGC, -10.122)<br>(AGATGCAC, -9.912)<br>(AAGACCCG, -10.742)<br>(ACTCTAGC, -9.272)<br>(AGCACAAA, -9.682)<br>(AGCATGGG, -10.852)<br>(AAAAGCAC, -9.282)<br>(ATAGAGGC, -9.272)<br>(AAACAGGG, -9.722)<br>(AGGTAGTT, -8.682)<br>(ACATCTGC, -9.912)<br>(ACGGTAAT, -9.172)<br>(AAAGTCCG, -9.902)<br>(AGACCAAT, -9.012)<br>(AAGAGAGG, -9.152)<br>(AATCCGCG, -11.472)<br>(ACGTATGT, -9.222)<br>(AGTTAGCT, -8.922)<br>(GAATAACC, -7.962)<br>(AAACAGGC, -10.122)<br>(AAGCAGAC, -9.862)<br>(AGAGGAAA, -8.822)<br>(ACTGTACC, -9.342)<br>(AGCGTAGC, -11.102)<br>(AAAGGACC, -9.572)<br>(GATTAAGC, -8.202)<br>(AAGCACCT, -10.352)<br>(AAATCGAG, -8.802)<br>(AATCGCGG, -11.472)<br>(ACGGTAGT, -10.012)<br>(AACGGAAC, -10.062)<br>(GATTTACC, -7.962)<br>(ACGTACAT, -9.222)<br>(AGGTCGTT, -10.292)<br>(AGGACATT, -9.012)<br>(AGAGAAGC, -9.552)<br>(ACCTATGT, -8.732)<br>(ATTACGCC, -10.022)<br>(GAGTTAAC, -7.962)<br>(AGCTTTTC, -8.972)<br>(AAGGTACC, -9.292)<br>(GAGTTATG, -7.852)<br>(ACATCACC, -9.672)<br>(ACGCTAAT, -9.412)<br>(GATAAACG, -8.292)<br>(ACCTAACT, -8.682)<br>(AGAGTGCG, -11.032)<br>(AACGGACT, -10.292)<br>(AACGCGTT, -11.282)<br>(AAGCCAAC, -10.122)<br>(AAATGTCC, -8.782)<br>(AGGTGATT, -9.012)<br>(AACAGGAC, -9.622)<br>(AGTGTCAT, -9.062)<br>(AGTAAGCT, -8.922)<br>(AAAGGTCC, -9.572) |

| | | | |
|---|---|---|---|
| | (GTACGG, -7.539)<br>(GAACAT, -6.089)<br>(CTAGCG, -7.619)<br>(GACAAT, -6.089)<br>(GGTAAC, -6.369)<br>(CAACTT, -6.189)<br>(TTCCCG, -8.169)<br>(GATTAG, -5.109)<br>(GATTTC, -5.549)<br>(GCACTG, -7.929)<br>(GACACG, -7.869)<br>(GAAAAC, -5.809)<br>(TTGCGT, -8.269) | (GTGATTC, -7.365)<br>(CTTGCTT, -8.195)<br>(GCTAGAA, -7.625)<br>(CGTCACT, -9.025)<br>(CTAAGTC, -6.875)<br>(GTAAACC, -7.295)<br>(CGTGTAA, -8.025)<br>(AACGTAG, -7.855)<br>(CGTTTTG, -8.055)<br>(CAAGTTC, -7.465)<br>(CCTAGTT, -7.365)<br>(AGGTTCG, -8.975)<br>(CGACTAA, -7.715) | (AGGATGAT, -8.752)<br>(AACCTGCG, -11.292)<br>(AGGACAAC, -9.622)<br>(AGCCTGAG, -10.542)<br>(AGGACAAT, -9.012)<br>(ACCCTAAT, -8.682)<br>(AACCTTCT, -8.962)<br>(AGCTGAAT, -9.252)<br>(AAACCCGG, -11.002)<br>(AAATGACG, -9.112)<br>(AGGTGAAT, -9.012)<br>(AGTCTCAA, -8.872)<br>(AGACGCCT, -11.372) |

**Table S6**: The 6-, 7-, and 8-nt handle sequences and their NUPACK 3.0[52] computed kcal/mol energies. Either the handle or complementary handle sequence from the above was appended to the 3′ end of the top- or bottom-staple strand sequence with a TT linker. The majority of megastructures formed in this study used permutations of the 32 7-nt handle sequences, versus the 256 6-, 7-, and 8-nt handle sequences which were used to determine growth and spontaneous nucleation using ribbons.

| 9-nt, x128 | 10-nt, x100 |
|---|---|
| (CAGCTACGA, -11.862) | (CTAGTCGACC, -12.404) |
| (TAATAGCAC, -9.932) | (TATTCATACG, -10.688) |
| (CAAAGTTTC, -9.318) | (CGCACATAAT, -11.814) |
| (CACATCAAC, -10.258) | (CCTGATCCTC, -12.244) |
| (TAACATCCC, -10.812) | (TATGAAACAC, -10.948) |
| (CGCATAAAA, -10.118) | (CCCTAGACCC, -13.014) |
| (TTTAAAAAA, -7.328) | (TAACATGGGA, -12.462) |
| (AATTCAAAC, -8.868) | (CTTTTCTGCA, -11.828) |
| (TTGTCATTC, -9.618) | (TTGACCAATG, -11.534) |
| (ATCAAAATC, -8.608) | (CAATATCGCA, -11.778) |
| (TTTTCAACT, -9.218) | (TAAATACATT, -9.168) |
| (CAAGACCCT, -11.228) | (TAATCAACGT, -11.618) |
| (ATGGCCTTT, -11.278) | (ATTACTCCCG, -12.054) |
| (ATTGTGCCT, -11.328) | (ATACCATCGG, -12.104) |
| (TCGAATCTG, -10.802) | (GACTAGCCGT, -13.294) |
| (TAACCAAAG, -10.072) | (TACGCCTAAA, -12.488) |
| (TCATCCATG, -10.672) | (CCCGTTTGAT, -12.644) |
| (CGTTTTCAA, -10.158) | (GGCTAACAAT, -11.434) |
| (CATGAAACG, -10.538) | (CTCCGATGTT, -12.384) |
| (CCAGACTTC, -10.738) | (TCACATTTGC, -12.258) |
| (CAGCATTCT, -10.678) | (CAATTTGATC, -10.034) |
| (CGACCGTAT, -11.618) | (CAACCAGTTG, -12.124) |
| (CAGGTCTGT, -11.278) | (TTGCAATAAC, -10.764) |
| (TTTTCTCCG, -10.688) | (CAGTCTGCCC, -13.894) |
| (ACACTTCAT, -9.988) | (TTTCGTATTC, -10.394) |
| (AACATTCGT, -10.428) | (TGGACCACTT, -12.818) |
| (TACAAAAGC, -10.472) | (CATTGAGCTC, -11.954) |
| (CCTTAGCTC, -10.648) | (TTGAATTTAA, -8.884) |
| (CGGGATATC, -10.638) | (TATCCACGTT, -12.458) |
| (TGAATTTCC, -9.672) | (CTCTCCTGAC, -12.244) |
| (CTACCAGCT, -11.188) | (ACGTTAACAC, -11.684) |
| (ACTTTCACC, -10.548) | (AACTCTTGAC, -11.214) |
| (CGCTTACGT, -12.118) | (GACCCGATCA, -13.348) |
| (TCTCAGACC, -11.312) | (ACCTACAACC, -12.034) |
| (TATTCAATC, -8.872) | (TCCTTTACCC, -12.168) |
| (TGCGTAAAC, -11.222) | (CACGTATCAC, -11.924) |
| (TTTGATTTT, -8.378) | (ACAAATGCAC, -12.074) |
| (GCGTTCAAT, -11.278) | (CTGGAGGCCC, -14.684) |
| (AATCTTACA, -8.782) | (ATCTTCAAAC, -10.374) |
| (ATCCACCAG, -11.278) | (CACATCAGCT, -12.494) |
| (TAACCCTTC, -10.762) | (CACCAATTTC, -11.134) |
| (AAACGATCC, -10.728) | (ACATTCAAAC, -10.684) |
| (CCTCTCACA, -11.242) | (CAGAACTCGT, -12.384) |
| (TCATACACC, -10.502) | (ATGCTCAATT, -11.154) |
| (TTTGCACAT, -10.658) | (TAACCAACTC, -11.738) |
| (ACAACATCG, -10.928) | (CGTGCAGATA, -13.198) |
| (CGTTCTCAA, -10.738) | (TCTACAGAGC, -12.198) |
| (CCATACCAC, -10.768) | (CGGACTATCC, -12.404) |
| (TGCAAATTT, -9.872) | (GCGTTCAAAT, -12.204) |
| (TAATTCTTT, -8.472) | (CACGAGAAAC, -12.154) |
| (ACTGGACCT, -11.618) | (TAAAAGACGG, -12.018) |
| (GGTACTCAC, -10.618) | (TATTTGTGGA, -11.622) |
| (CATCCAGGT, -11.278) | (TACATTTCCC, -11.738) |
| (ACAATATTC, -8.328) | (CTGTCGTCCT, -13.224) |
| (TACATTTCG, -10.302) | (CGACTACCCT, -12.894) |
| (AGGGTCACC, -12.228) | (AACCAAAATC, -10.634) |
| (ACCACATTC, -10.598) | (AAAGGCGTTT, -12.644) |
| (TAACTTGAT, -9.362) | (CGACTATGCA, -12.618) |
| (GAGGAGGTC, -11.428) | (CAATTCCATC, -10.874) |
| (TGTTTACCG, -10.822) | (TACTTTGTCG, -12.068) |
| (AACAAACAA, -9.528) | (AATTACGTTT, -10.184) |
| (AATTACACG, -9.758) | (AAAAACAAAC, -10.054) |
| (TAGTCCAAC, -10.812) | (ATTCCCTTGG, -12.154) |
| (GAGATCTCG, -10.658) | (CCAGTACCCT, -12.714) |

| | |
|---|---|
| (AAGCAACTC, -10.788)<br>(GTTCTTGCA, -11.062)<br>(ACATTACAT, -8.868)<br>(TAAAATTTC, -8.242)<br>(CTTTACACT, -9.268)<br>(TCGAAACTT, -10.562)<br>(AACAACGTT, -10.688)<br>(TATTATTCC, -8.542)<br>(CTCTGTAGC, -10.698)<br>(TAACAAATC, -9.132)<br>(TCAACTTCT, -10.122)<br>(TTAACATGC, -9.838)<br>(CACAATTCT, -9.598)<br>(GACTAGGTG, -10.458)<br>(CCATCTTCC, -10.738)<br>(TACAAATTA, -9.045)<br>(GTAGGATCG, -10.638)<br>(CCCCGAGGA, -13.532)<br>(CCCGGAACC, -13.118)<br>(ACGTACAAC, -10.758)<br>(CCCTCTTCA, -11.192)<br>(TTCATGACA, -10.122)<br>(CAGATCCCA, -11.242)<br>(TATTCTTCC, -9.662)<br>(CTACGGACT, -11.128)<br>(AACATACTC, -9.168)<br>(CTAAGAGCC, -10.648)<br>(CGCCAATTT, -11.378)<br>(CTGTTACCC, -10.718)<br>(GCCTGTACC, -11.958)<br>(CCGTTAACT, -10.548)<br>(ATCATTCAT, -8.888)<br>(CTAAGCCCT, -11.138)<br>(CTTTTGCGT, -11.378)<br>(TACTTCCAC, -10.812)<br>(TCGATTCCA, -11.415)<br>(TAATGTTGC, -10.522)<br>(CCCCAACTT, -11.488)<br>(AACTTCACC, -10.548)<br>(ATCACTGCA, -11.342)<br>(CCATCCTCA, -11.242)<br>(AATTGCTCC, -10.788)<br>(CAGTACCCC, -11.558)<br>(ACTTCAAAC, -9.708)<br>(CTCACAACT, -10.438)<br>(TTCGACATC, -10.638)<br>(ATACCTTTT, -8.768)<br>(ACGGCTAAT, -11.178)<br>(ACTTCGTAA, -9.958)<br>(CTACCCTGG, -11.398)<br>(TAACGAATT, -9.802)<br>(TAATTCCAG, -9.812)<br>(AAGATTTTC, -8.558)<br>(TTTCCTTTC, -9.518)<br>(CTAAGGCTC, -10.648)<br>(ACTTTAACC, -9.378)<br>(ACTCCTCTG, -10.968)<br>(TCTTATGGC, -10.692)<br>(TATATTGCA, -9.595)<br>(CTGGAGTCA, -11.242)<br>(TTTTTAACA, -8.322)<br>(CACCACTTT, -10.698)<br>(AATCAACAC, -9.758)<br>(CCTTCTCAG, -10.578) | (CCGTCTTTCA, -12.608)<br>(TAATTTCAAA, -9.568)<br>(TCGTACTACC, -12.118)<br>(CAAGCCATTT, -11.864)<br>(CGGTATCTCC, -12.404)<br>(AAAACTGTCA, -11.138)<br>(ACCTTAGACG, -12.054)<br>(TCTAAGCAAC, -11.618)<br>(CTTTAACACT, -10.194)<br>(AGAAAAAAAT, -9.134)<br>(TTCCAATTCA, -10.998)<br>(ATCAAAAATT, -9.184)<br>(GGAGACTTCC, -12.354)<br>(CGTTTTTGCA, -12.578)<br>(CTGACCGGAC, -13.834)<br>(AACTTGCAAC, -12.024)<br>(TCGCGATTAA, -12.638)<br>(AACAAAATGC, -11.184)<br>(GTATCAAAAT, -9.254)<br>(CAGTAGCTCC, -12.464)<br>(CCACCATTTG, -12.124)<br>(TTTGACAAAC, -10.804)<br>(TCATTTTTCA, -10.482)<br>(CTGGATGACC, -12.554)<br>(TCCTTTTCAC, -11.658)<br>(CACTACATCC, -11.434)<br>(CATTGTCCAT, -11.414)<br>(CTTGCCAGAT, -12.444)<br>(TAACGACATC, -11.968)<br>(TTAATTCACA, -9.878)<br>(ATGCAAACAT, -11.464)<br>(GTCGGGTTCT, -13.334)<br>(TACAATTACC, -10.618)<br>(AGTTGAAGCA, -12.218)<br>(TCGATTTTCA, -11.502)<br>(AATCATTTAA, -8.764) |

**Table S7**: The 9- and 10-nt handle sequences and their NUPACK 3.0[52] computed kcal/mol energies. Either the handle or complementary handle sequence from the above was appended to the 3′ end of the top- or bottom-staple strand sequence with no linker. These handle sequences were used to initially determine the viability of such length handle sequences for controlling nucleation of megastructures, as shown in Fig. S6.

<table>
<tr><th>original nanocube strands</th><th>modified nanocube strand</th></tr>
<tr><td rowspan="3">GTAAGTTGAAGTAGGAAGCTTTTTCTAGCCATAGCATCGACACTACGACCTGCTTTTCGACAC<br>GGACTGCATTCTGGACAGTAACTGCATTAACTACGTGCTCCCAACATAAGTGACGTCCTCAGCAG<br>TTGAAAATTATCTCGATAAGCAGAAGGACCTGTATAACTGGCAAGAGACAAGGCCGCTTCAGAA<br>AGGATAGCCGGACCGTATTAATGCCGCGCCAACGGTTTCCCGGACCTAGTGTCTATCAAGTCTA<br>TTCTATGAAACCATTCTCGGGTCGAGCGGGTCACTGTTGTGACCTACGAGAAGCGTATAGATGT<br>TCCGCGCGAATAGCTCACAGGCGAACTACGTATGAATTGGTTTAAACGCTCCTCGGGAATTAAT<br>ACGACAGGTGGCAAACCACCTCCGATGTCAGCGCCGCATACCCATTCACTGTGAATTTCCACAC<br>CGAGGATTCGCAGGTCCATGGGATTCACCAAGCTCGTATACACCCTGATTCTCCATGGCAGCGC<br>TTTTAGAATGCAGTCCGTGTCGAAAAGCATAGACACTCG<br>TTTTTAGACTTGAGGTCGTATTTT<br>TTTTCAGGTCCTTACTGTCCTTTT<br>TTTTTTGGCGCGGCTTCCTATTTT<br>TTTTTTCTGAAGCACTTATGTTTT<br>TTTTCTTCAACTTACCTGCTGAGGACGTCGGCCTTGACGGTCCGGGTTT<br>TTTTTTGGGAGCACGTAGTTTAGAAAAAGCATTAATTCTCTTGCTAAAC<br>TTTTGTGTCGATGCTATGGCAATGCAGTTCTGCTTATAGGTCCGGGTCA<br>TTTTCCGAGAATGCTATCCTTTTT<br>TTTTGGAGCGTTCAGTTATATTTT<br>TTTTTTCTCGTAGGAAACCGTTTT<br>TTTTCTGTGAGCATTTTCAATTTT<br>AGATATATTCGCGCGGAACATATGGAGAAGTTTGCCACCTGTCGTTTTT<br>CAACAGTGACCATCCCATGTGAATGGGTATGCGGCTTTT<br>CAATTCATACGCGGAGGTGTCAGGGTGTATACGAGTTTT<br>TTTTGCGCTGCCCTATACGCTTTT<br>TTTTGCTGACATTAGTTCGCTTTT<br>TTTTCTTGGTGACGCTCGACTTTT<br>TTTTGTGTGGAAATTCCCGATTTT</td><td>CATAGAAATTAATTCACAGGACCTGCGAATCCTCGTTTTTttttTGGAGTCACAGATTGG</td></tr>
<tr><th>complementary handle sequence</th></tr>
<tr><td>CCAATCTGTGACTCCA</td></tr>
</table>

**Table S8** Oligos for the nanocube, as adapted from as previously published[37]. The complementary 16-nt handle sequence was appended to the 6HB bottom staple strands so that nanocubes could be bound to the megastructures to create patterns.
